# Acid sphingomyelinase deactivation post-ischemia/ reperfusion promotes cerebral angiogenesis and brain remodeling via small extracellular vesicles

**DOI:** 10.1101/2021.03.01.433387

**Authors:** A Mohamud Yusuf, N Hagemann, X Zhang, M Zafar, T Hussner, C Bromkamp, C Martiny, T Tertel, V Börger, F Schumacher, FA Solari, M Hasenberg, C Kleinschnitz, TR Doeppner, B Kleuser, A Sickmann, M Gunzer, B Giebel, R Kolesnick, E Gulbins, DM Hermann

**Affiliations:** Department of Neurology, University Hospital Essen, Essen, Germany; Center for Translational and Behavioral Neurosciences, University Hospital Essen, Essen, Germany; Institute of Transfusion Medicine, University Hospital Essen, Essen, Germany; Institute of Molecular Biology, University Hospital Essen, Essen, Germany; Department of Toxicology, University of Potsdam, Nuthetal, Germany; Institute of Pharmacy, Freie Universität Berlin, Berlin, Germany; Leibniz-Institut für Analytische Wissenschaften-ISAS-e.V., Dortmund, Germany; Institute of Immunology and Experimental Imaging, University Hospital Essen, Essen, Germany; Department of Neurology, University Medicine Göttingen, Göttingen, Germany; Medizinisches Proteom-Center (MPC), Ruhr University, Bochum, Germany; Department of Chemistry, College of Physical Sciences, University of Aberdeen, Aberdeen, Scotland, U.K.; Memorial Sloan Kettering Cancer Center, New York, New York, U.S.A.

**Keywords:** Antidepressants, ceramide, exosome, focal cerebral ischemia, middle cerebral artery occlusion, sphingomyelin, stroke recovery

## Abstract

Functional inhibitors of acid sphingomyelinase are clinically used as anti-depressants since ∼60 years. Here, we show that acid sphingomyelinase inhibition by the antidepressants amitriptyline, fluoxetine and desipramine protects from ischemia/reperfusion and elicits a profound brain remodeling response with increased angiogenesis, improved blood-brain barrier integrity, reduced brain leukocyte infiltration and increased neuronal survival. Angiogenesis is promoted by small extracellular vesicles with *bona fide* characteristics of exosomes, which are released from endothelial cells and which constitute an elegant target for the amplification of stroke recovery.

---

Ischemic stroke is the leading cause of long-term disability and a major cause of death in humans ^1^. Treatment options are limited to the acute stroke phase, in which reopening of the occluded artery by thrombolysis ^2^ and/or mechanical thrombectomy ^3^ have been shown to promote clinical recovery. Owing to reperfusion therapies, the outcome of ischemic stroke patients has considerably improved. Yet, the majority of stroke patients still exhibit long-term neurological deficits. Strong efforts are currently made to establish restorative therapies that promote brain tissue remodeling and enhance neurological recovery in the post-acute stroke phase ^4, 5^. First treatments already entered clinical trials in human stroke patients ^6–8^. A safe treatment that enhances post-ischemic tissue remodeling and brain recovery would meet a major need in patient care.

In a randomized placebo-controlled multicenter study, the serotonin reuptake inhibitor and anti-depressant fluoxetine (20 mg/day) enhanced motor recovery over a follow-up of 90 days when administered to stroke patients with moderate to severe motor deficits ^9^. Improved activities of daily living and increased long-term survival independent of the presence of depressive symptoms have been reported in another randomized placebo-controlled study, in which the tricyclic antidepressant nortriptyline (100 mg/day), which is the active metabolite of the non-selective monoamine reuptake inhibitor amitriptyline, or fluoxetine (40 mg/day) were administered over 12 weeks ^10, 11^. In a placebo-controlled functional magnetic resonance imaging (fMRI) crossover study, fluoxetine (20 mg) improved hand motor performance and enhanced primary motor cortex activation in patients with pure motor hemiparesis ^9, 10, 12^. More recently, hopes were dampened by two randomized multicenter studies that did not detect significant recovery-promoting effects of fluoxetine over 6 months ^13, 14^. A possible reason for the failure of fluoxetine was a doubling of bone fractures due to increased falls.

Antidepressants have long been believed to act via the modulation of monoaminergic neurotransmitter systems ^15^. Experimental studies more recently showed an essential role of acid sphingomyelinase (abbreviated ASM for the human protein, abbreviated Asm for the murine protein) in mediating antidepressant drug effects ^15^. ASM hydrolytically cleaves sphingomyelin to ceramide, which is a constituent of membrane microdomains that profoundly control cell signaling processes ^16, 17^. In a model of unpredictable stress-induced depression, the antidepressants amitriptyline and fluoxetine, which are diverse in their neurotransmitter mode of action, inhibited cerebral Asm activity, restored neuronal proliferation and differentiation in the hippocampus and reversed depressive-like behaviors ^18^. In mice lacking Asm, both antidepressants did not increase neuronal proliferation and differentiation and did not reverse depressive-like behaviors ^18^: Hence, Asm inhibition is indispensable for the antidepressive effects of both drugs in stress-induced depression.

Following focal and global cerebral ischemia, Asm activity and ceramide level have been shown to be increased in the ischemic mouse and rat brain ^19–22^. After transient middle cerebral artery occlusion (MCAO), genetic Asm deficiency reduced infarct volume, neurological deficits and proinflammatory cytokine abundance in the acute stroke phase, that is, 24 hours post-stroke ^19^. The role of the Asm/ ceramide system in brain tissue remodeling in the post-acute stroke phase and the underlying mechanisms of brain responses to Asm inhibitory antidepressants requires definition. In the present study, we examined effects of Asm inhibitors, many of which are clinically used as antidepressants, on cerebral angiogenesis and brain tissue remodeling after focal cerebral ischemia/ reperfusion (I/R). Following observations that chemically and pharmacologically diverse antidepressants amplified cerebral angiogenesis, we discovered a novel mechanism via which ASM/ ceramide inhibition induces the formation and release of small extracellular vesicles (sEVs) by human cerebral microvascular endothelial cells, which have *bona fide* characteristics of exosomes and which, as we further show, promote angiogenesis. Increased angiogenesis sets the stage for successful brain parenchymal remodeling.

## Results

### I/R increases brain Asm activity and ceramide level *in vivo*, which is reversed by the ASM inhibitor amitriptyline

In the reperfused ischemic striatum of vehicle-treated mice we found increased Asm activity at 24 hours, but not 14 days post-MCAO (Fig. 1A). Asm inhibition by the tricyclic antidepressant amitriptyline (2 or 12 mg/kg b.i.d.) dose-dependently reduced Asm activity at both time-points (Fig. 1A). Both long-chain (C16, C18) and very long-chain (C20, C22, C24:1) ceramides were increased predominantly in cerebral microvessels upon I/R, as shown by liquid chromatography tandem-mass spectrometry (LC-MS/MS) combined with immunohistochemistry (Fig. 1B; **Suppl. Fig. 1**). Amitriptyline reduced ceramide levels (Fig. 1B; **Suppl. Fig. 1**). This effect similarly affected long-chain and very long-chain ceramides (Fig. 1B). Total sphingomyelin content did not change in response to I/R or amitriptyline (**Suppl. Fig. 2**). An increase of fusogenic C16 long-chain sphingomyelin and C22 and C24:1 very long-chain sphingomyelin at the expense of the most abundant C18 sphingomyelin was noted following I/R (**Suppl. Fig. 2**). Amitriptyline applied immediately after I/R reduced infarct volume, but did not alter brain edema or blood-brain barrier permeability assessed by IgG extravasation after 24 hours (**Suppl. Fig. 3A-C**). Delayed amitriptyline initiated after 24 hours did not influence infarct volume after 14 days (3.38±1.22, 3.59±1.17 and 2.89±1.09 mm³) in mice receiving vehicle, amitriptyline 2 mg/kg b.i.d. and amitriptyline 12 mg/kg b.i.d., respectively. At this time-point, infarct volume was resolved.

**Figure 1.**
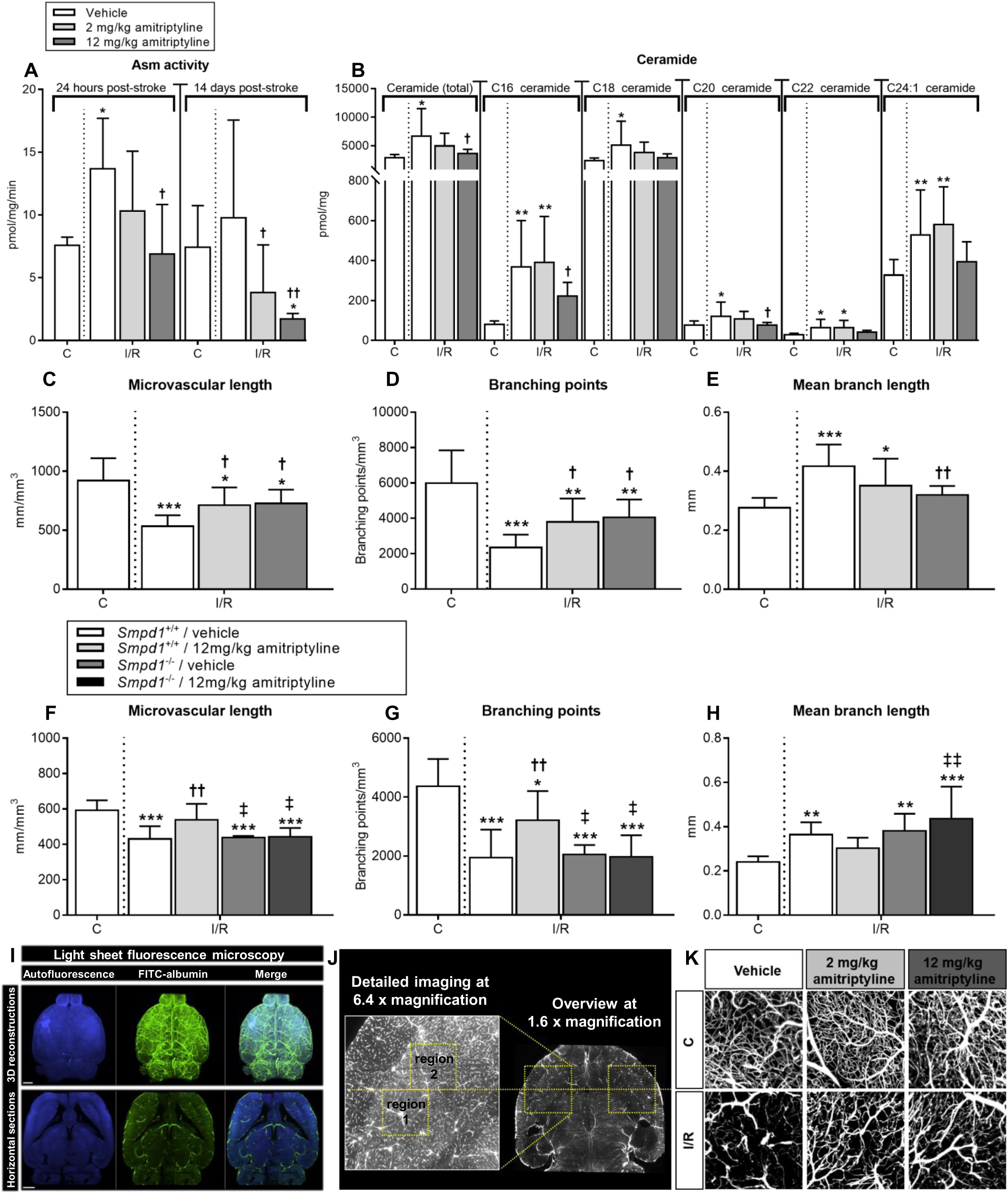
Amitriptyline induces angiogenesis after I/R *in vivo* in an Asm dependent way. **(A)** Asm activity in the reperfused ischemic striatum (labeled I/R) and contralateral non-ischemic striatum (labeled C), measured using BODIPY-labeled sphingomyelin in wildtype mice exposed to transient MCAO, which were intraperitoneally treated with vehicle or amitriptyline (2 or 12 mg/kg b.w., b.i.d.) immediately after MCAO or with 24 hours delay, followed by animal sacrifice after 24 hours or after 14 days. **(B)** Ceramide content, measured by LC-MS/MS in I/R and C of wildtype MCAO mice treated with vehicle or amitriptyline for 14 days as above. **(C)** Total microvascular length, **(D)** branching point density and **(E)** mean microvascular branch length, evaluated by LSM in I/R and C of wildtype MCAO mice treated with vehicle or amitriptyline for 14 days. **(F)** Microvascular length, **(G)** branching point density and **(H)** mean branch length in C and I/R of *Smpd1^+/+^* (wildtype) and *Smpd1^-/-^* (that is, ASM-deficient) MCAO mice treated with vehicle or amitriptyline for 14 days. Note that amitriptyline increases angiogenesis in wildtype but not *Smpd1^-/-^* mice. Representative 3D stacks post-I/R are shown in **(I)**, ROIs for the evaluation of microvascular networks in **(J)**, and maximum projection images inside these ROIs (100 µm-thick) in **(K)**. Data are means ± SD values. *p≤0.05/**p≤0.01/***p≤0.001 compared with non-ischemic C; ^†^p≤0.05/^††^p≤0.01 compared with corresponding vehicle; ^‡^p≤0.05/^‡‡^p≤0.01 compared with corresponding *Smpd1^+/+^* (n=4-7 animals/group [in **(A)**]; n=7-9 animals/group [in **(B)**]; n=7-8 animals/group [in **(C-E)**]; n=5-7 animals/ group [in **(F-H)**]; analyzed by one-way ANOVA, followed by LSD tests).

### Asm inhibition by amitriptyline induces angiogenesis after I/R *in vivo*

To evaluate consequences of Asm inhibition on neurovascular remodeling, we next examined the architecture of cerebral microvessels by 3D light sheet microscopy (LSM). LSM allows the quantitative analysis of microvascular networks labeled by the microvascular tracer FITC-albumin. Following optical clearing, image segmentation, skeletonization and 3D reconstruction, microvascular network characteristics can be quantified ^23, 24^. LSM has strongly advanced the evaluation of cerebral microvessels, which hitherto had been possible *ex vivo* only to limited extent in 2D sections. Administration of the ASM inhibitor amitriptyline for 14 days starting after 24 hours increased microvascular density post-I/R, as reflected by an increase in total microvascular length, increase in branching point number and reduction in mean branch length between branching points in the reperfused ischemic striatum (Fig. 1C-E), which is the core of the middle cerebral artery territory.

Amitriptyline is a functional ASM inhibitor, which also acts as non-selective monoamine reuptake inhibitor and besides has anticholinergic and antihistaminergic properties ^15^. To elucidate the role of Asm in amitriptyline’s actions, we next examined microvascular network characteristics in *Smpd1^-/-^* (that is, Asm deficient) mice, which were likewise treated with vehicle or amitriptyline as their wildtype counterparts. Amitriptyline increased microvascular density in the I/R striatum of *Smpd1^+/+^* but not *Smpd1^-/-^* mice, as revealed by an increase in the total microvascular length and the number of branching points (Fig. 1F, G). Mean branch length was not significantly influenced by amitriptyline in *Smpd1^+/+^* mice (Fig. 1H). These studies showed that amitriptyline induced the formation of cerebral microvessels via Asm/ ceramide pathway inhibition. Representative 3D stacks of transparent brains are shown in Figure 1I, regions of interest (ROI) used for the evaluation of microvascular network characteristics in Figure 1J and representative maximum projection images of microvessels inside these ROI of wildtype mice receiving vehicle or amitriptyline in Figure 1K.

Conventional immunohistochemistry of coronal 2D sections confirmed an increased number of CD31^+^ microvessels in the I/R striatum of wildtype mice by amitriptyline (Fig. 2A), indicative of enhanced angiogenesis. Asm inhibition reduced IgG extravasation in the brain parenchyma (Fig. 2B), reduced brain infiltrates of CD45^+^ leukocytes (Fig. 2C) and increased neuronal survival (Fig. 2D) in the I/R striatum. These data provide strong evidence that angiogenesis induced by Asm inhibition entrains a robust brain parenchymal remodeling response. Blood-brain barrier tightening, downscaling of the brain’s inflammatory response and promotion of long-term neuronal survival were hallmarks of the successful tissue recovery.

**Figure 2.**
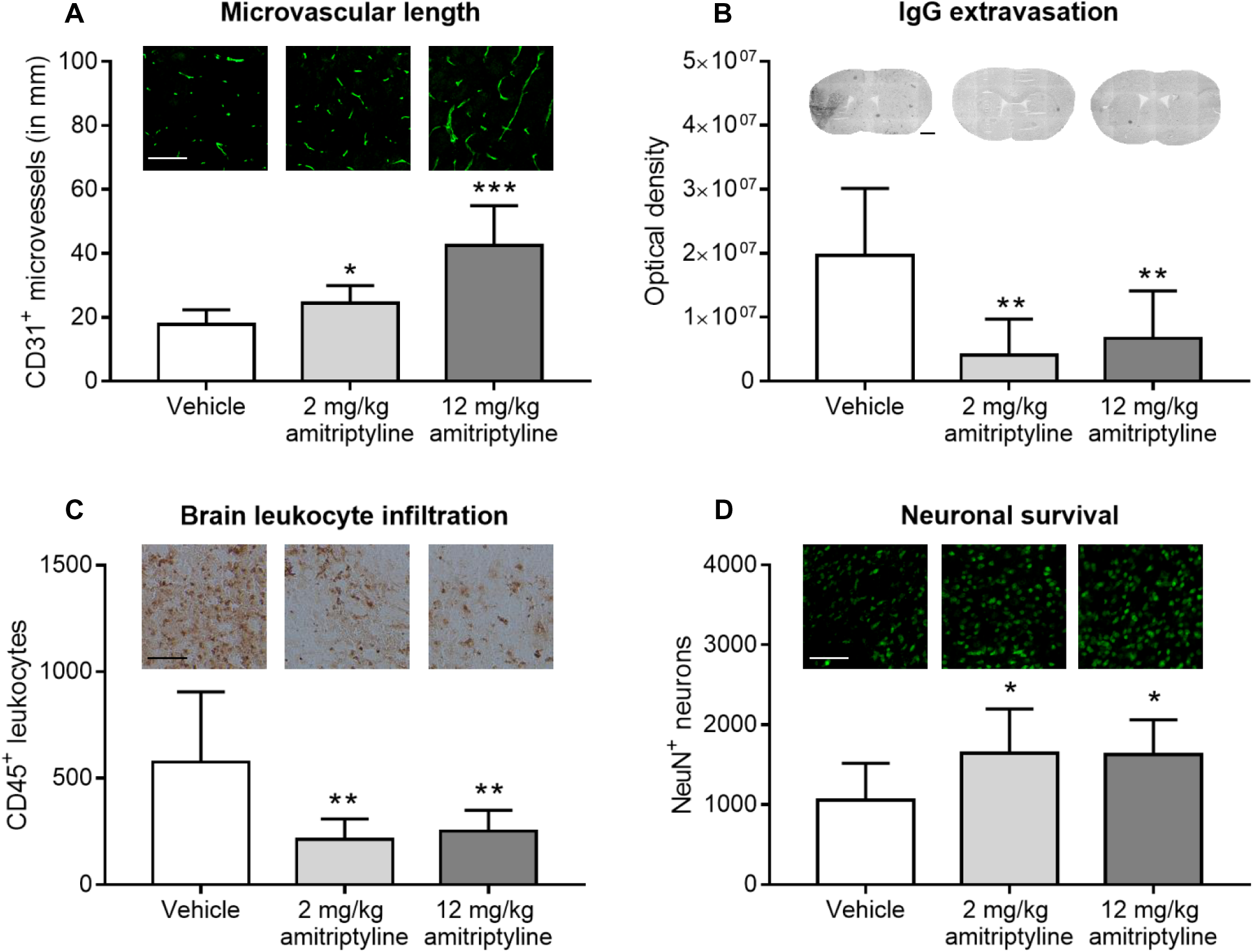
Asm inhibitor amitriptyline promotes brain remodeling after I/R *in vivo* when administered with 24 hours delay. **(A)** CD31^+^ microvessels, **(B)** serum IgG extravasation into brain parenchyma, **(C)** infiltrating CD45^+^ leukocytes and **(D)** surviving NeuN^+^ neurons evaluated by immunohistochemistry in the reperfused ischemic striatum of C57BL/6j mice exposed to transient MCAO, which were intraperitoneally treated with vehicle or amitriptyline (2 or 12 mg/kg b.w., b.i.d.) starting 24 hours after MCAO, followed by animal sacrifice after 14 days. Representative photographs are also shown. Data are means ± SD values. *p≤0.05/**p≤0.01/***p≤0.001 compared with corresponding vehicle (n=6-8 animals/group; analyzed by one-way ANOVA, followed by LSD tests). Scale bars, 100 µm (in **(A, C)**); 1000 µm (in **(B)**); 200 µm (in **(D)**).

### Amitriptyline reduces ASM activity of human cerebral microvascular endothelial cells *in vitro* and decreases the intracellular accumulation of ceramide-rich vesicles that are formed upon I/R

To further examine the effects of ASM inhibitors on cerebral microvessels, we next exposed human cerebral microvascular endothelial cells (hCMEC/D3) to non-ischemic control condition (C), oxygen-glucose deprivation (OGD) as *in vitro* model of ischemia (I), or OGD followed by reoxygenation/glucose re-supplementation as *in vitro* model of I/R. In contrast to *in vivo* conditions, ASM activity decreased after 24 hours ischemia (I) or 24 hours ischemia followed by 3 or 24 hours reoxygenation/glucose re-supplementation (I/R) (Fig. 3A, B). Upon I/R but not I only, ceramide accumulation was found in intracellular vesicles (Fig. 3C**, D**). The number of ceramide-rich vesicles reached maximum values at 3 hours post-I/R and then returned to baseline levels (**Suppl. Fig. 4A**). Ceramide-rich vesicles typically had a diameter of ∼200-250 nm (**Suppl. Fig. 4B**). Both ASM activity (Fig. 3A, B) and the number of intracellular ceramide-rich vesicles (Fig. 3C, D) were markedly reduced by amitriptyline. RT-qPCR studies revealed that *SMPD1* (i.e., *ASM*) mRNA was abundantly expressed in hCMEC/D3, whereas *SMPD2* (i.e., *NSM1*) and *SMPD3* (i.e., *NSM2*) mRNA levels were low (**Suppl. Fig. 5A**). A similar distribution, that is, abundant *SMPD1* mRNA but low *SMPD2* and *SMPD3* mRNAs, was detected in primary human brain microvascular endothelial cells (HBMECs) and human umbilical vein endothelial cells (HUVECs) (**Suppl. Fig. 5A**). In these studies, human peripheral blood mononuclear cells (hPBMC), which exhibited high *SMPD3* mRNA levels, served as positive control (**Suppl. Fig. 5A**). In hCMEC/D3, HBMECs and HUVECs, magnesium dependent NSM (that is, NSM2) activity was by a factor of 45.6±11.7, 58.9±15.3 and 91.4±49.3 lower than whole brain NSM activity (**Suppl. Fig. 5B**). These studies showed that *SMPD1* (ASM) is the predominant sphingomyelinase in endothelial cells.

**Figure 3.**
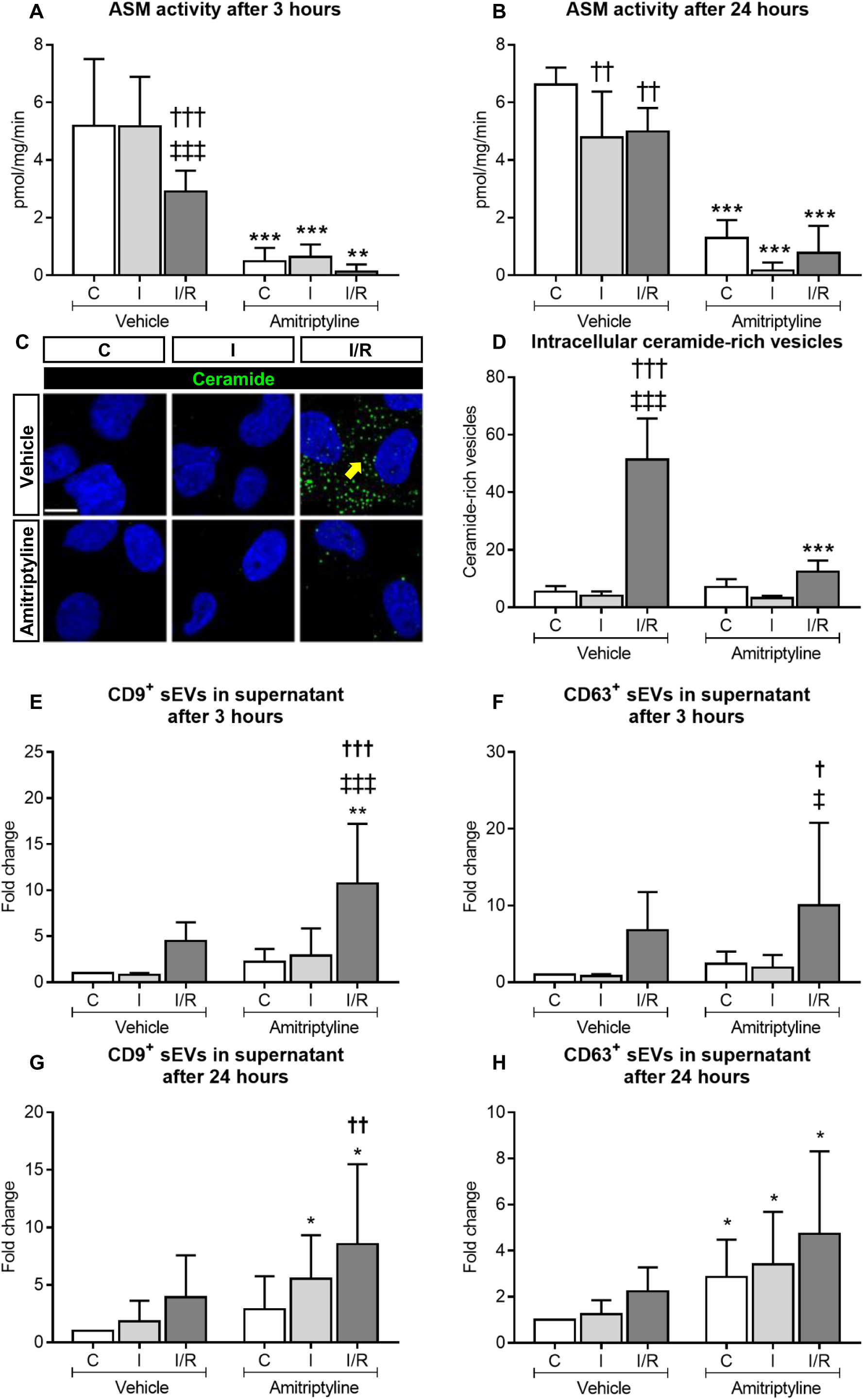
Amitriptyline reduces the intracellular accumulation of ceramide-rich vesicles after I/R *in vitro* and promotes the extracellular release of vesicles with immunofluorescence characteristics of exosomes. ASM activity, evaluated using BODIPY-labeled sphingomyelin in hCMEC/D3 exposed to **(A)** non-ischemic control condition (C), 3 hours OGD (that is, ischemia; I), or 24 hours ischemia followed by 3 hours reoxygenation/glucose re-supplementation (I/R), or to **(B)** non-ischemic C, 24 hours ischemia (I) or 24 hours ischemia followed by 24 hours reoxygenation/glucose re-supplementation (I/R), which were treated with vehicle or amitriptyline (50 µM) during 3 hours (in **(A)**) or 24 hours (in **(B)**). Note that ASM activity is reduced by I and I/R. **(C)** Immunohistochemistry for ceramide in hCMEC/D3 exposed to non-ischemic C, 24 hours I or 24 hours/3 hours I/R treated with vehicle or amitriptyline (50 µM). Note the intracellular accumulation of ceramide-rich vesicles (in green) upon I/R, which is reduced by amitriptyline (selected vesicles labeled with arrow; nuclei were counterstained in blue with DAPI). The number of vesicles evaluated by Cell Profiler is shown in **(D)**. Particle concentration of **(E,G)** CD9^+^ and **(F,H)** CD63^+^ sEVs in the supernatant of hCMEC/D3 exposed to 3 hours C, 3 hours I or 24 hours/3 hours I/R (in **(E,F)**) or 24 hours C, 24 hours I or 24 hours/24 hours I/R (in **(G,H)**) treated with vehicle or amitriptyline (50 µM). Particle concentration was evaluated by AMNIS flow cytometry. Note that the number of CD9^+^ and CD63^+^ sEVs, which is elevated upon I/R, further increases by amitriptyline. Data are means ± SD values. *p≤0.05/**p≤0.01/***p≤0.001 compared with corresponding vehicle; ^†^p≤0.05/^††^p≤0.01/^†††^p≤0.001 compared with corresponding C; ^‡^p≤0.05/^‡‡‡^p≤0.001 compared with corresponding I (n=4-5 independent samples/group [in **(A,B,D-F)**]; n=6-9 independent samples/group [in **(G,H)**]; analyzed by two-way ANOVA, followed by LSD tests).

### ASM inhibition by amitriptyline, fluoxetine and desipramine promotes angiogenesis *in vitro*

Subsequent studies showed that amitriptyline dose-dependently increased the tube formation and transwell migration of hCMEC/D3, increased VEGFR2 abundance on hCMEC/D3 and increased VEGF concentration in the supernatants of hCMEC/D3 (Fig. 4A-H). At the doses examined, amitriptyline did not influence hCMEC/D3 viability (**Suppl. Fig. 6A**). Similar findings, that is, elevation of VEGFR2 abundance on endothelial cells and VEGF concentration in supernatants, were made in the murine cerebral endothelial bEND5 cells (**Suppl. Fig. 7A, B**). Additional studies using the serotonin reuptake inhibitor fluoxetine and tricyclic desipramine, which similar to amitriptyline act as ASM inhibitors ^25^, revealed that fluoxetine and desipramine dose-dependently increased the tube formation, transwell migration and VEGFR2 abundance of hCMEC/D3 (Fig. 4I-L). Again, both inhibitors did not influence hCMEC/D3 viability at the doses administered (**Suppl. Fig. 6B, C**).

**Figure 4.**
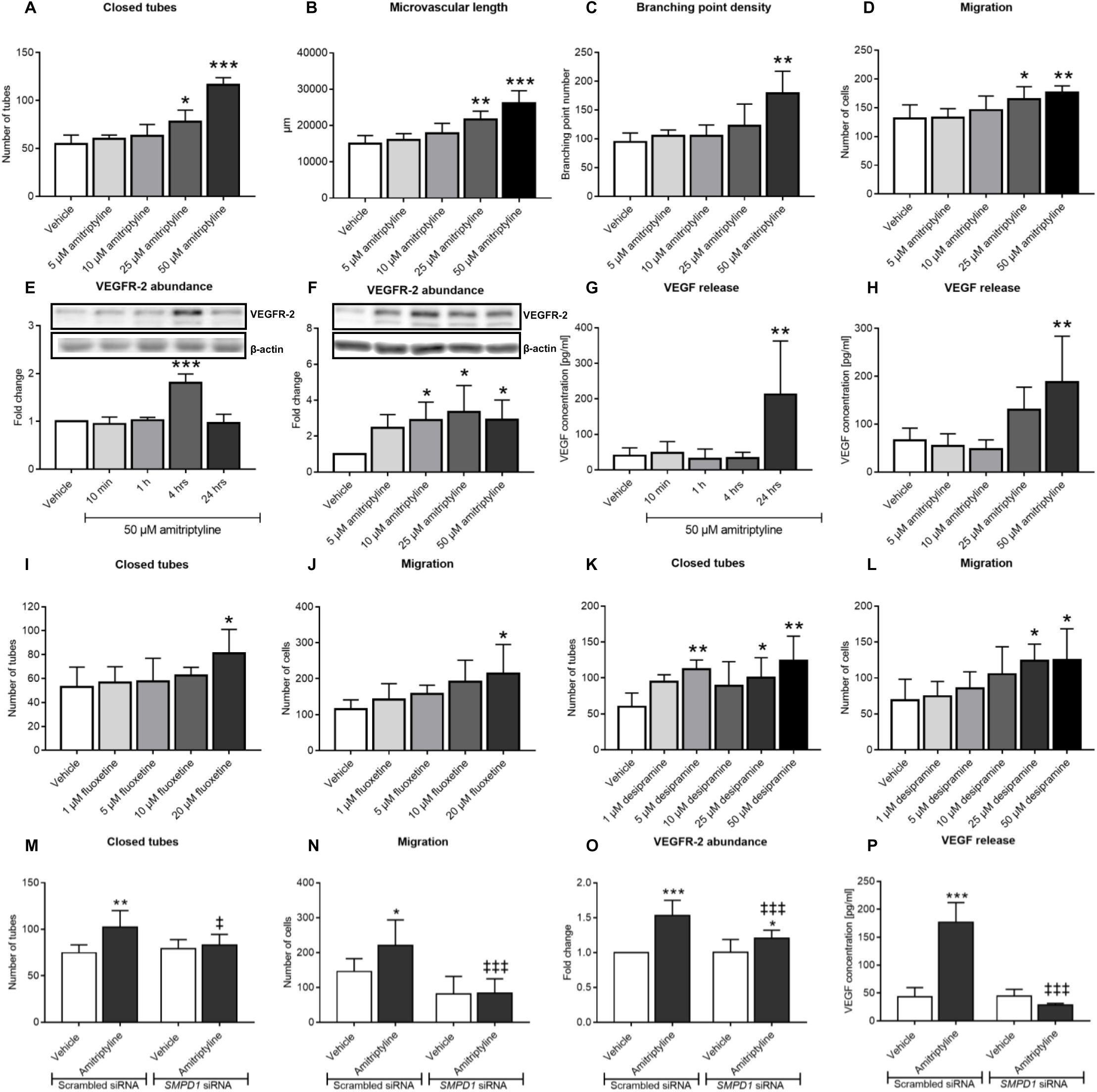
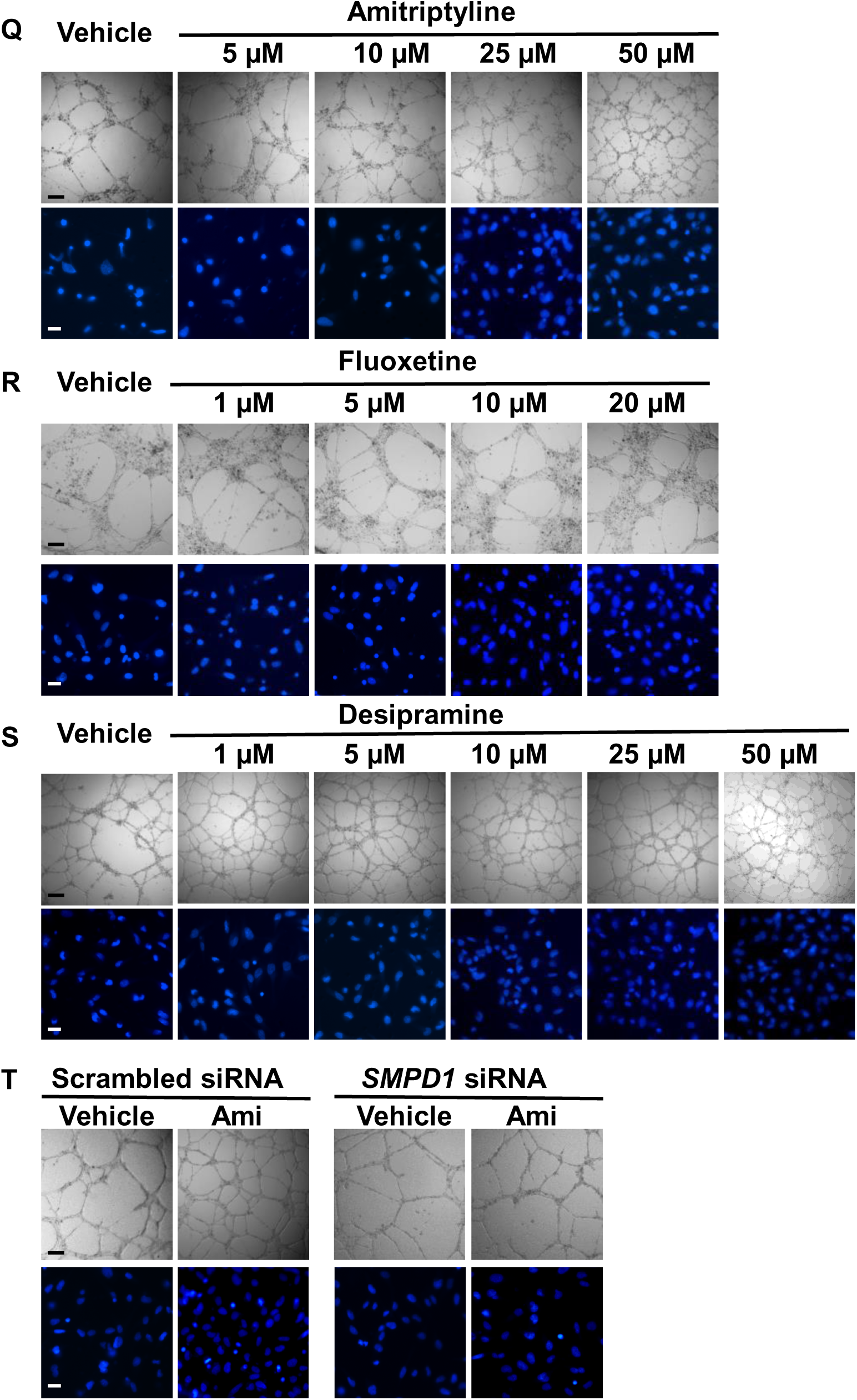
ASM inhibitors promote cerebral angiogenesis *in vitro* in an ASM dependent way. **(A-C)** Matrigel-based tube formation evaluated for the number of closed tubes, microvascular length and branching point density, **(D)** transwell migration, **(E,F)** VEGFR2 abundance examined by Western blot and **(G,H)** VEGF concentration in supernatants measured by enzyme-linked immunosorbent assay (ELISA) of hCMEC/D3 exposed to vehicle or amitriptyline (0-50 µM). In **(F)** and **(H)**, analyses were made after 4 and 24 hours amitriptyline exposure, respectively. **(I)** Tube formation and **(J)** transwell migration of hCMEC/D3 exposed to vehicle or fluoxetine (0-20 µM). **(K)** Tube formation and **(L)** transwell migration of hCMEC/D3 exposed to vehicle or desipramine (0-50 µM). Note that all three ASM inhibitors increase angiogenesis. **(M)** Tube formation, **(N)** transwell migration, **(O)** VEGFR2 abundance and **(P)** VEGF concentration in supernatants of hCMEC/D3 transfected with scrambled siRNA (used as control) or *SMPD1* siRNA which were exposed to vehicle or amitriptyline (0-50 µM). In **(O)** and **(P)**, measurements were made after 4 and 24 hours amitriptyline exposure, respectively. Representative examples of tube formation and migration assays are shown in **(Q-T)**. Data are means ± SD values. *p≤0.05/**p≤0.01/***p≤0.001 compared with corresponding vehicle; ^‡^p≤0.05/^‡‡^p≤0.01/^‡‡‡^p≤0.001 compared with corresponding scrambled siRNA (n=3-7 independent samples/group [in **(A-L)**]; n=5-8 independent samples/group [in **(M,N,O)**]; n=3 independent samples/group [in **(P)**]; analyzed by one-way ANOVA [in **(A-L)**] or two-way ANOVA [in **(M-P)**], followed by LSD tests). Bars in **(Q-T)**, 200 µm in tube formation photographs; 20 µm in transwell migration photographs.

In order to investigate if the effect of ASM inhibitors on angiogenesis *in vitro* was related to the inhibition of ASM, we next downregulated *SMPD1* expression by siRNA and examined ASM activity, sphingolipid levels, tube formation, migration and VEGFR2 abundance of hCMEC/D3 in response to amitriptyline. *SMPD1* mRNA level, ASM protein abundance and ASM activity were efficiently reduced by siRNA knockdown by 95.0±1.0%, 86.1±7.4% and 70.1±15.5%, respectively (**Suppl. Fig. 8A-C**). In response to I/R, *SMPD1* knockdown reduced the number of ceramide-rich intracellular vesicles by 65.6±19.5% (**Suppl. Fig. 9A, B**). *SMPD1* knockdown did not change the overall ceramide content of hCMEC/D3 assessed by LC-MS/MS (**Suppl. Fig. 10A-F**), but increased hCMEC/D3 sphingomyelin levels in cells cultured under non-ischemic control conditions (**Suppl. Fig. 11A-F**). This effect similarly affected long-chain (C16) and very long-chain (C20, C22, C24:1) sphingomyelins. Amitriptyline increased the tube formation, migration, VEGFR2 abundance and VEGF secretion of hCMEC/D3 exposed to scrambled (control) siRNA, but not in hCMEC/D3 exposed to *SMPD1* siRNA (Fig. 4M-P; **Suppl. Fig. 12A, B**). hCMEC/D3 viability was not influenced by amitriptyline after *SMPD1* knockdown (**Suppl. Fig. 6D**). These results confirmed that the induction of angiogenesis was ASM dependent.

### Ceramide-rich vesicles formed upon I/R are late endosomes/ multivesicular bodies (MVBs)

We next explored the nature of the ceramide-rich intracellular vesicles. In histochemical studies, ceramide immunoreactivity did not colocalize with markers of mitochondria (AIF), early endosomes (EEA1), lysosomes (Lamp1), autophagosomes (LC3b) or caveolae (caveolin) (**Suppl. Fig. 13**). In contrast, partial colocalization was found with markers of late endosomes (Rab7) (Fig. 5A) and MVBs (CD63) (Fig. 5B). MVBs are formed from maturing endosomes by the budding of their limiting membrane into their interior to form intraluminal vesicles called exosomes, which typically have a size of 70-150 nm and which, when released into the extracellular space, have important roles in cell communication ^26^.

**Figure 5.**
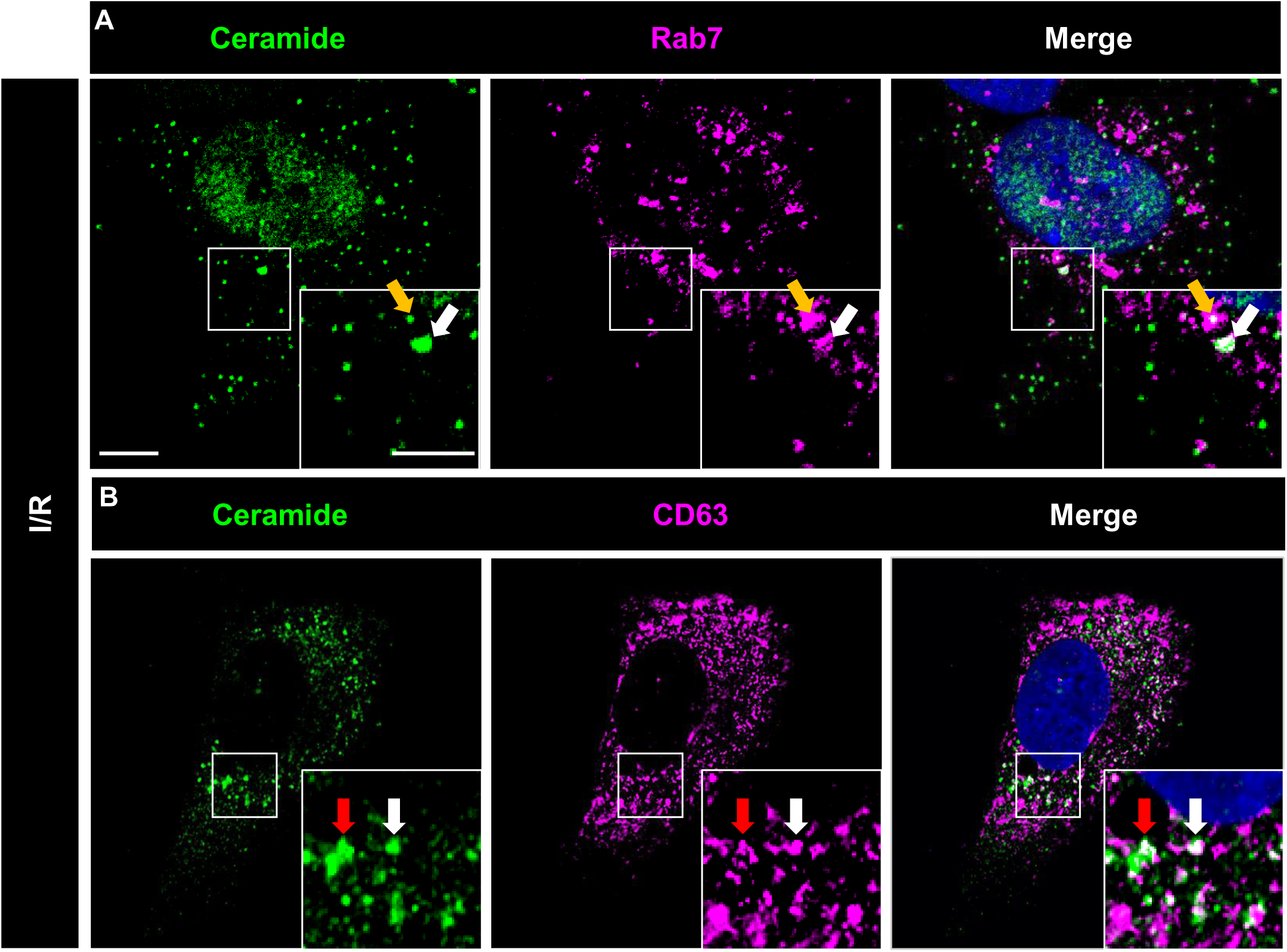
Intracellular ceramide-rich vesicles express markers of late endosomes and multivesicular bodies. Immunocytochemistry for **(A)** ceramide (in green) and the late endosome marker Rab7 (in magenta) and **(B)** ceramide (in green) and the multivesicular body marker CD63 (in magenta) of hCMEC/D3 exposed to 24 hours ischemia followed by 3 hours reoxygenation/glucose re-supplementation (I/R). In the merged photographs, double labeled cells are shown in white (selected cells labeled with arrow; nuclei were counterstained in blue with DAPI). Scale bar in overview photograph, 10 µm; in magnification, 5 µm. Data are representative for 3 independent studies.

### ASM inhibition or *SMPD1* knockdown induces the release of small extracellular vesicles (sEVs) by endothelial cells, which have *bona fide* characteristics of exosomes

Since the number of intracellular ceramide-rich vesicles decreased upon ASM inhibition (see above), we hypothesized that ASM inhibition may have stimulated exosome secretion resulting in the intracellular loss of vesicles. To clarify this hypothesis, we quantified sEVs in the supernatants of hCMEC/D3 exposed to non-ischemic control conditions (C), OGD (labeled I) or OGD followed by reoxygenation/glucose re-supplementation (labeled I/R) which were treated with amitriptyline for 3 or 24 hours or exposed to *SMPD1* siRNA knockdown by flow cytometry (**Suppl. Fig. 14**). I/R increased the number of tetraspanin CD9^+^ and CD63^+^ sEVs in supernatants (Fig. 3E-H). This number was further elevated after I/R by amitriptyline (Fig. 3E-H). Likewise, *SMPD1* siRNA knockdown increased the release of CD9^+^ and CD63^+^ sEVs by hCMEC/D3 (**Suppl. Fig. 9C-F**). Nanoparticle tracking analysis of sEV preparations revealed an average sEV size of 103±35 nm, which is in the range of exosomes (**Suppl. Fig. 15A**). Western blots revealed the presence of exosomal markers syntenin and CD9 and the absence of the cellular contamination marker calnexin in sEV preparations (**Suppl. Fig. 15B-D**). The overall protein content of sEVs was not influenced by functional ASM inhibition (vehicle: 7.38±3.64 pg/1000sEVs; amitriptyline: 8.87±9.04 pg/1000sEVs). Transmission electron microscopy confirmed that sEVs had the size and cup shape configuration of exosomes (**Suppl. Fig. 16**).

**Figure 6.**
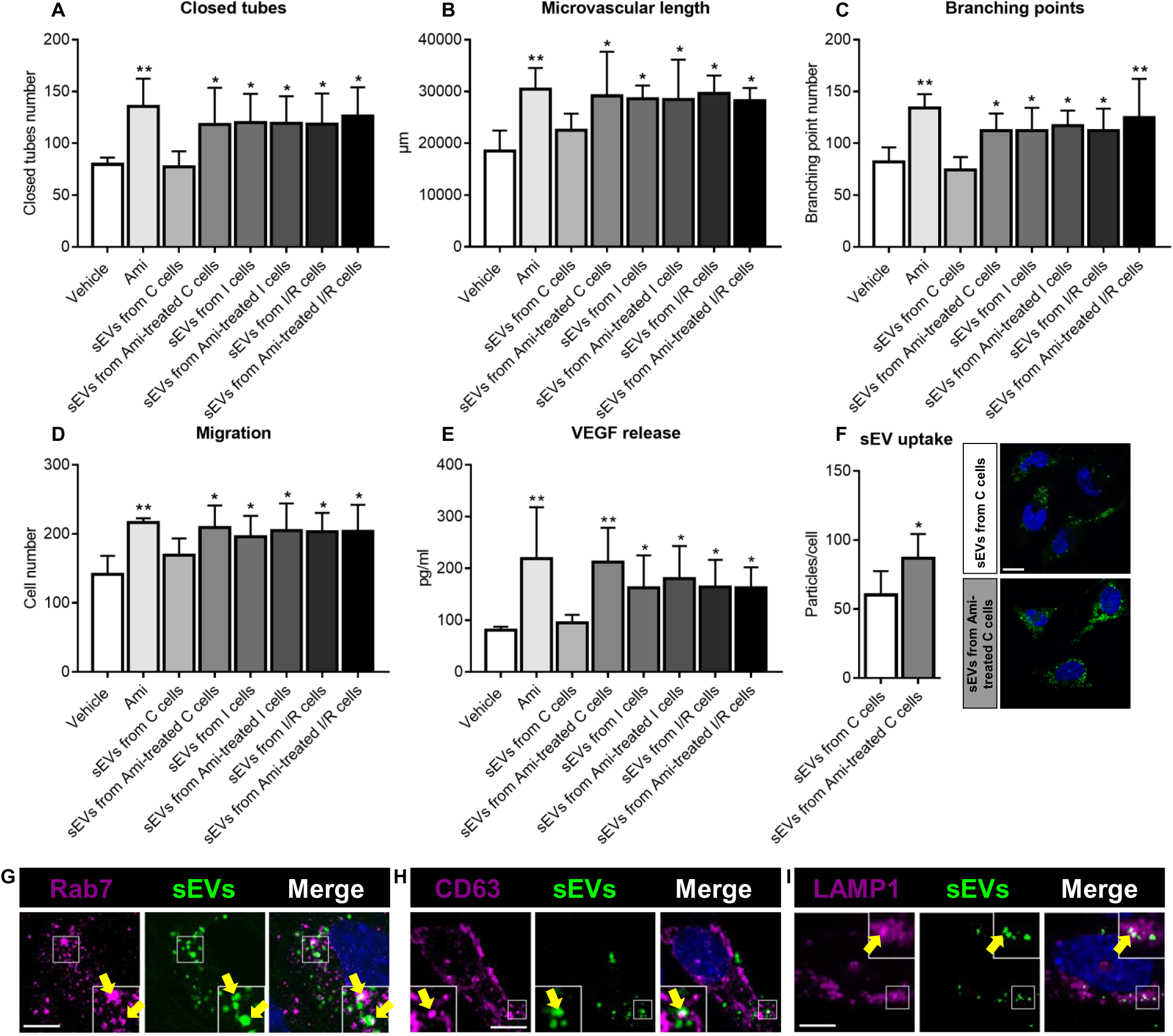
sEVs obtained from supernatants of endothelial cells exposed to ASM inhibitor amitriptyline have angiogenic activity that resembles sEVs released by endothelial cells during I/R. **(A-C)** Matrigel-based tube formation, **(D)** transwell migration and **(E)** VEGF release of hCMEC/D3, which were treated with vehicle, amitriptyline (Ami, 50 µM) or sEV preparations (25 µg protein/ml) isolated from the supernatants of hCMEC/D3 that had been cultured in non-ischemic control condition (C), ischemia (I) or ischemia followed by reoxygenation/glucose re-supplementation (I/R) and had been treated with vehicle or amitriptyline (50 µM) during cell cultivation. VEGF release was determined by ELISA. Representative tube formation and migration assays are shown in **Suppl. Fig. 16. (F)** hCMEC/D3 uptake of sEV preparations obtained from hCMEC/D3 treated with vehicle or amitriptyline (50 µM) evaluated by PKH67 dye. Representative photographs showing PKH67 dye (in green) in recipient hCMEC/D3 are depicted. The dye accumulated in intracellular vesicles, which based on their expression of **(G)** Rab7, **(H)** CD63 and **(I)** Lamp1 (in magenta) were vesicles belonging to the late endosomal compartment. In **(F)-(I)** nuclei were counterstained in blue with DAPI. Data are means ± SD values. *p≤0.05/**p≤0.01 compared with corresponding vehicle (n=4 independent samples/group [in **(A-C)**]; n=3 independent samples/group [in **(D)**]; n=3-4 independent samples/ group [in **(E)**]); n=5 independent samples/group [in **(F)**]; analyzed by one-way ANOVA, followed by LSD tests [in **(A-E)**] or two-tailed t tests [in **(F)**]).

### sEVs released upon ASM inhibitor exposure have angiogenic activity resembling sEVs released upon I/R

Depending on their source and pathophysiological conditions, sEVs have been shown to enhance neurological recovery, brain remodeling and angiogenesis post-I/R ^27, 28^. We thus hypothesized that sEVs mediated the angiogenic effects of ASM inhibitors. To test this assumption, we exposed hCMEC/D3 to sEV preparations obtained from supernatants of hCMEC/D3, which had been submitted to non-ischemic control conditions (C), ischemia (I) or ischemia followed by reoxygenation/glucose re-supplementation (I/R) and treatment with vehicle or amitriptyline. As positive control, amitriptyline was administered to hCMEC/D3. As reported above, amitriptyline increased the tube formation, migration and VEGF secretion of hCMEC/D3 (Fig. 6A-E; **Suppl. Fig. 17A, B**). sEVs isolated from supernatants of non-ischemic C hCMEC/D3 did not induce hCMEC/D3 tube formation, migration or VEGF secretion, whereas sEVs from amitriptyline treated control hCMEC/D3 increased hCMEC/D3 tube formation, migration and VEGF secretion similar to sEVs obtained from hCMEC/D3 exposed to I or I/R (Fig. 6A-E; **Suppl. Fig. 17A, B**). It had previously been shown that sEVs may obtain angiogenic characteristics when their parent cells are exposed to hypoxia ^29, 30^. Importantly, amitriptyline did not further elevate the angiogenic activity of sEVs from supernatants of cells exposed to I or I/R (Fig. 6A-E; **Suppl. Fig. 17A, B**), indicating that amitriptyline mimicked the effect of I or I/R but did not potentiate it.

### sEVs released upon ASM inhibitor exposure show enhanced uptake by endothelial cells

To elucidate how sEVs released upon ASM inhibitor exposure induce angiogenesis, we next studied sEV uptake by hCMEC/D3. For this purpose, we labeled sEV preparations obtained from vehicle and amitriptyline treated non-ischemic C hCMEC/D3 with a fluorescent dye (PKH67) and then exposed hCMEC/D3 with the labeled preparations. Although sEV preparations obtained from supernatants of endothelial cells exposed to vehicle were reproducibly taken up by hCMEC/D3, this uptake was significantly increased for preparations obtained from supernatants of cells exposed to amitriptyline (Fig. 6F). Immunostaining of recipient hCMEC/D3 revealed the accumulation of labeled particles in Rab7^+^ late endosomes (Fig. 6G), CD63^+^ MVBs (Fig. 6H) and Lamp1^+^ lysosomes (Fig. 6I).

### sEVs released by ASM inhibitor carry a protein signature associated with phagocytosis, protein export, extracellular matrix interaction and focal adhesion

By label free proteomics analysis we quantified 1163 proteins with ≤2 unique peptides in sEV preparations from supernatants of hCMEC/D3 cultured under normoxia that were treated with vehicle or amitriptyline. Of these, 111 proteins were up-regulated and 9 proteins were down-regulated by at least 2-fold with a p value ≤0.05 in sEVs from amitriptyline treated hCMEC/D3 compared with sEVs from vehicle treated hCMEC/D3 (**Suppl. Fig. 18**; **Suppl. Tables 1 and 2**). KEGG pathway database analysis (https://www.genome.jp/kegg/pathway.html, updated January 2021) showed that up-regulated proteins involved proteins with roles in phagosomes (VAMP3, CANX, ITGA5, LAMP1, STX7, HLA-C, HLA-B), protein export (SRP68, SRP72, SRP54) and lysosomes (CTSD, SCARB2, GUSB, GLB1, AP3S1, LAMP1, CTSB, AP1M1) (**Suppl. Tables 1 and 3**). Downregulated proteins were implicated in extracellular matrix -receptor interactions (TNC, COL4A2) and focal adhesion (TNC, COL4A2) (**Suppl. Tables 2 and 3**). A list of all quantified proteins containing original data of proteomic measurements is given in **Suppl. Table 4**.

## Discussion

We herein show that the antidepressants amitriptyline, fluoxetine and desipramine, which are diverse regarding their chemical structure and neurotransmitter mode of action, potently promote cerebral angiogenesis in an ASM dependent way, enabling successful remodeling of ischemic brain tissue. Reconstruction of microvascular anatomy using detailed 3D light sheet microscopy analysis ^23, 24^ allowed us to obtain a comprehensive set of microvascular network characteristics in mice exposed to MCAO that included microvascular density, branching point density and mean branch length ^23, 24^. The *in vivo* findings were complemented by well-established tube formation, migration and viability assays *in vitro* using hCMEC/D3 ^31, 32^. *Smpd1^-/-^* abolished angiogenic drug effects *in vivo*, as did siRNA-mediated *SMPD1* knockdown *in vitro*. These data proved that ASM inhibition mediated the angiogenic effects of the antidepressants. All three antidepressants, like several other antidepressants, had been previously shown to act as functional inhibitors of the ASM ^25^.

The observation that Asm inhibition promotes angiogenesis post-I/R and enhances post-ischemic brain remodeling expands previous findings and puts them in a new perspective. In rats, fluoxetine (10 mg/d) reduced infarct volume, nuclear factor-κB (NFκB) activation, microglial activation, brain leukocyte infiltration and neurological deficits, when administered with up to 9 hours delay in rats exposed to intraluminal MCAO ^33^. In mice, fluoxetine (10 mg/d) or sertraline (20 mg/d) reduced infarct volume when administered with 1 hour or 3 days delay in a photothrombotic stroke model ^22, 34^. Also in mice, fluoxetine (10 mg/d) increased neurogenesis in the dentate gyrus and spatial memory, when administered with one week delay after transient intraluminal MCAO for 4 weeks ^35^. Again in mice, nortriptyline (4 mg/kg) reduced infarct volume via mechanisms involving reduced mitochondrial cytochrome C, smac/Diablo and apoptosis-inducing factor release, when administered before transient intraluminal MCAO ^36^. To the best of our knowledge, recovery-promoting effects of nortriptyline, amitriptyline or other tricyclic or tetracyclic antidepressants have so far not been studied after I/R. This is surprising, since antidepressants are frequently prescribed in clinics for the treatment of post-stroke depressive disorder. The promotion of angiogenesis might represent the mechanism via which neurogenesis is induced by ASM inhibitors. Angiogenesis and neurogenesis are tightly linked post-stroke ^37–39^.

Post-I/R, Asm activity and ceramide level are known to be increased in the mouse and rat brain ^19–21^. LC-MS and MS imaging studies in mouse models of permanent or transient MCAO revealed that C18 ceramide, according to our data the most abundant brain ceramide species, was elevated in ischemic brain tissue ^40, 41^. One study after transient MCAO in mice found that specifically very long-chain (C22, C24) ceramides were elevated in the ischemic brain ^42^, whereas another study after photothrombotic stroke in mice also observed elevated long-chain (C16) ceramide ^22^. In immunohistochemical studies we now showed ceramide formation *in vivo* post-I/R in cerebral microvessels. Both long-chain (C16, C18) and very long-chain (C20, C22, C24:1) ceramides were increased, and amitriptyline reduced the ceramide levels in microvessels. Formation of ceramide in cerebral microvessels and release into the brain parenchyma has previously been observed in the mouse hippocampus in a model of glucocorticosterone-induced depression ^43^. In glucocorticosterone-induced depression, amitriptyline prevented the formation and release of ceramide from microvessels ^43^. The delivery of neutralizing anti-ceramide antibody reversed the effects of hippocampal extracts on PC12 cell proliferation ^43^. Overactivation of Asm has previously been described in cerebral endothelial cells in the aged mouse brain, associated with blood-brain barrier disturbance by increasing caveolae-mediated transcytosis ^44^. miR-induced *Smpd1* knockdown and tamoxifen-induced endothelial-specific *Smpd1* knockout reduced blood-brain barrier disturbance, neuronal degeneration in cortex and hippocampus and cognitive impairment in aged mice ^44^. It is interesting to note that I/R injury resembles glucocorticosterone-induced depression as well as brain ageing in that all three pathologies exhibit microvascular ASM/ ceramide pathway overactivity. It is tempting to speculate to which degree microvascular integrity is compromised in the depressed brain alongside Asm/ ceramide overactivation and whether the promotion of microvascular integrity and sprouting contributes to the mood-stabilizing effects of antidepressant drugs in line with enhanced neurogenesis.

In this study, cerebral human microvascular endothelial cells were found to form ceramide-rich late endosomes/ MVBs upon I/R, from which sEVs with *bona fide* exosome characteristics were released in response to ASM deactivation, which induced angiogenesis. In oligodendroglial precursor cells, the budding of exosomes from late endosomes was found to be independent of endosomal sorting complex required for transport (ESCRT), but ceramide and NSM2 dependent ^45, 46^. The NSM inhibitor GW4869 prevented exosome formation, as did siRNA-mediated NSM2 knockdown ^45, 46^. ASM and NSM are localized on the opposite endosomal membrane leaflets, ASM on the luminal leaflet and NSM on the cytosolic leaflet ^47^. Concentration gradients of ceramides on the two leaflets are thought to determine membrane curvature, possibly as a consequence of the cone-shaped physicochemical properties of ceramide ^45^. A negative curvature is induced on the side of ceramide accumulation which initiates membrane budding towards the opposite side ^45^. Further expanding these previous findings we propose that intraluminal membrane budding may be determined by the balance of sphingomyelinase activities on both membrane leaflets and that reduced ASM activity on the luminal leaflet in addition to increased NSM activity might promote intraluminal vesicle budding. According to our study, NSM2 (*SMPD3*) expression and NSM activity in endothelial cells is low. This observation might explain why ASM controls exosome release in endothelial cells. Besides ceramides, sphingomyelins critically control membrane fluidity and fusogenicity, increasing exosome release and uptake particularly at low pH ^48, 49^. Notably, *SMPD1* knockdown increased hCMEC/D3 sphingomyelin content. The concentration of sphingomyelins in hCMEC/D3 is ∼20 times higher than that of ceramides. It is therefore likely that the sphingomyelin changes contributed to the increased sEV release and uptake, generating a class of sEVs that in view of the inhibition of the proinflammatory ASM/ ceramide pathway confers restorative activity. These sEVs constitute an elegant target, via which stroke recovery can now be amplified.

## Online Methods

### Legal issues, animal husbandry and randomization

Experiments were performed with local government approval (State Agency for Nature, Environment and Consumer Protection, Recklinghausen) in accordance to E.U. guidelines (Directive 2010/63/EU) for the care and use of laboratory animals. Experiments were strictly randomized and blinded at all stages of the study. In the animal experiments, the investigator performing the surgeries and histochemical studies (A.M.Y.) was blinded by another researcher (N.H.) preparing the treatment solutions, which received dummy names (solution A, B, C) and were decoded after termination of the study. Animals were kept in a regular 12h:12h light/dark cycle in groups of 5 animals/cage with free access to food and water. Throughout the study, the animals had free access to food and water. The data that support the findings of this study are available from the corresponding author upon reasonable request.

### Middle cerebral artery occlusion (MCAO)

Focal cerebral ischemia was induced in male C57BL/6j wildtype mice (8-10 weeks; 22-25 g; Harlan Laboratories, Darmstadt, Germany), in *sphingomyelinase phosphodiesterase-1* [*Smpd1*]^-/-^) (that is, ASM deficient) mice on C57BL/6j background and their *Smpd1^+/+^* C57BL/6j littermates by 20 min left-sided intraluminal MCAO ^50, 51^. Mice were anesthetized with 1.0-1.5% isoflurane (30% O2, remainder N2O). Rectal temperature was maintained between 36.5 and 37.0°C using a feedback-controlled heating system. Cerebral blood flow was recorded by laser Doppler flow (LDF) measurement using a flexible probe with a diameter of 0.5 mm attached to the animals’ skull above the core of the middle cerebral artery territory. For MCAO, a midline neck incision was made and the left common and external carotid arteries were isolated and ligated. The internal carotid artery (ICA) was temporally clipped. A silicon-coated monofilament was introduced via a small incision into the common carotid artery (CCA) and advanced to the carotid bifurcation for MCAO. Reperfusion was initiated by monofilament removal. Starting immediately after reperfusion onset (animal sacrifice after 24 hours) or 24 hours after reperfusion (animal sacrifice after 14 days), vehicle or amitriptyline (2 or 12 mg/kg b.w.; Sigma-Aldrich, Deisenhofen, Germany) were intraperitoneally administered b.i.d. for up to 14 days. Wounds were carefully sutured. The opioid buprenorphine (0.1 mg/kg; Reckitt Benckiser, Slough, U.K.) was subcutaneously administered before surgery, and the antiphlogistic carprofen (4 mg/kg; Bayer Vital, Leverkusen, Germany) was subcutaneously administered daily for up to three days after surgery. At the indicated time-points, animals were deeply anesthetized with isoflurane and transcardially perfused with 40 ml 0.1 M phosphate-buffered saline (PBS) (animals used for histochemistry, activity assays and mass spectrometry) or with 40 ml 0.1 M PBS supplemented with heparin (50 U/ml) followed by 40 ml 4% paraformaldehyde (PFA) in 0.1 M PBS (animals used for light sheet microscopy).

### Infarct volumetry

Brains perfused with 0.1 M PBS were frozen on dry ice and cut into 20 µm thick coronal sections. Sections were collected at 1 mm intervals for cresyl violet staining. On these sections, the border between infarcted and non-infarcted tissue was outlined using Image J (National Institutes of Health [NIH], Bethesda, MD, U.S.A.). Infarct volume was determined by subtracting the volume of the non-lesioned ipsilateral hemisphere from the volume of the contralateral hemisphere ^51^. Edema volume was calculated as volume difference between the ipsilateral and the contralateral hemisphere ^51^.

### FITC-albumin hydrogel perfusion and whole-brain clearing

Immediately following transcardiac PFA fixation, 10 ml of a hand-warm (30°C) 2% gelatin hydrogel containing 0.1% FITC-conjugated albumin, which had been filtered using Whatman filter paper (GE Healthcare Life Science, Little Charfont, U.K.) and was protected from light, was transcardially infused into the animals’ aorta. Brains were subsequently removed, post-fixed overnight at 4°C in 4% PFA in 0.1 M PBS and dehydrated through a 30%, 60%, 80% and 100% tetrahydrofuran (THF; Sigma-Aldrich) gradient ^23^. Brain clearing was achieved with ethyl cinnamate (ECI; Sigma-Aldrich).

### 3D light sheet fluorescence microscopy (LSM) and microvasculature analysis

The FITC-albumin labeled vasculature of cleared brains was scanned by a light sheet microscope (Ultramicroscope II, LaVision BioTec, Göttingen, Germany) that was equipped with a 488 nm laser. Horizontal overview images of the cleared brain were taken using a 1.6x objective. Serial images of the striatum and cortex were acquired at 2 µm steps using a 6.4x objective. In each animal, two regions of interest (ROI) measuring 500 µm x 500 µm x 1000 µm (in the X, Y and Z planes, respectively) in the dorsolateral striatum were analyzed using Imaris (Bitplane, Zurich, Switzerland) software with 3D rendering software package, as described previously ^23^. Following image segmentation, skeletonization and 3D reconstruction, microvascular network characteristics, that is, microvascular length, branching point number and mean branch length between two branching points, were determined.

### Cell culture

Brain microvascular endothelial cells (hCMEC/D3) were cultured in endothelial basal medium (EBM-2, Lonza, Basel, Schweiz) supplemented with 5% fetal bovine serum (FBS, Life Technologies, Carlsbad, CA, U.S.A.), 100 U/ml penicillin/ streptomycin (Life Technologies), 1.4 µM hydrocortisone (Sigma-Aldrich), 5 µg/ml ascorbic acid (Sigma-Aldrich), 1% chemically defined lipid concentrate (Life Technologies), 10 mM HEPES (Life Technologies) and 1 ng/ml basic fibroblast growth factor (bFGF, Sigma-Aldrich). hCMEC/D3 were seeded on 150 µg/ml collagen (R&D Systems, Minneapolis, MN, U.S.A.) pre-coated flasks and kept at 37°C at 5% CO2. Oxygen-glucose deprivation (OGD) was induced by incubating the cells in a hypoxia chamber (1% O2, Toepffer Lab Systems, Göppingen, Germany) with glucose-free medium (Life Technologies). For comparative studies on sphingomyelinase expression and activities, primary human brain microvascular endothelial cells (HBMEC; catalog #1000; ScienCell^TM^ Research Laboratories, Carlsbad, CA, U.S.A.) cultured in endothelial cell growth medium MV (ECGM-MV, PromoCell, Heidelberg, Germany) containing 100 U/ml penicillin/ streptomycin (Life Technologies) and growth medium MV supplement mix (PromoCell) and human umbilical vein endothelial cells (HUVEC) cultured in endothelial cell growth medium (ECGM, PromoCell) containing 100 U/ml penicillin/ streptomycin (Life Technologies) and growth medium supplement mix (PromoCell) were used.

### Small interfering RNA (siRNA) knockdown

ASM knockdown *in vitro* was achieved by small interfering RNA (siRNA). Transfections were performed according to the manufacturer’s instructions using Dharmafect transfection reagents (Dharmacon, Lafayette, CO, U.S.A.). Scrambled siRNA was used as a negative control.

### Tube formation assay

To evaluate the formation of capillary-like tubular structures ^31^, 60 µl matrigel (Corning, NY, U.S.A.) were pipetted into 96-well plates. The gel solidified at 37°C for 30 minutes. 3×10^4^ cells were seeded and treated with ASM inhibitors and/or sEVs of interest. After 20 hours, photomicrographs were taken with a 2x objective using the EVOS digital inverted microscope (Advanced Microscopy Group, Bothell, WA, U.S.A.). Closed tubes were counted using Image J (NIH) in each well. Microvascular length and branching point density were evaluated. Experiments were done in triplicates for which mean values were formed.

### Transwell migration assay

Cell migration was assessed using a modified Boyden chamber. 3×10^4^ cells were seeded in the upper compartment of polycarbonate membrane inserts (8.0 μm pores) that contained serum-reduced (1.25% FBS) medium. Treatments of interest were administered into the lower compartment that contained 5% FBS. Cells that did not migrate were removed after 24 hours. Migrated cells were fixed with 4% PFA and stained with Hoechst 33342 (Thermo Fisher Scientific, Waltham, MA, U.S.A.). Photomicrographs were taken with a 20x objective using the EVOS digital inverted microscope (Advanced Microscopy Group). In each chamber, migrated cells were counted using Image J (NIH) in 8 ROIs measuring 530 x 400 µm. Experiments were done in duplicates for which mean values were formed.

### Cell viability assay

2×10^4^ hCMEC/D3 were seeded into 96-well plates and treated with ASM inhibitors and/or sEVs of interest. After 24 hours, cells were incubated with 0.5 mg/ml 3-(4,5-dimethyl-2-thiazolyl)-2,5-diphenyl-2H-tetrazolium bromide (MTT; Biomol, Hamburg, Germany) for 2 hours. Formazon formation was measured at 570 nm on a microplate reader (iMark Detection; Bio-Rad Laboratories, Hercules, CA, U.S.A.). Samples were analyzed in triplicates, of which mean values were formed.

### Sphingomyelinase activity assays

Brain samples obtained from the ischemic or contralateral middle cerebral artery territory or hCMEC/D3, HBMEC and HUVEC were lysed in 250 mM sodium acetate buffer (pH 5.0) containing 1% NP-40 detergent (Fluka BioChemika, Morristown, NJ, U.S.A.; for ASM activity measurement) or in 100 mM HEPES buffer (Life Technologies) (pH 7.4) containing 5 mM magnesium chloride (Sigma-Aldrich) and 0.1% NP-40 detergent (Fluka BioChemika; for neutral sphingomyelinase [NSM] activity measurement). The cellular membrane integrity was disrupted with a sonicator. After centrifugation for 5 minutes at 300 g at 4°C, supernatants were collected. Lysates were adjusted to a specific protein concentration and incubated with 100 pmol BODIPY-labeled sphingomyelin (Thermo Fisher Scientific) in 250 mM sodium acetate (pH 5.0) and 0.1% NP-40 for 1 hour at 37°C. Chloroform:methanol (2:1, v/v) was added, samples were vortexed and centrifuged for 5 minutes at 15,000 g to achieve a phase separation. The lower phase was collected and concentrated in a vacuum centrifuge (SPC111V, Thermo Fisher Scientific) for 45 minutes at 37°C. Lipids were dissolved in 20 µl chloroform:methanol (2:1, v/v) and spotted onto thin layer chromatography (TLC) plates (Macherey Nagel, Düren, Germany). The TLC run was performed with chloroform:methanol (80:20, v/v). TLC plates were analyzed with a Typhoon FLA 9500 scanner (GE Healthcare Life Sciences) and lipid spots were quantified with Image Quant (GE Healthcare Life Sciences).

### Real-time quantitative polymerase chain reaction

RNA was isolated according to the phase extraction method using Trizol (Life Technologies)/ Chloroform (Sigma-Aldrich) and treated with DNase I (Life Technologies). Complementary DNA (cDNA) was generated by reverse transcription with SuperScript II (Life Technologies) using oligo(dt) primers and random hexamers. Real-time quantitative polymerase chain reaction (RT-qPCR) was performed with SYBR-Green (Life Technologies) in a StepOnePlus real-time PCR system with primers for human *SMPD1* (Gene bank number: NM_000543, Biomol), human *SMPD2* (Gene bank number: NM_003080, Biomol), human *SMPD3* (Gene bank number: NM_018667, Biomol) and human *β-actin* (Gene bank number: NM_001101.2, Biomol) as a housekeeping gene. Results were quantified using the 2^−ΔΔCt^ method. Samples were analyzed in triplicates, of which mean values were formed.

### Liquid chromatography tandem-mass spectrometry (LC-MS/MS) of sphingolipids

Following lipid extraction with methanol:chloroform (2:1, v/v) as described ^52^, ceramides and sphingomyelins were quantified by LC-MS/MS using a 6490 triple-quadrupole mass spectrometer (Agilent Technologies, Waldbronn, Germany) operating in the positive electrospray ionization mode (ESI+) ^53^. Quantification was performed with MassHunter Software (Agilent Technologies). Sphingolipid amounts were normalized to protein content (*in vivo* experiments) or cell numbers (*in vitro* experiments. As such, concentrations per mg protein (*in vivo* experiments) or per 1 million cells (*in vitro* experiments) were calculated.

### Immunohistochemistry/ immunocytochemistry

Coronal brain sections obtained from the level of the midstriatum (bregma 0.0 mm; that is, the core of the middle cerebral artery territory) or hCMEC/D3 seeded on sterile coverslips were fixed with 4% PFA in 0.1 M PBS and immersed in 0.1 M PBS containing 0.1% Triton X-100, 5% normal donkey serum or 10% normal goat serum and 1 or 2.5% bovine serum albumin. Samples were incubated overnight at 4°C in mouse anti-NeuN (A60; Merck Millipore, Burlington, MA, U.S.A.), rat anti-CD31 (MEC 13.3; BD Biosciences, Franklin Lakes, NJ, U.S.A.), rat anti-CD45 (30-F11; BD Biosciences, Franklin Lakes, NJ, U.S.A.), biotinylated goat anti-IgG (sc-2039; Santa Cruz, Heidelberg, Germany), mouse anti-ceramide (S58-9; Glycobiotech, Kükels, Germany), rabbit anti-apoptosis-inducing factor (AIF; D39D2; Cell Signaling Technology, Danvers, MA, U.S.A.), rabbit anti-early endosome antigen-1 (EEA1; C45B10; Cell Signaling Technology), rabbit anti-lysosomal-associated membrane protein-1 (Lamp1; 1D4B; Abcam, Cambridge, U.K.), rabbit anti-microtubule-associated protein light chain-3b (LC3b; 2775; Cell Signaling Technology), rabbit anti-caveolin (3238S; Cell Signaling Technology), rabbit anti-Rab7 (D95F2; Cell Signaling Technology) or rabbit anti-CD63 (LS-C204227; Lifespan Biosciences, Seattle, WA, U.S.A.) antibody. Samples were rinsed and labeled with appropriate secondary Alexa Fluor-594, Alexa Fluor-488, Cy3 or biotinylated antibody. In sections stained with fluorescent antibody, nuclei were counterstained with Hoechst 33342 (Thermo Fisher Scientific). Sections stained with biotinylated antibody were revealed by 3,3’-diaminobenzidine (DAB) staining using a avidin-biotin complex (ABC) peroxidase kit (Vectastain Elite Kit Standard, Vector Laboratories, Burlingame, CA, U.S.A.). Sections were evaluated using an inverted microscope equipped with apotome optical sectioning (Axio Observer.Z1; Carl Zeiss, Oberkochen, Germany). Sections were analyzed by counting the density of labeled microvessels or cells (NeuN, CD45) in the ischemic and contralateral striatum (*in vivo* studies) or by counting intracellular vesicles (ceramide) in 8 ROIs measuring 135 x 135 µm on coverslips (*in vitro* studies). IgG Immunohistochemistry was examined by densitometry in the ischemic and contralateral striatum ^54^.

### Western blot analysis of cell lysates

Cells were lysed with NP-40 buffer containing protease and phosphatase inhibitors (Sigma-Aldrich). Lysates were centrifuged at 13,400 rpm at 4°C and the supernatant was collected. Equal amounts of protein were separated by sodium dodecyl sulfate-polyacrylamide gel electrophoresis (SDS-PAGE) and subsequently transferred to nitrocellulose membranes (GE Healthcare Life Science). Non-specific binding was blocked with 5% non-fat milk powder (Sigma-Aldrich) dissolved in 0.1% Tween in 0.1 M Tris-buffered saline (TBS-T). Membranes were incubated with rabbit anti-vascular endothelial growth factor (VEGF) receptor-2 (VEGFR2) (55B11; Cell Signaling Technology), goat anti-ASM (AF5348, R&D Systems) and rabbit anti-β-actin (4967; Cell Signaling Technology) antibody overnight at 4°C, rinsed and incubated in peroxidase-conjugated secondary antibodies (Santa Cruz, Heidelberg, Germany) for 1 hour at room temperature. Signals were detected by enhanced chemiluminescence using prime Western blotting detection reagent (GE Healthcare Life Science). VEGFR2 and ASM expression were normalized to β-actin abundance.

### Enzyme-linked immunosorbent assay (ELISA)

VEGF was quantified by ELISA (R&D Systems) in supernatants of hCMEC/D3 in accordance to the manufacturer’s instruction.

### Preparation of small extracellular vesicles (sEVs)

hCMEC/D3 were cultured in triple flasks (Thermo Fisher Scientific). Media were collected after exposure of cells to the different experimental conditions and centrifuged at 2,000 g for 15 minutes at 4°C, followed by centrifugation at 10,000 g for 45 minutes at 4°C (5810R centrifuge, Eppendorf, Hamburg, Germany). Supernatants were filtered through a 0.22 µm filter (Sartorius, Göttingen, Germany) and supplemented with NaCl at a final concentration of 75 mM and polyethylene glycol-6000 (PEG; Sigma-Aldrich) at a final concentration of 10%. sEVs were concentrated at 1,500 g for 30 minutes at 4°C (Avanti J-26 XP centrifuge, Beckmann Coulter, Brea, CA, U.S.A.). Pellets were then dissolved in 0.9% NaCl (Sigma-Aldrich), transferred to ultra-clear centrifuge bottles (Beckmann Coulter) and precipitated by ultracentrifugation at 110,000 g for 130 min at 4°C (Optima L7-65, k factor: 133, Beckmann Coulter). sEV pellets were resuspended in 0.9% NaCl supplemented with 10 mM HEPES (Life Technologies) and stored in low retention microcentrifuge tubes (Kisker Biotech, Steinfurt, Germany) at −80°C until further use.

### Amnis ImagestreamX flow cytometry of sEVs

sEVs were quantified with an ImageStreamX MkII instrument (Merck Millipore) as described previously ^55^ after CD9-FITC (MEM-61; Exbio, Vestec, Czech Republic) and CD63-APC (MEM-259; Exbio) antibody staining. All samples were appropriately diluted in order to avoid coincidence or swarm detection. Data analysis was performed using Amnis IDEAS software (version 6.1).

### sEV uptake analysis

Following sEV preparation by PEG precipitation and ultracentrifugation from the supernatant of hCMEC/D3, as described, sEV preparations were labeled with PKH67 membrane dye (Sigma-Aldrich). Briefly, sEV preparations were stained with 2 µM PKH67 diluted in diluent C for 5 min at room temperature. Excess dye was removed by washing twice with PBS using Amicon-Ultra Centrifugal Filter Units (Merck Millipore). Labeled sEVs were incubated with hCMEC/D3 for 24 hours. Cells were fixed with PFA, labeled with rabbit anti-EEA1 (C45B10; Cell Signaling Technology), rabbit anti-Rab7 (D95F2; Cell Signaling Technology), rabbit anti-CD63 (LS-C204227; Lifespan Biosciences) or rabbit anti-Lamp1 (1D4B; Abcam) antibody, detected by appropriate secondary antibody (as above) and counterstained with Hoechst 33342 (Thermo Fisher Scientific).

### Characterization of sEV preparations by nanoparticle tracking analysis (NTA)

According to recently updated guidelines for the characterization of small extracellular vesicles ^56^, sEV preparations were analyzed for concentration and size by nanoparticle tracking analysis (NTA; Particle Metrix, Meerbusch, Germany), as described previously ^57^.

### Western blot analysis of sEV preparations

Protein concentrations in sEV preparations were determined by a standardized bicinchoninic acid (BCA) assay according the manufacturer’s protocol (Pierce, Rockford, IL, U.S.A.). For Western blot, 30 µg protein samples were solubilised with Laemmli sample buffer under reducing (containing dithiothreitol [DTT]; AppliChem, Darmstadt, Germany) or non-reducing (not containing DTT) conditions and separated on SDS-PAGE gels before transfer to polyvinylidene fluoride membranes (PVDF; Millipore, Darmstadt, Germany). Membranes were blocked in TBS-T supplemented with 5% (w/v) skim milk powder (Sigma-Aldrich). Membranes were stained with rabbit anti-syntenin (clone EPR8102; Abcam), rabbit anti-calnexin (ab10286; Abcam) or mouse anti-CD9 (clone VJ1/20.3.1; kindly provided by Francisco Sánchez, Madrid, Spain) antibodies. Anti-syntenin and anti-CD9 antibodies were used as sEV markers, anti-calnexin antibody as cellular contamination marker. Membranes were washed and counterstained with appropriate horseradish peroxidase-conjugated secondary antibodies (Santa Cruz) that were detected by enhanced chemiluminescence using prime Western blotting detection reagent (GE Healthcare Life Science).

### Transmission electron microscopy of sEV preparations

200 mesh copper grids (Plano, Wetzlar, Germany) were physically charged by a glow discharge procedure to allow strong adherence of particles to the electron microscopy grid during further processing. 4.5 µl of sEV preparations were added to the pre-treated grid surface and dried at room temperature. For subsequent removal of salts, grids were successively transferred on three droplets of deionized water. Samples were incubated with 1.5% phosphotungstate acid (Electron Microscopy Science, Hatfield, PA, U.S.A.) and dried. Images were acquired using a JEM 1400Plus electron microscope (JEOL, Tokyo, Japan) operating at 120 kV that was equipped with a 4096×4096 pixel CMOS camera (TemCam-F416; TVIPS, Gauting, Germany). Image acquisition software EMMENU (Version 4.09.83) was used for taking 16 bit images. Image post-processing was carried out using ImageJ software (Version 1.52b; NIH).

### LC-MS-based proteomics

For label free proteome analysis, sEV preparations obtained from supernatants of hCMEC/D3 cultured under normoxic conditions were exposed to vehicle or amitriptyline (50 µM) and lysed in 50 mM Tris-HCl, 150 mM NaCl and 1% sodium dodecyl sulfate (SDS) at pH 7.8 supplemented with 1 tablet cOmplete Mini and 1 tablet PhosSTOP (Roche, Basel, Switzerland) per 10 ml. Protein concentrations were determined using the bicinchoninic acid assay (Pierce, Thermo Fisher Scientific). Afterwards, cysteines were reduced by 30 min incubation at 56°C with 10 mM dithiothreitol and free sulfhydryl groups were alkylated with 30 mM iodoacetamide for 30 min at room temperature in the dark. Samples were processed using S-trap Micro Column Digestion Protocol (PROTIFI, Farmingdale, NY, USA) according to the manufacturer’s instructions ^58^ with slight modifications. In brief, carbamidomethylated samples were diluted with 10% SDS to a final concentration of 5% SDS. Afterwards, 4.43 µl of 12% phosphoric acid were added, followed by the addition of 292.38 µl of S-trap binding buffer (90% methanol, 100 mM triethylammonium bicarbonate (TEAB), pH 7.1). 165 µl of the acidified lysate/S-trap buffer mix was placed into the spin column and spun down in a bench-top centrifuge in a 1.5 ml tube at 4,000 g until all the solution had passed through. The flow through was discarded and the rest of the acidified lysate/S-trap buffer mix was loaded into the spin column and the procedure explained before was repeated.

Afterwards, 3 washing steps with 150 µl of S-trap binding buffer each were performed by centrifugation at 4,000 g. Then, sequencing grade modified trypsin (Promega, Madison, WI, U.S.A.) was added in an enzyme to a sample ratio of 1:10 (w/w) in 25 µl of 50 mM ammonium bicarbonate (ABC) containing 2 mM CaCl_2_. Spin columns were incubated for 1 h at 47°C. After incubation, peptides were recovered by centrifugation prior addition of 40 µl of 50 mM ABC to the spin columns. Further peptide recovery was done by adding 40 µl 0.1% of trifluoroacetic acid (TFA) and 35 µl of 50% acetonitrile (for recovering hydrophobic peptides) to the spin columns followed by centrifugation. Finally, samples were dried under vacuum and resuspended in 0.1% TFA. Digestion quality control was performed via a monolithic column-HPLC ^59^. NanoLC-MS/MS analysis was done using a U3000 RSLCnano online-coupled to a Q Exactive HF mass spectrometer (Thermo Fisher Scientific). Thus, peptides samples were loaded onto the trap column (Acclaim PepMap100 C18; 100 µm x 2 cm) in 0.1% TFA at a flow rate of 20 µl/min. After 5 min, the pre-column was switched in line with the main column (Acclaim PepMap100 C18; 75 μm × 50 cm) and peptides were separated using a 100 min binary gradient ranging from 5-24-40 of 84% acetonitrile in 0.1% formic acid at 60°C and a flow rate of 250 nl/min. The MS was operated in data dependent acquisition (DDA) mode with survey scans acquired at a resolution of 60,000 followed by 15 MS/MS scans at a resolution of 15,000 (top15). Selected precursor ions (highest intense) were isolated in a 1.6 *m/z* window and subjected to fragmentation by higher energy collision induced dissociation using a normalized collision energy of 27 eV. Automatic gain control target values were set to 1×10^6^ and 5×10^4^ and the maximum ion injection was set to 120 ms and 250 ms for MS and MS/MS, respectively, with 20 sec dynamic exclusion. Polysiloxane at *m/z* 371.1012 was used as internal calibrant ^60^. Raw data and label free quantification analysis were done with Proteome Discoverer 2.3 (Thermo Fisher Scientific). In the consensus workflow, the Feature Mapper, the Precursor Ion Quantifier and the Protein False Discovery Rate (FDR) Validator nodes were used. With the Feature Mapper node chromatographic runs were aligned based on the retention time. For the Precursor Ion Quantifier node parameters for peptides were set to Unique and Razor, precursor abundances were based on intensity, and sample normalization was based on total peptide amount. Target FDR was set to 0.01. For the processing workflow the Minora Feature Detector was used. Raw data were searched against the Uniprot human database (November 2019; 20,609 protein sequences). Mascot and Sequest were applied as search algorithms with the following settings: (1) trypsin as enzyme allowing two missed cleavages, (2) carbamidomethylation of cysteins (+57.0214 Da) as fixed modification, (3) oxidation of methionine (+15.9949 Da) as variable modification, and (4) mass tolerance for MS and MS/MS were set to 10 ppm and 0.02 Da, respectively. Raw data and Proteome Discoverer search results have been deposited in the ProteomeXchange repository with identifier PXD024106 (username, password: OYobSSEO).

### Statistical analysis

Data are expressed as mean ± standard deviation (SD). In case of multiple group comparisons, one-way or two-way analysis of variance (ANOVA) was used, followed by least significant difference (LSD) tests as posthoc tests. In case of comparisons between 2 groups, two-tailed t tests were used. All values were normally distributed. P values ≤0.05 were defined to indicate statistical significance. The statistical details (n numbers, mean ± SD and tests) are given in the figure legends. Sample size planning was conducted with G*Power version 3.1.7 software of the University of Düsseldorf, Germany. Statistical analyses were performed using GraphPad Prism version 7.0 software.

### Data and software ability

G*Power version 3.1.7 software of the University of Düsseldorf, Germany and Image J software are public programs available online without charge. GraphPad Prism version 7.0 software is commercially available via Graphpad Software (San Diego, CA, U.S.A.).

## Acknowledgments

The authors thank Britta Kaltwasser for technical assistance. Supported by the German Research Foundation (GRK2098 project 259317790 to D.M.H., E.G., R.K. and B.K.; project 389030878 to D.M.H. and M.G.; FOR-2879 project 405358801 to D.M.H., M.G. and C.K.; SFB/TR240 project 374031971 to C.K., SFB-1116 project 236177352 to A.S. and F.A.S.), German Federal Ministry of Education and Science (3DOS, to D.M.H. and B.G.; to A.S. and F.A.S.), North Rhine Westphalian Ministry for Culture and Science (to A.S. and F.A.S.) and E.U. (ERA-NET EuroTransBio-11 project EVTrust [031B0332B] to D.M.H. and B.G.).

## Author contributions

D.M.H., E.G. and R.K. designed the study. A.M.Y., N.H. and C.B. performed the animal experiments. A.M.Y., N.H., X.Z., M.Z. and T.H. conducted the cell culture studies. A.M.Y., N.H., X.Z., M.Z., T.H. and D.M.H. were responsible for histochemical and molecular biological experiments on tissues and cells, A.M.Y., N.H., M.G. and D.M.H. for light sheet microscopy studies on cleared whole brains. A.M.Y., X.Z., T.T., V.B., M.H. and B.G. performed the in-depth characterization of small extracellular vesicles. F.Sch. and B.K. conducted the sphingolipid mass spectrometry studies. F.A.S. and A.S. provided the proteome mass spectrometry analysis. D.M.H., A.M.Y. and E.G. drafted the manuscript, all authors revised it.

## Legends to supplementary figures

**Supplementary Figure 1. Acid sphingomyelinase (Asm) product ceramide accumulates in cerebral microvessels after ischemia/ reperfusion (I/R) *in vivo*.** Immunohistochemistry for ceramide in the contralateral non-ischemic control striatum (C) and reperfused ischemic striatum (I/R) of C57BL/6j mice exposed to transient middle cerebral artery occlusion (MCAO), which were intraperitoneally treated with vehicle or amitriptyline (2 or 12 mg/kg b.w., b.i.d.) immediately after I/R, followed by animal sacrifice after 24 hours. Bar, 50 µm. Data are representative for 3 independent studies.

**Supplementary Figure 2. Asm inhibitor amitriptyline does not change cerebral sphingomyelin levels after I/R *in vivo*.** Total sphingomyelin, C16 sphingomyelin, C18 sphingomyelin, C20 sphingomyelin, C22 sphingomyelin and C24:1 sphingomyelin content in the contralateral non-ischemic control striatum (C) and reperfused ischemic striatum (I/R) measured by liquid chromatography tandem-mass spectrometry (LC-MS/MS) in C57BL/6j mice exposed to transient MCAO, which were intraperitoneally treated with vehicle or amitriptyline (2 or 12 mg/kg b.w., b.i.d.) starting 24 hours post-MCAO, followed by animal sacrifice after 14 days. Data are means ± SD values. *p≤0.05/**p≤0.01 compared with non-ischemic C (n=7-9 animals/group; analyzed by one-way ANOVA, followed by LSD tests).

**Supplementary Figure 3. Asm inhibitor amitriptyline reduces ischemic injury *in vivo*, when administered immediately after I/R. (A)** Infarct volume and **(B)** brain edema evaluated on cresyl violet-stained brain sections, and **(C)** serum IgG extravasation as a marker of blood-brain barrier permeability by immunohistochemistry in the reperfused ischemic striatum of mice exposed to transient MCAO, which were intraperitoneally treated with vehicle or amitriptyline (2 or 12 mg/kg b.w., b.i.d.) immediately after MCAO, followed by animal sacrifice after 24 hours. Representative brain sections are also shown. Data are means ± SD values. *p≤0.05/**p≤0.01 compared with corresponding vehicle (n=7-8 animals/group; analyzed by one-way ANOVA, followed by LSD tests). Scale bar, 1000 µm.

**Supplementary Figure 4. I/R induces the intracellular formation of ceramide-rich vesicles in cerebral microvascular endothelial cells *in vitro*. (A)** Number and (**B**) size of intracellular ceramide-rich vesicles evaluated by Cell Profiler in hCMEC/D3 exposed to non-ischemic control condition (C), 24 hours OGD (that is, ischemia; I) or 24 hours ischemia followed by different durations of reoxygenation/glucose re-supplementation (I/R). Data are means ± SD values. ^†^p≤0.05/^†††^p≤0.001 compared with corresponding C; ^‡‡‡^p≤0.001 compared with corresponding I (n=3-8 independent samples/group [in **(A)**]; n=3-4 independent samples/group [in **(B)**]; analyzed by one-way ANOVA, followed by LSD tests).

**Supplementary Figure 5. ASM is the predominant sphingomyelinase in endothelial cells. (A)** *SMPD1*, *SMPD2* and *SMPD3* mRNA level evaluated by polymerase-chain reaction (PCR) in hCMEC/D3, primary human brain microvascular endothelial cells (HBMEC), human umbilical vein endothelial cells (HUVEC) and human peripheral blood mononuclear cells (hPBMC) and **(B)** magnesium dependent NSM activity examined using BODIPY-labeled sphingomyelin as substrate in whole brain tissue, hCMEC/D3, HBMEC and HUVEC. Data are means ± SD values (n=3 independent samples/group).

**Supplementary Figure 6. ASM inhibitors do not influence the survival of human cerebral microvascular endothelial cells *in vitro*.** Endothelial viability assessed by the 3-(4,5-dimethylthiazol-2-yl)-2,5-diphenyltetrazolium bromide (MTT) assay in non-ischemic hCMEC/D3 treated with **(A)** vehicle or amitriptyline at defined doses (0-50 µM), **(B)** vehicle or fluoxetine at defined doses (0-20 µM) or **(C)** vehicle or desipramine at defined doses (0-50 µM), or in **(D)** in non-ischemic hCMEC/D3 transfected with scrambled siRNA or *SMPD1* siRNA that were treated with vehicle or amitriptyline (50 µM). Data are means ± SD values. *p≤0.05 compared with scrambled siRNA/ vehicle (n=3 independent samples/group [in **(A, B)**]; n=4 independent samples/group [in **(C, D)**]; analyzed by one-way ANOVA, followed by LSD tests).

**Supplementary Figure 7. ASM inhibitor amitriptyline increases VEGFR2 abundance and VEGF secretion by mouse cerebral microvascular endothelial cells. (A)** VEGFR2 abundance evaluated by Western blot and **(B)** VEGF concentration in supernatant evaluated by enzyme-linked immunosorbent assay (ELISA) of non-ischemic mouse brain endothelial cells belonging to the cell line bEND5, which were treated with vehicle or amitriptyline for defined exposure times (0-24 hours) at a 50 µM concentration. Data are means ± SD values. **p≤0.01/***p≤0.001 compared with corresponding vehicle (n=4 independent samples/group [in **(A-B)**]; analyzed by one-way ANOVA, followed by LSD tests).

**Supplementary Figure 8. *SMPD1* mRNA level, ASM abundance and ASM activity are effectively downregulated by small-interfering RNA (siRNA) *in vitro*. (A)** *SMPD1* mRNA expression, evaluated by polymerase-chain reaction (PCR), (B) ASM abundance, determined by Western blot and (C) ASM activity, examined using BODIPY-labeled sphingomyelin as substrate, in hCMEC/D3 transfected with scrambled siRNA and *SMPD1* siRNA for 24-72 hours. Data are means ± SD values. *p≤0.05/***p≤0.001 compared with scrambled siRNA (n=3 independent samples/ group [in (A)]; n=4 independent samples/ group [in (B)]; n=3-6 independent samples/ group [in (C)]; analyzed by two-tailed t tests).

**Supplementary Figure 9. *SMPD1* knockdown reduces the intracellular accumulation of ceramide-rich vesicles after I/R *in vitro* and promotes the extracellular release of vesicles with immunofluorescence exosome characteristics. (A)** Immunohistochemistry for ceramide (in magenta) in hCMEC/D3 exposed to 24 hours OGD (that is, ischemia) followed by 3 hours reoxygenation/glucose re-supplementation, which were transfected with scrambled siRNA (used as control) or *SMPD1* siRNA. Nuclei were counterstained with DAPI (in blue). Note that the number of intracellular ceramide-rich vesicles is reduced by *SMPD1* knockdown. Shown is a representative result from 3 independent studies. The density of vesicles evaluated by Cell Profiler is shown in **(B)**. Particle concentration of **(C, E)** CD9^+^ and **(D, F)** CD63^+^ sEVs in the supernatant of hCMEC/D3 exposed to 24 hours ischemia followed by 3 hours reperfusion (in **(C, D)**) or 24 hours reperfusion (in **(E, F)**), which were transfected with scrambled siRNA (used as control) or *SMPD1* siRNA. The particle concentration was evaluated by AMNIS image flow cytometry. Note that the number of CD9^+^ and CD63^+^ sEVs is increased by *SMPD1* knockdown. Data are means ± SD values. *p≤0.05/**p≤0.01 compared with corresponding scrambled siRNA (n=3 independent samples/ group [in **(B)**]; n=4 independent samples/ group [in **(C-F)**]; analyzed by two-tailed t tests).

**Supplementary Figure 10. *SMPD1* knockdown does not change overall ceramide levels in cerebral microvascular endothelial cells *in vitro*. (A)** Total ceramide, **(B)** C16 ceramide, **(C)** C18 ceramide, **(D)** C20 ceramide, **(E)** C22 ceramide and **(F)** C24:1 ceramide content of hCMEC/D3 exposed to non-ischemic control condition (C), OGD (that is, ischemia; I) or ischemia followed by reoxygenation/glucose re-supplementation (I/R), which were transfected with scrambled siRNA (used as control) or *SMPD1* siRNA. Ceramide levels were measured by LC-MS/MS. Data are means ± SD values. ^†^p≤0.05 compared with corresponding C; ^‡^p≤0.05/^‡‡^p≤0.001 compared with corresponding I (n=3 independent samples/ group [in **(A-F)**]; analyzed by two-way ANOVA, followed by LSD tests).

**Supplementary Figure 11. *SMPD1* knockdown increases sphingomyelin levels in cerebral microvascular endothelial cells *in vitro*. (A)** Total sphingomyelin, **(B)** C16 sphingomyelin, **(C)** C18 sphingomyelin, **(D)** C20 sphingomyelin, **(E)** C22 sphingomyelin and **(F)** C24:1 sphingomyelin content of hCMEC/D3 exposed to non-ischemic control condition (C), OGD (that is, ischemia; I) or ischemia followed by reoxygenation/glucose re-supplementation (I/R), which were transfected with scrambled siRNA (used as control) or *SMPD1* siRNA. Sphingomyelin levels were measured by LC-MS/MS. Data are means ± SD values. *p≤0.05/**p≤0.01 compared with corresponding scrambled siRNA; ^†^p≤0.05/^††^p≤0.01 compared with corresponding C (n=3 independent samples/ group [in **(A-F)**]; analyzed by two-way ANOVA, followed by LSD tests).

**Supplementary Figure 12. Amitriptyline promotes cerebral angiogenesis *in vitro* in an ASM dependent way. (A)** Microvascular length and **(B)** branching point density evaluated in the matrigel-based tube formation assay of non-ischemic hCMEC/D3 transfected with scrambled siRNA (used as control) or *SMPD1* siRNA which were exposed to vehicle or amitriptyline (50 µM). Representative photographs for tube formation assays are shown in Figure 4. Data are means ± SD values. *p≤0.05/***p≤0.001 compared with corresponding vehicle; ^‡^p≤0.05/^‡‡‡^p≤0.001 compared with corresponding scrambled siRNA (n=5-8 independent samples/group; analyzed by two-way ANOVA, followed by LSD tests).

**Supplementary Figure 13. Intracellular ceramide-rich vesicles do not express mitochondrial, early endosome, lysosome, autophagosome and caveolae markers.** Immunocytochemistry for ceramide (in green) and **(A)** the mitochondrial marker apoptosis-inducing factor (AIF), **(B)** the early endosome marker early endosome antigen-1 (EEA1), **(C)** the lysosome marker lysosomal-associated membrane protein-1 (LAMP1), **(D)** the autophagosome marker microtubule-associated protein light chain-3b (LC3b) and **(E)** the caveolae marker caveolin (all in magenta) of hCMEC/D3 exposed to 24 hours OGD (that is, ischemia) followed by 3 hours reoxygenation/glucose re-supplementation (I/R). In the merged photomicrographs, no double labeled cells are visible. Nuclei were counterstained with DAPI (in blue). Scale bar, 5 µm. Shown are representative results from 4 independent studies.

**Supplementary Figure 14. Gating strategy for evaluating sEVs by ImageStreamX flow cytometry.** From all recorded signals (1^st^ plot from left), signals not showing spot count signal or signal multiplets were excluded (2^nd^ plot from left). In the two representative plots on the right, side scatter (SSC) intensities of single objects are plotted against the fluorescence intensities of CD9^+^ (labeled with FITC) or CD63^+^ (labeled with APC) objects. Shown are representative results under control conditions from 4-9 independent studies.

**Supplementary Figure 15. sEVs released from cerebral microvascular endothelial cells have the physicochemical properties and protein expression characteristics of exosomes. (A)** Particle size evaluated by nanoparticle tracking analysis (NTA) of sEVs obtained from supernatants of hCMEC/D3 exposed to non-ischemic control condition (C), OGD (that is, ischemia; I) or ischemia followed by reoxygenation/glucose re-supplementation (I/R) which were treated with vehicle or amitriptyline (Ami, 50 µM). Western blots for **(B, C)** the exosomal markers synthenin and CD9 and **(D)** the cellular contamination marker calnexin using protein samples obtained from (1) hCMEC/D3 lysates, (2) sEVs from vehicle-treated non-ischemic control hCMEC/D3, (3) sEVs from amitriptyline-treated control hCMEC/D3, (4) sEVs from vehicle-treated ischemic hCMEC/D3, (5) sEVs from amitriptyline-treated ischemic hCMEC/D3, (6) sEVs from vehicle-treated I/R hCMEC/D3 and (7) sEVs from amitriptyline-treated I/R hCMEC/D3. Note the presence of 32 kDa synthenin and 25 kDa CD9, but absence of 90 kDa/ 67 kDa calnexin in sEVs samples. Data are means ± SD values. No significant differences were noted between groups (n=3 independent samples/group [in **(A)**]; analyzed by one-way ANOVA).

**Supplementary Figure 16. sEVs released from cerebral microvascular endothelial cells have the electron microscopic size and appearance of exosomes.** Transmission electron microscopy showing representative sEVs obtained from supernatants of hCMEC/D3 exposed to non-ischemic control condition (C), OGD (that is, ischemia; I) or ischemia followed by reoxygenation/glucose re-supplementation (I/R), which had been treated with vehicle or amitriptyline (50 µM), while cells were cultured. The typical cup shape double contrast is suggestive for a double-layered membrane structure that is found in sEVs. Scale bar, 100 nm.

**Supplementary Figure 17. sEVs obtained from supernatants of cerebral microvascular endothelial cells exposed to ASM inhibitor amitriptyline have angiogenic activity that resembles sEVs released by endothelial cells during I/R.** Representative photographs of **(A)** matrigel-based tube formation and **(B)** transwell migration assays of hCMEC/D3, which were treated with vehicle, amitriptyline (Ami, 50 µM) or sEVs (25 µg protein/ ml) isolated from the supernatants of hCMEC/D3 that had been cultured in non-ischemic control condition (C), OGD (that is, ischemia; I) or ischemia followed by reoxygenation/glucose re-supplementation (I/R) and had been treated with vehicle or amitriptyline (50 µM) while supernatants were collected. A quantitative analysis is shown in Figure 4. Scale bar, 200 µm in **(A)**; 20 µm in **(B)**.

**Supplementary Figure 18.** Volcano plot showing differentially regulated proteins in sEVs released from hCMEC/D3 cultured under non-ischemic control conditions that had been exposed to vehicle or amitriptyline (50 µM). Proteins were evaluated by LC-MS-based proteomics. Proteins with statistically significant differential regulation (≥2-fold, p value ≤0.05; analyzed by two-tailed t tests) are located in the top left and right sectors (n=6 independent samples/group).

