## Supplementary Figures 1 - 18 for "Acid sphingomyelinase deactivation post-ischemia/ reperfusion promotes cerebral angiogenesis and brain remodeling via small extracellular vesicles"

**Supplementary Figure 1: Acid sphingomyelinase (Asm) product ceramide accumulates in cerebral microvessels after ischemia/ reperfusion (I/R) *in vivo*.**

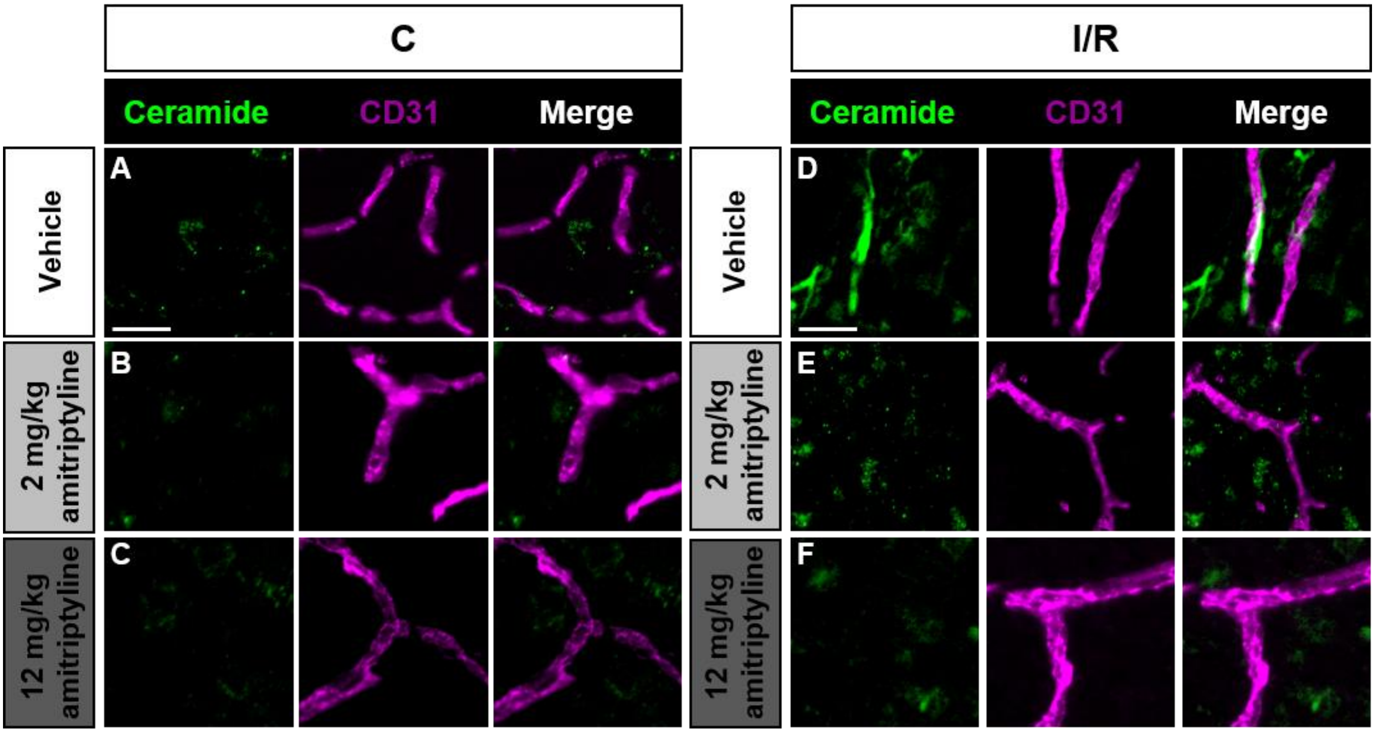

**Supplementary Figure 2: Asm inhibitor amitriptyline does not change cerebral sphingomyelin levels after I/R *in vivo*.**

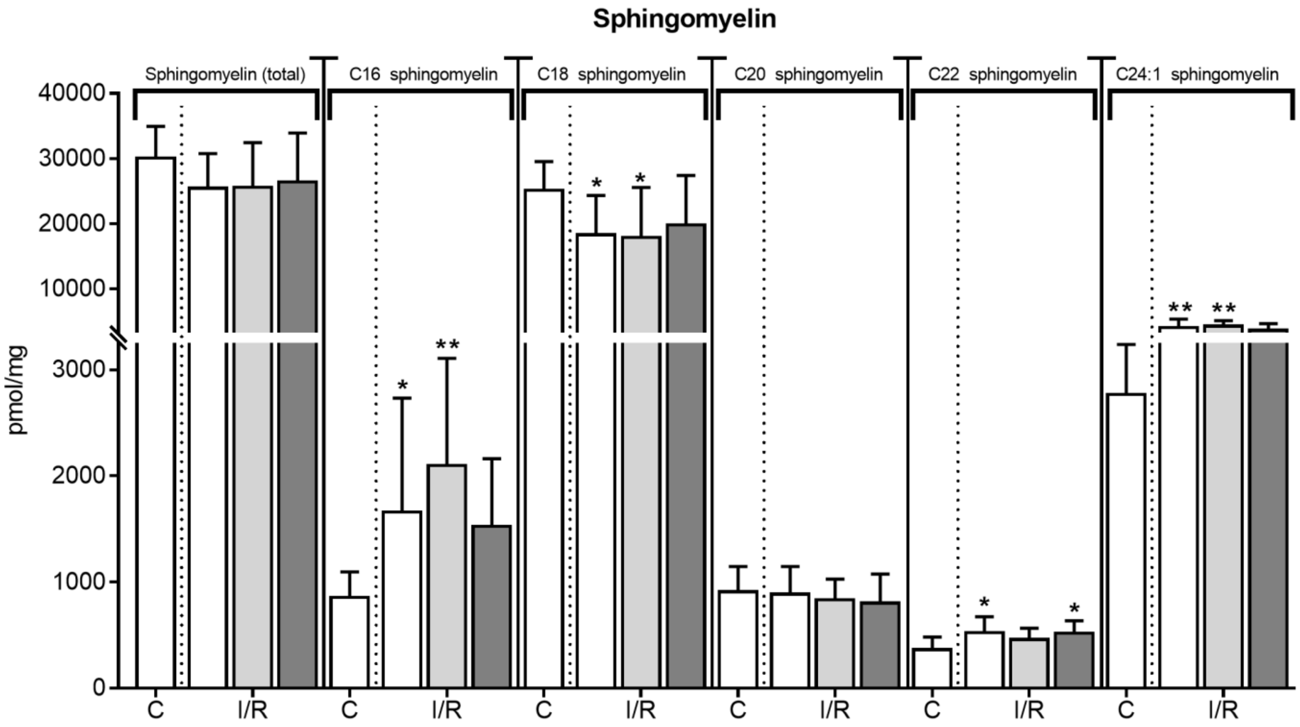

**Supplementary Figure 3: Asm inhibitor amitriptyline reduces ischemic injury *in vivo*, when administered immediately after I/R.**

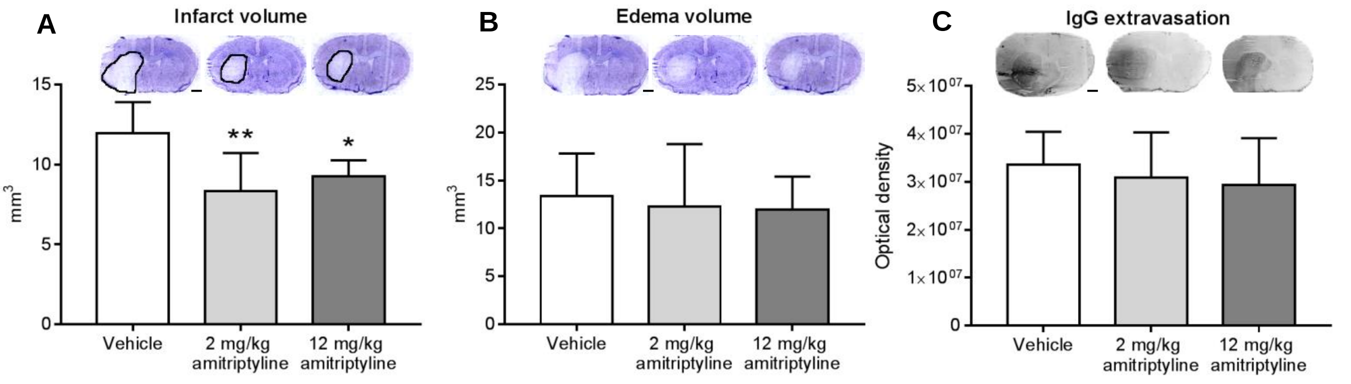

**Supplementary Figure 4: I/R induces the intracellular formation of ceramide-rich vesicles in cerebral microvascular endothelial cells *in vitro*.**

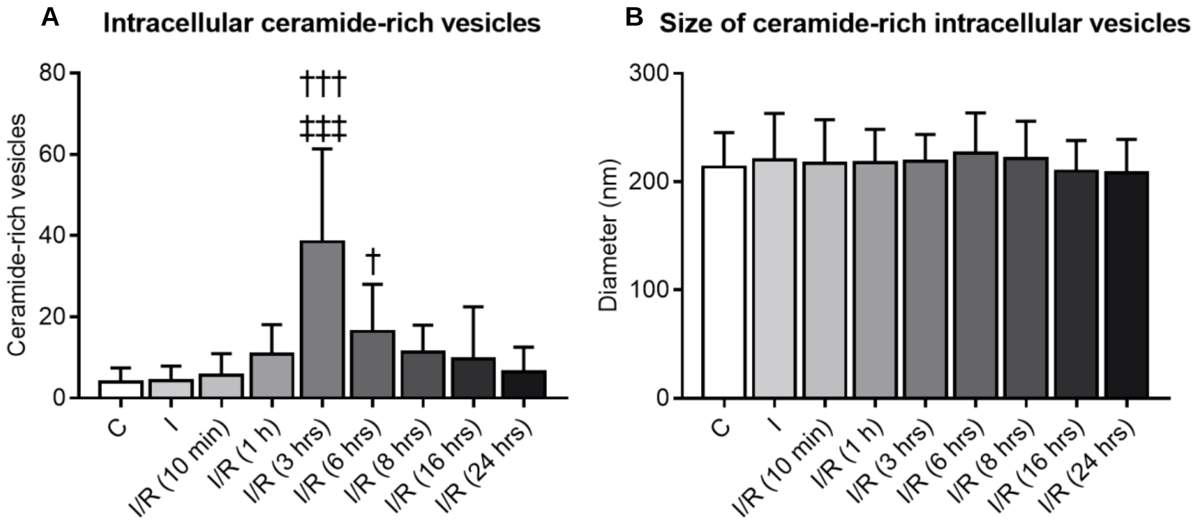

**Supplementary Figure 5: ASM is the predominant sphingomyelinase in endothelial cells.**

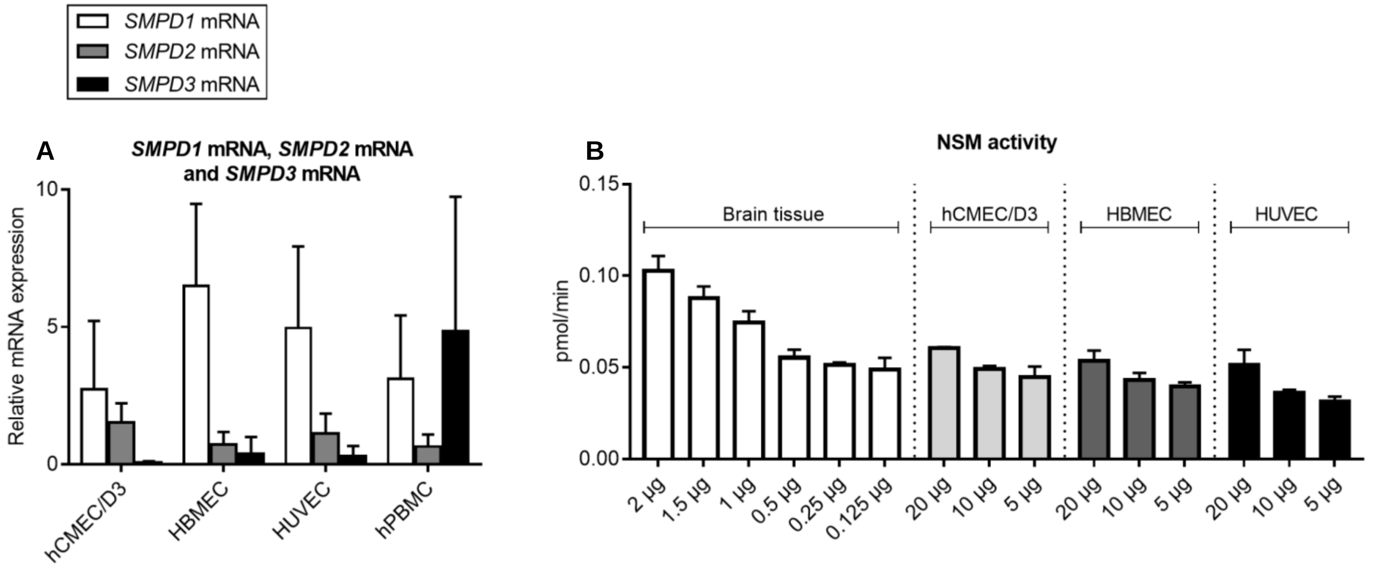

**Supplementary Figure 6: ASM inhibitors do not influence the survival of human cerebral microvascular endothelial cells *in vitro*.**

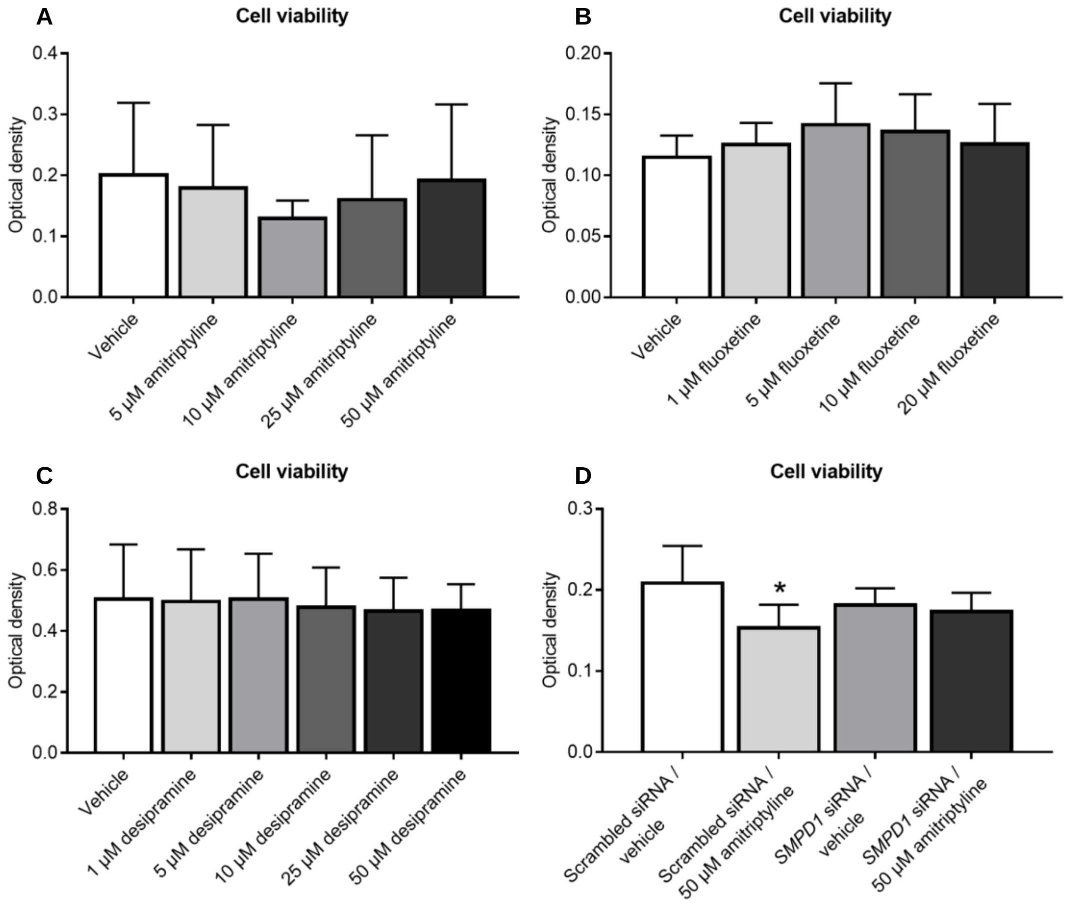

**Supplementary Figure 7: ASM inhibitor amitriptyline increases VEGFR2 abundance and VEGF secretion by mouse cerebral microvascular endothelial cells.**

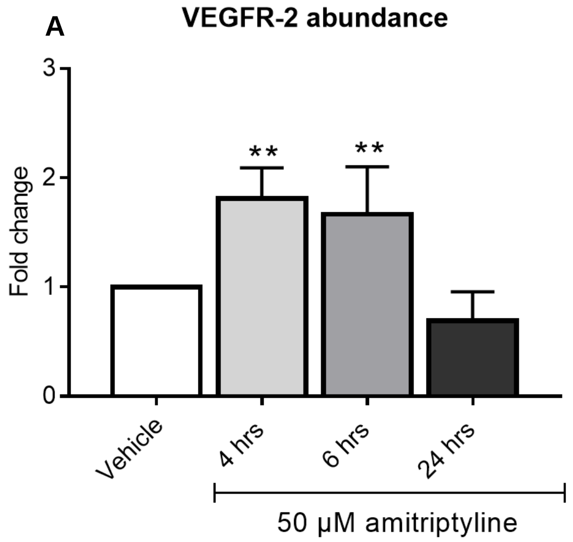

**Supplementary Figure 8: *SMPD1* mRNA level, ASM abundance and ASM activity are effectively downregulated by small-interfering RNA (siRNA) *in vitro*.**

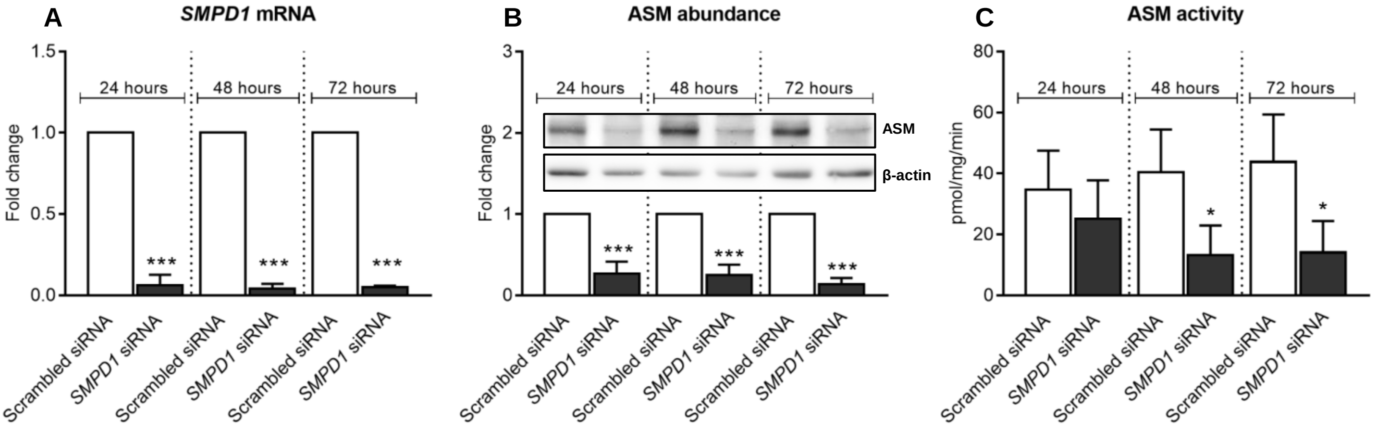

**Supplementary Figure 9: *SMPD1* knockdown reduces the intracellular accumulation of ceramide-rich vesicles after I/R *in vitro* and promotes the extracellular release of vesicles with immunofluorescence exosome characteristics.**

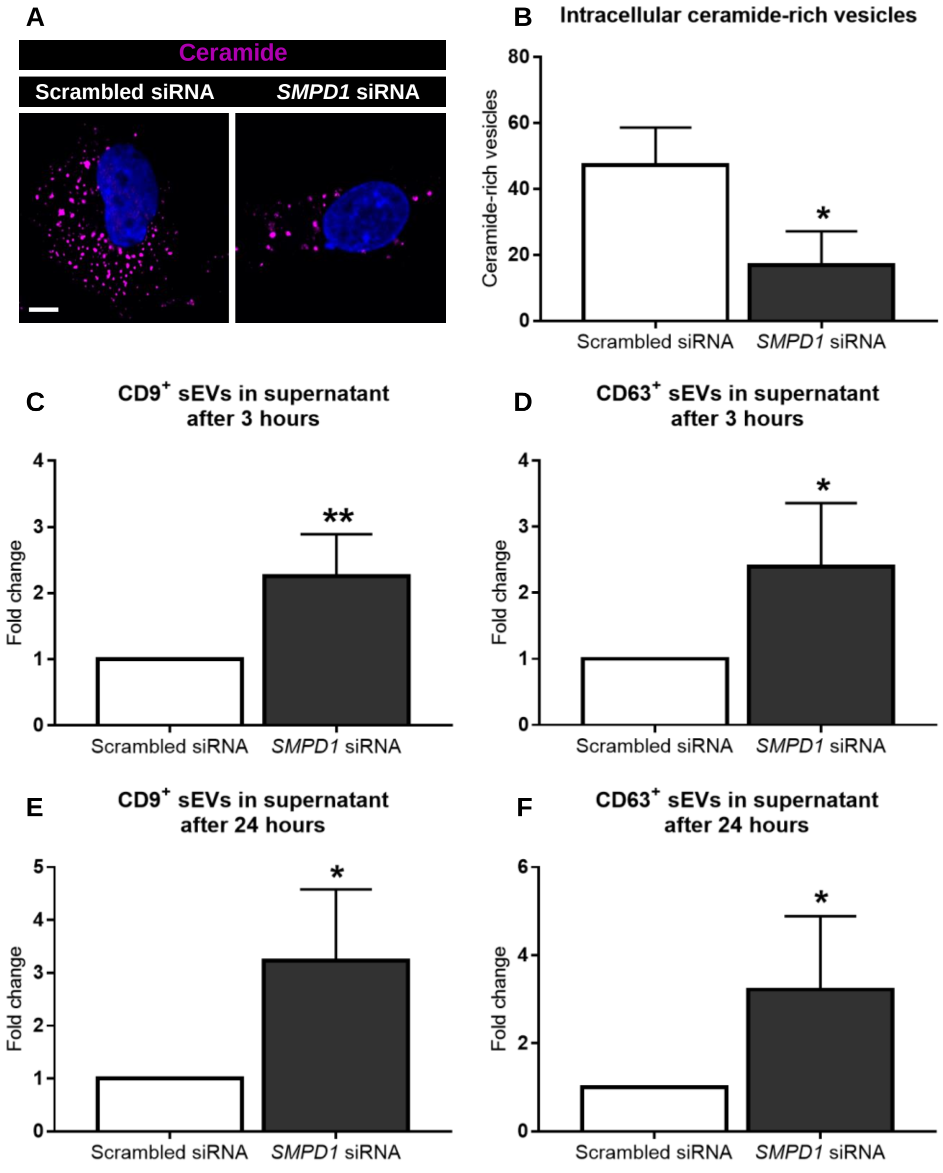

**Supplementary Figure 10: *SMPD1* knockdown does not change overall ceramide levels in cerebral microvascular endothelial cells *in vitro*.**

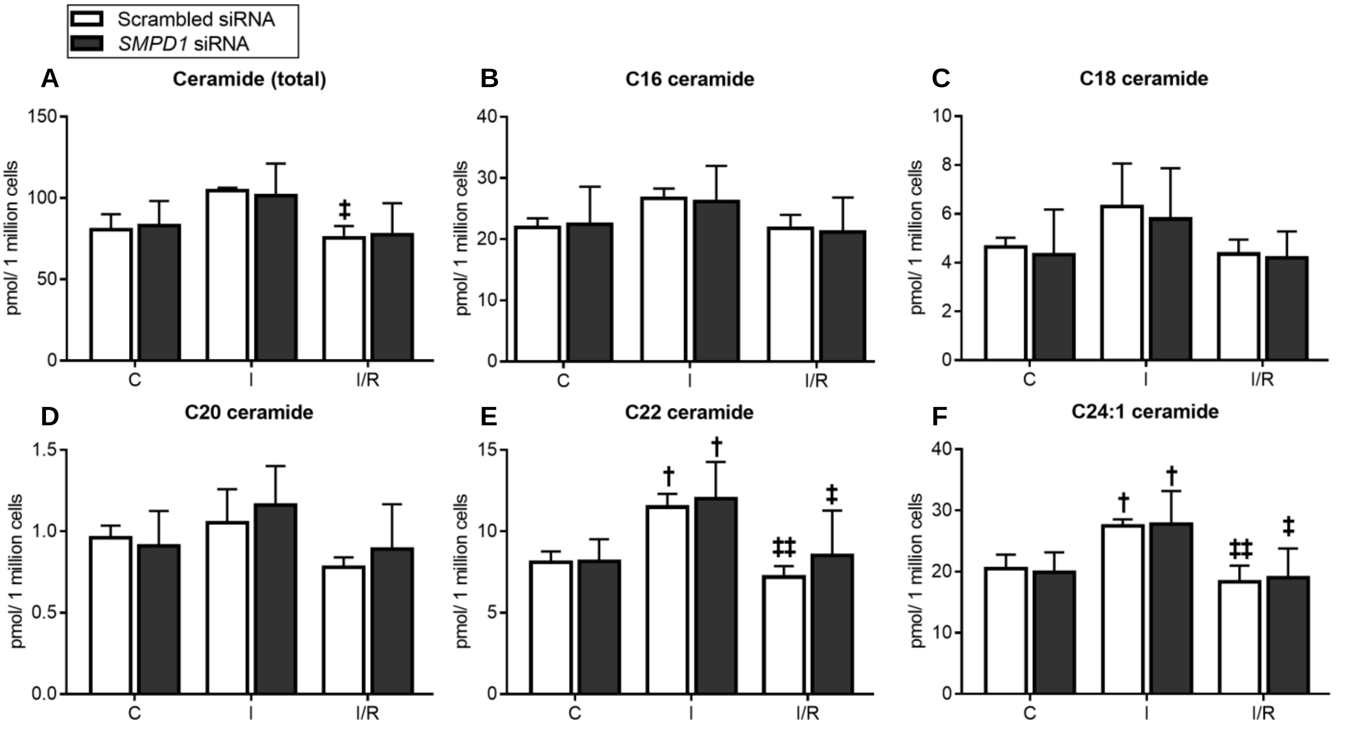

**Supplementary Figure 11: *SMPD1* knockdown increases sphingomyelin levels in cerebral microvascular endothelial cells *in vitro*.**

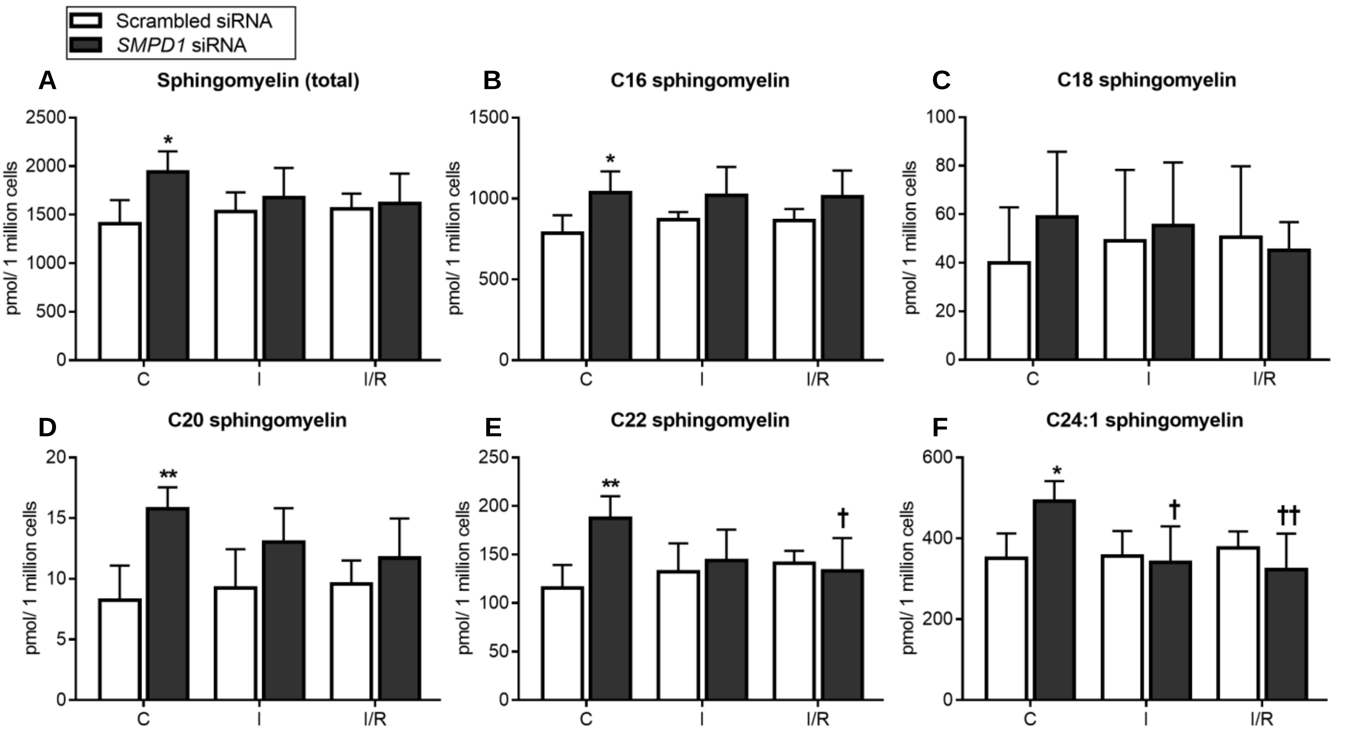

**Supplementary Figure 12: Amitriptyline promotes cerebral angiogenesis *in vitro* in an ASM dependent way.**

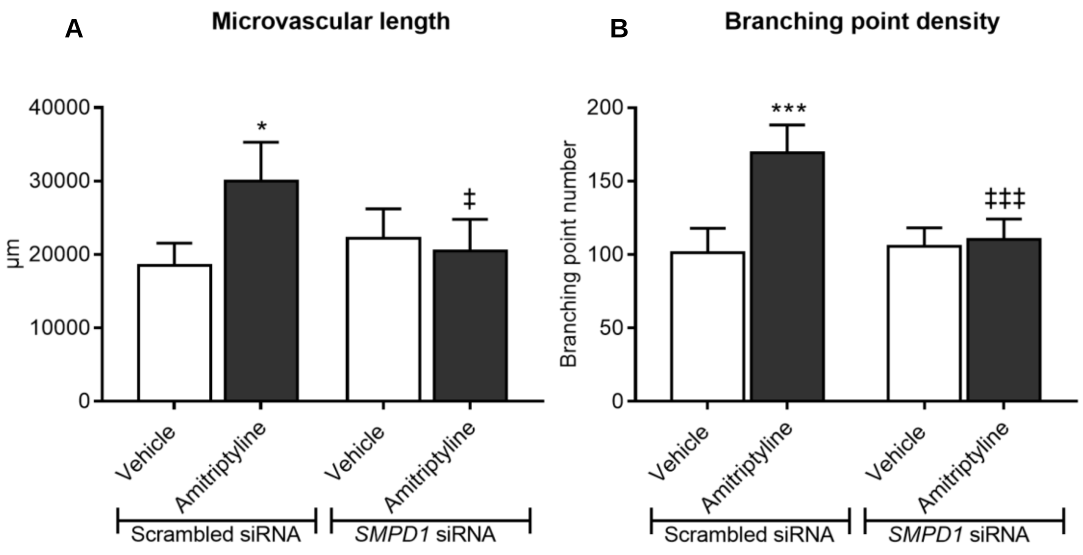

**Supplementary Figure 13: Intracellular ceramide-rich vesicles do not express mitochondrial, early endosome, lysosome, autophagosome and caveolae markers.**

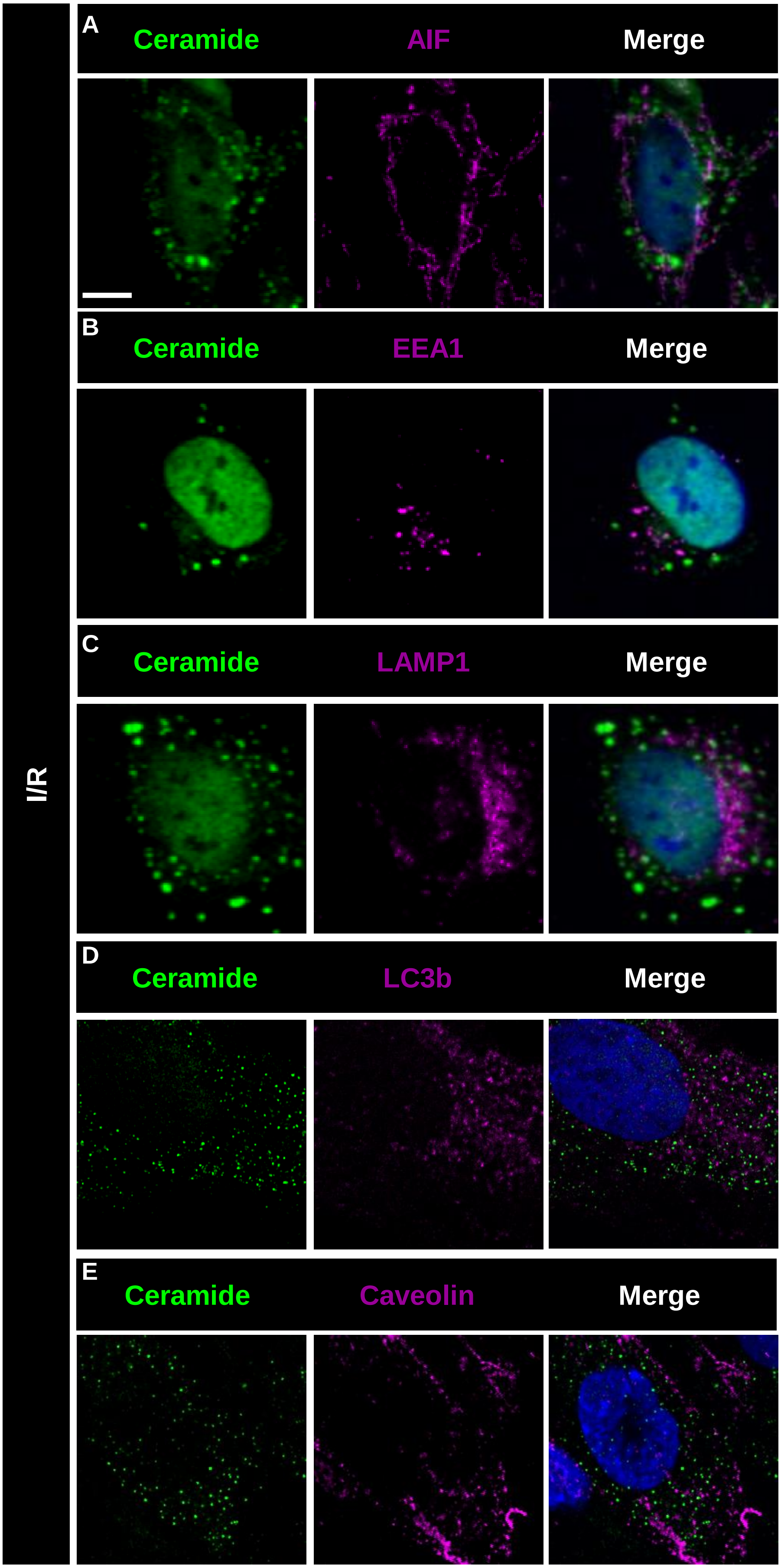

**Supplementary Figure 14: Gating strategy for evaluating sEVs by ImageStreamX flow cytometry.**

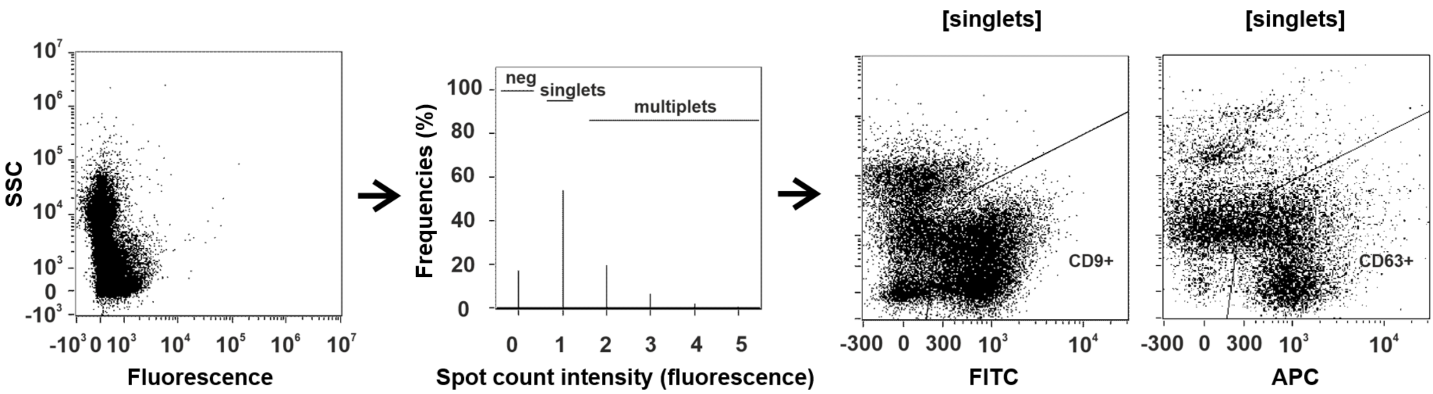

**Supplementary Figure 15: sEVs released from cerebral microvascular endothelial cells have the physicochemical properties and protein expression characteristics of exosomes.**

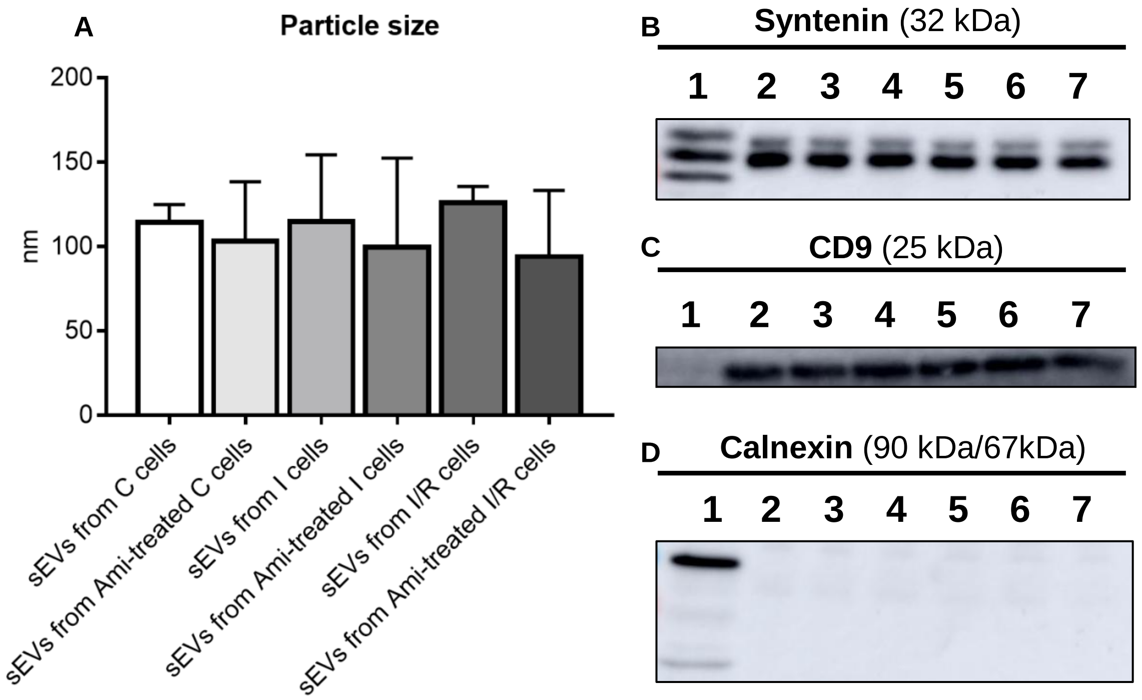

**Supplementary Figure 16: sEVs released from cerebral microvascular endothelial cells have the electron microscopic size and appearance of exosomes.**

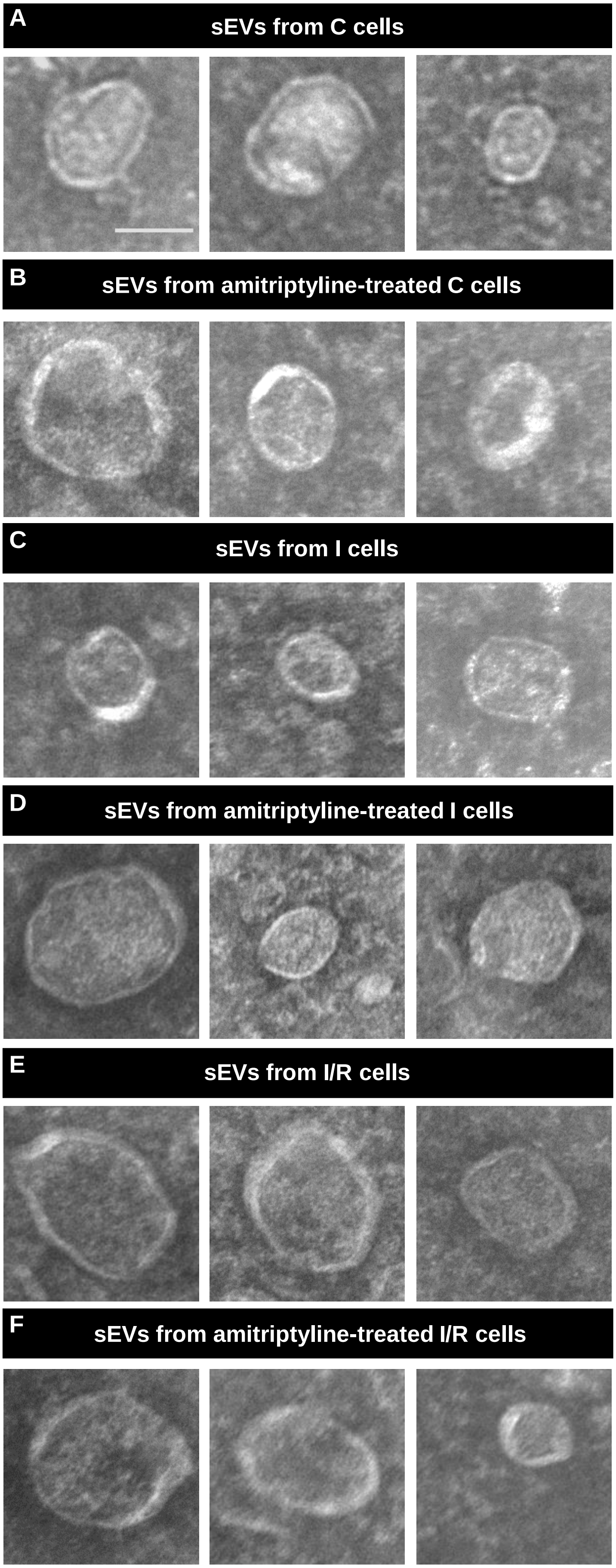

**Supplementary Figure 17: sEVs obtained from supernatants of cerebral microvascular endothelial cells exposed to ASM inhibitor amitriptyline have angiogenic activity that resembles sEVs released by endothelial cells during I/R.**

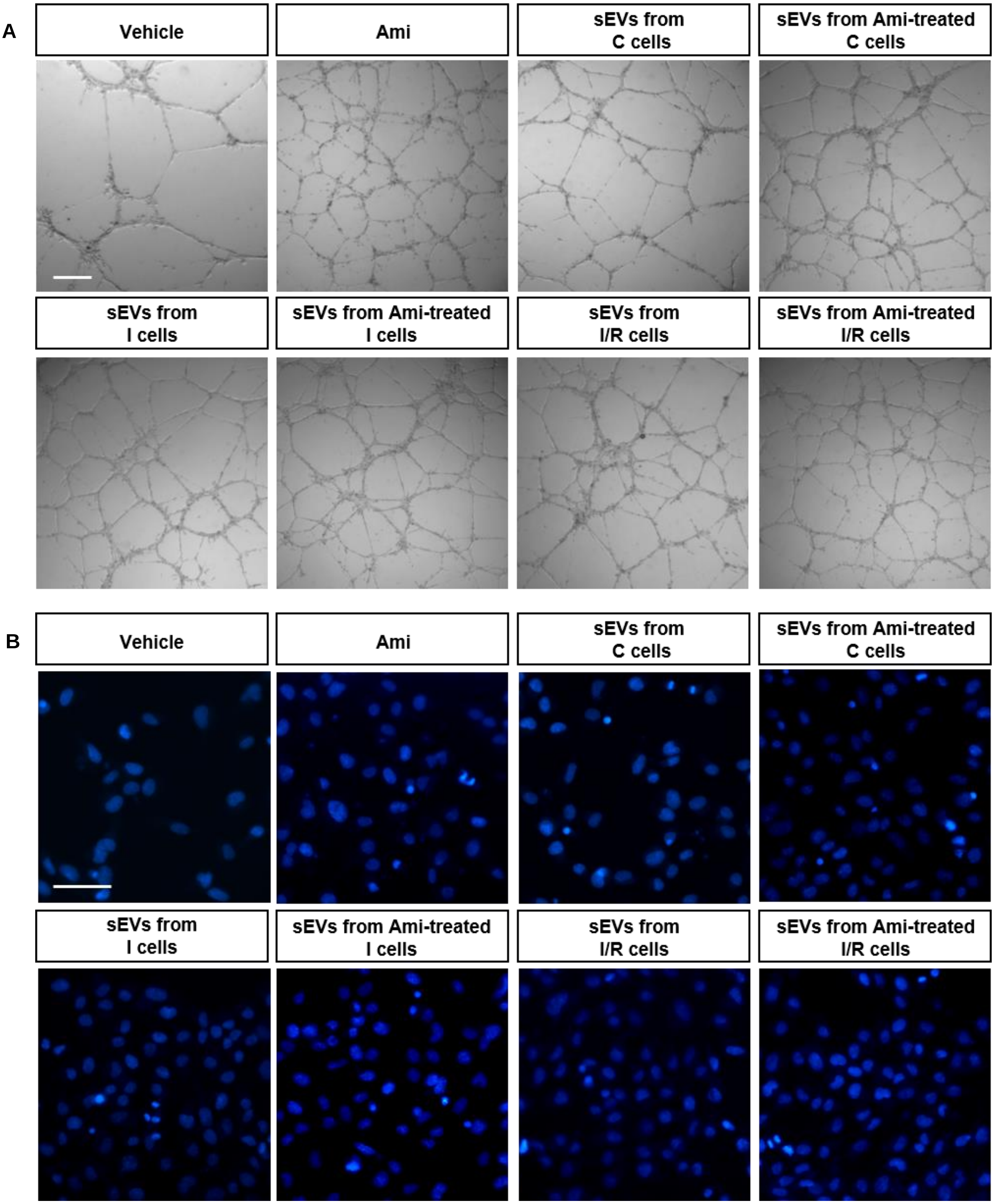

**Supplementary Figure 18: Volcano plot showing differentially regulated proteins in sEVs released from hCMEC/D3 exposed to vehicle or amitriptyline.**

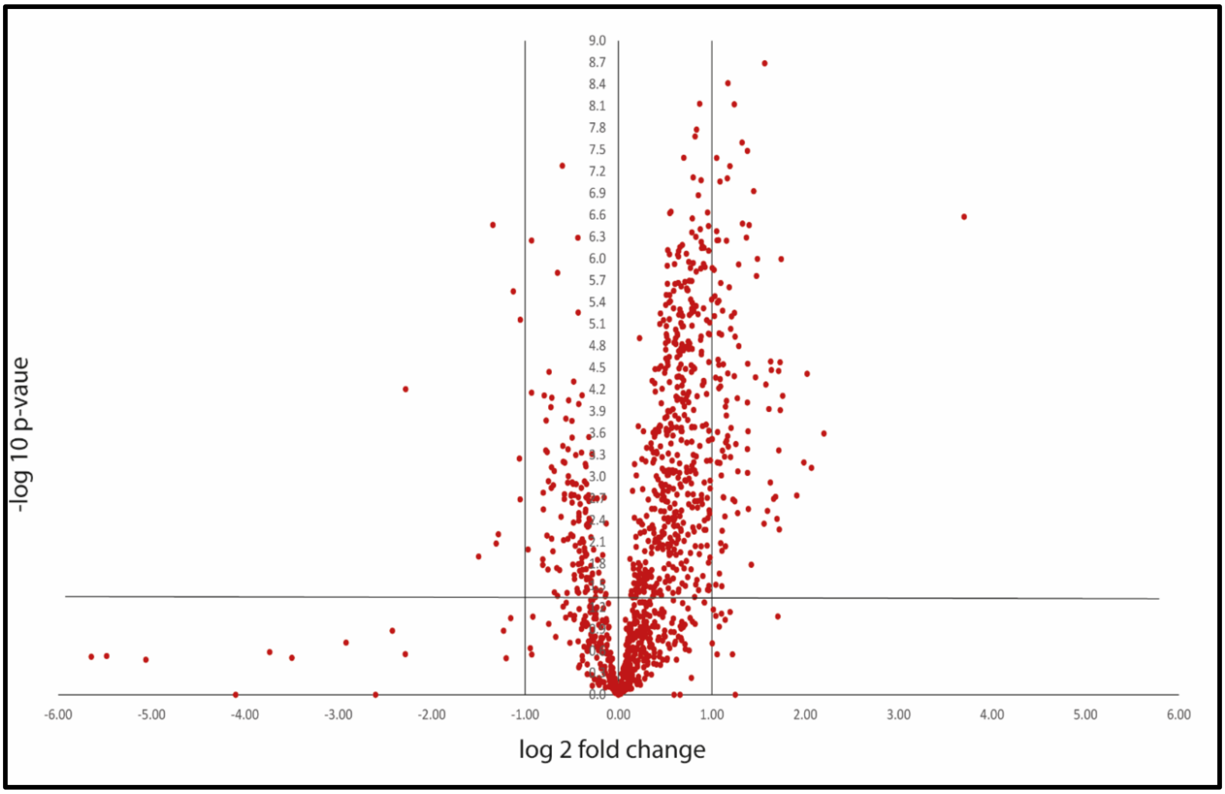
