## Supplementary Tables 1 - 4 for "Acid sphingomyelinase deactivation post-ischemia/ reperfusion promotes cerebral angiogenesis and brain remodeling via small extracellular vesicles"

### Supplementary Table 1. Proteins significantly upregulated in sEVs released by hCMEC/D3 exposed to amitriptyline.

| Association according to KEGG pathway database: |
|---|
| Phagosome |
| Lysosome |
| Protein export |
| Antigen processing and presentation |
| Epstein-Barr virus infection |
| Viral myocarditis |

| Accession | Description | Protein name | Log 2 fold change | P-value |
|---|---|---|---|---|
| Q8NE71 | ATP-binding cassette sub-family F member 1 | ABCF1 | 1.12 | 0.00 |
| P11310 | Medium-chain specific acyl-CoA dehydrogenase, mitochondrial | ACADM | 1.73 | 0.00 |
| Q13443 | Disintegrin and metalloproteinase domain-containing protein 9 | ADAM9 | 1.38 | 0.00 |
| Q13155 | Aminoacyl tRNA synthase complex-interacting multifunctional protein 2 | AIMP2 | 1.09 | 0.00 |
| Q13740 | CD166 antigen | ALCAM | 1.20 | 0.00 |
| P09972 | Fructose-bisphosphate aldolase C | ALDOC | 1.04 | 0.03 |
| P15144 | Aminopeptidase N | AN | 1.10 | 0.00 |
| Q9BXS5 | AP-1 complex subunit mu-1 | AP1M1 | 1.06 | 0.00 |
| Q92572 | AP-3 complex subunit sigma-1 | AP3S1 | 1.14 | 0.00 |
| P05026 | Sodium/potassium-transporting ATPase subunit beta-1 | ATP1B1 | 1.00 | 0.00 |
| P61769 | Beta-2-microglobulin | B2M | 1.21 | 0.00 |
| Q05682 | Caldesmon | CALD1 | 1.14 | 0.00 |
| P27824 | Calnexin | CANX | 1.28 | 0.00 |
| Q6NZI2 | Caveolae-associated protein 1 | CAVIN1 | 1.63 | 0.00 |
| P45973 | Chromobox protein homolog 5 | CBX5 | 1.33 | 0.00 |
| P16070 | CD44 antigen | CD44 | 1.09 | 0.00 |
| P15529 | Membrane cofactor protein | CD46 | 1.28 | 0.00 |
| P08174 | Complement decay-accelerating factor | CD55 | 1.38 | 0.00 |
| P48960 | CD97 antigen | CD97 | 1.17 | 0.00 |
| Q14008 | Cytoskeleton-associated protein 5 | CKAP5 | 1.11 | 0.00 |
| Q6UVK1 | Chondroitin sulfate proteoglycan 4 | CSPG4 | 1.40 | 0.00 |
| P07858 | Cathepsin B | CTSB | 1.06 | 0.00 |
| P07339 | Cathepsin D | CTSD | 1.07 | 0.00 |
| Q14247 | Src substrate cortactin | CTTN | 1.42 | 0.02 |
| P17844 | Probable ATP-dependent RNA helicase DDX5 | DDX5 | 1.03 | 0.00 |
| P31689 | DnaJ homolog subfamily A member 1 | DNAJA1 | 1.09 | 0.00 |
| P23919 | Thymidylate kinase | DTYMK | 1.56 | 0.00 |
| Q9BY44 | Eukaryotic translation initiation factor 2A | EIF2A | 1.23 | 0.00 |
| P19525 | Interferon-induced, double-stranded RNA-activated protein kinase | EIF2AK2 | 1.74 | 0.00 |
| O75821 | Eukaryotic translation initiation factor 3 subunit G | EIF3G | 1.11 | 0.01 |
| Q7L2H7 | Eukaryotic translation initiation factor 3 subunit M | EIF3M | 1.66 | 0.00 |
| Q04637 | Eukaryotic translation initiation factor 4 gamma 1 | EIF4G1 | 1.05 | 0.00 |
| P13804 | Electron transfer flavoprotein subunit alpha, mitochondrial | ETFA | 1.16 | 0.00 |
| Q96A65 | Exocyst complex component 4 | EXOC4 | 2.02 | 0.00 |
| Q13451 | Peptidyl-prolyl cis-trans isomerase FKBP5 | FKBP5 | 1.09 | 0.00 |
| P35637 | RNA-binding protein FUS | FUS | 1.47 | 0.00 |
| P51114 | Fragile X mental retardation syndrome-related protein 1 | FXR1 | 1.08 | 0.00 |
| P60520 | Gamma-aminobutyric acid receptor-associated protein-like 2 | GABARAPL2 | 1.58 | 0.00 |
| Q96RP9 | Elongation factor G, mitochondrial | GFM1 | 1.06 | 0.00 |
| P16278 | Beta-galactosidase | GLB1 | 1.68 | 0.00 |
| Q14344 | Guanine nucleotide-binding protein subunit alpha-13 | GNA13 | 2.20 | 0.00 |
| P08236 | Beta-glucuronidase | GUSB | 1.98 | 0.00 |
| P19367 | Hexokinase-1 | HK1 | 1.64 | 0.00 |
| P01889 | HLA class I histocompatibility antigen, B alpha chain | HLA-B | 1.24 | 0.00 |
| P10321 | HLA class I histocompatibility antigen, C alpha chain | HLA-C | 1.25 | 0.00 |
| P51991 | Heterogeneous nuclear ribonucleoprotein A3 | HNRNPA3 | 1.63 | 0.00 |
| P31943 | Heterogeneous nuclear ribonucleoprotein H | HNRNPH1 | 1.08 | 0.00 |
| P14866 | Heterogeneous nuclear ribonucleoprotein L | HNRNPL | 1.00 | 0.00 |
| Q00839 | Heterogeneous nuclear ribonucleoprotein U | HNRNPU | 1.07 | 0.00 |
| Q16666 | Gamma-interferon-inducible protein 16 | IFI16 | 1.38 | 0.00 |
| Q9Y6M1 | Insulin-like growth factor 2 mRNA-binding protein 2 | IGF2BP2 | 1.29 | 0.00 |
| P08648 | Integrin alpha-5 | ITGA5 | 1.38 | 0.00 |
| P33176 | Kinesin-1 heavy chain | KIF5B | 1.19 | 0.00 |
| P52292 | Importin subunit alpha-1 | KPNA2 | 1.39 | 0.00 |
| P11279 | Lysosome-associated membrane glycoprotein 1 | LAMP1 | 1.24 | 0.00 |
| Q9UHB6 | LIM domain and actin-binding protein 1 | LIMA1 | 1.38 | 0.00 |
| Q8N1G4 | Leucine-rich repeat-containing protein 47 | LRRC47 | 1.11 | 0.01 |
| P27816 | Microtubule-associated protein 4 | MAP4 | 1.32 | 0.00 |
| P43243 | Matrin-3 | MATR3 | 1.73 | 0.00 |
| P43121 | Cell surface glycoprotein MUC18 | MCAM | 1.20 | 0.00 |
| Q14566 | DNA replication licensing factor MCM6 | MCM6 | 1.26 | 0.00 |
| O95297 | Myelin protein zero-like protein 1 | MPZL1 | 1.15 | 0.00 |
| P42285 | Exosome RNA helicase MTR4 | MTREX | 1.61 | 0.00 |
| P20591 | Interferon-induced GTP-binding protein Mx1 | MX1 | 1.37 | 0.00 |
| Q92542 | Nicastrin | NCSTN | 1.72 | 0.00 |
| P09874 | Poly [ADP-ribose] polymerase 1 | PARP1 | 1.06 | 0.00 |

Comparison of n=6 different samples/group obtained from 10 independent cell cultures.

**Supplementary Table 1. Proteins significantly upregulated in sEVs released by hCMEC/D3 exposed to amitriptyline.**

| Association according to KEGG pathway database: |
|---|
| Phagosome |
| Lysosome |
| Protein export |
| Antigen processing and presentation |
| Epstein-Barr virus infection |
| Viral myocarditis |

| Accession | Description | Protein name | Log 2 fold change | P-value |
|---|---|---|---|---|
| O60664 | Perilipin-3 | PLIN3 | 1.04 | 0.00 |
| P30876 | DNA-directed RNA polymerase II subunit RPB2 | POLR2B | 1.15 | 0.00 |
| Q06830 | Peroxiredoxin-1 | PRDX1 | 1.05 | 0.00 |
| O75475 | PC4 and SFRS1-interacting protein | PSIP1 | 1.57 | 0.00 |
| P28066 | Proteasome subunit alpha type-5 | PSMA5 | 1.70 | 0.00 |
| Q06323 | Proteasome activator complex subunit 1 | PSME1 | 1.39 | 0.00 |
| Q9UKM9 | RNA-binding protein Raly | RALY | 1.06 | 0.00 |
| Q14498 | RNA-binding protein 39 | RBM39 | 1.27 | 0.00 |
| Q04206 | Transcription factor p65 | RELA | 1.18 | 0.00 |
| P13489 | Ribonuclease inhibitor | RNH1 | 1.17 | 0.00 |
| P27635 | 60S ribosomal protein L10 | RPL10 | 1.00 | 0.00 |
| P35268 | 60S ribosomal protein L22 | RPL22 | 1.17 | 0.00 |
| P36578 | 60S ribosomal protein L4 | RPL4 | 1.11 | 0.00 |
| P62841 | 40S ribosomal protein S15 | RPS15 | 1.72 | 0.01 |
| Q9P2E9 | Ribosome-binding protein 1 | RRBP1 | 1.10 | 0.00 |
| Q14108 | Lysosome membrane protein 2 | SCARB2 | 1.48 | 0.00 |
| P53985 | Monocarboxylate transporter 1 | SLC16A1 | 1.12 | 0.00 |
| Q9H2H9 | Sodium-coupled neutral amino acid transporter 1 | SLC38A1 | 1.21 | 0.00 |
| Q96QD8 | Sodium-coupled neutral amino acid transporter 2 | SLC38A2 | 1.07 | 0.00 |
| Q15043 | Zinc transporter ZIP14 | SLC39A14 | 1.13 | 0.00 |
| P08195 | 4F2 cell-surface antigen heavy chain | SLC3A2 | 1.45 | 0.00 |
| P30825 | High affinity cationic amino acid transporter 1 | SLC7A1 | 1.08 | 0.02 |
| Q01650 | Large neutral amino acids transporter small subunit 1 | SLC7A5 | 1.16 | 0.00 |
| O60264 | SWI/SNF-related matrix-associated actin-dependent regulator of chromatin subfamily A member 5 | SMARCA5 | 1.03 | 0.00 |
| Q8TAQ2 | SWI/SNF complex subunit SMARCC2 | SMARCC2 | 1.16 | 0.00 |
| O95347 | Structural maintenance of chromosomes protein 2 | SMC2 | 1.76 | 0.00 |
| Q9UQE7 | Structural maintenance of chromosomes protein 3 | SMC3 | 1.91 | 0.00 |
| Q13501 | Sequestosome-1 | SQSTM1 | 3.70 | 0.00 |
| P61011 | Signal recognition particle 54 kDa protein | SRP54 | 1.15 | 0.00 |
| Q9UHB9 | Signal recognition particle subunit SRP68 | SRP68 | 1.17 | 0.00 |
| O76094 | Signal recognition particle subunit SRP72 | SRP72 | 1.09 | 0.00 |
| P05455 | Lupus La protein | SSB | 1.00 | 0.00 |
| O15400 | Syntaxin-7 | STX7 | 1.15 | 0.01 |
| Q13428 | Treacle protein | TCOF1 | 1.49 | 0.00 |
| Q9Y5L0 | Transportin-3 | TNPO3 | 1.60 | 0.00 |
| P06753 | Tropomyosin alpha-3 chain | TPM3 | 1.09 | 0.01 |
| Q13263 | Transcription intermediary factor 1-beta | TRIM28 | 1.29 | 0.00 |
| P49411 | Elongation factor Tu, mitochondrial | TUFM | 1.02 | 0.00 |
| P26368 | Splicing factor U2AF 65 kDa subunit | U2AF2 | 1.72 | 0.00 |
| O94874 | E3 UFM1-protein ligase 1 | UFL1 | 1.08 | 0.00 |
| Q16739 | Ceramide glucosyltransferase | UGCG | 1.11 | 0.03 |
| Q08AM6 | Protein VAC14 homolog | VAC14 | 1.24 | 0.00 |
| Q15836 | Vesicle-associated membrane protein 3 | VAMP3 | 2.07 | 0.00 |
| P08670 | Vimentin | VIM | 1.19 | 0.00 |
| O75436 | Vacuolar protein sorting-associated protein 26A | VPS26A | 1.25 | 0.00 |

Comparison of n=6 different samples/group obtained from 10 independent cell cultures.

**Supplementary Table 2. Proteins significantly downregulated in sEVs released by hCMEC/D3 exposed to amitriptyline.**

| Association according to KEGG pathway database: | |
|---|---|
| (yellow) | Extracellular matrix - receptor interaction and focal adhesion |

| Accession | Description | Protein name | Log 2 fold change | P-value |
|---|---|---|---|---|
| P24821 | Tenascin | TNC | -1.05 | 0.00 |
| P02647 | Apolipoprotein A-I | APOA1 | -1.05 | 0.00 |
| P08572 | Collagen alpha-2(IV) chain | COL4A2 | -1.06 | 0.00 |
| O00622 | CCN family member 1 | CCN1 | -1.13 | 0.00 |
| Q96N76 | Urocanate hydratase | UROC1 | -1.29 | 0.01 |
| P58107 | Epiplakin | EPPK1 | -1.31 | 0.01 |
| P02788 | Lactotransferrin | LTF | -1.34 | 0.00 |
| O76074 | cGMP-specific 3',5'-cyclic phosphodiesterase | PDE5A | -1.50 | 0.01 |
| P26022 | Pentraxin-related protein PTX3 | PTX3 | -2.28 | 0.00 |

Comparison of n=6 different samples/group obtained from 10 independent cell cultures.

### Supplementary Table 3. Differentially regulated protein networks identified by KEGG pathway database analysis.

| Term ID | Term description | Observed gene count | Background gene count | Strength | False discovery rate | Matching proteins in the network |
|---|---|---|---|---|---|---|
| hsa04145 | Phagosome | 7 | 145 | 0.93 | 0.0014 | VAMP3, CANX, ITGA5, LAMP1, STX7, HLA-C, HLA-B |
| hsa04142 | Lysosome | 8 | 123 | 1.06 | 0.00013 | CTSD, SCARB2, GUSB, GLB1, AP3S1, LAMP1,CTSB, AP1M1 |
| hsa03060 | Protein export | 3 | 23 | 1.36 | 0.0123 | SRP68, SRP72, SRP54 |
| hsa04612 | Antigen processing and presentation | 6 | 66 | 1.2 | 0.00027 | CANX, CTSB, HLA-C, PSME1, HLA-B, B2M |
| hsa05169 | Epstein-Barr virus infection | 7 | 194 | 0.8 | 0.0061 | EIF2AK2, HLA-C, POLR2B, RELA, CD44, HLA-B, VIM |
| hsa05416 | Viral myocarditis | 4 | 56 | 1.1 | 0.0123 | CD55, HLA-C, HLA-B, EIF4G1 |

| Term ID | Term description | Observed gene count | Background gene count | Strength | False discovery rate | Matching proteins in the network |
|---|---|---|---|---|---|---|
| hsa04512 | ECM-receptor interaction | 2 | 81 | 1.73 | 0.0125 | TNC,COL4A2 |
| hsa04510 | Focal adhesion | 2 | 197 | 1.34 | 0.0353 | TNC,COL4A2 |

Comparison of n=6 different samples/group obtained from 10 independent cell cultures.

### Supplementary Table 4. List of all proteins identified by label free proteomics analysis.

| Accession | Description | Coverage [%] | # PSMs | # Unique Peptides | Amitriptyline 1 | Amitriptyline 2 | Amitriptyline 3 | Amitriptyline 4 | Amitriptyline 5 | Amitriptyline 6 | Vehicle 1 | Vehicle 2 | Vehicle 3 | Vehicle 4 | Vehicle 5 | Vehicle 6 | Average amitriptyline | Average vehicle | Ratio amitriptyline/ vehicle | Log 2 fold change (amitriptyline /vehicle) | P-value |
|---|---|---|---|---|---|---|---|---|---|---|---|---|---|---|---|---|---|---|---|---|---|
| P68371 | Tubulin beta-4B chain OS=Homo sapiens OX=9606 GN=TUBB4B PE=1 SV=1 | 81 | 5349 | 2 | 1.70E+09 | 1.58E+09 | 1.79E+09 | 1.67E+09 | 1.72E+09 | 1.81E+09 | 1.39E+09 | 1.39E+09 | 1.58E+09 | 1.43E+09 | 1.55E+09 | 1.68E+09 | 1.71E+09 | 1.50E+09 | 1.14 | 0.19 | 0.00672027 |
| Q71U36 | Tubulin alpha-1A chain OS=Homo sapiens OX=9606 GN=TUBA1A PE=1 SV=1 | 77 | 3804 | 2 | 7.25E+06 | 8.77E+06 | 1.75E+07 | 1.02E+07 | 1.02E+07 | 1.47E+07 | 1.08E+07 | 1.02E+07 | 1.36E+07 | 6.14E+06 | 1.58E+07 | 1.32E+07 | 1.14E+07 | 1.16E+07 | 0.98 | -0.02 | 0.927398077 |
| Q9BQE3 | Tubulin alpha-1C chain OS=Homo sapiens OX=9606 GN=TUBA1C PE=1 SV=1 | 73 | 3200 | 2 | 1.78E+09 | 1.89E+09 | 1.99E+09 | 1.71E+09 | 1.82E+09 | 1.86E+09 | 1.55E+09 | 1.60E+09 | 1.61E+09 | 1.62E+09 | 1.75E+09 | 1.55E+09 | 1.84E+09 | 1.61E+09 | 1.14 | 0.19 | 0.000963406 |
| Q7L7L0 | Histone H2A type 3 OS=Homo sapiens OX=9606 GN=HIST3H2A PE=1 SV=3 | 64 | 2655 | 2 | 2.31E+08 | 2.19E+08 | 3.22E+08 | 2.42E+08 | 2.00E+08 | 1.76E+08 | 2.05E+08 | 2.78E+08 | 2.58E+08 | 3.62E+08 | 2.67E+08 | 1.64E+08 | 2.32E+08 | 2.55E+08 | 0.91 | -0.14 | 0.505091271 |
| P10412 | Histone H1.4 OS=Homo sapiens OX=9606 GN=H1-4 PE=1 SV=2 | 39 | 706 | 2 | 1.89E+09 | 1.91E+09 | 1.70E+09 | 1.82E+09 | 1.81E+09 | 1.75E+09 | 1.60E+09 | 1.69E+09 | 1.70E+09 | 1.62E+09 | 1.60E+09 | 1.58E+09 | 1.81E+09 | 1.63E+09 | 1.11 | 0.15 | 0.001566557 |
| P04259 | Keratin, type II cytoskeletal 6B OS=Homo sapiens OX=9606 GN=KRT6B PE=1 SV=5 | 31 | 563 | 2 | 4.07E+06 | 9.78E+05 | 1.35E+06 |  |  | 2.30E+06 | 6.20E+05 | 4.04E+06 | 2.78E+06 | 2.15E+06 | 1.07E+06 |  | 2.17E+06 | 2.13E+06 | 1.02 | 0.03 | 0.966063547 |
| Q562R1 | Beta-actin-like protein 2 OS=Homo sapiens OX=9606 GN=ACTBL2 PE=1 SV=2 | 32 | 1264 | 2 | 1.15E+06 | 4.90E+05 |  | 1.31E+06 | 3.10E+05 | 1.15E+06 | 1.75E+05 | 8.19E+05 | 1.05E+06 | 2.51E+05 |  |  | 8.83E+05 | 5.74E+05 | 1.54 | 0.62 | 0.329252562 |
| Q13310 | Polyadenylate-binding protein 4 OS=Homo sapiens OX=9606 GN=PABPC4 PE=1 SV=1 | 14 | 88 | 2 | 1.88E+06 | 2.49E+06 | 1.84E+06 | 2.07E+06 | 2.14E+06 | 1.65E+06 | 1.74E+06 | 1.85E+06 | 1.61E+06 | 1.73E+06 | 1.23E+06 | 1.49E+06 | 2.01E+06 | 1.61E+06 | 1.25 | 0.32 | 0.023859109 |
| P62820 | Ras-related protein Rab-1A OS=Homo sapiens OX=9606 GN=RAB1A PE=1 SV=3 | 54 | 91 | 2 | 6.67E+06 | 8.09E+06 | 4.82E+06 | 5.06E+06 | 4.88E+06 | 4.99E+06 | 6.53E+06 | 8.04E+06 | 4.64E+06 | 5.16E+06 | 4.89E+06 | 5.34E+06 | 5.75E+06 | 5.77E+06 | 1.00 | 0.00 | 0.984813058 |
| P08754 | Guanine nucleotide-binding protein G(i) subunit alpha OS=Homo sapiens OX=9606 GN=GNAI3 PE=1 SV=3 | 21 | 120 | 2 |  | 1.66E+06 | 2.31E+06 | 2.28E+06 | 1.12E+06 | 1.23E+06 | 1.53E+06 | 1.67E+06 | 1.26E+06 | 1.62E+06 |  | 9.11E+05 | 1.72E+06 | 1.40E+06 | 1.23 | 0.30 | 0.304238845 |
| P62873 | Guanine nucleotide-binding protein G(I)/G(S)/G(T) subunit beta-1 OS=Homo sapiens OX=9606 GN=GNB1 PE=1 SV=3 | 21 | 174 | 2 | 7.07E+06 | 6.56E+06 | 6.88E+06 | 7.16E+06 | 7.33E+06 | 8.12E+06 | 7.13E+06 | 7.19E+06 | 6.71E+06 | 8.14E+06 | 6.80E+06 | 6.19E+06 | 7.19E+06 | 7.03E+06 | 1.02 | 0.03 | 0.652075449 |
| O43488 | Aflatoxin B1 aldehyde reductase member 2 OS=Homo sapiens OX=9606 GN=AKR7A2 PE=1 SV=3 | 15 | 34 | 2 | 1.25E+07 | 1.47E+07 | 1.20E+07 | 1.29E+07 | 1.13E+07 | 9.13E+06 | 1.03E+07 | 8.81E+06 | 1.01E+07 | 7.81E+06 | 7.67E+06 | 6.67E+06 | 1.21E+07 | 8.57E+06 | 1.41 | 0.49 | 0.004860741 |
| Q15836 | Vesicle-associated membrane protein 3 OS=Homo sapiens OX=9606 GN=VAMP3 PE=1 SV=3 | 24 | 56 | 2 | 3.17E+06 | 4.53E+06 | 5.86E+06 | 6.42E+06 | 5.78E+06 | 8.20E+06 | 8.46E+05 |  | 8.62E+05 | 1.61E+06 | 7.29E+05 | 2.70E+06 | 5.66E+06 | 1.35E+06 | 4.19 | 2.07 | 0.000754519 |
| P50148 | Guanine nucleotide-binding protein G(q) subunit alpha OS=Homo sapiens OX=9606 GN=GNAQ PE=1 SV=4 | 16 | 32 | 2 | 4.48E+06 | 6.55E+06 | 4.74E+06 | 4.71E+06 | 3.69E+06 | 4.47E+06 | 2.77E+06 | 3.21E+06 | 3.17E+06 | 3.90E+06 | 2.29E+06 | 3.44E+06 | 4.77E+06 | 3.13E+06 | 1.53 | 0.61 | 0.006232056 |
| P28482 | Mitogen-activated protein kinase 1 OS=Homo sapiens OX=9606 GN=MAPK1 PE=1 SV=3 | 14 | 74 | 2 | 7.49E+06 | 7.48E+06 | 7.84E+06 | 8.75E+06 | 7.28E+06 | 7.87E+06 | 8.94E+06 | 8.37E+06 | 9.63E+06 | 8.19E+06 | 9.13E+06 | 8.29E+06 | 7.79E+06 | 8.76E+06 | 0.89 | -0.17 | 0.01196294 |
| Q16629 | Serine/arginine-rich splicing factor 7 OS=Homo sapiens OX=9606 GN=SRSF7 PE=1 SV=1 | 16 | 72 | 2 | 1.47E+07 | 1.20E+07 | 8.62E+06 | 9.13E+06 | 9.02E+06 | 9.60E+06 | 9.31E+06 | 8.31E+06 | 4.70E+06 | 8.15E+06 | 6.44E+06 | 6.95E+06 | 1.05E+07 | 7.31E+06 | 1.44 | 0.52 | 0.023709968 |
| P09972 | Fructose-bisphosphate aldolase C OS=Homo sapiens OX=9606 GN=ALDOC PE=1 SV=2 | 17 | 54 | 2 | 2.39E+06 |  |  |  |  | 2.78E+06 |  |  |  | 1.29E+06 | 1.48E+06 | 9.99E+05 | 2.58E+06 | 1.26E+06 | 2.06 | 1.04 | 0.030700266 |
| P31040 | Succinate dehydrogenase [ubiquinone] flavoprotein subunit, mitochondrial OS=Homo sapiens OX=9606 GN=SDHA PE=1 SV=2 | 6 | 20 | 2 | 4.37E+06 | 2.70E+06 | 2.37E+06 | 1.85E+06 | 1.80E+06 | 3.67E+06 | 3.10E+06 | 1.77E+06 | 6.09E+05 | 2.17E+06 | 2.00E+06 | 2.20E+06 | 2.79E+06 | 1.97E+06 | 1.41 | 0.50 | 0.158411389 |
| O43809 | Cleavage and polyadenylation specificity factor subunit 5 OS=Homo sapiens OX=9606 GN=NUDT21 PE=1 SV=1 | 21 | 14 | 2 | 1.71E+06 | 2.76E+06 | 2.91E+06 | 2.78E+06 | 2.99E+06 | 1.50E+06 | 1.40E+06 | 1.31E+06 | 1.98E+06 | 1.57E+06 | 8.67E+05 | 1.16E+06 | 2.44E+06 | 1.38E+06 | 1.77 | 0.82 | 0.00904725 |
| Q9NRY6 | Phospholipid scramblase 3 OS=Homo sapiens OX=9606 GN=PLSCR3 PE=1 SV=2 | 11 | 25 | 2 | 6.49E+06 | 3.68E+06 | 5.38E+06 | 1.77E+06 | 3.85E+06 | 5.26E+06 | 2.48E+06 | 3.15E+06 | 5.13E+06 | 2.55E+06 | 4.17E+06 | 3.60E+06 | 4.40E+06 | 3.51E+06 | 1.25 | 0.33 | 0.293607572 |
| P58107 | Epiplakin OS=Homo sapiens OX=9606 GN=EPPK1 PE=1 SV=3 | 5 | 34 | 2 | 1.11E+06 | 7.30E+05 | 1.85E+06 | 1.10E+06 | 7.03E+05 | 1.82E+06 | 5.03E+06 | 2.50E+06 | 2.12E+06 | 2.07E+06 | 3.13E+06 | 3.26E+06 | 1.22E+06 | 3.02E+06 | 0.40 | -1.31 | 0.008304223 |
| Q01650 | Large neutral amino acids transporter small subunit 1 OS=Homo sapiens OX=9606 GN=SLC7A5 PE=1 SV=2 | 6 | 34 | 2 | 5.84E+06 | 6.43E+06 | 7.64E+06 | 7.02E+06 | 6.73E+06 | 6.97E+06 | 3.22E+06 | 3.04E+06 | 3.40E+06 | 3.69E+06 | 2.48E+06 | 2.39E+06 | 6.77E+06 | 3.04E+06 | 2.23 | 1.16 | 5.65275E-07 |
| P28161 | Glutathione S-transferase Mu 2 OS=Homo sapiens OX=9606 GN=GSTM2 PE=1 SV=2 | 13 | 54 | 2 | 2.88E+07 | 2.43E+07 | 2.28E+07 | 2.08E+07 | 2.16E+07 | 2.31E+07 | 4.14E+07 | 3.48E+07 | 3.19E+07 | 2.76E+07 | 3.23E+07 | 3.26E+07 | 2.36E+07 | 3.34E+07 | 0.71 | -0.50 | 0.001766335 |
| P46778 | 60S ribosomal protein L21 OS=Homo sapiens OX=9606 GN=RPL21 PE=1 SV=2 | 14 | 50 | 2 | 2.27E+07 | 2.98E+07 | 3.00E+07 | 3.77E+07 | 2.95E+07 | 2.83E+07 | 1.55E+07 | 1.71E+07 | 1.72E+07 | 1.68E+07 | 1.32E+07 | 1.13E+07 | 2.97E+07 | 1.52E+07 | 1.96 | 0.97 | 0.000232639 |
| Q13765 | Nascent polypeptide-associated complex subunit alpha OS=Homo sapiens OX=9606 GN=NACA PE=1 SV=1 | 13 | 36 | 2 | 1.03E+07 | 1.14E+07 | 8.21E+06 | 1.21E+07 | 1.28E+07 | 1.03E+07 | 7.53E+06 | 7.70E+06 | 7.42E+06 | 7.13E+06 | 6.17E+06 | 5.83E+06 | 1.08E+07 | 6.96E+06 | 1.56 | 0.64 | 0.001031881 |
| Q15286 | Ras-related protein Rab-35 OS=Homo sapiens OX=9606 GN=RAB35 PE=1 SV=1 | 23 | 63 | 2 | 1.62E+06 | 1.57E+06 | 2.59E+06 | 2.60E+06 | 2.68E+06 | 2.96E+06 | 1.20E+06 | 1.56E+06 | 1.57E+06 | 2.25E+06 | 1.60E+06 | 1.15E+06 | 2.34E+06 | 1.56E+06 | 1.50 | 0.59 | 0.025175228 |
| P21964 | Catechol O-methyltransferase OS=Homo sapiens OX=9606 GN=COMT PE=1 SV=2 | 12 | 16 | 2 | 7.52E+05 | 5.21E+06 | 1.38E+06 | 3.17E+06 | 1.29E+06 | 5.96E+06 | 2.23E+06 | 8.44E+06 | 3.10E+06 | 5.18E+06 | 7.75E+06 | 6.81E+06 | 2.96E+06 | 5.59E+06 | 0.53 | -0.92 | 0.084567692 |
| P53999 | Activated RNA polymerase II transcriptional coactivator p15 OS=Homo sapiens OX=9606 GN=SUB1 PE=1 SV=3 | 19 | 28 | 2 | 8.04E+06 | 6.16E+06 | 3.29E+06 | 6.73E+06 | 3.76E+06 | 6.25E+06 | 6.15E+06 | 6.27E+06 | 3.29E+06 | 1.57E+06 | 3.10E+06 | 4.00E+06 | 5.71E+06 | 4.06E+06 | 1.40 | 0.49 | 0.15204 |
| O14617 | AP-3 complex subunit delta-1 OS=Homo sapiens OX=9606 GN=AP3D1 PE=1 SV=1 | 3 | 32 | 2 | 6.69E+06 | 7.81E+06 | 6.39E+06 | 4.07E+06 | 5.34E+06 | 5.52E+06 | 3.17E+06 | 4.50E+06 | 4.22E+06 | 3.47E+06 | 4.84E+06 | 3.47E+06 | 5.97E+06 | 3.95E+06 | 1.51 | 0.60 | 0.010057096 |
| Q16772 | Glutathione S-transferase A3 OS=Homo sapiens OX=9606 GN=GSTA3 PE=1 SV=3 | 21 | 263 | 2 | 3.54E+07 | 4.39E+07 | 6.24E+07 | 6.21E+07 | 6.53E+07 | 6.47E+07 | 4.47E+07 | 5.02E+07 | 8.98E+07 | 8.96E+07 | 9.45E+07 | 7.06E+07 | 5.56E+07 | 7.32E+07 | 0.76 | -0.40 | 0.124095599 |
| P10301 | Ras-related protein R-Ras OS=Homo sapiens OX=9606 GN=RRAS PE=1 SV=1 | 20 | 28 | 2 | 3.74E+06 | 4.33E+06 | 3.71E+06 | 4.14E+06 | 3.36E+06 | 4.89E+06 | 3.35E+06 | 2.13E+06 | 1.82E+06 | 2.75E+06 | 2.59E+06 | 1.56E+06 | 4.03E+06 | 2.36E+06 | 1.70 | 0.77 | 0.000838833 |
| Q08170 | Serine/arginine-rich splicing factor 4 OS=Homo sapiens OX=9606 GN=SRSF4 PE=1 SV=2 | 11 | 75 | 2 | 2.55E+07 | 2.80E+07 | 3.10E+07 | 3.31E+07 | 3.72E+07 | 3.08E+07 | 2.53E+07 | 2.62E+07 | 2.83E+07 | 2.63E+07 | 2.41E+07 | 2.37E+07 | 3.09E+07 | 2.57E+07 | 1.21 | 0.27 | 0.022541621 |
| P13693 | Translationally-controlled tumor protein OS=Homo sapiens OX=9606 GN=TPT1 PE=1 SV=1 | 31 | 10 | 2 | 4.54E+06 | 3.91E+06 | 3.52E+06 | 4.06E+06 | 1.68E+06 | 4.73E+06 | 1.96E+06 | 3.99E+06 | 3.09E+06 | 2.22E+06 | 6.05E+06 | 5.26E+06 | 3.74E+06 | 3.76E+06 | 0.99 | -0.01 | 0.980630094 |
| Q9HC07 | Transmembrane protein 165 OS=Homo sapiens OX=9606 GN=TMEM165 PE=1 SV=1 | 14 | 18 | 2 | 6.87E+06 | 7.12E+06 | 5.89E+06 | 7.69E+06 | 7.63E+06 | 6.85E+06 | 5.41E+06 | 5.87E+06 | 7.03E+06 | 5.82E+06 | 7.84E+06 | 3.76E+06 | 7.01E+06 | 5.95E+06 | 1.18 | 0.24 | 0.139131246 |
| Q6EMK4 | Vasorin OS=Homo sapiens OX=9606 GN=VASN PE=1 SV=1 | 4 | 18 | 2 | 4.60E+06 | 3.95E+06 | 2.46E+06 | 3.63E+06 | 3.18E+06 | 3.00E+06 | 3.64E+06 | 4.95E+06 | 8.11E+05 | 1.64E+06 | 5.59E+05 | 2.76E+06 | 3.47E+06 | 2.39E+06 | 1.45 | 0.54 | 0.203006177 |
| P48059 | LIM and senescent cell antigen-like-containing domain protein 1 OS=Homo sapiens OX=9606 GN=LIMS1 PE=1 SV=4 | 12 | 46 | 2 | 1.12E+07 | 7.79E+06 | 7.85E+06 | 9.96E+06 | 9.95E+06 | 9.23E+06 | 1.31E+07 | 1.16E+07 | 1.58E+07 | 1.02E+07 | 1.67E+07 | 1.03E+07 | 9.33E+06 | 1.29E+07 | 0.72 | -0.47 | 0.022645577 |
| Q9BWD1 | Acetyl-CoA acetyltransferase, cytosolic OS=Homo sapiens OX=9606 GN=ACAT2 PE=1 SV=2 | 13 | 6 | 2 | 2.76E+06 | 2.23E+06 | 1.64E+06 | 2.47E+06 | 1.71E+06 | 2.31E+06 | 4.73E+06 | 2.91E+06 | 1.23E+06 | 1.69E+06 | 1.10E+06 | 9.05E+05 | 2.19E+06 | 2.10E+06 | 1.04 | 0.06 | 0.889511115 |
| P55809 | Succinyl-CoA:3-ketoacid coenzyme A transferase 1, mitochondrial OS=Homo sapiens OX=9606 GN=OXCT1 PE=1 SV=1 | 9 | 10 | 2 | 3.56E+06 | 2.72E+06 | 2.61E+06 | 2.80E+06 | 2.33E+06 | 4.23E+06 | 2.57E+06 | 7.20E+05 | 1.48E+06 | 1.49E+06 | 2.42E+06 | 2.03E+06 | 3.04E+06 | 1.78E+06 | 1.71 | 0.77 | 0.01120172 |
| P11310 | Medium-chain specific acyl-CoA dehydrogenase, mitochondrial OS=Homo sapiens OX=9606 GN=ACADM PE=1 SV=1 | 10 | 14 | 2 | 1.59E+06 | 2.26E+06 | 1.85E+06 | 1.62E+06 | 1.50E+06 | 1.77E+06 | 3.89E+05 | 4.10E+05 | 9.44E+05 | 2.55E+05 | 8.95E+05 | 2.98E+05 | 1.76E+06 | 5.32E+05 | 3.32 | 1.73 | 2.66998E-05 |
| Q9NRV9 | Heme-binding protein 1 OS=Homo sapiens OX=9606 GN=HEBP1 PE=1 SV=1 | 17 | 10 | 2 | 4.40E+06 | 9.23E+05 | 2.82E+06 | 4.35E+06 | 3.96E+06 | 3.64E+06 | 6.30E+06 | 5.55E+06 | 4.23E+06 | 3.86E+06 | 4.06E+06 | 4.00E+06 | 3.35E+06 | 4.67E+06 | 0.72 | -0.48 | 0.083183856 |
| P31942 | Heterogeneous nuclear ribonucleoprotein H3 OS=Homo sapiens OX=9606 GN=HNRNPH3 PE=1 SV=2 | 8 | 32 | 2 | 5.41E+06 | 4.94E+06 | 4.51E+06 | 4.36E+06 | 4.76E+06 | 3.97E+06 | 3.30E+06 | 2.78E+06 | 2.89E+06 | 3.07E+06 | 3.05E+06 | 1.86E+06 | 4.66E+06 | 2.82E+06 | 1.65 | 0.72 | 8.53264E-05 |
| P35222 | Catenin beta-1 OS=Homo sapiens OX=9606 GN=CTNNB1 PE=1 SV=1 | 6 | 52 | 2 | 4.48E+06 | 3.52E+06 | 3.39E+06 | 3.57E+06 | 5.25E+06 | 5.79E+06 | 2.76E+06 | 2.80E+06 | 4.04E+06 | 2.93E+06 | 2.78E+06 | 2.05E+06 | 4.33E+06 | 2.89E+06 | 1.50 | 0.58 | 0.017518357 |
| Q15833 | Syntaxin-binding protein 2 OS=Homo sapiens OX=9606 GN=STXBP2 PE=1 SV=2 | 6 | 32 | 2 | 4.64E+06 | 4.42E+06 | 4.54E+06 | 4.25E+06 | 3.87E+06 | 4.20E+06 | 5.12E+06 | 5.17E+06 | 3.30E+06 | 4.11E+06 | 3.56E+06 | 4.40E+06 | 4.32E+06 | 4.28E+06 | 1.01 | 0.01 | 0.902249295 |
| P29966 | Myristoylated alanine-rich C-kinase substrate OS=Homo sapiens OX=9606 GN=MARCKS PE=1 SV=4 | 12 | 18 | 2 | 2.29E+06 | 3.07E+06 | 6.00E+06 | 6.44E+06 | 4.80E+06 | 4.95E+06 | 4.49E+06 | 4.97E+06 | 5.67E+06 | 4.65E+06 | 4.08E+06 | 4.04E+06 | 4.59E+06 | 4.65E+06 | 0.99 | -0.02 | 0.936491304 |
| Q14520 | Hyaluronan-binding protein 2 OS=Homo sapiens OX=9606 GN=HABP2 PE=1 SV=1 | 4 | 30 | 2 | 8.43E+06 | 8.76E+06 | 8.48E+06 | 1.31E+07 | 9.24E+06 | 4.91E+06 | 1.40E+07 | 6.73E+06 | 1.37E+07 | 1.13E+07 | 8.29E+05 | 3.31E+06 | 8.82E+06 | 8.29E+06 | 1.06 | 0.09 | 0.839489266 |
| Q13247 | Serine/arginine-rich splicing factor 6 OS=Homo sapiens OX=9606 GN=SRSF6 PE=1 SV=2 | 15 | 51 | 2 | 4.53E+06 | 5.09E+06 | 4.19E+06 | 4.62E+06 | 4.02E+06 | 3.95E+06 | 4.84E+06 | 4.42E+06 | 3.99E+06 | 4.59E+06 | 3.83E+06 | 3.51E+06 | 4.40E+06 | 4.20E+06 | 1.05 | 0.07 | 0.466719361 |
| P22105 | Tenascin-X OS=Homo sapiens OX=9606 GN=TNXB PE=1 SV=5 | 2 | 34 | 2 | 4.40E+06 | 4.90E+06 | 6.75E+06 | 6.34E+06 | 4.84E+06 | 5.15E+06 |  | 7.79E+06 | 7.59E+06 | 7.40E+06 | 6.61E+06 | 6.81E+06 | 5.40E+06 | 7.24E+06 | 0.75 | -0.42 | 0.003179203 |
| P60903 | Protein S100-A10 OS=Homo sapiens OX=9606 GN=S100A10 PE=1 SV=2 | 28 | 47 | 2 | 1.54E+07 | 1.90E+07 | 1.29E+07 | 1.63E+07 | 1.48E+07 | 1.13E+07 | 9.55E+06 | 1.09E+07 | 5.99E+06 | 1.01E+07 | 6.56E+06 | 7.49E+06 | 1.49E+07 | 8.42E+06 | 1.77 | 0.83 | 0.00090011 |
| P02042 | Hemoglobin subunit delta OS=Homo sapiens OX=9606 GN=HBD PE=1 SV=2 | 13 | 1536 | 2 | 1.17E+09 | 1.12E+09 | 1.00E+09 | 9.20E+08 | 1.15E+09 | 1.11E+09 | 1.24E+09 | 1.28E+09 | 1.23E+09 | 1.36E+09 | 1.24E+09 | 1.43E+09 | 1.08E+09 | 1.30E+09 | 0.83 | -0.26 | 0.001988061 |
| P19827 | Inter-alpha-trypsin inhibitor heavy chain H1 OS=Homo sapiens OX=9606 GN=ITIH1 PE=1 SV=3 | 3 | 28 | 2 | 3.22E+06 | 3.37E+06 | 5.34E+06 | 6.63E+06 | 2.51E+06 | 5.38E+06 | 7.86E+06 | 7.66E+06 | 3.15E+06 | 6.84E+06 | 6.66E+06 | 5.64E+06 | 4.41E+06 | 6.30E+06 | 0.70 | -0.52 | 0.078461488 |
| P48147 | Prolyl endopeptidase OS=Homo sapiens OX=9606 GN=PREP PE=1 SV=2 | 4 | 37 | 2 | 3.93E+06 | 4.10E+06 | 2.74E+06 | 2.99E+06 | 2.92E+06 | 4.78E+06 | 4.01E+06 | 4.84E+06 | 3.66E+06 | 3.73E+06 | 3.99E+06 | 4.24E+06 | 3.57E+06 | 4.08E+06 | 0.88 | -0.19 | 0.219969684 |
| P52565 | Rho GDP-dissociation inhibitor 1 OS=Homo sapiens OX=9606 GN=ARHGDIA PE=1 SV=3 | 16 | 28 | 2 | 4.13E+06 | 4.81E+06 | 6.03E+06 | 4.53E+06 | 5.70E+06 | 4.80E+06 | 3.62E+06 | 3.84E+06 | 4.07E+06 | 3.88E+06 | 2.84E+06 | 5.55E+06 | 5.00E+06 | 3.97E+06 | 1.26 | 0.33 | 0.052468929 |
| Q01844 | RNA-binding protein EWS OS=Homo sapiens OX=9606 GN=EWSR1 PE=1 SV=1 | 4 | 26 | 2 | 7.39E+06 | 7.37E+06 | 8.02E+06 | 7.27E+06 | 6.40E+06 | 6.57E+06 | 3.99E+06 | 4.55E+06 | 3.26E+06 | 4.09E+06 | 3.52E+06 | 3.46E+06 | 7.17E+06 | 3.81E+06 | 1.88 | 0.91 | 1.18302E-06 |
| Q09028 | Histone-binding protein RBBP4 OS=Homo sapiens OX=9606 GN=RBBP4 PE=1 SV=3 | 16 | 41 | 2 |  |  | 2.95E+06 | 2.93E+06 |  | 1.37E+06 |  |  |  | 1.49E+06 | 2.49E+06 |  | 2.42E+06 | 1.99E+06 | 1.22 | 0.28 | 0.599353373 |
| P28065 | Proteasome subunit beta type-9 OS=Homo sapiens OX=9606 GN=PSMB9 PE=1 SV=2 | 14 | 20 | 2 | 3.83E+06 | 3.88E+06 | 3.01E+06 | 3.84E+06 | 3.44E+06 | 3.53E+06 | 3.62E+06 | 4.36E+06 | 3.17E+06 | 3.74E+06 | 3.36E+06 | 1.18E+06 | 3.59E+06 | 3.24E+06 | 1.11 | 0.15 | 0.480994747 |
| Q96N76 | Urocanate hydratase OS=Homo sapiens OX=9606 GN=UROC1 PE=1 SV=1 | 5 | 14 | 2 | 7.91E+05 | 8.26E+05 | 1.86E+06 | 1.30E+06 | 1.44E+06 | 8.62E+05 | 4.06E+06 | 1.94E+06 | 3.61E+06 | 3.11E+06 | 3.03E+06 | 1.51E+06 | 1.18E+06 | 2.88E+06 | 0.41 | -1.29 | 0.006216674 |
| P23258 | Tubulin gamma-1 chain OS=Homo sapiens OX=9606 GN=TUBG1 PE=1 SV=2 | 11 | 8 | 2 | 6.33E+05 | 9.53E+05 | 2.14E+06 | 9.58E+05 | 1.75E+06 | 3.56E+05 | 6.86E+05 | 3.61E+05 |  | 4.35E+05 |  |  | 1.13E+06 | 4.94E+05 | 2.29 | 1.19 | 0.072918266 |
| Q14112 | Nidogen-2 OS=Homo sapiens OX=9606 GN=NID2 PE=1 SV=3 | 3 | 6 | 2 | 6.45E+05 |  | 1.88E+06 | 1.28E+06 | 5.37E+05 |  | 1.71E+06 |  | 4.62E+05 | 5.83E+05 | 6.67E+05 | 1.99E+06 | 1.08E+06 | 1.08E+06 | 1.00 | 0.00 | 0.995317394 |
| P15531 | Nucleoside diphosphate kinase A OS=Homo sapiens OX=9606 GN=NME1 PE=1 SV=1 | 30 | 32 | 2 | 6.10E+06 | 7.47E+06 | 7.81E+06 | 9.56E+06 | 8.11E+06 | 8.98E+06 | 8.61E+06 | 8.24E+06 | 7.75E+06 | 7.02E+06 | 7.72E+06 | 6.69E+06 | 8.01E+06 | 7.67E+06 | 1.04 | 0.06 | 0.574551007 |
| P07203 | Glutathione peroxidase 1 OS=Homo sapiens OX=9606 GN=GPX1 PE=1 SV=4 | 18 | 21 | 2 | 1.48E+06 | 2.98E+06 | 2.97E+06 | 4.07E+06 | 2.45E+06 | 2.23E+06 | 5.10E+06 | 5.03E+06 | 5.28E+06 | 3.75E+06 | 4.42E+06 | 2.97E+06 | 2.70E+06 | 4.42E+06 | 0.61 | -0.71 | 0.007129705 |
| Q9BXN1 | Asporin OS=Homo sapiens OX=9606 GN=ASPN PE=1 SV=2 | 13 | 4 | 2 | 1.57E+06 | 5.51E+05 |  | 6.24E+05 |  | 9.06E+05 | 1.63E+06 | 1.43E+06 |  | 1.22E+06 | 5.88E+05 | 6.97E+05 | 9.12E+05 | 1.11E+06 | 0.82 | -0.29 | 0.538505832 |
| Q9BZQ8 | Protein Niban 1 OS=Homo sapiens OX=9606 GN=NIBAN1 PE=1 SV=1 | 4 | 16 | 2 | 2.94E+06 | 1.48E+06 | 2.01E+06 | 2.53E+06 | 2.11E+06 | 2.42E+06 | 2.82E+06 | 2.21E+06 | 2.32E+06 | 2.62E+06 | 2.07E+06 | 1.51E+06 | 2.25E+06 | 2.26E+06 | 1.00 | -0.01 | 0.973296315 |
| Q96DI7 | U5 small nuclear ribonucleoprotein 40 kDa protein OS=Homo sapiens OX=9606 GN=SNRNP40 PE=1 SV=1 | 11 | 8 | 2 |  | 1.37E+06 | 6.47E+05 |  |  | 2.75E+06 | 5.11E+05 | 7.03E+05 |  | 6.81E+05 | 1.06E+06 | 4.48E+05 | 1.59E+06 | 6.81E+05 | 2.33 | 1.22 | 0.277857593 |
| P32754 | 4-hydroxyphenylpyruvate dioxygenase OS=Homo sapiens OX=9606 GN=HPD PE=1 SV=2 | 9 | 18 | 2 | 2.93E+06 | 2.99E+06 | 2.78E+06 | 3.08E+06 | 2.49E+06 | 2.97E+06 | 4.76E+06 | 4.71E+06 | 4.54E+06 | 3.45E+06 | 3.89E+06 | 2.53E+06 | 2.87E+06 | 3.98E+06 | 0.72 | -0.47 | 0.02561566 |
| Q96T76 | MMS19 nucleotide excision repair protein homolog OS=Homo sapiens OX=9606 GN=MMS19 PE=1 SV=2 | 4 | 6 | 2 | 1.08E+06 | 2.00E+06 | 1.48E+06 | 1.77E+06 | 1.45E+06 | 1.58E+06 | 1.25E+06 | 1.52E+06 | 1.36E+06 | 8.46E+05 | 1.09E+06 | 7.99E+05 | 1.56E+06 | 1.14E+06 | 1.36 | 0.45 | 0.037089734 |
| O14763 | Tumor necrosis factor receptor superfamily member 10B OS=Homo sapiens OX=9606 GN=TNFRSF10B PE=1 SV=2 | 7 | 17 | 2 | 7.29E+06 | 4.71E+06 | 5.46E+06 | 4.62E+06 | 3.90E+06 | 4.62E+06 | 3.38E+06 | 2.75E+06 | 2.96E+06 | 2.33E+06 | 3.67E+06 | 3.18E+06 | 5.10E+06 | 3.05E+06 | 1.67 | 0.74 | 0.006265415 |
| P68402 | Platelet-activating factor acetylhydrolase IB subunit beta OS=Homo sapiens OX=9606 GN=PAFAH1B2 PE=1 SV=1 | 19 | 6 | 2 |  |  |  | 8.67E+05 | 1.03E+06 |  | 1.02E+06 | 1.32E+06 | 1.18E+06 |  | 1.75E+06 | 7.05E+05 | 9.47E+05 | 1.20E+06 | 0.79 | -0.34 | 0.246945279 |
| P07954 | Fumarate hydratase, mitochondrial OS=Homo sapiens OX=9606 GN=FH PE=1 SV=3 | 7 | 18 | 2 | 6.53E+06 | 7.77E+06 | 4.23E+06 | 5.83E+06 | 5.59E+06 | 6.40E+06 | 1.00E+07 | 8.59E+06 | 7.71E+06 | 7.69E+06 | 7.55E+06 | 7.02E+06 | 6.06E+06 | 8.10E+06 | 0.75 | -0.42 | 0.010547801 |
| P20645 | Cation-dependent mannose-6-phosphate receptor OS=Homo sapiens OX=9606 GN=M6PR PE=1 SV=1 | 12 | 42 | 2 | 5.35E+06 | 5.03E+06 | 3.93E+06 | 4.41E+06 | 3.92E+06 | 4.58E+06 | 4.56E+06 | 4.01E+06 | 4.26E+06 | 3.58E+06 | 3.90E+06 | 2.38E+06 | 4.54E+06 | 3.78E+06 | 1.20 | 0.26 | 0.084349745 |
| P04637 | Cellular tumor antigen p53 OS=Homo sapiens OX=9606 GN=TP53 PE=1 SV=4 | 9 | 17 | 2 | 3.95E+06 | 5.32E+06 | 4.00E+06 | 3.69E+06 | 3.02E+06 | 5.58E+06 | 2.95E+06 | 3.14E+06 | 1.68E+06 | 2.14E+06 | 2.56E+06 | 6.86E+05 | 4.26E+06 | 2.19E+06 | 1.94 | 0.96 | 0.003724725 |
| P61081 | NEDD8-conjugating enzyme Ubc12 OS=Homo sapiens OX=9606 GN=UBE2M PE=1 SV=1 | 15 | 16 | 2 | 2.91E+06 | 2.68E+06 | 1.92E+06 | 2.43E+06 | 2.85E+06 | 3.30E+06 | 2.90E+06 | 2.40E+06 | 2.34E+06 | 3.00E+06 | 2.19E+06 | 1.91E+06 | 2.68E+06 | 2.46E+06 | 1.09 | 0.13 | 0.403916909 |
| P47914 | 60S ribosomal protein L29 OS=Homo sapiens OX=9606 GN=RPL29 PE=1 SV=2 | 14 | 30 | 2 | 1.21E+07 | 1.20E+07 | 1.28E+07 | 1.45E+07 | 1.19E+07 | 1.24E+07 | 7.23E+06 | 7.29E+06 | 5.61E+06 | 7.08E+06 | 6.66E+06 | 6.18E+06 | 1.26E+07 | 6.68E+06 | 1.89 | 0.92 | 7.07417E-07 |
| O15127 | Secretory carrier-associated membrane protein 2 OS=Homo sapiens OX=9606 GN=SCAMP2 PE=1 SV=2 | 10 | 4 | 2 | 2.76E+06 | 2.60E+06 | 2.88E+06 | 2.68E+06 | 3.05E+06 | 3.01E+06 | 2.28E+06 | 2.11E+06 | 1.92E+06 |  | 2.11E+06 | 1.83E+06 | 2.83E+06 | 2.05E+06 | 1.38 | 0.47 | 6.17135E-05 |
| P50851 | Lipopolysaccharide-responsive and beige-like anchor protein OS=Homo sapiens OX=9606 GN=LRBA PE=1 SV=4 | 1 | 4 | 2 | 1.44E+06 | 7.10E+05 |  |  | 5.52E+05 | 5.34E+05 | 1.15E+06 |  | 4.95E+05 | 7.81E+05 | 3.82E+05 | 3.61E+05 | 8.10E+05 | 6.34E+05 | 1.28 | 0.35 | 0.528086604 |
| Q9UI10 | Translation initiation factor eIF-2B subunit delta OS=Homo sapiens OX=9606 GN=EIF2B4 PE=1 SV=2 | 9 | 6 | 2 | 3.14E+06 |  | 1.79E+06 | 1.94E+06 | 2.70E+06 |  |  |  |  |  |  |  | 2.39E+06 |  |  |  |  |
| Q9H0D6 | 5'-3' exoribonuclease 2 OS=Homo sapiens OX=9606 GN=XRN2 PE=1 SV=1 | 3 | 14 | 2 | 4.29E+06 | 4.27E+06 | 3.79E+06 | 3.75E+06 | 8.78E+05 | 1.98E+06 | 2.98E+06 | 2.93E+06 | 2.27E+06 | 1.41E+06 | 3.69E+06 | 2.01E+06 | 3.16E+06 | 2.55E+06 | 1.24 | 0.31 | 0.382087325 |
| Q9UIA9 | Exportin-7 OS=Homo sapiens OX=9606 GN=XPO7 PE=1 SV=3 | 3 | 18 | 2 | 2.70E+06 | 2.98E+06 | 3.04E+06 | 2.32E+06 | 3.76E+06 | 2.99E+06 | 4.10E+06 | 1.69E+06 | 3.32E+06 | 2.46E+06 | 3.52E+06 | 2.52E+06 | 2.97E+06 | 2.94E+06 | 1.01 | 0.01 | 0.942642798 |
| Q6XQN6 | Nicotinate phosphoribosyltransferase OS=Homo sapiens OX=9606 GN=NAPRT PE=1 SV=2 | 6 | 20 | 2 | 4.82E+06 | 2.34E+06 | 2.24E+06 | 3.09E+06 | 2.03E+06 | 2.89E+06 | 4.96E+06 | 4.31E+06 | 1.21E+06 | 1.89E+06 | 2.36E+06 | 2.54E+06 | 2.90E+06 | 2.88E+06 | 1.01 | 0.01 | 0.974287847 |
| P27824 | Calnexin OS=Homo sapiens OX=9606 GN=CANX PE=1 SV=2 | 5 | 12 | 2 | 1.27E+06 | 1.33E+06 | 2.15E+06 | 2.26E+06 | 1.66E+06 | 1.91E+06 | 1.10E+06 |  | 5.60E+05 | 5.43E+05 | 8.90E+05 | 5.42E+05 | 1.76E+06 | 7.28E+05 | 2.42 | 1.28 | 0.000847091 |
| P30086 | Phosphatidylethanolamine-binding protein 1 OS=Homo sapiens OX=9606 GN=PEBP1 PE=1 SV=3 | 15 | 24 | 2 | 7.72E+06 | 6.61E+06 | 7.36E+06 | 9.24E+06 | 7.41E+06 | 6.86E+06 | 8.16E+06 | 8.99E+06 | 9.48E+06 | 9.81E+06 | 9.56E+06 | 8.31E+06 | 7.53E+06 | 9.05E+06 | 0.83 | -0.27 | 0.010091167 |
| Q9C005 | Protein dpy-30 homolog OS=Homo sapiens OX=9606 GN=DPY30 PE=1 SV=1 | 36 | 20 | 2 | 2.12E+06 | 1.66E+06 | 3.49E+06 | 3.40E+06 | 2.11E+06 | 1.64E+06 | 3.13E+06 | 8.72E+05 | 1.69E+06 | 1.82E+06 | 1.52E+06 | 5.19E+05 | 2.40E+06 | 1.59E+06 | 1.51 | 0.59 | 0.137075982 |
| Q5VW32 | BRO1 domain-containing protein BROX OS=Homo sapiens OX=9606 GN=BROX PE=1 SV=1 | 10 | 16 | 2 | 1.80E+06 | 2.31E+06 | 2.54E+06 | 2.00E+06 | 2.40E+06 | 2.06E+06 | 1.77E+06 | 1.73E+06 | 1.26E+06 | 1.44E+06 | 1.70E+06 | 1.49E+06 | 2.19E+06 | 1.57E+06 | 1.40 | 0.48 | 0.001555842 |
| Q8WWI5 | Choline transporter-like protein 1 OS=Homo sapiens OX=9606 GN=SLC44A1 PE=1 SV=1 | 4 | 40 | 2 | 7.70E+06 | 8.24E+06 | 4.04E+06 | 5.64E+06 | 5.93E+06 | 5.33E+06 | 5.94E+06 | 5.93E+06 | 2.77E+06 | 3.89E+06 | 3.81E+06 | 4.38E+06 | 6.15E+06 | 4.45E+06 | 1.38 | 0.46 | 0.066931701 |
| Q92544 | Transmembrane 9 superfamily member 4 OS=Homo sapiens OX=9606 GN=TM9SF4 PE=1 SV=2 | 6 | 16 | 2 | 4.61E+06 | 2.15E+06 | 2.70E+06 | 2.03E+06 | 1.10E+06 | 2.30E+06 | 1.55E+06 | 1.52E+06 | 1.60E+06 | 6.96E+05 | 1.75E+06 | 1.63E+06 | 2.48E+06 | 1.46E+06 | 1.70 | 0.77 | 0.087713963 |
| Q9UJ70 | N-acetyl-D-glucosamine kinase OS=Homo sapiens OX=9606 GN=NAGK PE=1 SV=4 | 7 | 20 | 2 | 2.15E+06 | 2.52E+06 | 2.20E+06 | 2.20E+06 | 2.44E+06 | 2.28E+06 | 7.29E+05 | 2.60E+06 | 2.12E+06 | 2.09E+06 | 1.16E+06 | 1.87E+06 | 2.30E+06 | 1.76E+06 | 1.31 | 0.39 | 0.115645377 |
| Q16739 | Ceramide glucosyltransferase OS=Homo sapiens OX=9606 GN=UGCG PE=1 SV=1 | 7 | 6 | 2 | 2.26E+06 | 2.70E+06 | 1.44E+06 | 2.18E+06 | 8.03E+05 | 2.13E+06 | 5.61E+05 | 1.69E+06 | 4.32E+05 |  | 1.51E+06 | 2.71E+05 | 1.92E+06 | 8.92E+05 | 2.15 | 1.11 | 0.032208519 |
| Q9Y295 | Developmentally-regulated GTP-binding protein 1 OS=Homo sapiens OX=9606 GN=DRG1 PE=1 SV=1 | 11 | 6 | 2 | 1.33E+06 | 1.52E+06 | 1.18E+06 | 1.30E+06 | 1.01E+06 | 9.53E+05 |  |  | 7.79E+05 |  |  | 1.08E+06 | 1.22E+06 | 9.31E+05 | 1.31 | 0.38 | 0.265019588 |
| P00750 | Tissue-type plasminogen activator OS=Homo sapiens OX=9606 GN=PLAT PE=1 SV=1 | 6 | 4 | 2 | 1.55E+06 | 2.11E+06 | 1.91E+06 | 1.79E+06 | 1.48E+06 | 1.55E+06 | 2.15E+06 | 2.35E+06 | 2.12E+06 | 2.09E+06 | 1.43E+06 | 1.54E+06 | 1.73E+06 | 1.95E+06 | 0.89 | -0.17 | 0.26612819 |
| Q14103 | Heterogeneous nuclear ribonucleoprotein D0 OS=Homo sapiens OX=9606 GN=HNRNPD PE=1 SV=1 | 8 | 76 | 2 | 1.85E+07 | 2.07E+07 | 2.27E+07 | 1.96E+07 | 1.95E+07 | 2.05E+07 | 1.66E+07 | 1.49E+07 | 1.02E+07 | 1.08E+07 | 1.49E+07 | 1.11E+07 | 2.03E+07 | 1.31E+07 | 1.55 | 0.63 | 0.000503618 |
| P10909 | Clusterin OS=Homo sapiens OX=9606 GN=CLU PE=1 SV=1 | 8 | 29 | 2 | 6.61E+06 | 4.29E+06 | 2.88E+06 | 2.10E+06 | 4.92E+06 | 4.03E+06 | 7.13E+06 | 6.44E+06 | 5.49E+06 | 3.84E+06 | 3.55E+06 | 1.31E+07 | 4.14E+06 | 6.59E+06 | 0.63 | -0.67 | 0.160151503 |
| P13797 | Plastin-3 OS=Homo sapiens OX=9606 GN=PLS3 PE=1 SV=4 | 9 | 23 | 2 | 2.54E+06 | 2.84E+06 | 4.43E+06 | 4.71E+06 | 1.03E+06 | 1.59E+06 | 2.42E+06 | 1.48E+06 | 1.45E+06 | 3.81E+06 | 4.31E+06 | 1.53E+06 | 2.86E+06 | 2.50E+06 | 1.14 | 0.19 | 0.664528672 |
| Q14699 | Raftlin OS=Homo sapiens OX=9606 GN=RFTN1 PE=1 SV=4 | 6 | 4 | 2 | 2.35E+06 | 4.91E+05 | 1.81E+06 | 1.07E+06 | 1.36E+06 | 1.57E+06 |  | 6.41E+05 | 1.04E+06 | 1.18E+06 | 5.33E+05 | 4.25E+05 | 1.44E+06 | 7.63E+05 | 1.89 | 0.92 | 0.05350748 |
| P51608 | Methyl-CpG-binding protein 2 OS=Homo sapiens OX=9606 GN=MECP2 PE=1 SV=1 | 7 | 26 | 2 | 5.35E+06 | 4.72E+06 | 4.04E+06 | 3.46E+06 | 2.94E+06 | 4.17E+06 | 2.18E+06 | 2.97E+06 | 2.08E+06 | 2.89E+06 | 2.85E+06 | 2.55E+06 | 4.11E+06 | 2.59E+06 | 1.59 | 0.67 | 0.005574675 |
| Q9P2E9 | Ribosome-binding protein 1 OS=Homo sapiens OX=9606 GN=RRBP1 PE=1 SV=5 | 2 | 24 | 2 | 1.99E+06 | 2.82E+06 | 3.23E+06 | 4.05E+06 | 3.20E+06 | 3.47E+06 | 2.02E+06 | 1.18E+06 | 2.15E+06 | 1.39E+06 | 8.80E+05 | 1.13E+06 | 3.13E+06 | 1.46E+06 | 2.14 | 1.10 | 0.000961755 |
| O14817 | Tetraspanin-4 OS=Homo sapiens OX=9606 GN=TSPAN4 PE=1 SV=1 | 14 | 8 | 2 | 2.87E+06 | 2.75E+06 | 2.00E+06 |  | 3.28E+06 | 2.60E+06 | 2.65E+06 | 3.56E+06 | 3.39E+06 |  | 3.05E+06 | 2.58E+06 | 2.70E+06 | 3.05E+06 | 0.89 | -0.17 | 0.25665275 |
| Q92859 | Neogenin OS=Homo sapiens OX=9606 GN=NEO1 PE=1 SV=2 | 2 | 4 | 2 | 2.91E+06 | 3.33E+06 | 1.68E+06 | 3.49E+06 | 2.49E+06 | 2.20E+06 | 3.16E+06 | 4.01E+06 | 3.47E+06 | 3.06E+06 | 2.64E+06 | 3.18E+06 | 2.68E+06 | 3.26E+06 | 0.82 | -0.28 | 0.12702878 |
| O75874 | Isocitrate dehydrogenase [NADP] cytoplasmic OS=Homo sapiens OX=9606 GN=IDH1 PE=1 SV=2 | 8 | 26 | 2 | 3.60E+06 | 3.88E+06 | 4.37E+06 | 3.83E+06 | 3.84E+06 | 5.83E+06 | 4.92E+06 | 4.88E+06 | 3.06E+06 | 4.10E+06 | 5.61E+06 | 3.94E+06 | 4.22E+06 | 4.42E+06 | 0.96 | -0.06 | 0.706768167 |
| Q9NVJ2 | ADP-ribosylation factor-like protein 8B OS=Homo sapiens OX=9606 GN=ARL8B PE=1 SV=1 | 14 | 14 | 2 | 3.93E+06 | 6.29E+06 | 4.90E+06 | 4.42E+06 | 3.13E+06 | 6.22E+06 | 3.00E+06 | 3.34E+06 | 2.76E+06 | 2.44E+06 | 3.78E+06 | 2.79E+06 | 4.82E+06 | 3.02E+06 | 1.60 | 0.67 | 0.015670394 |

### Supplementary Table 4. List of all proteins identified by label free proteomics analysis.

| Accession | Description | Coverage [%] | # PSMs | # Unique Peptides | Amitriptyline 1 | Amitriptyline 2 | Amitriptyline 3 | Amitriptyline 4 | Amitriptyline 5 | Amitriptyline 6 | Vehicle 1 | Vehicle 2 | Vehicle 3 | Vehicle 4 | Vehicle 5 | Vehicle 6 | Average amitriptyline | Average vehicle | Ratio amitriptyline/ vehicle | Log 2 fold change (amitriptyline /vehicle) | P-value |
|---|---|---|---|---|---|---|---|---|---|---|---|---|---|---|---|---|---|---|---|---|---|
| P08237 | ATP-dependent 6-phosphofructokinase, muscle type OS=Homo sapiens OX=9606 GN=PFKM PE=1 SV=2 | 4 | 31 | 2 | 3.36E+06 | 3.55E+06 | 3.32E+06 | 3.58E+06 | 3.29E+06 | 2.64E+06 | 3.18E+06 | 3.14E+06 | 3.20E+06 | 3.29E+06 | 3.08E+06 | 2.54E+06 | 3.29E+06 | 3.07E+06 | 1.07 | 0.10 | 0.248096493 |
| P49888 | Sulfotransferase 1E1 OS=Homo sapiens OX=9606 GN=SULT1E1 PE=1 SV=1 | 6 | 70 | 2 | 1.12E+07 | 1.10E+07 | 2.36E+07 | 2.45E+07 | 2.68E+07 | 1.84E+07 | 2.86E+07 | 2.99E+07 | 4.45E+07 | 3.68E+07 | 2.91E+07 | 2.73E+07 | 1.92E+07 | 3.27E+07 | 0.59 | -0.76 | 0.006451522 |
| Q92974 | Rho guanine nucleotide exchange factor 2 OS=Homo sapiens OX=9606 GN=ARHGEF2 PE=1 SV=4 | 3 | 6 | 2 | | 4.36E+05 | 1.83E+06 | 1.48E+06 | 9.46E+05 | 2.16E+06 | 1.24E+06 | 3.49E+05 | 6.85E+05 | 4.71E+05 | 3.60E+05 | 7.24E+05 | 1.37E+06 | 6.38E+05 | 2.15 | 1.10 | 0.076597122 |
| O60664 | Perilipin-3 OS=Homo sapiens OX=9606 GN=PLIN3 PE=1 SV=3 | 6 | 18 | 2 | 2.96E+06 | 2.74E+06 | 2.63E+06 | 3.40E+06 | 2.92E+06 | 2.30E+06 | 2.02E+06 | 1.29E+06 | 9.25E+05 | 1.41E+06 | 1.40E+06 | 1.20E+06 | 2.82E+06 | 1.37E+06 | 2.06 | 1.04 | 4.33151E-05 |
| P23468 | Receptor-type tyrosine-protein phosphatase delta OS=Homo sapiens OX=9606 GN=PTPRD PE=1 SV=2 | 1 | 26 | 2 | 2.83E+06 | 2.39E+06 | 3.21E+06 | 2.65E+06 | 2.95E+06 | 1.93E+06 | 3.83E+06 | 3.83E+06 | 3.17E+06 | 2.58E+06 | 3.22E+06 | 2.98E+06 | 2.66E+06 | 3.27E+06 | 0.81 | -0.30 | 0.049632035 |
| O95297 | Myelin protein zero-like protein 1 OS=Homo sapiens OX=9606 GN=MPZL1 PE=1 SV=1 | 10 | 18 | 2 | 2.53E+06 | 2.47E+06 | 3.28E+06 | 3.67E+06 | 3.31E+06 | 2.95E+06 | 1.31E+06 | 1.33E+06 | 1.75E+06 | 1.40E+06 | 1.09E+06 | 1.34E+06 | 3.03E+06 | 1.37E+06 | 2.21 | 1.15 | 0.000108535 |
| P60866 | 40S ribosomal protein S20 OS=Homo sapiens OX=9606 GN=RPS20 PE=1 SV=1 | 19 | 54 | 2 | 1.83E+07 | 1.75E+07 | 1.57E+07 | 1.92E+07 | 1.64E+07 | 1.70E+07 | 1.26E+07 | 1.42E+07 | 1.47E+07 | 1.10E+07 | 1.11E+07 | 1.60E+07 | 1.73E+07 | 1.33E+07 | 1.31 | 0.39 | 0.002775952 |
| P52272 | Heterogeneous nuclear ribonucleoprotein M OS=Homo sapiens OX=9606 GN=HNRNPM PE=1 SV=3 | 3 | 34 | 2 | 1.05E+07 | 1.20E+07 | 7.04E+06 | 8.88E+06 | 6.65E+06 | 7.03E+06 | 6.09E+06 | 5.54E+06 | 3.63E+06 | 3.48E+06 | 3.90E+06 | 3.82E+06 | 8.68E+06 | 4.41E+06 | 1.97 | 0.98 | 0.003213492 |
| Q16836 | Hydroxyacyl-coenzyme A dehydrogenase, mitochondrial OS=Homo sapiens OX=9606 GN=HADH PE=1 SV=3 | 10 | 22 | 2 | 3.70E+06 | 4.23E+06 | 4.87E+06 | 2.96E+06 | 4.27E+06 | 4.96E+06 | 3.84E+06 | 4.09E+06 | 4.18E+06 | 4.14E+06 | 4.20E+06 | 4.48E+06 | 4.16E+06 | 4.15E+06 | 1.00 | 0.00 | 0.978286341 |
| P13489 | Ribonuclease inhibitor OS=Homo sapiens OX=9606 GN=RNH1 PE=1 SV=2 | 6 | 12 | 2 | 3.03E+06 | 3.61E+06 | 2.81E+06 | 3.12E+06 | 3.87E+06 | 2.69E+06 | 9.20E+05 | 1.65E+06 | 1.67E+06 | 1.90E+06 | 9.40E+05 | 1.42E+06 | 3.19E+06 | 1.42E+06 | 2.25 | 1.17 | 3.7909E-05 |
| Q14444 | Caprin-1 OS=Homo sapiens OX=9606 GN=CAPRIN1 PE=1 SV=2 | 5 | 8 | 2 | 2.74E+06 | 1.99E+06 | 3.38E+06 | 2.83E+06 | 3.38E+06 | 3.04E+06 | 2.46E+06 | 1.79E+06 | 2.11E+06 | 1.20E+06 | 1.91E+06 | 1.54E+06 | 2.89E+06 | 1.83E+06 | 1.58 | 0.66 | 0.003485861 |
| P35637 | RNA-binding protein FUS OS=Homo sapiens OX=9606 GN=FUS PE=1 SV=1 | 6 | 14 | 2 | 5.74E+06 | 5.13E+06 | 4.98E+06 | 4.81E+06 | 4.08E+06 | 3.61E+06 | 2.27E+06 | 1.56E+06 | 1.70E+06 | 1.85E+06 | 1.76E+06 | 1.12E+06 | 4.72E+06 | 1.71E+06 | 2.76 | 1.47 | 4.28385E-05 |
| Q7RTV2 | Glutathione S-transferase A5 OS=Homo sapiens OX=9606 GN=GSTA5 PE=1 SV=1 | 7 | 270 | 2 | 5.97E+07 | 5.78E+07 | 5.22E+07 | 5.68E+07 | 5.27E+07 | 4.54E+07 | 7.24E+07 | 6.94E+07 | 7.65E+07 | 8.07E+07 | 6.28E+07 | 4.92E+07 | 5.41E+07 | 6.85E+07 | 0.79 | -0.34 | 0.024471466 |
| Q8TBC4 | NEDD8-activating enzyme E1 catalytic subunit OS=Homo sapiens OX=9606 GN=UBA3 PE=1 SV=2 | 9 | 13 | 2 | | 1.03E+06 | 7.96E+05 | 3.26E+06 | 8.38E+05 | 2.29E+06 | 5.20E+05 | 6.24E+05 | 1.75E+06 | 1.76E+06 | 3.05E+06 | 1.33E+06 | 1.64E+06 | 1.51E+06 | 1.09 | 0.12 | 0.831342996 |
| P61966 | AP-1 complex subunit sigma-1A OS=Homo sapiens OX=9606 GN=AP1S1 PE=1 SV=1 | 15 | 5 | 2 | | | | | | | | | | | | | | | | | |
| Q7L2E3 | ATP-dependent RNA helicase DHX30 OS=Homo sapiens OX=9606 GN=DHX30 PE=1 SV=1 | 4 | 4 | 2 | | 3.88E+05 | 1.56E+06 | 2.77E+05 | 1.44E+06 | 8.54E+05 | 1.02E+06 | 3.29E+05 | 4.23E+05 | 3.67E+05 | 5.17E+05 | 5.50E+05 | 9.04E+05 | 5.35E+05 | 1.69 | 0.76 | 0.245743931 |
| O14879 | Interferon-induced protein with tetratricopeptide repeats 3 OS=Homo sapiens OX=9606 GN=IFIT3 PE=1 SV=1 | 5 | 6 | 2 | | 1.43E+06 | 1.18E+06 | 1.58E+06 | 2.09E+06 | 1.35E+06 | | | 5.32E+05 | | | 9.50E+05 | 1.53E+06 | 7.41E+05 | 2.06 | 1.04 | 0.082308324 |
| P46060 | Ran GTPase-activating protein 1 OS=Homo sapiens OX=9606 GN=RANGAP1 PE=1 SV=1 | 4 | 4 | 2 | 2.40E+06 | 3.19E+06 | 1.62E+06 | 2.89E+06 | 2.75E+06 | 1.33E+06 | 1.96E+06 | 6.28E+05 | 1.63E+06 | 1.49E+06 | 9.12E+05 | 5.74E+05 | 2.37E+06 | 1.20E+06 | 1.97 | 0.98 | 0.012914068 |
| P35244 | Replication protein A 14 kDa subunit OS=Homo sapiens OX=9606 GN=RPA3 PE=1 SV=1 | 30 | 13 | 2 | 4.84E+06 | 2.05E+06 | 1.42E+06 | 1.85E+06 | 1.62E+06 | 2.12E+06 | 2.84E+06 | 1.86E+06 | 1.48E+06 | 1.58E+06 | 1.56E+06 | 1.36E+06 | 2.31E+06 | 1.78E+06 | 1.30 | 0.38 | 0.373051958 |
| P11233 | Ras-related protein Ral-A OS=Homo sapiens OX=9606 GN=RALA PE=1 SV=1 | 19 | 8 | 2 | 1.40E+06 | 1.95E+06 | 2.30E+06 | 2.06E+06 | 1.39E+06 | 2.13E+06 | 1.34E+06 | 1.04E+06 | 7.84E+05 | | 1.83E+06 | 6.05E+05 | 1.87E+06 | 1.12E+06 | 1.67 | 0.74 | 0.024126592 |
| P46783 | 40S ribosomal protein S10 OS=Homo sapiens OX=9606 GN=RPS10 PE=1 SV=1 | 15 | 58 | 2 | 1.40E+07 | 9.03E+06 | 5.81E+06 | 1.05E+07 | 6.21E+06 | 1.02E+07 | 1.30E+07 | 5.92E+06 | 6.69E+06 | 3.93E+06 | 5.19E+06 | 5.44E+06 | 9.30E+06 | 6.69E+06 | 1.39 | 0.47 | 0.180156815 |
| O75821 | Eukaryotic translation initiation factor 3 subunit G OS=Homo sapiens OX=9606 GN=EIF3G PE=1 SV=2 | 13 | 6 | 2 | 1.42E+06 | 5.93E+05 | 1.07E+06 | 1.53E+06 | 1.20E+06 | 1.87E+06 | 7.54E+05 | 4.96E+05 | 6.39E+05 | 3.85E+05 | 6.48E+05 | 6.40E+05 | 1.28E+06 | 5.94E+05 | 2.16 | 1.11 | 0.01045088 |
| Q99439 | Calponin-2 OS=Homo sapiens OX=9606 GN=CNN2 PE=1 SV=4 | 13 | 25 | 2 | 7.21E+06 | 6.48E+06 | 6.20E+06 | 6.41E+06 | 6.43E+06 | 5.39E+06 | 7.33E+06 | 3.22E+06 | 3.80E+06 | 3.37E+06 | 4.59E+06 | 4.59E+06 | 6.35E+06 | 4.48E+06 | 1.42 | 0.50 | 0.028127179 |
| P22694 | cAMP-dependent protein kinase catalytic subunit beta OS=Homo sapiens OX=9606 GN=PRKACB PE=1 SV=2 | 9 | 10 | 2 | 2.68E+06 | 2.59E+06 | 2.14E+06 | 1.39E+06 | 1.04E+06 | 1.82E+06 | 3.33E+06 | 3.16E+06 | 3.04E+06 | 1.56E+06 | 3.15E+06 | 1.97E+06 | 1.95E+06 | 2.70E+06 | 0.72 | -0.47 | 0.091664469 |
| P42167 | Lamina-associated polypeptide 2, isoforms beta/gamma OS=Homo sapiens OX=9606 GN=TMPO PE=1 SV=2 | 9 | 12 | 2 | 1.81E+06 | 4.74E+06 | 4.73E+06 | 5.04E+06 | 1.91E+06 | 1.74E+06 | 4.04E+06 | 4.56E+06 | 9.86E+05 | 4.55E+06 | 9.12E+05 | 3.37E+06 | 3.33E+06 | 3.07E+06 | 1.08 | 0.12 | 0.795342912 |
| P62847 | 40S ribosomal protein S24 OS=Homo sapiens OX=9606 GN=RPS24 PE=1 SV=1 | 20 | 53 | 2 | 1.47E+07 | 1.35E+07 | 8.13E+06 | 1.12E+07 | 1.06E+07 | 1.10E+07 | 8.79E+06 | 1.03E+07 | 4.09E+06 | 4.07E+06 | 6.75E+06 | 5.43E+06 | 1.15E+07 | 6.57E+06 | 1.75 | 0.81 | 0.005679177 |
| Q12805 | EGF-containing fibulin-like extracellular matrix protein 1 OS=Homo sapiens OX=9606 GN=EFEMP1 PE=1 SV=2 | 7 | 38 | 2 | 6.75E+06 | 5.72E+06 | 5.52E+06 | 5.18E+06 | 5.93E+06 | 5.52E+06 | 9.72E+06 | 7.99E+06 | 9.32E+06 | 7.23E+06 | 8.43E+06 | 7.68E+06 | 5.77E+06 | 8.39E+06 | 0.69 | -0.54 | 0.000419632 |
| O75436 | Vacuolar protein sorting-associated protein 26A OS=Homo sapiens OX=9606 GN=VPS26A PE=1 SV=2 | 9 | 7 | 2 | 2.20E+06 | 3.25E+06 | 2.25E+06 | 2.56E+06 | 3.64E+06 | 2.60E+06 | 1.27E+06 | | 5.58E+05 | 1.91E+06 | 4.91E+05 | 1.57E+06 | 2.75E+06 | 1.16E+06 | 2.37 | 1.25 | 0.002173935 |
| Q9UHB6 | LIM domain and actin-binding protein 1 OS=Homo sapiens OX=9606 GN=LIMA1 PE=1 SV=1 | 3 | 8 | 2 | 1.71E+06 | 3.05E+06 | 3.19E+06 | 3.00E+06 | 2.93E+06 | 2.57E+06 | | | 1.16E+06 | 1.03E+06 | 1.13E+06 | 8.99E+05 | 2.74E+06 | 1.05E+06 | 2.61 | 1.38 | 0.000417889 |
| Q15637 | Splicing factor 1 OS=Homo sapiens OX=9606 GN=SF1 PE=1 SV=4 | 5 | 10 | 2 | | 4.52E+05 | 2.67E+06 | 6.63E+05 | 1.91E+06 | 9.01E+05 | 3.24E+05 | 5.89E+05 | 4.22E+05 | 1.23E+06 | 8.76E+05 | 5.05E+05 | 1.32E+06 | 6.58E+05 | 2.01 | 1.00 | 0.196465508 |
| P01116 | GTPase KRas OS=Homo sapiens OX=9606 GN=KRAS PE=1 SV=1 | 20 | 4 | 2 | 5.94E+06 | 5.75E+06 | 3.17E+06 | 3.74E+06 | 3.16E+06 | 6.39E+06 | 4.13E+06 | 5.64E+06 | 2.71E+06 | 3.31E+06 | 2.80E+06 | 3.38E+06 | 4.69E+06 | 3.66E+06 | 1.28 | 0.36 | 0.205712945 |
| Q9Y2V2 | Calcium-regulated heat-stable protein 1 OS=Homo sapiens OX=9606 GN=CARHSP1 PE=1 SV=2 | 18 | 33 | 2 | 3.58E+06 | 5.70E+06 | 5.02E+06 | 6.87E+06 | 5.12E+06 | 4.47E+06 | 4.94E+06 | 5.38E+06 | 4.99E+06 | 3.25E+06 | 5.12E+06 | 4.48E+06 | 5.13E+06 | 4.69E+06 | 1.09 | 0.13 | 0.451481331 |
| P62306 | Small nuclear ribonucleoprotein F OS=Homo sapiens OX=9606 GN=SNRPF PE=1 SV=1 | 24 | 28 | 2 | 6.90E+06 | 5.69E+06 | 4.66E+06 | 5.54E+06 | 4.52E+06 | 5.70E+06 | 4.08E+06 | 4.06E+06 | 3.17E+06 | 4.53E+06 | 3.65E+06 | 2.24E+06 | 5.50E+06 | 3.62E+06 | 1.52 | 0.60 | 0.003112069 |
| O95747 | Serine/threonine-protein kinase OSR1 OS=Homo sapiens OX=9606 GN=OXSR1 PE=1 SV=1 | 8 | 3 | 2 | | | | 2.38E+05 | | 8.58E+05 | 3.47E+05 | | | | | | 5.48E+05 | 3.47E+05 | 1.58 | 0.66 | |
| O43149 | Zinc finger ZZ-type and EF-hand domain-containing protein 1 OS=Homo sapiens OX=9606 GN=ZZEF1 PE=1 SV=6 | 1 | 8 | 2 | 4.25E+06 | 4.57E+06 | | | | | 9.68E+05 | | 6.27E+06 | 4.77E+06 | | | 4.41E+06 | 4.00E+06 | 1.10 | 0.14 | 0.821824898 |
| P35080 | Profilin-2 OS=Homo sapiens OX=9606 GN=PFN2 PE=1 SV=3 | 27 | 8 | 2 | 2.41E+06 | 3.90E+05 | 2.63E+06 | 1.91E+06 | 1.89E+06 | 1.48E+06 | 5.44E+05 | 1.79E+06 | 1.36E+06 | 5.26E+05 | 1.87E+06 | | 1.78E+06 | 1.22E+06 | 1.46 | 0.55 | 0.228073902 |
| P23919 | Thymidylate kinase OS=Homo sapiens OX=9606 GN=DTYMK PE=1 SV=4 | 11 | 6 | 2 | 2.94E+06 | 4.06E+06 | 2.44E+06 | 3.19E+06 | 1.38E+06 | 2.23E+06 | 8.28E+05 | 9.57E+05 | 7.09E+05 | 1.12E+06 | 9.84E+05 | | 2.71E+06 | 9.19E+05 | 2.95 | 1.56 | 0.004428635 |
| P55735 | Protein SEC13 homolog OS=Homo sapiens OX=9606 GN=SEC13 PE=1 SV=3 | 7 | 25 | 2 | 5.31E+06 | 4.51E+06 | 5.39E+06 | 5.92E+06 | 5.09E+06 | 4.94E+06 | 3.58E+06 | 2.43E+06 | 3.88E+06 | 3.48E+06 | 3.24E+06 | 2.83E+06 | 5.20E+06 | 3.24E+06 | 1.60 | 0.68 | 5.55916E-05 |
| P61077 | Ubiquitin-conjugating enzyme E2 D3 OS=Homo sapiens OX=9606 GN=UBE2D3 PE=1 SV=1 | 20 | 13 | 2 | 6.07E+06 | 7.24E+06 | 3.40E+06 | 6.96E+06 | 3.47E+06 | 4.51E+06 | 3.16E+06 | 5.01E+06 | 1.23E+06 | 1.97E+06 | 3.62E+06 | 2.70E+06 | 5.27E+06 | 2.95E+06 | 1.79 | 0.84 | 0.02630549 |
| Q04828 | Aldo-keto reductase family 1 member C1 OS=Homo sapiens OX=9606 GN=AKR1C1 PE=1 SV=1 | 6 | 53 | 2 | 2.32E+07 | 2.60E+07 | 2.85E+07 | 3.19E+07 | 2.89E+07 | 2.66E+07 | 3.91E+07 | 4.12E+07 | 4.39E+07 | 4.26E+07 | 3.83E+07 | 3.38E+07 | 2.75E+07 | 3.98E+07 | 0.69 | -0.53 | 8.91519E-05 |
| P61970 | Nuclear transport factor 2 OS=Homo sapiens OX=9606 GN=NUTF2 PE=1 SV=1 | 28 | 18 | 2 | 8.46E+06 | 7.74E+06 | 7.80E+06 | 1.19E+07 | 8.57E+06 | 8.73E+06 | 6.61E+06 | 8.30E+06 | 8.46E+06 | 8.39E+06 | 9.86E+06 | 7.82E+06 | 8.86E+06 | 8.24E+06 | 1.08 | 0.11 | 0.433027879 |
| Q9UBI6 | Guanine nucleotide-binding protein G(I)/G(S)/G(O) subunit gamma-12 OS=Homo sapiens OX=9606 GN=GNG12 PE=1 SV=3 | 32 | 24 | 2 | 4.59E+06 | 3.99E+06 | 3.58E+06 | 4.50E+06 | 3.76E+06 | 4.63E+06 | 2.32E+06 | 3.57E+06 | 3.32E+06 | 3.42E+06 | 2.77E+06 | 2.48E+06 | 4.18E+06 | 2.98E+06 | 1.40 | 0.49 | 0.001910277 |
| P11245 | Arylamine N-acetyltransferase 2 OS=Homo sapiens OX=9606 GN=NAT2 PE=1 SV=1 | 9 | 18 | 2 | 6.90E+06 | 3.30E+06 | 6.59E+06 | 7.16E+06 | 5.92E+06 | 5.19E+06 | 1.23E+07 | 6.10E+06 | 7.96E+06 | 9.15E+06 | 6.90E+06 | 6.13E+06 | 5.84E+06 | 8.09E+06 | 0.72 | -0.47 | 0.081767135 |
| Q9UN36 | Protein NDRG2 OS=Homo sapiens OX=9606 GN=NDRG2 PE=1 SV=2 | 7 | 19 | 2 | 2.13E+06 | 2.60E+06 | 3.21E+06 | 3.78E+06 | 1.98E+06 | 3.26E+06 | 5.65E+06 | 5.04E+06 | 5.54E+06 | 5.37E+06 | 5.14E+06 | 2.86E+06 | 2.83E+06 | 4.93E+06 | 0.57 | -0.80 | 0.002814837 |
| P48960 | CD97 antigen OS=Homo sapiens OX=9606 GN=CD97 PE=1 SV=4 | 4 | 15 | 2 | 3.11E+06 | 2.37E+06 | 1.90E+06 | 2.29E+06 | 2.48E+06 | 2.87E+06 | 1.30E+06 | 9.24E+05 | | | 1.12E+06 | | 2.50E+06 | 1.11E+06 | 2.25 | 1.17 | 0.000277016 |
| Q9Y383 | Putative RNA-binding protein Luc7-like 2 OS=Homo sapiens OX=9606 GN=LUC7L2 PE=1 SV=2 | 6 | 6 | 2 | 7.96E+05 | 9.21E+05 | 1.14E+06 | 1.45E+06 | 1.35E+06 | 1.06E+06 | 1.00E+06 | 6.95E+05 | 7.99E+05 | 1.31E+06 | 7.97E+05 | 6.12E+05 | 1.12E+06 | 8.68E+05 | 1.29 | 0.37 | 0.11362792 |
| P02788 | Lactotransferrin OS=Homo sapiens OX=9606 GN=LTF PE=1 SV=6 | 3 | 122 | 2 | 2.54E+07 | 2.56E+07 | 2.74E+07 | 3.06E+07 | 2.61E+07 | 3.00E+07 | 6.43E+07 | 6.73E+07 | 7.68E+07 | 6.72E+07 | 7.56E+07 | 6.82E+07 | 2.75E+07 | 6.99E+07 | 0.39 | -1.34 | 3.44529E-07 |
| O75531 | Barrier-to-autointegration factor OS=Homo sapiens OX=9606 GN=BANF1 PE=1 SV=1 | 40 | 6 | 2 | 2.14E+06 | 3.80E+06 | 1.96E+06 | 1.73E+06 | 2.08E+06 | 2.98E+06 | 2.13E+06 | 2.88E+06 | 1.99E+06 | 1.03E+06 | 3.16E+06 | 2.03E+06 | 2.45E+06 | 2.20E+06 | 1.11 | 0.15 | 0.592104179 |
| Q7Z4W1 | L-xylulose reductase OS=Homo sapiens OX=9606 GN=DCXR PE=1 SV=2 | 9 | 66 | 2 | 2.28E+07 | 2.69E+07 | 2.01E+07 | 2.15E+07 | 2.00E+07 | 2.04E+07 | 3.37E+07 | 3.36E+07 | 2.52E+07 | 2.79E+07 | 3.69E+07 | 2.67E+07 | 2.20E+07 | 3.07E+07 | 0.72 | -0.48 | 0.004278787 |
| P08603 | Complement factor H OS=Homo sapiens OX=9606 GN=CFH PE=1 SV=4 | 2 | 4 | 2 | | | | | | | | | | | | 2.09E+06 | | | | | |
| P06727 | Apolipoprotein A-IV OS=Homo sapiens OX=9606 GN=APOA4 PE=1 SV=3 | 6 | 44 | 2 | 4.21E+06 | 4.04E+06 | 4.54E+06 | 6.71E+06 | 5.07E+06 | 5.06E+06 | 6.30E+06 | 8.30E+06 | 8.69E+06 | 7.81E+06 | 6.53E+06 | 6.60E+06 | 4.94E+06 | 7.37E+06 | 0.67 | -0.58 | 0.001739952 |
| P19525 | Interferon-induced, double-stranded RNA-activated protein kinase OS=Homo sapiens OX=9606 GN=EIF2AK2 PE=1 SV=2 | 4 | 4 | 2 | 2.21E+06 | 2.34E+06 | 2.64E+06 | 2.95E+06 | 2.56E+06 | 2.93E+06 | 6.32E+05 | 8.45E+05 | 5.70E+05 | 1.30E+06 | 4.45E+05 | 8.84E+05 | 2.60E+06 | 7.79E+05 | 3.35 | 1.74 | 1.01492E-06 |
| Q00796 | Sorbitol dehydrogenase OS=Homo sapiens OX=9606 GN=SORD PE=1 SV=4 | 6 | 38 | 2 | 8.68E+06 | 9.75E+06 | 1.12E+07 | 1.06E+07 | 9.75E+06 | 9.22E+06 | 1.33E+07 | 1.29E+07 | 1.38E+07 | 1.31E+07 | 1.05E+07 | 1.08E+07 | 9.87E+06 | 1.24E+07 | 0.80 | -0.33 | 0.004862077 |
| P08183 | ATP-dependent translocase ABCB1 OS=Homo sapiens OX=9606 GN=ABCB1 PE=1 SV=3 | 3 | 10 | 2 | 1.00E+06 | 2.49E+06 | 7.25E+05 | 1.28E+06 | | 2.06E+06 | 8.01E+05 | 6.22E+05 | 1.23E+06 | 2.11E+06 | | 5.00E+05 | 1.51E+06 | 1.05E+06 | 1.43 | 0.52 | 0.331419497 |
| Q9BPW8 | Protein NipSnap homolog 1 OS=Homo sapiens OX=9606 GN=NIPSNAP1 PE=1 SV=1 | 13 | 9 | 2 | 3.97E+06 | 3.39E+06 | 1.64E+06 | 2.18E+06 | 9.27E+05 | 2.77E+06 | | 2.24E+06 | 1.01E+06 | 1.38E+06 | 8.77E+05 | 2.14E+06 | 2.48E+06 | 1.53E+06 | 1.62 | 0.69 | 0.117038791 |
| P09455 | Retinol-binding protein 1 OS=Homo sapiens OX=9606 GN=RBP1 PE=1 SV=2 | 14 | 19 | 2 | 4.84E+06 | 5.09E+06 | 4.06E+06 | 5.06E+06 | 3.31E+06 | 4.20E+06 | 4.87E+06 | 7.70E+06 | 2.18E+06 | 3.56E+06 | 3.04E+06 | 3.93E+06 | 4.43E+06 | 4.21E+06 | 1.05 | 0.07 | 0.80855918 |
| P48723 | Heat shock 70 kDa protein 13 OS=Homo sapiens OX=9606 GN=HSPA13 PE=1 SV=1 | 5 | 18 | 2 | 2.93E+06 | 2.64E+06 | 2.47E+06 | 2.83E+06 | 2.49E+06 | 2.11E+06 | 3.52E+06 | 2.93E+06 | 3.38E+06 | 3.44E+06 | 2.98E+06 | 2.50E+06 | 2.58E+06 | 3.13E+06 | 0.82 | -0.28 | 0.022127497 |
| P08236 | Beta-glucuronidase OS=Homo sapiens OX=9606 GN=GUSB PE=1 SV=2 | 4 | 14 | 2 | 3.52E+06 | 3.21E+06 | 4.11E+06 | 3.52E+06 | 4.38E+06 | 1.77E+06 | 8.65E+05 | 8.62E+05 | | 4.56E+05 | 1.19E+06 | 9.47E+05 | 3.42E+06 | 8.64E+05 | 3.96 | 1.98 | 0.00063788 |
| P28838 | Cytosol aminopeptidase OS=Homo sapiens OX=9606 GN=LAP3 PE=1 SV=3 | 5 | 14 | 2 | 2.97E+06 | 2.06E+06 | 2.65E+06 | 3.45E+06 | 2.63E+06 | 2.18E+06 | 1.56E+06 | 4.87E+06 | 1.57E+06 | 3.13E+06 | 3.12E+06 | 2.11E+06 | 2.66E+06 | 2.73E+06 | 0.97 | -0.04 | 0.904474887 |
| Q06323 | Proteasome activator complex subunit 1 OS=Homo sapiens OX=9606 GN=PSME1 PE=1 SV=1 | 8 | 11 | 2 | 3.15E+06 | 3.02E+06 | 3.58E+06 | 2.95E+06 | 3.45E+06 | 3.14E+06 | 8.69E+05 | 9.79E+05 | 9.76E+05 | 2.07E+06 | 1.25E+06 | | 3.22E+06 | 1.23E+06 | 2.62 | 1.39 | 0.00023719 |
| Q8N1G4 | Leucine-rich repeat-containing protein 47 OS=Homo sapiens OX=9606 GN=LRRC47 PE=1 SV=1 | 4 | 4 | 2 | 2.66E+06 | 2.54E+06 | 3.20E+06 | 3.10E+06 | 3.19E+06 | 1.18E+06 | 4.57E+05 | 1.60E+06 | 1.66E+06 | 1.38E+06 | | 1.03E+06 | 2.64E+06 | 1.23E+06 | 2.16 | 1.11 | 0.005533421 |
| P30825 | High affinity cationic amino acid transporter 1 OS=Homo sapiens OX=9606 GN=SLC7A1 PE=1 SV=1 | 4 | 6 | 2 | 8.61E+05 | 1.56E+06 | 1.75E+06 | 2.34E+06 | 2.62E+06 | 1.12E+06 | 8.21E+05 | 7.94E+05 | 1.17E+06 | 8.02E+05 | 6.91E+05 | 5.73E+05 | 1.71E+06 | 8.09E+05 | 2.11 | 1.08 | 0.021586988 |
| P26583 | High mobility group protein B2 OS=Homo sapiens OX=9606 GN=HMGB2 PE=1 SV=2 | 14 | 24 | 2 | 3.40E+06 | 3.29E+06 | 3.53E+06 | 4.45E+06 | 3.25E+06 | 2.88E+06 | 3.81E+06 | 3.89E+06 | 3.65E+06 | 3.77E+06 | 4.29E+06 | 3.04E+06 | 3.47E+06 | 3.74E+06 | 0.93 | -0.11 | 0.332486326 |
| P23284 | Peptidyl-prolyl cis-trans isomerase B OS=Homo sapiens OX=9606 GN=PPIB PE=1 SV=2 | 12 | 31 | 2 | 6.58E+06 | 4.97E+06 | 5.85E+06 | 3.87E+06 | 5.19E+06 | 4.59E+06 | 4.40E+06 | 1.96E+06 | 4.36E+06 | 3.27E+06 | 2.63E+06 | 3.33E+06 | 5.17E+06 | 3.32E+06 | 1.56 | 0.64 | 0.00721025 |
| Q9H2H9 | Sodium-coupled neutral amino acid transporter 1 OS=Homo sapiens OX=9606 GN=SLC38A1 PE=1 SV=1 | 4 | 24 | 2 | 4.85E+06 | 6.02E+06 | 6.52E+06 | 6.73E+06 | 8.38E+06 | 5.55E+06 | 3.48E+06 | 2.18E+06 | 1.38E+06 | 3.40E+06 | 3.13E+06 | 2.93E+06 | 6.34E+06 | 2.75E+06 | 2.31 | 1.21 | 0.000214129 |
| Q9Y3Z3 | Deoxynucleoside triphosphate triphosphohydrolase SAMHD1 OS=Homo sapiens OX=9606 GN=SAMHD1 PE=1 SV=2 | 4 | 24 | 2 | 2.51E+06 | 3.02E+06 | 4.35E+06 | 4.30E+06 | 2.52E+06 | 2.18E+06 | 3.35E+06 | 3.10E+06 | 1.83E+06 | 2.21E+06 | 1.94E+06 | 2.49E+06 | 3.15E+06 | 2.49E+06 | 1.26 | 0.34 | 0.191912525 |
| P45877 | Peptidyl-prolyl cis-trans isomerase C OS=Homo sapiens OX=9606 GN=PPIC PE=1 SV=1 | 10 | 32 | 2 | 4.38E+06 | 6.43E+06 | 3.27E+06 | 4.40E+06 | 3.70E+06 | 3.77E+06 | 7.85E+06 | 1.04E+07 | 5.31E+06 | 5.18E+06 | 5.22E+06 | 4.37E+06 | 4.33E+06 | 6.39E+06 | 0.68 | -0.56 | 0.086702087 |
| P55769 | NHP2-like protein 1 OS=Homo sapiens OX=9606 GN=SNU13 PE=1 SV=3 | 16 | 29 | 2 | 3.02E+06 | 3.52E+06 | 4.35E+06 | 5.34E+06 | 4.13E+06 | 3.61E+06 | 3.98E+06 | 3.92E+06 | 3.42E+06 | 3.24E+06 | 2.14E+06 | 3.22E+06 | 4.00E+06 | 3.32E+06 | 1.20 | 0.27 | 0.144910012 |
| P61201 | COP9 signalosome complex subunit 2 OS=Homo sapiens OX=9606 GN=COPS2 PE=1 SV=1 | 5 | 12 | 2 | 5.65E+05 | 1.21E+06 | 2.01E+06 | 1.18E+06 | 1.49E+06 | 1.33E+06 | 9.50E+05 | 1.24E+06 | 1.50E+06 | 8.85E+05 | 9.75E+05 | 8.74E+05 | 1.30E+06 | 1.07E+06 | 1.21 | 0.28 | 0.325838248 |
| P48509 | CD151 antigen OS=Homo sapiens OX=9606 GN=CD151 PE=1 SV=3 | 8 | 30 | 2 | 1.37E+07 | 1.24E+07 | 1.22E+07 | 1.28E+07 | 1.33E+07 | 1.98E+07 | 8.62E+06 | 7.80E+06 | 9.69E+06 | 9.74E+06 | 9.31E+06 | 6.86E+06 | 1.40E+07 | 8.67E+06 | 1.62 | 0.69 | 0.004306027 |
| Q14160 | Protein scribble homolog OS=Homo sapiens OX=9606 GN=SCRIB PE=1 SV=4 | 2 | 12 | 2 | 2.59E+08 | 2.91E+08 | 2.61E+08 | 1.92E+08 | 1.82E+08 | 2.63E+08 | 1.95E+08 | 2.75E+08 | 2.03E+08 | 2.11E+08 | 2.50E+08 | 2.38E+08 | 2.41E+08 | 2.29E+08 | 1.06 | 0.08 | 0.577298634 |
| P00742 | Coagulation factor X OS=Homo sapiens OX=9606 GN=F10 PE=1 SV=2 | 6 | 42 | 2 | 7.75E+06 | 8.56E+06 | 8.00E+06 | 5.52E+06 | 6.97E+06 | 8.48E+06 | 1.15E+07 | 9.68E+06 | 1.10E+07 | 7.97E+06 | 1.16E+07 | 5.60E+06 | 7.55E+06 | 9.56E+06 | 0.79 | -0.34 | 0.102753823 |
| Q08AM6 | Protein VAC14 homolog OS=Homo sapiens OX=9606 GN=VAC14 PE=1 SV=1 | 4 | 14 | 2 | 2.20E+06 | 2.13E+06 | 2.42E+06 | 2.53E+06 | 2.05E+06 | 2.37E+06 | | | 1.16E+06 | 1.08E+06 | 9.34E+05 | 6.96E+05 | 2.29E+06 | 9.67E+05 | 2.36 | 1.24 | 4.15152E-05 |
| Q04206 | Transcription factor p65 OS=Homo sapiens OX=9606 GN=RELA PE=1 SV=2 | 4 | 9 | 2 | 3.38E+06 | 4.00E+06 | 3.77E+06 | 2.78E+06 | 3.24E+06 | 4.47E+06 | 9.47E+05 | 2.56E+06 | 1.02E+06 | 2.18E+06 | 1.91E+06 | 9.59E+05 | 3.61E+06 | 1.60E+06 | 2.26 | 1.18 | 0.000386348 |
| P25685 | DnaJ homolog subfamily B member 1 OS=Homo sapiens OX=9606 GN=DNAJB1 PE=1 SV=4 | 6 | 30 | 2 | 4.40E+06 | 4.34E+06 | 4.34E+06 | 5.44E+06 | 5.07E+06 | 5.05E+06 | 2.79E+06 | 3.03E+06 | 2.86E+06 | 2.77E+06 | 2.96E+06 | 3.13E+06 | 4.77E+06 | 2.92E+06 | 1.63 | 0.71 | 0.000107835 |
| P08572 | Collagen alpha-2(IV) chain OS=Homo sapiens OX=9606 GN=COL4A2 PE=1 SV=4 | 1 | 38 | 2 | 2.91E+06 | 2.64E+06 | 2.21E+06 | 2.95E+06 | 2.23E+06 | 2.42E+06 | 6.06E+06 | 7.01E+06 | 4.80E+06 | 4.52E+06 | 4.88E+06 | 4.77E+06 | 2.56E+06 | 5.34E+06 | 0.48 | -1.06 | 0.000562783 |
| P30048 | Thioredoxin-dependent peroxide reductase, mitochondrial OS=Homo sapiens OX=9606 GN=PRDX3 PE=1 SV=3 | 10 | 6 | 2 | 4.51E+06 | 3.53E+06 | 2.06E+06 | 1.92E+06 | 2.31E+06 | 2.43E+06 | 2.48E+06 | 2.01E+06 | 1.32E+06 | 1.13E+06 | 1.98E+06 | 1.48E+06 | 2.79E+06 | 1.73E+06 | 1.61 | 0.69 | 0.055012773 |
| P00450 | Ceruloplasmin OS=Homo sapiens OX=9606 GN=CP PE=1 SV=1 | 3 | 26 | 2 | 2.55E+06 | 2.72E+06 | 2.83E+06 | 2.59E+06 | 2.11E+06 | 3.20E+06 | 3.42E+06 | 2.66E+06 | 2.94E+06 | 3.65E+06 | 2.76E+06 | 3.34E+06 | 2.67E+06 | 3.13E+06 | 0.85 | -0.23 | 0.062951218 |
| P35590 | Tyrosine-protein kinase receptor Tie-1 OS=Homo sapiens OX=9606 GN=TIE1 PE=1 SV=1 | 3 | 15 | 2 | 2.73E+06 | 2.67E+06 | 4.49E+06 | 3.13E+06 | 2.94E+06 | 2.25E+06 | 3.41E+06 | 3.48E+06 | 5.03E+06 | 3.90E+06 | 3.37E+06 | 2.19E+06 | 3.04E+06 | 3.56E+06 | 0.85 | -0.23 | 0.308773458 |
| Q14558 | Phosphoribosyl pyrophosphate synthase-associated protein 1 OS=Homo sapiens OX=9606 GN=PRPSAP1 PE=1 SV=2 | 7 | 11 | 2 | 5.47E+06 | 1.15E+07 | 6.53E+06 | 6.14E+06 | 6.62E+06 | 1.09E+07 | 4.95E+06 | 1.50E+07 | 5.59E+06 | 6.41E+06 | 7.79E+06 | 6.07E+06 | 7.85E+06 | 7.64E+06 | 1.03 | 0.04 | 0.913759857 |
| P29279 | CCN family member 2 OS=Homo sapiens OX=9606 GN=CCN2 PE=1 SV=2 | 5 | 36 | 2 | 1.43E+07 | 1.22E+07 | 1.60E+07 | 1.24E+07 | 1.24E+07 | 1.34E+07 | 2.15E+07 | 2.36E+07 | 2.65E+07 | 2.07E+07 | 2.24E+07 | 1.88E+07 | 1.35E+07 | 2.22E+07 | 0.61 | -0.72 | 0.000110528 |
| Q3LXA3 | Triokinase/FMN cyclase OS=Homo sapiens OX=9606 GN=TKFC PE=1 SV=2 | 4 | 12 | 2 | 1.43E+06 | 2.41E+06 | 1.84E+06 | 1.95E+06 | 2.15E+06 | 1.70E+06 | 3.25E+06 | 2.77E+06 | 3.30E+06 | 2.67E+06 | 2.55E+06 | 2.64E+06 | 1.91E+06 | 2.86E+06 | 0.67 | -0.58 | 0.000629046 |
| Q13618 | Cullin-3 OS=Homo sapiens OX=9606 GN=CUL3 PE=1 SV=2 | 5 | 4 | 2 | 1.97E+06 | 7.76E+05 | 1.13E+06 | 7.69E+05 | 7.00E+05 | | 9.65E+05 | | | | | 6.01E+05 | 1.07E+06 | 7.83E+05 | 1.37 | 0.45 | 0.389921565 |
| P19623 | Spermidine synthase OS=Homo sapiens OX=9606 GN=SRM PE=1 SV=1 | 6 | 26 | 2 | 3.81E+06 | 3.74E+06 | 3.44E+06 | 4.18E+06 | 3.12E+06 | 2.67E+06 | 5.37E+06 | 4.56E+06 | 2.69E+06 | 3.13E+06 | 3.49E+06 | 2.64E+06 | 3.49E+06 | 3.65E+06 | 0.96 | -0.06 | 0.765110479 |
| Q9NZA1 | Chloride intracellular channel protein 5 OS=Homo sapiens OX=9606 GN=CLIC5 PE=1 SV=3 | 4 | 6 | 2 | | 6.13E+05 | 4.42E+05 | 6.84E+05 | | | 6.63E+05 | 4.60E+05 | 3.96E+05 | 3.25E+05 | 5.56E+05 | | 5.79E+05 | 4.80E+05 | 1.21 | 0.27 | 0.33925833 |
| Q03167 | Transforming growth factor beta receptor type 3 OS=Homo sapiens OX=9606 GN=TGFBR3 PE=1 SV=3 | 3 | 6 | 2 | 2.77E+06 | 2.34E+06 | 3.49E+06 | 2.65E+06 | 2.50E+06 | 2.41E+06 | 3.31E+06 | 1.61E+06 | 1.46E+06 | 2.25E+06 | 1.54E+06 | 2.60E+06 | 2.69E+06 | 2.13E+06 | 1.26 | 0.34 | 0.141342199 |
| O00186 | Syntaxin-binding protein 3 OS=Homo sapiens OX=9606 GN=STXBP3 PE=1 SV=2 | 5 | 13 | 2 | | 4.14E+05 | 1.05E+06 | | 6.73E+05 | 7.52E+05 | | | 2.00E+05 | 3.84E+05 | 4.86E+05 | | 7.22E+05 | 3.57E+05 | 2.02 | 1.02 | 0.067471556 |
| P08559 | Pyruvate dehydrogenase E1 component subunit alpha, somatic form, mitochondrial OS=Homo sapiens OX=9606 GN=PDHA1 PE=1 SV=3 | 6 | 5 | 2 | 1.33E+08 | 1.56E+08 | 1.24E+08 | 1.83E+08 | 1.08E+08 | 1.35E+08 | 2.23E+08 | 2.36E+08 | 1.82E+08 | 2.11E+08 | 1.97E+08 | 1.46E+08 | 1.40E+08 | 1.99E+08 | 0.70 | -0.51 | 0.006281812 |
| Q7Z2Z2 | Elongation factor-like GTPase 1 OS=Homo sapiens OX=9606 GN=EFL1 PE=1 SV=2 | 3 | 5 | 2 | 2.00E+06 | 1.63E+06 | 1.28E+06 | 1.33E+06 | 1.11E+06 | 1.78E+06 | 1.33E+06 | 1.07E+06 | 7.69E+05 | 1.66E+06 | 1.62E+06 | 1.16E+06 | 1.52E+06 | 1.27E+06 | 1.20 | 0.26 | 0.225343263 |
| P05455 | Lupus La protein OS=Homo sapiens OX=9606 GN=SSB PE=1 SV=2 | 4 | 8 | 2 | 2.11E+06 | 1.97E+06 | 3.04E+06 | 2.78E+06 | 2.22E+06 | 2.21E+06 | 1.16E+06 | 1.01E+06 | 1.64E+06 | 1.34E+06 | 1.18E+06 | 8.16E+05 | 2.39E+06 | 1.19E+06 | 2.01 | 1.00 | 0.000303836 |
| Q63HN8 | E3 ubiquitin-protein ligase RNF213 OS=Homo sapiens OX=9606 GN=RNF213 PE=1 SV=3 | 0 | 8 | 2 | | | | | | | | | | | | | | | | | |
| P04066 | Tissue alpha-L-fucosidase OS=Homo sapiens OX=9606 GN=FUCA1 PE=1 SV=4 | 5 | 14 | 2 | 2.52E+06 | 2.26E+06 | 2.08E+06 | 2.21E+06 | 1.94E+06 | 2.08E+06 | 3.80E+06 | 3.71E+06 | 3.25E+06 | 1.40E+06 | 2.43E+06 | 1.84E+06 | 2.18E+06 | 2.74E+06 | 0.80 | -0.33 | 0.236693727 |
| Q5JTH9 | RRP12-like protein OS=Homo sapiens OX=9606 GN=RRP12 PE=1 SV=2 | 4 | 4 | 2 | 5.22E+05 | | 5.30E+05 | 4.15E+05 | | 4.89E+05 | | | | | | | 4.89E+05 | | | | |
| P13497 | Bone morphogenetic protein 1 OS=Homo sapiens OX=9606 GN=BMP1 PE=1 SV=2 | 3 | 8 | 2 | 1.48E+07 | 1.68E+07 | 1.05E+07 | 1.17E+07 | 1.08E+07 | 1.08E+07 | 1.26E+07 | 1.29E+07 | 9.53E+06 | 8.84E+06 | 8.31E+06 | 9.10E+06 | 1.26E+07 | 1.02E+07 | 1.23 | 0.30 | 0.114051355 |
| P10646 | Tissue factor pathway inhibitor OS=Homo sapiens OX=9606 GN=TFPI PE=1 SV=1 | 6 | 28 | 2 | 4.13E+06 | 4.84E+06 | 3.00E+06 | 3.07E+06 | 3.10E+06 | 3.15E+06 | 5.09E+06 | 5.70E+06 | 4.10E+06 | 3.73E+06 | 3.33E+06 | 3.59E+06 | 3.55E+06 | 4.26E+06 | 0.83 | -0.26 | 0.182847344 |
| Q16563 | Synaptophysin-like protein 1 OS=Homo sapiens OX=9606 GN=SYPL1 PE=1 SV=1 | 15 | 16 | 2 | 2.02E+06 | | 1.67E+06 | 3.59E+06 | 4.07E+06 | 2.67E+06 | 1.53E+06 | 1.72E+06 | 1.02E+06 | 8.16E+05 | 1.75E+06 | 1.77E+06 | 2.80E+06 | 1.44E+06 | 1.95 | 0.97 | 0.036330681 |
| Q93009 | Ubiquitin carboxyl-terminal hydrolase 7 OS=Homo sapiens OX=9606 GN=USP7 PE=1 SV=2 | 3 | 4 | 2 | 2.36E+06 | 1.64E+06 | 1.18E+06 | 7.41E+05 | 1.43E+06 | 1.42E+06 | 1.23E+06 | 1.34E+06 | 1.29E+06 | 5.47E+05 | 5.87E+05 | 9.14E+05 | 1.46E+06 | 9.84E+05 | 1.49 | 0.57 | 0.102716953 |
| O60610 | Protein diaphanous homolog 1 OS=Homo sapiens OX=9606 GN=DIAPH1 PE=1 SV=2 | 2 | 12 | 2 | 9.77E+05 | 7.62E+05 | 5.38E+05 | 7.86E+05 | 5.68E+05 | 6.13E+05 | 7.43E+05 | 4.54E+05 | 3.75E+05 | 5.27E+05 | | 6.12E+05 | 7.07E+05 | 5.42E+05 | 1.30 | 0.38 | 0.110086066 |
| P13473 | Lysosome-associated membrane glycoprotein 2 OS=Homo sapiens OX=9606 GN=LAMP2 PE=1 SV=2 | 7 | 8 | 2 | 2.10E+06 | | 1.52E+06 | 1.66E+06 | 1.85E+06 | 1.31E+06 | | | 4.56E+05 | 4.41E+05 | 1.63E+06 | 1.37E+06 | 1.69E+06 | 9.74E+05 | 1.73 | 0.79 | 0.098951116 |

### Supplementary Table 4. List of all proteins identified by label free proteomics analysis.

| Accession | Description | Coverage [%] | # PSMs | # Unique Peptides | Amitriptyline 1 | Amitriptyline 2 | Amitriptyline 3 | Amitriptyline 4 | Amitriptyline 5 | Amitriptyline 6 | Vehicle 1 | Vehicle 2 | Vehicle 3 | Vehicle 4 | Vehicle 5 | Vehicle 6 | Average amitriptyline | Average vehicle | Ratio amitriptyline/ vehicle | Log 2 fold change (amitriptyline /vehicle) | P-value |
|---|---|---|---|---|---|---|---|---|---|---|---|---|---|---|---|---|---|---|---|---|---|
| Q86UP2 | Kinectin OS=Homo sapiens OX=9606 GN=KTN1 PE=1 SV=1 | 2 | 24 | 2 | 2.46E+06 | 2.63E+06 | 1.77E+06 | 1.98E+06 | 2.40E+06 | 2.08E+06 | 3.20E+06 | 3.46E+06 | 4.07E+06 | 2.85E+06 | 3.49E+06 | 2.99E+06 | 2.22E+06 | 3.34E+06 | 0.66 | -0.59 | 0.000622413 |
| P61011 | Signal recognition particle 54 kDa protein OS=Homo sapiens OX=9606 GN=SRP54 PE=1 SV=1 | 4 | 14 | 2 | 5.49E+06 | 5.85E+06 | 4.31E+06 | 4.41E+06 | 4.56E+06 | 3.88E+06 | 2.99E+06 | 3.16E+06 | 2.52E+06 | 1.93E+06 | 1.35E+06 | 8.93E+05 | 4.75E+06 | 2.14E+06 | 2.22 | 1.15 | 0.000338126 |
| P27144 | Adenylate kinase 4, mitochondrial OS=Homo sapiens OX=9606 GN=AK4 PE=1 SV=1 | 13 | 10 | 2 | | 1.90E+06 | 1.78E+06 | 1.90E+06 | 1.74E+06 | 1.57E+06 | 8.20E+05 | 1.08E+06 | 8.41E+05 | 9.82E+05 | 1.13E+06 | 6.88E+05 | 1.78E+06 | 9.23E+05 | 1.93 | 0.95 | 7.00256E-06 |
| P07947 | Tyrosine-protein kinase Yes OS=Homo sapiens OX=9606 GN=YES1 PE=1 SV=3 | 4 | 61 | 2 | 2.61E+06 | 2.97E+06 | 1.83E+06 | 3.08E+06 | 1.80E+06 | 3.72E+06 | 2.46E+06 | 1.78E+06 | 1.78E+06 | 2.20E+06 | 2.29E+06 | 3.06E+06 | 2.67E+06 | 2.26E+06 | 1.18 | 0.24 | 0.293676934 |
| P26358 | DNA (cytosine-5)-methyltransferase 1 OS=Homo sapiens OX=9606 GN=DNMT1 PE=1 SV=2 | 2 | 5 | 2 | 9.56E+06 | 1.31E+07 | 9.21E+06 | 1.39E+07 | 1.16E+07 | 8.94E+06 | 1.14E+07 | 1.67E+07 | 1.03E+07 | 1.82E+07 | 9.85E+06 | 8.86E+06 | 1.10E+07 | 1.26E+07 | 0.88 | -0.18 | 0.4317191 |
| Q53EL6 | Programmed cell death protein 4 OS=Homo sapiens OX=9606 GN=PDCD4 PE=1 SV=2 | 7 | 3 | 2 | 7.20E+05 | 7.47E+05 | 1.05E+06 | 7.92E+05 | 6.87E+05 | 6.48E+05 | 6.10E+05 | 5.09E+05 | 4.83E+05 | 5.63E+05 | 4.54E+05 | 5.47E+05 | 7.74E+05 | 5.28E+05 | 1.47 | 0.55 | 0.006947303 |
| P62750 | 60S ribosomal protein L23a OS=Homo sapiens OX=9606 GN=RPL23A PE=1 SV=1 | 16 | 13 | 2 | 2.01E+06 | 3.20E+06 | 4.58E+06 | 3.61E+06 | 4.56E+06 | 3.52E+06 | 3.35E+06 | 2.58E+06 | 3.53E+06 | 2.07E+06 | 2.50E+06 | 2.39E+06 | 3.58E+06 | 2.74E+06 | 1.31 | 0.39 | 0.100604684 |
| P51610 | Host cell factor 1 OS=Homo sapiens OX=9606 GN=HCFC1 PE=1 SV=2 | 1 | 6 | 2 | 5.52E+05 | 6.93E+05 | 7.11E+05 | 8.62E+05 | 1.00E+06 | 6.99E+05 | 3.41E+05 | 4.28E+05 | 4.96E+05 | | 3.59E+05 | | 7.53E+05 | 4.06E+05 | 1.85 | 0.89 | 0.001781073 |
| Q14247 | Src substrate cortactin OS=Homo sapiens OX=9606 GN=CTTN PE=1 SV=2 | 4 | 6 | 2 | 4.76E+05 | 2.40E+06 | 3.21E+06 | 3.12E+06 | 3.31E+06 | 2.73E+06 | | 1.74E+06 | | 4.98E+05 | 5.49E+05 | 1.00E+06 | 2.54E+06 | 9.48E+05 | 2.68 | 1.42 | 0.016237802 |
| P22352 | Glutathione peroxidase 3 OS=Homo sapiens OX=9606 GN=GPX3 PE=1 SV=2 | 8 | 393 | 2 | 1.95E+07 | 4.04E+07 | 4.12E+07 | 2.03E+07 | 2.94E+07 | 3.63E+07 | 3.69E+07 | 3.53E+07 | 6.21E+07 | 3.54E+07 | 4.91E+07 | 4.96E+07 | 3.06E+07 | 4.47E+07 | 0.68 | -0.55 | 0.039728575 |
| Q9UKK3 | Protein mono-ADP-ribosyltransferase PARP4 OS=Homo sapiens OX=9606 GN=PARP4 PE=1 SV=3 | 2 | 8 | 2 | 2.21E+06 | 2.90E+06 | 3.27E+06 | 1.90E+06 | 9.43E+05 | 2.16E+06 | 1.46E+06 | 1.95E+06 | | 4.65E+05 | 1.21E+06 | 6.30E+05 | 2.23E+06 | 1.14E+06 | 1.96 | 0.97 | 0.031716787 |
| Q9HA64 | Ketosamine-3-kinase OS=Homo sapiens OX=9606 GN=FN3KRP PE=1 SV=2 | 10 | 8 | 2 | 3.07E+06 | 2.92E+06 | 2.94E+06 | 2.53E+06 | 2.44E+06 | 2.32E+06 | 2.26E+06 | 2.04E+06 | 2.70E+06 | 2.27E+06 | 3.12E+06 | 2.64E+06 | 2.70E+06 | 2.50E+06 | 1.08 | 0.11 | 0.357277828 |
| Q9UDY4 | DnaJ homolog subfamily B member 4 OS=Homo sapiens OX=9606 GN=DNAJB4 PE=1 SV=1 | 6 | 16 | 2 | 2.79E+06 | 4.11E+06 | 2.55E+06 | 3.76E+06 | 2.43E+06 | 3.97E+06 | 3.23E+06 | 2.79E+06 | 2.62E+06 | 2.55E+06 | 2.18E+06 | 2.82E+06 | 3.27E+06 | 2.70E+06 | 1.21 | 0.28 | 0.136321765 |
| P63173 | 60S ribosomal protein L38 OS=Homo sapiens OX=9606 GN=RPL38 PE=1 SV=2 | 33 | 24 | 2 | 6.91E+06 | 8.91E+06 | 1.10E+07 | 1.27E+07 | 1.11E+07 | 1.09E+07 | 6.86E+06 | 6.07E+06 | 9.12E+06 | 7.97E+06 | 8.14E+06 | 4.21E+06 | 1.03E+07 | 7.06E+06 | 1.45 | 0.54 | 0.015947836 |
| Q96KG9 | N-terminal kinase-like protein OS=Homo sapiens OX=9606 GN=SCYL1 PE=1 SV=1 | 4 | 4 | 2 | 9.96E+05 | 1.25E+06 | 1.15E+06 | 1.06E+06 | 9.06E+05 | 9.96E+05 | 7.44E+05 | 7.87E+05 | 8.02E+05 | 8.63E+05 | 8.29E+05 | 5.91E+05 | 1.06E+06 | 7.69E+05 | 1.38 | 0.46 | 0.001253744 |
| Q8NBI5 | Solute carrier family 43 member 3 OS=Homo sapiens OX=9606 GN=SLC43A3 PE=1 SV=2 | 5 | 18 | 2 | 5.74E+06 | 5.90E+06 | 5.41E+06 | 5.13E+06 | 5.29E+06 | 5.06E+06 | 4.20E+06 | 3.98E+06 | 3.67E+06 | 4.13E+06 | 3.34E+06 | 3.41E+06 | 5.42E+06 | 3.79E+06 | 1.43 | 0.52 | 1.18136E-05 |
| P39059 | Collagen alpha-1(XV) chain OS=Homo sapiens OX=9606 GN=COL15A1 PE=1 SV=2 | 1 | 34 | 2 | 1.13E+07 | 1.23E+07 | 8.66E+06 | 9.25E+06 | 9.67E+06 | 9.40E+06 | 1.42E+07 | 1.56E+07 | 1.13E+07 | 1.32E+07 | 1.14E+07 | 1.19E+07 | 1.01E+07 | 1.29E+07 | 0.78 | -0.36 | 0.010780158 |
| P04180 | Phosphatidylcholine-sterol acyltransferase OS=Homo sapiens OX=9606 GN=LCAT PE=1 SV=1 | 4 | 71 | 2 | 1.35E+07 | 1.52E+07 | 2.86E+07 | 1.52E+07 | 1.37E+07 | 1.39E+07 | 2.26E+07 | 2.35E+07 | 2.54E+07 | 2.44E+07 | 1.63E+07 | 1.86E+07 | 1.67E+07 | 2.18E+07 | 0.77 | -0.38 | 0.105951358 |
| Q15046 | Lysine--tRNA ligase OS=Homo sapiens OX=9606 GN=KARS1 PE=1 SV=3 | 5 | 8 | 2 | 2.99E+06 | 4.24E+06 | 4.48E+06 | 5.18E+06 | 4.11E+06 | 3.27E+06 | 3.57E+06 | 3.99E+06 | 4.41E+06 | 4.29E+06 | 2.60E+06 | 9.07E+05 | 4.04E+06 | 3.29E+06 | 1.23 | 0.30 | 0.2725313 |
| Q8N8N7 | Prostaglandin reductase 2 OS=Homo sapiens OX=9606 GN=PTGR2 PE=1 SV=1 | 6 | 5 | 2 | | 5.54E+05 | 4.93E+05 | 4.03E+05 | 3.07E+05 | 4.17E+05 | 1.29E+06 | | | | 5.07E+05 | 7.07E+05 | 4.35E+05 | 8.36E+05 | 0.52 | -0.94 | 0.228612787 |
| Q9BRL6 | Serine/arginine-rich splicing factor 8 OS=Homo sapiens OX=9606 GN=SRSF8 PE=1 SV=1 | 5 | 30 | 2 | 9.24E+06 | 1.16E+07 | 1.06E+07 | 1.15E+07 | 1.13E+07 | 9.80E+06 | 1.13E+07 | 1.27E+07 | 1.15E+07 | 1.28E+07 | 1.07E+07 | 1.10E+07 | 1.07E+07 | 1.17E+07 | 0.91 | -0.13 | 0.091034611 |
| Q8N1F7 | Nuclear pore complex protein Nup93 OS=Homo sapiens OX=9606 GN=NUP93 PE=1 SV=2 | 5 | 24 | 2 | 2.36E+06 | 3.34E+06 | 1.81E+06 | 3.20E+06 | 3.51E+06 | 2.74E+06 | 1.09E+06 | 2.32E+06 | 2.73E+06 | 3.10E+06 | 2.76E+06 | 6.49E+05 | 2.83E+06 | 2.11E+06 | 1.34 | 0.42 | 0.176452279 |
| Q13428 | Treacle protein OS=Homo sapiens OX=9606 GN=TCOF1 PE=1 SV=3 | 1 | 12 | 2 | 4.03E+06 | 3.93E+06 | 3.10E+06 | 4.14E+06 | 3.65E+06 | 3.43E+06 | 1.06E+06 | 1.18E+06 | 7.72E+05 | 1.52E+06 | 1.61E+06 | 1.81E+06 | 3.71E+06 | 1.32E+06 | 2.80 | 1.49 | 1.00794E-06 |
| P59998 | Actin-related protein 2/3 complex subunit 4 OS=Homo sapiens OX=9606 GN=ARPC4 PE=1 SV=3 | 16 | 40 | 2 | 9.60E+06 | 9.65E+06 | 6.17E+06 | 7.92E+06 | 6.43E+06 | 6.02E+06 | 9.66E+06 | 8.99E+06 | 6.65E+06 | 6.65E+06 | 6.90E+06 | 5.87E+06 | 7.63E+06 | 7.45E+06 | 1.02 | 0.03 | 0.849522163 |
| Q9UKG1 | DCC-interacting protein 13-alpha OS=Homo sapiens OX=9606 GN=APPL1 PE=1 SV=1 | 4 | 11 | 2 | 1.28E+06 | 1.73E+06 | 1.73E+06 | 1.90E+06 | 1.90E+06 | 1.61E+06 | 1.45E+06 | | 1.68E+06 | | 2.16E+06 | | 1.69E+06 | 1.76E+06 | 0.96 | -0.06 | 0.775288131 |
| Q92572 | AP-3 complex subunit sigma-1 OS=Homo sapiens OX=9606 GN=AP3S1 PE=1 SV=1 | 12 | 17 | 2 | 3.86E+06 | 4.72E+06 | 2.72E+06 | 3.73E+06 | 3.31E+06 | 2.87E+06 | 1.23E+06 | 1.14E+06 | 1.98E+06 | 2.44E+06 | 1.21E+06 | 1.64E+06 | 3.53E+06 | 1.61E+06 | 2.20 | 1.14 | 0.000531902 |
| P19367 | Hexokinase-1 OS=Homo sapiens OX=9606 GN=HK1 PE=1 SV=3 | 3 | 4 | 2 | 3.13E+06 | 2.81E+06 | 2.68E+06 | 2.41E+06 | 2.91E+06 | 3.17E+06 | 1.54E+06 | 5.78E+05 | 6.01E+05 | 4.57E+05 | 7.80E+05 | 1.55E+06 | 2.85E+06 | 9.18E+05 | 3.11 | 1.64 | 3.41232E-05 |
| Q08722 | Leukocyte surface antigen CD47 OS=Homo sapiens OX=9606 GN=CD47 PE=1 SV=1 | 5 | 48 | 2 | 9.50E+06 | 1.04E+07 | 1.48E+07 | 1.07E+07 | 1.34E+07 | 9.27E+06 | 8.13E+06 | 8.44E+06 | 8.77E+06 | 7.23E+06 | 7.90E+06 | 5.30E+06 | 1.13E+07 | 7.63E+06 | 1.49 | 0.57 | 0.007746523 |
| O76074 | cGMP-specific 3',5'-cyclic phosphodiesterase OS=Homo sapiens OX=9606 GN=PDE5A PE=1 SV=2 | 3 | 9 | 2 | 1.03E+06 | | 3.54E+05 | | 4.80E+05 | 3.33E+05 | 2.10E+06 | | 1.71E+06 | | 1.27E+06 | 1.12E+06 | 5.49E+05 | 1.55E+06 | 0.35 | -1.50 | 0.012534034 |
| Q9UKM9 | RNA-binding protein Raly OS=Homo sapiens OX=9606 GN=RALY PE=1 SV=1 | 6 | 15 | 2 | 7.09E+06 | 7.58E+06 | 8.06E+06 | 8.79E+06 | 8.26E+06 | 7.94E+06 | 4.93E+06 | 5.13E+06 | 2.10E+06 | 1.68E+06 | 4.87E+06 | 4.17E+06 | 7.95E+06 | 3.81E+06 | 2.08 | 1.06 | 0.000636003 |
| Q96JB2 | Conserved oligomeric Golgi complex subunit 3 OS=Homo sapiens OX=9606 GN=COG3 PE=1 SV=3 | 4 | 4 | 2 | | | | | | | | | | | | | | | | | |
| Q13451 | Peptidyl-prolyl cis-trans isomerase FKBP5 OS=Homo sapiens OX=9606 GN=FKBP5 PE=1 SV=2 | 4 | 6 | 2 | 1.49E+06 | 1.10E+06 | 1.05E+06 | 1.35E+06 | 1.26E+06 | | 5.90E+05 | 5.41E+05 | 6.33E+05 | | | | 1.25E+06 | 5.88E+05 | 2.12 | 1.09 | 0.000650481 |
| Q12846 | Syntaxin-4 OS=Homo sapiens OX=9606 GN=STX4 PE=1 SV=2 | 7 | 16 | 2 | 2.94E+06 | 3.41E+06 | 3.81E+06 | 3.40E+06 | 3.31E+06 | 4.01E+06 | 2.67E+06 | 2.81E+06 | 2.33E+06 | 2.51E+06 | 1.94E+06 | 7.44E+05 | 3.48E+06 | 3.28E+06 | 1.06 | 0.08 | 0.82510637 |
| Q9Y2T2 | AP-3 complex subunit mu-1 OS=Homo sapiens OX=9606 GN=AP3M1 PE=1 SV=1 | 4 | 14 | 2 | 1.09E+06 | 9.78E+05 | 2.17E+06 | 1.73E+06 | 1.85E+06 | 2.01E+06 | 7.76E+05 | 1.34E+06 | 1.43E+06 | 6.68E+05 | 1.07E+06 | 9.94E+05 | 1.64E+06 | 1.05E+06 | 1.56 | 0.65 | 0.035494254 |
| Q06278 | Aldehyde oxidase OS=Homo sapiens OX=9606 GN=AOX1 PE=1 SV=2 | 2 | 16 | 2 | 9.74E+05 | 1.28E+06 | 1.51E+06 | 1.18E+06 | | | 1.63E+06 | 2.22E+06 | 2.07E+06 | 2.19E+06 | 2.52E+06 | 1.84E+06 | 1.24E+06 | 2.08E+06 | 0.59 | -0.75 | 0.001151674 |
| Q9UJU6 | Drebrin-like protein OS=Homo sapiens OX=9606 GN=DBNL PE=1 SV=1 | 6 | 15 | 2 | | | | | | | | | | | | | | | | | |
| P10619 | Lysosomal protective protein OS=Homo sapiens OX=9606 GN=CTSA PE=1 SV=2 | 5 | 6 | 2 | 3.91E+06 | 5.76E+06 | 4.01E+06 | 3.69E+06 | 2.68E+06 | 1.03E+06 | 2.19E+06 | 2.38E+06 | 2.46E+06 | 2.56E+06 | 7.96E+05 | 7.39E+05 | 3.51E+06 | 1.85E+06 | 1.89 | 0.92 | 0.053986254 |
| Q7L1Q6 | Basic leucine zipper and W2 domain-containing protein 1 OS=Homo sapiens OX=9606 GN=BZW1 PE=1 SV=1 | 5 | 21 | 2 | 6.16E+06 | 2.55E+06 | 6.23E+06 | 1.76E+06 | 6.88E+06 | 1.66E+06 | 5.53E+06 | 1.66E+06 | 5.89E+06 | 5.84E+06 | 5.14E+06 | 4.38E+06 | 4.21E+06 | 4.74E+06 | 0.89 | -0.17 | 0.666941036 |
| O15400 | Syntaxin-7 OS=Homo sapiens OX=9606 GN=STX7 PE=1 SV=4 | 8 | 5 | 2 | 2.07E+06 | 2.04E+06 | 1.94E+06 | 1.86E+06 | 1.79E+06 | 5.45E+05 | 7.99E+05 | 8.03E+05 | 9.77E+05 | 8.06E+05 | 4.67E+05 | | 1.71E+06 | 7.71E+05 | 2.21 | 1.15 | 0.009068842 |
| Q7L9L4 | MOB kinase activator 1B OS=Homo sapiens OX=9606 GN=MOB1B PE=1 SV=3 | 9 | 28 | 2 | 1.60E+06 | 2.26E+06 | 1.99E+06 | 2.92E+06 | 2.35E+06 | 2.73E+06 | 2.53E+06 | 2.65E+06 | 2.71E+06 | 2.90E+06 | 3.58E+06 | 2.06E+06 | 2.31E+06 | 2.74E+06 | 0.84 | -0.25 | 0.158502927 |
| Q9UNF0 | Protein kinase C and casein kinase substrate in neurons protein 2 OS=Homo sapiens OX=9606 GN=PACSIN2 PE=1 SV=2 | 4 | 7 | 2 | 1.82E+06 | 1.92E+06 | 2.44E+06 | 2.62E+06 | 2.35E+06 | 1.78E+06 | 1.07E+06 | 1.04E+06 | 1.06E+06 | 9.77E+05 | 1.45E+06 | 1.03E+06 | 2.16E+06 | 1.10E+06 | 1.95 | 0.96 | 0.000317653 |
| P53634 | Dipeptidyl peptidase 1 OS=Homo sapiens OX=9606 GN=CTSC PE=1 SV=2 | 5 | 6 | 2 | 1.93E+06 | 2.17E+06 | 8.51E+05 | 2.28E+06 | 2.32E+06 | 2.47E+06 | 1.92E+06 | 1.99E+06 | 2.85E+05 | 1.35E+06 | 1.72E+06 | 1.25E+06 | 2.00E+06 | 1.42E+06 | 1.41 | 0.50 | 0.130759669 |
| P15529 | Membrane cofactor protein OS=Homo sapiens OX=9606 GN=CD46 PE=1 SV=3 | 4 | 14 | 2 | 2.92E+06 | 2.41E+06 | 2.21E+06 | 4.32E+06 | 2.92E+06 | 4.21E+06 | 1.43E+06 | 1.30E+06 | 1.15E+06 | 1.65E+06 | 1.34E+06 | 9.66E+05 | 3.17E+06 | 1.31E+06 | 2.42 | 1.28 | 0.003167779 |
| Q99986 | Serine/threonine-protein kinase VRK1 OS=Homo sapiens OX=9606 GN=VRK1 PE=1 SV=1 | 6 | 8 | 2 | | 4.57E+05 | 8.95E+05 | 7.30E+05 | 6.34E+05 | 6.32E+05 | 5.35E+05 | 4.17E+05 | 4.46E+05 | 3.39E+05 | 4.14E+05 | 3.16E+05 | 6.70E+05 | 4.11E+05 | 1.63 | 0.70 | 0.01831036 |
| O43837 | Isocitrate dehydrogenase [NAD] subunit beta, mitochondrial OS=Homo sapiens OX=9606 GN=IDH3B PE=1 SV=2 | 5 | 18 | 2 | 2.31E+06 | 2.34E+06 | 1.66E+06 | 2.41E+06 | 2.21E+06 | 1.33E+06 | 1.25E+06 | 1.26E+06 | 1.05E+06 | 1.32E+06 | | 6.05E+05 | 2.04E+06 | 1.10E+06 | 1.86 | 0.90 | 0.00237797 |
| Q9H267 | Vacuolar protein sorting-associated protein 33B OS=Homo sapiens OX=9606 GN=VPS33B PE=1 SV=2 | 5 | 3 | 2 | 2.68E+06 | 1.28E+06 | 1.52E+06 | 1.25E+06 | 8.49E+05 | 2.30E+06 | 2.32E+06 | 1.91E+06 | 9.45E+05 | 1.12E+06 | 1.37E+06 | 1.44E+06 | 1.65E+06 | 1.52E+06 | 1.09 | 0.12 | 0.720299849 |
| P56199 | Integrin alpha-1 OS=Homo sapiens OX=9606 GN=ITGA1 PE=1 SV=2 | 2 | 11 | 2 | 3.56E+06 | 3.33E+06 | 3.52E+06 | 4.00E+06 | 3.47E+06 | 2.57E+06 | 2.54E+06 | 2.42E+06 | 2.15E+06 | 1.10E+06 | 1.52E+06 | 1.97E+06 | 3.41E+06 | 1.95E+06 | 1.75 | 0.81 | 0.000633867 |
| O60313 | Dynamin-like 120 kDa protein, mitochondrial OS=Homo sapiens OX=9606 GN=OPA1 PE=1 SV=3 | 3 | 7 | 2 | 3.26E+06 | 2.05E+06 | 1.52E+06 | 3.14E+06 | 2.96E+06 | 2.13E+06 | | 4.76E+06 | | 2.54E+06 | 4.96E+06 | 1.23E+06 | 2.51E+06 | 3.37E+06 | 0.74 | -0.43 | 0.418604386 |
| Q14C86 | GTPase-activating protein and VPS9 domain-containing protein 1 OS=Homo sapiens OX=9606 GN=GAPVD1 PE=1 SV=2 | 2 | 6 | 2 | 1.90E+06 | 1.89E+06 | 1.45E+06 | 1.98E+06 | 2.29E+06 | 2.15E+06 | | | | 1.29E+06 | | | 1.94E+06 | 1.29E+06 | 1.51 | 0.60 | |
| P13987 | CD59 glycoprotein OS=Homo sapiens OX=9606 GN=CD59 PE=1 SV=1 | 16 | 12 | 2 | | 5.12E+05 | | 2.30E+07 | 2.05E+07 | 1.84E+07 | | 5.11E+05 | 1.77E+07 | | | | 1.56E+07 | 9.08E+06 | 1.72 | 0.78 | 0.58845335 |
| P36915 | Guanine nucleotide-binding protein-like 1 OS=Homo sapiens OX=9606 GN=GNL1 PE=1 SV=2 | 3 | 6 | 2 | 3.16E+06 | 3.45E+06 | 2.61E+06 | 2.94E+06 | 2.06E+06 | 2.55E+06 | 2.44E+06 | 2.67E+06 | 2.31E+06 | 1.78E+06 | 2.04E+06 | 1.57E+06 | 2.79E+06 | 2.14E+06 | 1.31 | 0.39 | 0.032533996 |
| Q15102 | Platelet-activating factor acetylhydrolase IB subunit gamma OS=Homo sapiens OX=9606 GN=PAFAH1B3 PE=1 SV=1 | 8 | 16 | 2 | 4.44E+06 | 4.22E+06 | 4.61E+06 | 4.05E+06 | 3.86E+06 | 4.90E+06 | 4.54E+06 | 4.23E+06 | 4.02E+06 | 3.70E+06 | 3.30E+06 | 4.24E+06 | 4.35E+06 | 4.00E+06 | 1.09 | 0.12 | 0.18410744 |
| P09012 | U1 small nuclear ribonucleoprotein A OS=Homo sapiens OX=9606 GN=SNRPA PE=1 SV=3 | 6 | 28 | 2 | 5.24E+06 | 5.09E+06 | 7.11E+06 | 7.16E+06 | 5.57E+06 | 4.25E+06 | 1.86E+06 | 1.64E+06 | 5.17E+06 | 4.29E+06 | 2.12E+06 | 3.12E+06 | 5.74E+06 | 3.03E+06 | 1.89 | 0.92 | 0.005371269 |
| Q13620 | Cullin-4B OS=Homo sapiens OX=9606 GN=CUL4B PE=1 SV=4 | 2 | 6 | 2 | 7.34E+05 | 7.60E+05 | 8.12E+05 | 9.73E+05 | 7.13E+05 | 7.44E+05 | 9.05E+05 | 6.15E+05 | 5.64E+05 | 4.52E+05 | 7.21E+05 | 3.67E+05 | 7.89E+05 | 6.04E+05 | 1.31 | 0.39 | 0.070792404 |
| P22061 | Protein-L-isoaspartate(D-aspartate) O-methyltransferase OS=Homo sapiens OX=9606 GN=PCMT1 PE=1 SV=4 | 7 | 28 | 2 | 4.27E+06 | 5.45E+06 | 5.38E+06 | 5.84E+06 | 5.45E+06 | 4.84E+06 | 5.86E+06 | 5.83E+06 | 5.64E+06 | 5.84E+06 | 5.08E+06 | 4.28E+06 | 5.21E+06 | 5.42E+06 | 0.96 | -0.06 | 0.548697345 |
| Q9UGM3 | Deleted in malignant brain tumors 1 protein OS=Homo sapiens OX=9606 GN=DMBT1 PE=1 SV=2 | 9 | 3 | 2 | | | | | | | 2.04E+06 | 2.04E+06 | 1.40E+06 | 2.65E+06 | | | | 2.03E+06 | | | |
| P45973 | Chromobox protein homolog 5 OS=Homo sapiens OX=9606 GN=CBX5 PE=1 SV=1 | 11 | 22 | 2 | 8.46E+06 | 7.86E+06 | 9.13E+06 | 7.26E+06 | 9.28E+06 | 8.75E+06 | 3.86E+06 | 2.68E+06 | 4.45E+06 | 2.70E+06 | 3.35E+06 | 3.17E+06 | 8.46E+06 | 3.37E+06 | 2.51 | 1.33 | 3.30351E-07 |
| P42765 | 3-ketoacyl-CoA thiolase, mitochondrial OS=Homo sapiens OX=9606 GN=ACAA2 PE=1 SV=2 | 5 | 6 | 2 | 3.78E+06 | | 3.59E+06 | 2.04E+06 | 3.66E+06 | 4.32E+06 | 1.08E+06 | 1.43E+06 | 1.03E+06 | 3.22E+06 | 3.34E+06 | 1.05E+06 | 3.48E+06 | 1.86E+06 | 1.87 | 0.90 | 0.023318943 |
| P11172 | Uridine 5'-monophosphate synthase OS=Homo sapiens OX=9606 GN=UMPS PE=1 SV=1 | 4 | 8 | 2 | 3.69E+06 | 3.95E+06 | 3.36E+06 | 3.94E+06 | 2.77E+06 | 4.25E+06 | 3.23E+06 | 3.40E+06 | 3.23E+06 | | 3.16E+06 | 3.15E+06 | 3.66E+06 | 3.23E+06 | 1.13 | 0.18 | 0.104645707 |
| Q04446 | 1,4-alpha-glucan-branching enzyme OS=Homo sapiens OX=9606 GN=GBE1 PE=1 SV=3 | 5 | 3 | 2 | 1.14E+06 | | 1.08E+06 | 2.57E+05 | 1.06E+06 | 6.14E+05 | | | | | | | 8.32E+05 | | | | |
| O15305 | Phosphomannomutase 2 OS=Homo sapiens OX=9606 GN=PMM2 PE=1 SV=1 | 7 | 21 | 2 | 9.22E+06 | 1.01E+07 | 1.03E+07 | 1.26E+07 | 9.85E+06 | 8.21E+06 | 1.27E+07 | 1.35E+07 | 1.28E+07 | 1.25E+07 | 1.34E+07 | 1.16E+07 | 1.01E+07 | 1.27E+07 | 0.79 | -0.34 | 0.004652396 |
| Q9BVJ7 | Dual specificity protein phosphatase 23 OS=Homo sapiens OX=9606 GN=DUSP23 PE=1 SV=1 | 15 | 5 | 2 | 2.13E+06 | 2.06E+06 | 1.73E+06 | 1.56E+06 | 9.82E+05 | 1.92E+06 | | 1.63E+06 | 5.14E+05 | 1.73E+06 | 7.60E+05 | 5.30E+05 | 1.73E+06 | 1.03E+06 | 1.67 | 0.74 | 0.065199107 |
| Q5M9N0 | Coiled-coil domain-containing protein 158 OS=Homo sapiens OX=9606 GN=CCDC158 PE=2 SV=2 | 3 | 35 | 2 | 1.09E+08 | 1.23E+08 | 1.98E+08 | 2.29E+08 | 1.87E+08 | 1.79E+08 | 2.47E+08 | 2.41E+08 | 2.82E+08 | 2.53E+08 | 2.39E+08 | 2.31E+08 | 1.71E+08 | 2.49E+08 | 0.69 | -0.54 | 0.007174991 |
| P43246 | DNA mismatch repair protein Msh2 OS=Homo sapiens OX=9606 GN=MSH2 PE=1 SV=1 | 2 | 21 | 2 | 1.78E+07 | 2.09E+07 | 1.92E+07 | 2.14E+07 | 2.05E+07 | 1.85E+07 | 2.56E+07 | 2.57E+07 | 3.10E+07 | 2.97E+07 | 2.55E+07 | 2.13E+07 | 1.97E+07 | 2.65E+07 | 0.75 | -0.42 | 0.003534954 |
| Q9H845 | Complex I assembly factor ACAD9, mitochondrial OS=Homo sapiens OX=9606 GN=ACAD9 PE=1 SV=1 | 3 | 12 | 2 | 3.51E+08 | 3.57E+08 | 3.41E+08 | 2.94E+08 | 2.82E+08 | 3.23E+08 | 6.42E+08 | 6.35E+08 | 4.64E+08 | 5.78E+08 | 5.60E+08 | 4.58E+08 | 3.25E+08 | 5.56E+08 | 0.58 | -0.78 | 0.000434855 |
| P60520 | Gamma-aminobutyric acid receptor-associated protein-like 2 OS=Homo sapiens OX=9606 GN=GABARAPL2 PE=1 SV=1 | 12 | 17 | 2 | 4.19E+06 | 5.77E+06 | 4.83E+06 | 5.73E+06 | 6.14E+06 | 4.06E+06 | 9.02E+05 | 1.01E+06 | 2.73E+06 | 1.79E+06 | 2.81E+06 | 1.05E+06 | 5.12E+06 | 1.72E+06 | 2.98 | 1.58 | 5.40329E-05 |
| Q14289 | Protein-tyrosine kinase 2-beta OS=Homo sapiens OX=9606 GN=PTK2B PE=1 SV=2 | 2 | 20 | 2 | 8.39E+06 | 8.13E+06 | 1.06E+07 | 1.33E+07 | 1.23E+07 | 1.08E+07 | 1.66E+07 | 1.63E+07 | 2.13E+07 | 1.70E+07 | 1.75E+07 | 1.33E+07 | 1.06E+07 | 1.70E+07 | 0.62 | -0.69 | 0.00083523 |
| P09496 | Clathrin light chain A OS=Homo sapiens OX=9606 GN=CLTA PE=1 SV=1 | 6 | 24 | 2 | 1.60E+07 | 1.83E+07 | 1.36E+07 | 1.18E+07 | 1.48E+07 | 1.46E+07 | 2.00E+07 | 1.98E+07 | 2.43E+07 | 1.70E+07 | 2.13E+07 | 1.52E+07 | 1.49E+07 | 1.96E+07 | 0.76 | -0.40 | 0.015601422 |
| P46976 | Glycogenin-1 OS=Homo sapiens OX=9606 GN=GYG1 PE=1 SV=4 | 5 | 4 | 2 | 3.34E+06 | 3.24E+06 | 3.65E+06 | 4.75E+06 | 3.99E+06 | 1.91E+06 | 3.89E+06 | 4.69E+06 | 4.44E+06 | 5.55E+06 | 3.48E+06 | 3.38E+06 | 3.48E+06 | 4.24E+06 | 0.82 | -0.28 | 0.169241516 |
| O95347 | Structural maintenance of chromosomes protein 2 OS=Homo sapiens OX=9606 GN=SMC2 PE=1 SV=2 | 2 | 5 | 2 | 1.17E+06 | 1.59E+06 | 1.54E+06 | 1.12E+06 | 1.24E+06 | 1.17E+06 | 5.02E+05 | 3.87E+05 | | | | 2.70E+05 | 1.31E+06 | 3.86E+05 | 3.38 | 1.76 | 7.71827E-05 |
| Q8WZ42 | Titin OS=Homo sapiens OX=9606 GN=TTN PE=1 SV=4 | 0 | 11 | 2 | 2.97E+06 | 3.63E+06 | 2.74E+06 | 3.23E+06 | 2.04E+06 | 1.35E+06 | 5.28E+06 | 2.98E+06 | 4.08E+06 | 2.07E+06 | 5.35E+06 | 4.30E+06 | 2.66E+06 | 4.01E+06 | 0.66 | -0.59 | 0.061343705 |
| Q12882 | Dihydropyrimidine dehydrogenase [NADP(+)] OS=Homo sapiens OX=9606 GN=DPYD PE=1 SV=2 | 2 | 6 | 2 | 3.33E+05 | 4.99E+05 | 4.48E+05 | 6.44E+05 | 5.78E+05 | 3.11E+05 | 2.70E+05 | 9.20E+05 | 4.27E+05 | 8.64E+05 | | 5.78E+05 | 4.69E+05 | 6.12E+05 | 0.77 | -0.38 | 0.336009406 |
| P25398 | 40S ribosomal protein S12 OS=Homo sapiens OX=9606 GN=RPS12 PE=1 SV=3 | 14 | 6 | 2 | 3.22E+06 | 2.87E+06 | 3.77E+06 | 3.92E+06 | 3.47E+06 | 2.74E+06 | 3.46E+06 | 2.58E+06 | 3.42E+06 | 2.45E+06 | 2.90E+06 | 2.61E+06 | 3.33E+06 | 2.90E+06 | 1.15 | 0.20 | 0.135191235 |
| Q96SN8 | CDK5 regulatory subunit-associated protein 2 OS=Homo sapiens OX=9606 GN=CDK5RAP2 PE=1 SV=5 | 1 | 5 | 2 | 1.14E+08 | 1.23E+08 | 1.08E+08 | 1.14E+08 | 1.18E+08 | 1.08E+08 | 1.49E+08 | 1.47E+08 | 1.50E+08 | 1.55E+08 | 1.36E+08 | 1.20E+08 | 1.14E+08 | 1.43E+08 | 0.80 | -0.32 | 0.001797956 |
| P04350 | Tubulin beta-4A chain OS=Homo sapiens OX=9606 GN=TUBB4A PE=1 SV=2 | 78 | 4364 | 3 | 7.59E+07 | 7.21E+07 | 4.38E+07 | 5.07E+07 | 5.37E+07 | 6.12E+07 | 4.05E+07 | 3.75E+07 | 3.17E+07 | 4.15E+07 | 4.38E+07 | 3.76E+07 | 5.96E+07 | 3.88E+07 | 1.54 | 0.62 | 0.008213349 |
| Q13885 | Tubulin beta-2A chain OS=Homo sapiens OX=9606 GN=TUBB2A PE=1 SV=1 | 62 | 4377 | 3 | 3.49E+06 | 4.02E+06 | 4.07E+06 | 4.28E+06 | 4.93E+06 | 4.06E+06 | 3.19E+06 | 5.09E+06 | 3.48E+06 | 4.60E+06 | 4.13E+06 | 4.17E+06 | 4.14E+06 | 4.11E+06 | 1.01 | 0.01 | 0.933456377 |
| Q16778 | Histone H2B type 2-E OS=Homo sapiens OX=9606 GN=HIST2H2BE PE=1 SV=3 | 56 | 6292 | 3 | 1.26E+09 | 1.32E+09 | 1.07E+09 | 1.26E+09 | 1.21E+09 | 1.24E+09 | 1.31E+09 | 1.40E+09 | 1.41E+09 | 1.67E+09 | 1.82E+09 | 1.11E+09 | 1.23E+09 | 1.45E+09 | 0.84 | -0.25 | 0.082991654 |
| Q8IUE6 | Histone H2A type 2-B OS=Homo sapiens OX=9606 GN=HIST2H2AB PE=1 SV=3 | 63 | 414 | 3 | 2.24E+08 | 2.40E+08 | 2.14E+08 | 1.39E+08 | 1.82E+08 | 2.06E+08 | 2.44E+08 | 2.82E+08 | 2.21E+08 | 1.66E+08 | 2.50E+08 | 2.12E+08 | 2.01E+08 | 2.29E+08 | 0.88 | -0.19 | 0.221812514 |
| Q14240 | Eukaryotic initiation factor 4A-II OS=Homo sapiens OX=9606 GN=EIF4A2 PE=1 SV=2 | 36 | 429 | 3 | 1.33E+07 | 1.11E+07 | 9.42E+06 | 1.35E+07 | 1.08E+07 | 1.08E+07 | 1.14E+07 | 1.22E+07 | 9.48E+06 | 1.01E+07 | 8.77E+06 | 8.16E+06 | 1.18E+07 | 1.00E+07 | 1.17 | 0.23 | 0.083816263 |
| P10321 | HLA class I histocompatibility antigen, C alpha chain OS=Homo sapiens OX=9606 GN=HLA-C PE=1 SV=3 | 30 | 444 | 3 | 5.37E+07 | 5.50E+07 | 4.31E+07 | 4.16E+07 | 4.37E+07 | 4.51E+07 | 2.27E+07 | 2.35E+07 | 2.00E+07 | 1.88E+07 | 1.58E+07 | 1.79E+07 | 4.70E+07 | 1.98E+07 | 2.38 | 1.25 | 1.18481E-05 |
| P31150 | Rab GDP dissociation inhibitor alpha OS=Homo sapiens OX=9606 GN=GDI1 PE=1 SV=2 | 31 | 206 | 3 | 5.97E+06 | 8.43E+06 | 8.86E+06 | 9.74E+06 | 8.40E+06 | 8.98E+06 | 1.08E+07 | 6.25E+06 | 8.77E+06 | 1.04E+07 | 8.32E+06 | 7.38E+06 | 8.40E+06 | 8.65E+06 | 0.97 | -0.04 | 0.781109985 |
| P62136 | Serine/threonine-protein phosphatase PP1-alpha catalytic subunit OS=Homo sapiens OX=9606 GN=PPP1CA PE=1 SV=1 | 44 | 160 | 3 | 4.28E+07 | 3.98E+07 | 4.04E+07 | 4.01E+07 | 4.45E+07 | 4.47E+07 | 4.19E+07 | 4.19E+07 | 2.97E+07 | 3.50E+07 | 3.25E+07 | 3.09E+07 | 4.21E+07 | 3.53E+07 | 1.19 | 0.25 | 0.026594184 |
| P20742 | Pregnancy zone protein OS=Homo sapiens OX=9606 GN=PZP PE=1 SV=4 | 6 | 2368 | 3 | 5.93E+08 | 4.52E+08 | 3.65E+08 | 4.71E+08 | 4.11E+08 | 6.98E+08 | 6.05E+08 | 5.68E+08 | 3.27E+08 | 3.76E+08 | 5.59E+08 | 5.40E+08 | 4.98E+08 | 4.96E+08 | 1.00 | 0.01 | 0.972491362 |
| P18085 | ADP-ribosylation factor 4 OS=Homo sapiens OX=9606 GN=ARF4 PE=1 SV=3 | 45 | 332 | 3 | 2.34E+07 | 2.46E+07 | 2.71E+07 | 3.04E+07 | 2.81E+07 | 2.50E+07 | 2.34E+07 | 2.29E+07 | 2.34E+07 | 2.47E+07 | 2.34E+07 | 2.03E+07 | 2.64E+07 | 2.30E+07 | 1.15 | 0.20 | 0.023175988 |
| Q9H0U4 | Ras-related protein Rab-1B OS=Homo sapiens OX=9606 GN=RAB1B PE=1 SV=1 | 58 | 123 | 3 | 7.61E+07 | 8.93E+07 | 7.79E+07 | 6.34E+07 | 6.94E+07 | 7.01E+07 | 6.29E+07 | 6.07E+07 | 4.33E+07 | 4.74E+07 | 5.42E+07 | 4.61E+07 | 7.44E+07 | 5.25E+07 | 1.42 | 0.50 | 0.001287557 |
| P35443 | Thrombospondin-4 OS=Homo sapiens OX=9606 GN=THBS4 PE=1 SV=2 | 9 | 168 | 3 | 8.44E+06 | 7.79E+06 | 6.97E+06 | 1.03E+07 | 7.00E+06 | 9.97E+06 | 1.47E+07 | 1.18E+07 | 8.34E+06 | 8.61E+06 | 8.61E+06 | 9.05E+06 | 8.41E+06 | 1.02E+07 | 0.83 | -0.28 | 0.178884902 |
| P62879 | Guanine nucleotide-binding protein G(I)/G(S)/G(T) subunit beta-2 OS=Homo sapiens OX=9606 GN=GNB2 PE=1 SV=3 | 27 | 206 | 3 | 6.34E+07 | 6.20E+07 | 5.83E+07 | 6.54E+07 | 6.18E+07 | 5.23E+07 | 6.54E+07 | 5.83E+07 | 5.55E+07 | 5.56E+07 | 4.78E+07 | 5.13E+07 | 6.05E+07 | 5.56E+07 | 1.09 | 0.12 | 0.148702095 |
| P08567 | Pleckstrin OS=Homo sapiens OX=9606 GN=PLEK PE=1 SV=3 | 10 | 40 | 3 | 2.12E+07 | 8.55E+06 | 1.36E+07 | 6.25E+06 | 1.93E+07 | 1.82E+07 | 3.75E+07 | 2.43E+07 | 1.36E+07 | 1.35E+07 | 1.81E+07 | 1.78E+07 | 1.45E+07 | 2.08E+07 | 0.70 | -0.52 | 0.193424095 |
| Q07021 | Complement component 1 Q subcomponent-binding protein, mitochondrial OS=Homo sapiens OX=9606 GN=C1QBP PE=1 SV=1 | 25 | 40 | 3 | 3.06E+06 | 2.77E+06 | 6.81E+06 | 4.91E+06 | 7.85E+06 | 6.38E+06 | 2.20E+06 | 5.69E+06 | 1.92E+06 | 1.90E+06 | 2.77E+06 | 1.91E+06 | 5.29E+06 | 2.73E+06 | 1.94 | 0.95 | 0.036029525 |
| P31153 | S-adenosylmethionine synthase isoform type-2 OS=Homo sapiens OX=9606 GN=MAT2A PE=1 SV=1 | 17 | 124 | 3 | 1.93E+07 | 2.27E+07 | 1.98E+07 | 2.10E+07 | 2.01E+07 | 1.93E+07 | 2.59E+07 | 2.38E+07 | 2.56E+07 | 1.56E+07 | 2.27E+07 | 2.40E+07 | 2.04E+07 | 2.29E+07 | 0.89 | -0.17 | 0.167193047 |
| P20020 | Plasma membrane calcium-transporting ATPase 1 OS=Homo sapiens OX=9606 GN=ATP2B1 PE=1 SV=4 | 8 | 62 | 3 | 6.19E+06 | 7.00E+06 | 6.35E+06 | 7.64E+06 | 6.93E+06 | 6.86E+06 | 4.62E+06 | 3.90E+06 | 4.73E+06 | 4.80E+06 | 4.14E+06 | 3.80E+06 | 6.83E+06 | 4.33E+06 | 1.58 | 0.66 | 4.977E-06 |
| P15169 | Carboxypeptidase N catalytic chain OS=Homo sapiens OX=9606 GN=CPN1 PE=1 SV=1 | 8 | 352 | 3 | 1.55E+08 | 9.48E+07 | 1.50E+08 | 9.23E+07 | 1.34E+08 | 1.22E+08 | 2.62E+08 | 1.32E+08 | 2.37E+08 | 1.39E+08 | 2.03E+08 | 2.10E+08 | 1.25E+08 | 1.97E+08 | 0.63 | -0.66 | 0.018125241 |
| Q9H2A2 | 2-aminomuconic semialdehyde dehydrogenase OS=Homo sapiens OX=9606 GN=ALDH8A1 PE=1 SV=1 | 10 | 97 | 3 | 2.31E+07 | 2.10E+07 | 1.22E+07 | 1.37E+07 | 1.22E+07 | 1.45E+07 | 2.75E+07 | 2.52E+07 | 1.48E+07 | 1.74E+07 | 1.82E+07 | 1.77E+07 | 1.61E+07 | 2.01E+07 | 0.80 | -0.32 | 0.184816247 |
| P35858 | Insulin-like growth factor-binding protein complex acid labile subunit OS=Homo sapiens OX=9606 GN=IGFALS PE=1 SV=1 | 4 | 99 | 3 | 1.93E+07 | 1.75E+07 | 1.84E+07 | 1.74E+07 | 1.57E+07 | 1.97E+07 | 1.97E+07 | 2.27E+07 | 1.56E+07 | 2.03E+07 | 1.73E+07 | 1.87E+07 | 1.80E+07 | 1.91E+07 | 0.95 | -0.08 | 0.407016903 |
| Q15417 | Calponin-3 OS=Homo sapiens OX=9606 GN=CNN3 PE=1 SV=1 | 21 | 26 | 3 | 3.52E+06 | 4.34E+06 | 8.02E+06 | 5.89E+06 | 5.16E+06 | 7.82E+06 | 4.58E+06 | 3.37E+06 | 4.60E+06 | 3.62E+06 | 3.19E+06 | 2.91E+06 | 5.79E+06 | 3.71E+06 | 1.56 | 0.64 | 0.038394801 |
| Q92747 | Actin-related protein 2/3 complex subunit 1A OS=Homo sapiens OX=9606 GN=ARPC1A PE=2 SV=2 | 17 | 44 | 3 | 9.27E+06 | 6.10E+06 | 8.02E+06 | 9.03E+06 | 7.54E+06 | 7.88E+06 | 1.00E+07 | 9.17E+06 | 9.54E+06 | 1.13E+07 | 5.31E+06 | 7.56E+06 | 7.97E+06 | 8.82E+06 | 0.90 | -0.15 | 0.412274795 |
| P43007 | Neutral amino acid transporter A OS=Homo sapiens OX=9606 GN=SLC1A4 PE=1 SV=1 | 12 | 36 | 3 | 4.56E+06 | 5.10E+06 | 1.15E+07 | 5.65E+06 | 5.21E+06 | 4.98E+06 | 5.70E+06 | 5.74E+06 | 5.09E+06 | 3.62E+06 | 3.03E+06 | 6.61E+06 | 6.17E+06 | 4.96E+06 | 1.24 | 0.31 | 0.352695405 |
| P00747 | Plasminogen OS=Homo sapiens OX=9606 GN=PLG PE=1 SV=2 | 6 | 80 | 3 | 2.67E+07 | 2.37E+07 | 3.02E+07 | 3.31E+07 | 3.00E+07 | 2.73E+07 | 4.01E+07 | 4.24E+07 | 4.21E+07 | 4.27E+07 | 3.47E+07 | 3.20E+07 | 2.85E+07 | 3.90E+07 | 0.73 | -0.45 | 0.001267297 |
| P12277 | Creatine kinase B-type OS=Homo sapiens OX=9606 GN=CKB PE=1 SV=1 | 18 | 28 | 3 | 3.69E+06 | 4.36E+05 | 1.40E+06 | 1.56E+06 | 1.76E+06 | 2.26E+06 | 2.96E+06 | 1.22E+06 | 1.67E+06 | 4.66E+06 | 1.42E+06 | 2.91E+06 | 1.85E+06 | 2.47E+06 | 0.75 | -0.42 | 0.393617959 |

### Supplementary Table 4. List of all proteins identified by label free proteomics analysis.

| Accession | Description | Coverage [%] | # PSMs | # Unique Peptides | Amitriptyline 1 | Amitriptyline 2 | Amitriptyline 3 | Amitriptyline 4 | Amitriptyline 5 | Amitriptyline 6 | Vehicle 1 | Vehicle 2 | Vehicle 3 | Vehicle 4 | Vehicle 5 | Vehicle 6 | Average amitriptyline | Average vehicle | Ratio amitriptyline/ vehicle | Log 2 fold change (amitriptyline /vehicle) | P-value |
|---|---|---|---|---|---|---|---|---|---|---|---|---|---|---|---|---|---|---|---|---|---|
| P63000 | Ras-related C3 botulinum toxin substrate 1 OS=Homo sapiens OX=9606 GN=RAC1 PE=1 SV=1 | 32 | 167 | 3 | 1.79E+07 | 1.57E+07 | 2.01E+07 | 1.65E+07 | 2.04E+07 | 2.06E+07 | 1.59E+07 | 1.37E+07 | 1.34E+07 | 2.28E+07 | 1.93E+07 | 1.74E+07 | 1.85E+07 | 1.71E+07 | 1.08 | 0.12 | 0.42026623 |
| P62841 | 40S ribosomal protein S15 OS=Homo sapiens OX=9606 GN=RPS15 PE=1 SV=2 | 47 | 26 | 3 | 3.27E+06 | 5.01E+06 | 8.01E+06 | 7.10E+06 | 1.06E+07 | 6.31E+06 | 1.46E+06 | 2.96E+06 | 1.61E+06 | 1.78E+06 | 2.61E+06 | 1.79E+06 | 6.71E+06 | 2.04E+06 | 3.30 | 1.72 | 0.005324168 |
| Q15404 | Ras suppressor protein 1 OS=Homo sapiens OX=9606 GN=RSU1 PE=1 SV=3 | 18 | 29 | 3 | 7.41E+06 | 4.48E+06 | 4.96E+06 | 4.55E+06 | 5.36E+06 | 7.44E+06 | 1.04E+07 | 6.71E+06 | 5.89E+06 | 5.43E+06 | 5.33E+06 | 8.81E+06 | 5.70E+06 | 7.09E+06 | 0.80 | -0.31 | 0.20287752 |
| P31949 | Protein S100-A11 OS=Homo sapiens OX=9606 GN=S100A11 PE=1 SV=2 | 54 | 24 | 3 | 8.61E+06 | 1.14E+07 | 9.65E+06 | 1.50E+07 | 1.41E+07 | 1.13E+07 | 7.54E+06 | 6.68E+06 | 7.26E+06 | 6.10E+06 | 6.39E+06 | 5.01E+06 | 1.17E+07 | 6.50E+06 | 1.80 | 0.85 | 0.002666145 |
| P84095 | Rho-related GTP-binding protein RhoG OS=Homo sapiens OX=9606 GN=RHOG PE=1 SV=1 | 24 | 33 | 3 | 4.73E+06 | 5.01E+06 | 3.07E+06 | 2.99E+06 | 3.57E+06 | 4.28E+06 | 3.83E+06 | 3.86E+06 | 3.58E+06 | 3.38E+06 | 2.18E+06 | 3.15E+06 | 3.94E+06 | 3.33E+06 | 1.18 | 0.24 | 0.191194835 |
| Q9UBF2 | Coatomer subunit gamma-2 OS=Homo sapiens OX=9606 GN=COPG2 PE=1 SV=1 | 8 | 40 | 3 | 6.94E+05 | 2.83E+05 | 8.91E+05 | 6.19E+05 | 1.25E+06 | 5.74E+05 | | | | 3.02E+05 | | | 7.18E+05 | 3.02E+05 | 2.38 | 1.25 | #DIV/0! |
| Q9GZS3 | WD repeat-containing protein 61 OS=Homo sapiens OX=9606 GN=WDR61 PE=1 SV=1 | 23 | 16 | 3 | 2.14E+06 | 5.11E+06 | 2.03E+06 | 4.60E+06 | 3.37E+06 | 6.92E+06 | 2.87E+06 | 1.57E+06 | 2.31E+06 | 1.58E+06 | 4.04E+06 | 2.78E+06 | 4.03E+06 | 2.53E+06 | 1.59 | 0.67 | 0.122496316 |
| P15927 | Replication protein A 32 kDa subunit OS=Homo sapiens OX=9606 GN=RPA2 PE=1 SV=1 | 22 | 56 | 3 | 1.06E+07 | 9.71E+06 | 9.30E+06 | 1.04E+07 | 9.73E+06 | 1.28E+07 | 1.02E+07 | 9.23E+06 | 6.87E+06 | 9.28E+06 | 1.10E+07 | 5.91E+06 | 1.04E+07 | 8.76E+06 | 1.19 | 0.25 | 0.117672167 |
| P62330 | ADP-ribosylation factor 6 OS=Homo sapiens OX=9606 GN=ARF6 PE=1 SV=2 | 26 | 42 | 3 | 2.15E+06 | 1.06E+07 | 7.30E+06 | 4.05E+06 | 7.36E+06 | 4.72E+06 | 8.79E+06 | 9.68E+06 | 7.09E+06 | 4.89E+06 | 1.00E+07 | 8.31E+06 | 6.03E+06 | 8.13E+06 | 0.74 | -0.43 | 0.184684461 |
| Q8TAQ2 | SWI/SNF complex subunit SMARCC2 OS=Homo sapiens OX=9606 GN=SMARCC2 PE=1 SV=1 | 4 | 14 | 3 | 6.48E+06 | 5.92E+06 | 5.77E+06 | 7.20E+06 | 4.67E+06 | 5.94E+06 | 2.73E+06 | 2.63E+06 | 2.21E+06 | 2.82E+06 | 3.00E+06 | 2.74E+06 | 6.00E+06 | 2.69E+06 | 2.23 | 1.16 | 9.06918E-05 |
| P21810 | Biglycan OS=Homo sapiens OX=9606 GN=BGN PE=1 SV=2 | 13 | 34 | 3 | 4.09E+06 | 4.35E+06 | 3.99E+06 | 5.36E+06 | 4.59E+06 | 6.28E+06 | 1.21E+07 | 5.59E+06 | 4.99E+06 | 8.68E+06 | 6.73E+06 | 7.98E+06 | 4.78E+06 | 7.68E+06 | 0.62 | -0.68 | 0.038928016 |
| Q9UHX1 | Poly(U)-binding-splicing factor PUF60 OS=Homo sapiens OX=9606 GN=PUF60 PE=1 SV=1 | 12 | 18 | 3 | 3.79E+06 | 3.77E+06 | 2.48E+06 | 2.61E+06 | 2.48E+06 | 3.11E+06 | 2.28E+06 | 2.02E+06 | 1.27E+06 | 1.26E+06 | 1.45E+06 | 1.35E+06 | 3.04E+06 | 1.61E+06 | 1.89 | 0.92 | 0.001200311 |
| P42285 | Exosome RNA helicase MTR4 OS=Homo sapiens OX=9606 GN=MTREX PE=1 SV=3 | 5 | 12 | 3 | 3.09E+06 | 3.37E+06 | 4.06E+06 | 4.75E+06 | 5.08E+06 | 3.52E+06 | 1.65E+06 | 1.33E+06 | 1.20E+06 | 1.65E+06 | | 6.84E+05 | 3.98E+06 | 1.30E+06 | 3.06 | 1.61 | 0.000117013 |
| P21291 | Cysteine and glycine-rich protein 1 OS=Homo sapiens OX=9606 GN=CSRP1 PE=1 SV=3 | 25 | 62 | 3 | 1.04E+07 | 9.40E+06 | 8.88E+06 | 7.95E+06 | 8.89E+06 | 9.35E+06 | 1.14E+07 | 9.83E+06 | 6.78E+06 | 9.81E+06 | 8.28E+06 | 8.63E+06 | 9.15E+06 | 9.11E+06 | 1.00 | 0.01 | 0.960538045 |
| P17858 | ATP-dependent 6-phosphofructokinase, liver type OS=Homo sapiens OX=9606 GN=PFKL PE=1 SV=6 | 9 | 49 | 3 | 1.82E+07 | 1.79E+07 | 1.92E+07 | 1.88E+07 | 1.73E+07 | 1.68E+07 | 2.21E+07 | 2.23E+07 | 2.23E+07 | 2.20E+07 | 1.92E+07 | 1.73E+07 | 1.80E+07 | 2.09E+07 | 0.86 | -0.21 | 0.020948453 |
| P32970 | CD70 antigen OS=Homo sapiens OX=9606 GN=CD70 PE=1 SV=2 | 20 | 30 | 3 | 1.86E+07 | 1.13E+07 | 1.19E+07 | 1.16E+07 | 1.21E+07 | 1.14E+07 | 1.12E+07 | 8.49E+06 | 6.30E+06 | 5.81E+06 | 8.42E+06 | 9.84E+06 | 1.28E+07 | 8.34E+06 | 1.54 | 0.62 | 0.01192165 |
| P62304 | Small nuclear ribonucleoprotein E OS=Homo sapiens OX=9606 GN=SNRPE PE=1 SV=1 | 52 | 155 | 3 | 2.05E+07 | 1.82E+07 | 1.41E+07 | 1.18E+07 | 1.39E+07 | 2.10E+07 | 1.32E+07 | 1.50E+07 | 1.21E+07 | 1.20E+07 | 1.17E+07 | 1.49E+07 | 1.66E+07 | 1.31E+07 | 1.26 | 0.34 | 0.082594046 |
| P42892 | Endothelin-converting enzyme 1 OS=Homo sapiens OX=9606 GN=ECE1 PE=1 SV=2 | 7 | 31 | 3 | 5.52E+06 | 6.38E+06 | 3.66E+06 | 5.03E+06 | 5.27E+06 | 6.74E+06 | 4.78E+06 | 4.95E+06 | 1.77E+06 | 2.15E+06 | 4.78E+06 | 1.90E+06 | 5.43E+06 | 3.39E+06 | 1.60 | 0.68 | 0.029490098 |
| P08708 | 40S ribosomal protein S17 OS=Homo sapiens OX=9606 GN=RPS17 PE=1 SV=2 | 29 | 14 | 3 | 1.13E+07 | 1.01E+07 | 7.76E+06 | 8.73E+06 | 9.87E+06 | 9.01E+06 | 6.50E+06 | 4.98E+06 | 3.76E+06 | 4.95E+06 | 5.07E+06 | 5.15E+06 | 9.46E+06 | 5.07E+06 | 1.87 | 0.90 | 5.49507E-05 |
| O14828 | Secretory carrier-associated membrane protein 3 OS=Homo sapiens OX=9606 GN=SCAMP3 PE=1 SV=3 | 16 | 26 | 3 | 9.22E+06 | 8.90E+06 | 6.76E+06 | 5.95E+06 | 5.74E+06 | 5.75E+06 | 6.17E+06 | 4.88E+06 | 3.37E+06 | 4.07E+06 | 3.78E+06 | 6.36E+05 | 7.05E+06 | 3.82E+06 | 1.85 | 0.89 | 0.009062845 |
| Q99715 | Collagen alpha-1(XII) chain OS=Homo sapiens OX=9606 GN=COL12A1 PE=1 SV=2 | 2 | 18 | 3 | 2.71E+06 | 2.87E+06 | 1.81E+06 | 1.95E+06 | 1.27E+06 | 1.85E+06 | 4.92E+06 | 1.88E+06 | 3.35E+06 | 4.70E+06 | 5.32E+06 | 4.21E+06 | 2.08E+06 | 4.06E+06 | 0.51 | -0.97 | 0.010035328 |
| P54578 | Ubiquitin carboxyl-terminal hydrolase 14 OS=Homo sapiens OX=9606 GN=USP14 PE=1 SV=3 | 11 | 13 | 3 | 4.88E+06 | 8.00E+05 | 3.31E+06 | 4.02E+06 | 4.04E+06 | 3.42E+06 | 4.46E+06 | 9.48E+05 | 2.06E+06 | 4.26E+06 | 3.44E+06 | 2.48E+06 | 3.41E+06 | 2.94E+06 | 1.16 | 0.21 | 0.567349181 |
| Q13443 | Disintegrin and metalloproteinase domain-containing protein 9 OS=Homo sapiens OX=9606 GN=ADAM9 PE=1 SV=1 | 7 | 11 | 3 | 6.02E+06 | 3.13E+06 | 4.14E+06 | 4.21E+06 | 5.99E+06 | 5.46E+06 | 1.78E+06 | 2.37E+06 | 1.52E+06 | 2.39E+06 | 1.23E+06 | 1.82E+06 | 4.82E+06 | 1.85E+06 | 2.61 | 1.38 | 0.000883313 |
| P48735 | Isocitrate dehydrogenase [NADP], mitochondrial OS=Homo sapiens OX=9606 GN=IDH2 PE=1 SV=2 | 12 | 16 | 3 | 8.07E+06 | 7.62E+06 | 5.75E+06 | 5.16E+06 | 7.08E+06 | 5.65E+06 | 5.15E+06 | 2.95E+06 | 3.15E+06 | 4.54E+06 | 4.41E+06 | 2.38E+06 | 6.56E+06 | 3.76E+06 | 1.74 | 0.80 | 0.001747295 |
| Q99627 | COP9 signalosome complex subunit 8 OS=Homo sapiens OX=9606 GN=COPS8 PE=1 SV=1 | 28 | 18 | 3 | 1.59E+06 | 1.82E+06 | 9.21E+05 | 9.16E+05 | 7.90E+05 | 1.01E+06 | 7.07E+05 | | | | 6.16E+05 | | 1.17E+06 | 6.62E+05 | 1.77 | 0.83 | 0.030582465 |
| Q86Y82 | Syntaxin-12 OS=Homo sapiens OX=9606 GN=STX12 PE=1 SV=1 | 15 | 46 | 3 | 7.71E+06 | 6.97E+06 | 8.61E+06 | 1.01E+07 | 8.83E+06 | 6.06E+06 | 5.94E+06 | 4.74E+06 | 6.24E+06 | 5.87E+06 | 4.26E+06 | 4.68E+06 | 8.05E+06 | 5.29E+06 | 1.52 | 0.61 | 0.003671413 |
| Q12906 | Interleukin enhancer-binding factor 3 OS=Homo sapiens OX=9606 GN=ILF3 PE=1 SV=3 | 8 | 14 | 3 | 4.27E+06 | 5.29E+06 | 6.83E+06 | 4.64E+06 | 5.31E+06 | 3.90E+06 | 2.77E+06 | 2.79E+06 | 2.01E+06 | 2.61E+06 | 2.40E+06 | 2.74E+06 | 5.04E+06 | 2.55E+06 | 1.97 | 0.98 | 0.001455152 |
| P14923 | Junction plakoglobin OS=Homo sapiens OX=9606 GN=JUP PE=1 SV=3 | 10 | 39 | 3 | 3.99E+06 | 5.24E+06 | 2.88E+06 | 3.26E+06 | 5.90E+06 | 3.71E+06 | 1.97E+06 | 5.34E+06 | 1.43E+06 | 3.03E+06 | 2.72E+06 | 2.13E+06 | 4.16E+06 | 2.77E+06 | 1.50 | 0.59 | 0.089176842 |
| P02748 | Complement component C9 OS=Homo sapiens OX=9606 GN=C9 PE=1 SV=2 | 4 | 132 | 3 | 6.27E+07 | 6.17E+07 | 4.65E+07 | 5.40E+07 | 4.64E+07 | 5.22E+07 | 8.79E+07 | 8.75E+07 | 6.38E+07 | 6.45E+07 | 6.17E+07 | 6.05E+07 | 5.39E+07 | 7.10E+07 | 0.76 | -0.40 | 0.023226034 |
| Q13425 | Beta-2-syntrophin OS=Homo sapiens OX=9606 GN=SNTB2 PE=1 SV=1 | 10 | 18 | 3 | 4.64E+06 | 5.69E+06 | 4.10E+06 | 3.35E+06 | 3.46E+06 | 4.92E+06 | 3.22E+06 | 4.35E+06 | 1.65E+06 | 1.92E+06 | 3.20E+06 | 2.69E+06 | 4.36E+06 | 2.84E+06 | 1.54 | 0.62 | 0.019035898 |
| P62318 | Small nuclear ribonucleoprotein Sm D3 OS=Homo sapiens OX=9606 GN=SNRPD3 PE=1 SV=1 | 32 | 68 | 3 | 3.52E+07 | 3.71E+07 | 3.50E+07 | 3.16E+07 | 4.61E+07 | 5.00E+07 | 2.68E+07 | 2.82E+07 | 2.68E+07 | 2.99E+07 | 2.18E+07 | 2.23E+07 | 3.92E+07 | 2.60E+07 | 1.51 | 0.59 | 0.00469065 |
| P12821 | Angiotensin-converting enzyme OS=Homo sapiens OX=9606 GN=ACE PE=1 SV=1 | 2 | 20 | 3 | 9.35E+06 | 8.17E+06 | 4.80E+06 | 8.74E+06 | 7.11E+06 | 6.21E+06 | 9.16E+06 | 7.56E+06 | 6.54E+06 | 6.35E+06 | 7.71E+06 | 5.37E+06 | 7.40E+06 | 7.12E+06 | 1.04 | 0.06 | 0.756807776 |
| P52788 | Spermine synthase OS=Homo sapiens OX=9606 GN=SMS PE=1 SV=2 | 20 | 10 | 3 | 1.42E+06 | 1.07E+06 | 6.34E+05 | 1.06E+06 | 8.10E+05 | 1.64E+06 | 1.33E+06 | | | 5.46E+05 | 8.68E+05 | 6.52E+05 | 1.11E+06 | 8.49E+05 | 1.30 | 0.38 | 0.304421139 |
| P56537 | Eukaryotic translation initiation factor 6 OS=Homo sapiens OX=9606 GN=EIF6 PE=1 SV=1 | 22 | 20 | 3 | 1.09E+07 | 1.14E+07 | 7.61E+06 | 9.90E+06 | 8.97E+06 | 1.01E+07 | 8.15E+06 | 8.81E+06 | 8.57E+06 | 7.42E+06 | 8.71E+06 | 6.76E+06 | 9.82E+06 | 8.07E+06 | 1.22 | 0.28 | 0.027317347 |
| P25325 | 3-mercaptopyruvate sulfurtransferase OS=Homo sapiens OX=9606 GN=MPST PE=1 SV=3 | 16 | 23 | 3 | 4.51E+06 | 8.40E+06 | 2.51E+06 | 3.48E+06 | 3.86E+06 | 5.10E+06 | 5.29E+06 | 4.51E+06 | 2.40E+06 | 5.08E+06 | 4.45E+06 | 4.96E+06 | 4.64E+06 | 4.45E+06 | 1.04 | 0.06 | 0.841475641 |
| P62070 | Ras-related protein R-Ras2 OS=Homo sapiens OX=9606 GN=RRAS2 PE=1 SV=1 | 21 | 56 | 3 | 1.52E+07 | 1.58E+07 | 1.69E+07 | 1.71E+07 | 1.59E+07 | 1.48E+07 | 1.20E+07 | 1.18E+07 | 5.86E+06 | 1.33E+07 | 1.19E+07 | 8.65E+06 | 1.60E+07 | 1.06E+07 | 1.51 | 0.59 | 0.004012113 |
| P28066 | Proteasome subunit alpha type-5 OS=Homo sapiens OX=9606 GN=PSMA5 PE=1 SV=3 | 17 | 48 | 3 | 2.50E+07 | 2.49E+07 | 2.12E+07 | 2.53E+07 | 2.56E+07 | 7.09E+06 | 6.80E+06 | 7.02E+06 | 7.89E+06 | 6.72E+06 | 6.19E+06 | 5.17E+06 | 2.15E+07 | 6.63E+06 | 3.24 | 1.70 | 0.003796693 |
| P28062 | Proteasome subunit beta type-8 OS=Homo sapiens OX=9606 GN=PSMB8 PE=1 SV=3 | 13 | 36 | 3 | 9.14E+06 | 7.92E+06 | 1.14E+07 | 1.01E+07 | 1.03E+07 | 9.85E+06 | 7.86E+06 | 9.38E+06 | 8.91E+06 | 9.82E+06 | 8.68E+06 | 7.70E+06 | 9.77E+06 | 8.72E+06 | 1.12 | 0.16 | 0.104466351 |
| Q15366 | Poly(rC)-binding protein 2 OS=Homo sapiens OX=9606 GN=PCBP2 PE=1 SV=1 | 17 | 68 | 3 | 1.36E+07 | 1.16E+07 | 9.14E+06 | 9.14E+06 | 8.63E+06 | 9.61E+06 | 9.56E+06 | 5.60E+06 | 5.48E+06 | 8.69E+06 | 5.86E+06 | 7.08E+06 | 1.03E+07 | 7.04E+06 | 1.46 | 0.55 | 0.011962552 |
| P62995 | Transformer-2 protein homolog beta OS=Homo sapiens OX=9606 GN=TRA2B PE=1 SV=1 | 13 | 36 | 3 | 9.27E+06 | 1.01E+07 | 1.05E+07 | 1.26E+07 | 1.03E+07 | 1.01E+07 | 9.34E+06 | 1.21E+07 | 1.05E+07 | 1.07E+07 | 1.22E+07 | 7.62E+06 | 1.05E+07 | 1.04E+07 | 1.00 | 0.01 | 0.964546162 |
| Q9BT78 | COP9 signalosome complex subunit 4 OS=Homo sapiens OX=9606 GN=COPS4 PE=1 SV=1 | 13 | 30 | 3 | 5.20E+06 | 5.59E+06 | 4.81E+06 | 5.14E+06 | 4.67E+06 | 4.07E+06 | 4.67E+06 | 4.81E+06 | 4.62E+06 | 3.31E+06 | 4.76E+06 | 2.20E+06 | 4.91E+06 | 4.06E+06 | 1.21 | 0.27 | 0.123988575 |
| P24752 | Acetyl-CoA acetyltransferase, mitochondrial OS=Homo sapiens OX=9606 GN=ACAT1 PE=1 SV=1 | 11 | 14 | 3 | 5.76E+06 | 3.55E+06 | 4.13E+06 | 4.34E+06 | 3.78E+06 | 3.40E+06 | 3.80E+06 | 2.60E+06 | 2.36E+06 | 3.48E+06 | 9.22E+05 | 1.91E+06 | 4.16E+06 | 2.51E+06 | 1.66 | 0.73 | 0.014460507 |
| Q14498 | RNA-binding protein 39 OS=Homo sapiens OX=9606 GN=RBM39 PE=1 SV=2 | 8 | 28 | 3 | 3.47E+06 | 3.25E+06 | 4.14E+06 | 5.20E+06 | 3.96E+06 | 3.94E+06 | 1.82E+06 | 1.99E+06 | 1.69E+06 | 2.31E+06 | 1.44E+06 | 6.58E+05 | 3.99E+06 | 1.65E+06 | 2.42 | 1.27 | 8.35261E-05 |
| P60660 | Myosin light polypeptide 6 OS=Homo sapiens OX=9606 GN=MYL6 PE=1 SV=2 | 25 | 70 | 3 | 2.36E+07 | 2.42E+07 | 2.38E+07 | 2.49E+07 | 2.46E+07 | 2.26E+07 | 1.82E+07 | 2.06E+07 | 1.87E+07 | 2.20E+07 | 2.00E+07 | 1.99E+07 | 2.39E+07 | 1.99E+07 | 1.20 | 0.27 | 0.00023778 |
| P51570 | Galactokinase OS=Homo sapiens OX=9606 GN=GALK1 PE=1 SV=1 | 8 | 187 | 3 | 6.26E+07 | 7.38E+07 | 6.10E+07 | 5.45E+07 | 5.89E+07 | 5.70E+07 | 8.83E+07 | 9.21E+07 | 8.21E+07 | 8.70E+07 | 9.88E+07 | 6.72E+07 | 6.13E+07 | 8.59E+07 | 0.71 | -0.49 | 0.001252825 |
| Q06828 | Fibromodulin OS=Homo sapiens OX=9606 GN=FMOD PE=1 SV=2 | 9 | 76 | 3 | 1.52E+07 | 1.40E+07 | 1.14E+07 | 1.51E+07 | 1.54E+07 | 1.70E+07 | 1.92E+07 | 2.41E+07 | 1.89E+07 | 1.72E+07 | 1.94E+07 | 1.66E+07 | 1.47E+07 | 1.92E+07 | 0.76 | -0.39 | 0.007278878 |
| Q16822 | Phosphoenolpyruvate carboxykinase [GTP], mitochondrial OS=Homo sapiens OX=9606 GN=PCK2 PE=1 SV=4 | 6 | 24 | 3 | 4.59E+06 | 3.85E+06 | 3.89E+06 | 4.80E+06 | 4.64E+06 | 5.28E+06 | 3.83E+06 | 9.54E+05 | 1.75E+06 | 2.22E+06 | 2.30E+06 | 2.64E+06 | 4.51E+06 | 2.28E+06 | 1.97 | 0.98 | 0.00113164 |
| Q8NFW8 | N-acylneuraminate cytidylyltransferase OS=Homo sapiens OX=9606 GN=CMAS PE=1 SV=2 | 10 | 30 | 3 | 7.39E+06 | 8.88E+06 | 7.60E+06 | 8.82E+06 | 4.68E+06 | 7.06E+06 | 4.70E+06 | 5.79E+06 | 6.25E+06 | 6.44E+06 | 6.90E+06 | 5.80E+06 | 7.40E+06 | 5.98E+06 | 1.24 | 0.31 | 0.078832666 |
| P24534 | Elongation factor 1-beta OS=Homo sapiens OX=9606 GN=EEF1B2 PE=1 SV=3 | 16 | 42 | 3 | 1.08E+07 | 1.17E+07 | 1.48E+07 | 1.08E+07 | 1.07E+07 | 1.34E+07 | 9.42E+06 | 1.03E+07 | 1.01E+07 | 7.86E+06 | 8.19E+06 | 9.06E+06 | 1.20E+07 | 9.16E+06 | 1.31 | 0.39 | 0.007319431 |
| Q93088 | Betaine--homocysteine S-methyltransferase 1 OS=Homo sapiens OX=9606 GN=BHMT PE=1 SV=2 | 12 | 34 | 3 | 4.15E+06 | 4.29E+06 | 4.55E+06 | 4.46E+06 | 3.69E+06 | 3.26E+06 | 5.98E+06 | 5.90E+06 | 5.62E+06 | 5.52E+06 | 5.57E+06 | 4.22E+06 | 4.07E+06 | 5.47E+06 | 0.74 | -0.43 | 0.001907594 |
| Q14344 | Guanine nucleotide-binding protein subunit alpha-13 OS=Homo sapiens OX=9606 GN=GNA13 PE=1 SV=2 | 16 | 133 | 3 | 3.73E+06 | 3.80E+06 | 3.37E+06 | 4.01E+06 | 3.33E+06 | 1.92E+06 | 7.67E+05 | 8.10E+05 | 6.35E+05 | | 9.18E+05 | 5.27E+05 | 3.36E+06 | 7.31E+05 | 4.59 | 2.20 | 0.00025449 |
| P42357 | Histidine ammonia-lyase OS=Homo sapiens OX=9606 GN=HAL PE=1 SV=1 | 9 | 17 | 3 | 9.51E+05 | 2.44E+06 | 1.65E+06 | 2.54E+06 | 3.19E+06 | 2.51E+06 | 2.04E+06 | 1.81E+06 | 2.50E+06 | 2.06E+06 | 4.44E+06 | 1.40E+06 | 2.21E+06 | 2.38E+06 | 0.93 | -0.10 | 0.773003589 |
| P06753 | Tropomyosin alpha-3 chain OS=Homo sapiens OX=9606 GN=TPM3 PE=1 SV=2 | 16 | 16 | 3 | 3.55E+06 | 1.82E+06 | 3.35E+06 | 3.65E+06 | 2.68E+06 | 1.59E+06 | 1.47E+06 | 1.08E+06 | 1.64E+06 | 8.20E+05 | 1.48E+06 | | 2.78E+06 | 1.30E+06 | 2.14 | 1.09 | 0.008213688 |
| Q15043 | Zinc transporter ZIP14 OS=Homo sapiens OX=9606 GN=SLC39A14 PE=1 SV=3 | 9 | 18 | 3 | 2.23E+06 | 3.02E+06 | 3.21E+06 | 2.99E+06 | 3.05E+06 | 4.53E+06 | 1.55E+06 | 1.54E+06 | 1.60E+06 | 1.09E+06 | 1.56E+06 | 1.38E+06 | 3.17E+06 | 1.45E+06 | 2.18 | 1.13 | 0.001938642 |
| Q14258 | E3 ubiquitin/ISG15 ligase TRIM25 OS=Homo sapiens OX=9606 GN=TRIM25 PE=1 SV=2 | 7 | 26 | 3 | 4.94E+06 | 7.26E+06 | 6.45E+06 | 4.37E+06 | 6.24E+06 | 6.73E+06 | 2.70E+06 | 3.11E+06 | 4.68E+06 | 4.62E+06 | 2.41E+06 | 3.82E+06 | 6.00E+06 | 3.56E+06 | 1.69 | 0.75 | 0.002399013 |
| P30876 | DNA-directed RNA polymerase II subunit RPB2 OS=Homo sapiens OX=9606 GN=POLR2B PE=1 SV=1 | 4 | 16 | 3 | 3.38E+06 | 2.89E+06 | 2.35E+06 | 3.11E+06 | 3.10E+06 | 2.10E+06 | 1.79E+06 | 1.26E+06 | 1.07E+06 | 1.30E+06 | 1.35E+06 | 8.47E+05 | 2.82E+06 | 1.27E+06 | 2.23 | 1.15 | 0.000144073 |
| P50213 | Isocitrate dehydrogenase [NAD] subunit alpha, mitochondrial OS=Homo sapiens OX=9606 GN=IDH3A PE=1 SV=1 | 9 | 44 | 3 | 7.38E+06 | 6.77E+06 | 5.93E+06 | 7.00E+06 | 6.61E+06 | 6.31E+06 | 4.43E+06 | 4.86E+06 | 3.82E+06 | 4.38E+06 | 4.07E+06 | 3.60E+06 | 6.66E+06 | 4.19E+06 | 1.59 | 0.67 | 5.50661E-06 |
| P26368 | Splicing factor U2AF 65 kDa subunit OS=Homo sapiens OX=9606 GN=U2AF2 PE=1 SV=4 | 9 | 20 | 3 | 7.50E+06 | 6.12E+06 | 4.55E+06 | 4.56E+06 | 4.30E+06 | 7.44E+06 | 2.80E+06 | 2.74E+06 | 1.22E+06 | 6.51E+05 | 1.01E+06 | 2.06E+06 | 5.75E+06 | 1.75E+06 | 3.29 | 1.72 | 0.000434999 |
| Q9UBQ5 | Eukaryotic translation initiation factor 3 subunit K OS=Homo sapiens OX=9606 GN=EIF3K PE=1 SV=1 | 13 | 48 | 3 | 1.11E+07 | 1.30E+07 | 1.14E+07 | 9.71E+06 | 1.12E+07 | 1.12E+07 | 5.96E+06 | 8.94E+06 | 6.15E+06 | 5.96E+06 | 5.50E+06 | 6.26E+06 | 1.13E+07 | 6.46E+06 | 1.74 | 0.80 | 3.12453E-05 |
| O00743 | Serine/threonine-protein phosphatase 6 catalytic subunit OS=Homo sapiens OX=9606 GN=PPP6C PE=1 SV=1 | 11 | 23 | 3 | 7.37E+06 | 8.13E+06 | 4.96E+06 | 4.59E+06 | 6.09E+06 | 4.63E+06 | 5.66E+06 | 5.41E+06 | 3.98E+06 | 4.36E+06 | 4.42E+06 | 4.88E+06 | 5.96E+06 | 4.78E+06 | 1.25 | 0.32 | 0.123621273 |
| Q01581 | Hydroxymethylglutaryl-CoA synthase, cytoplasmic OS=Homo sapiens OX=9606 GN=HMGCS1 PE=1 SV=2 | 8 | 12 | 3 | 3.44E+06 | 2.87E+06 | 2.07E+06 | 3.21E+06 | 3.50E+06 | 3.33E+06 | 3.84E+06 | 3.10E+06 | 2.46E+06 | 3.01E+06 | 4.17E+06 | 3.08E+06 | 3.07E+06 | 3.28E+06 | 0.94 | -0.09 | 0.551378719 |
| Q8WXF1 | Paraspeckle component 1 OS=Homo sapiens OX=9606 GN=PSPC1 PE=1 SV=1 | 9 | 34 | 3 | 2.04E+06 | 1.41E+06 | 2.19E+06 | 3.01E+06 | 1.66E+06 | 1.87E+06 | 1.92E+06 | 8.06E+05 | 1.06E+06 | | 1.06E+06 | 1.07E+06 | 2.03E+06 | 1.18E+06 | 1.71 | 0.78 | 0.018844376 |
| Q9H4G4 | Golgi-associated plant pathogenesis-related protein 1 OS=Homo sapiens OX=9606 GN=GLIPR2 PE=1 SV=3 | 31 | 46 | 3 | 4.73E+06 | 5.24E+06 | 4.39E+06 | 5.07E+06 | 4.68E+06 | 4.59E+06 | 3.83E+06 | 4.21E+06 | 4.69E+06 | 4.85E+06 | 3.89E+06 | 3.52E+06 | 4.78E+06 | 4.16E+06 | 1.15 | 0.20 | 0.036839379 |
| O43681 | ATPase ASNA1 OS=Homo sapiens OX=9606 GN=ASNA1 PE=1 SV=2 | 11 | 38 | 3 | 5.02E+06 | 8.14E+06 | 7.14E+06 | 6.94E+06 | 4.99E+06 | 6.08E+06 | 4.61E+06 | 5.92E+06 | 4.61E+06 | 4.63E+06 | 4.02E+06 | 5.95E+06 | 6.39E+06 | 4.96E+06 | 1.29 | 0.37 | 0.044075559 |
| Q9Y266 | Nuclear migration protein nudC OS=Homo sapiens OX=9606 GN=NUDC PE=1 SV=1 | 12 | 26 | 3 | 9.67E+06 | 1.09E+07 | 1.25E+07 | 9.19E+06 | 1.11E+07 | 7.72E+06 | 5.47E+06 | 6.95E+06 | 5.23E+06 | 7.00E+06 | 4.94E+06 | 3.78E+06 | 1.02E+07 | 5.56E+06 | 1.83 | 0.87 | 0.000378411 |
| P24928 | DNA-directed RNA polymerase II subunit RPB1 OS=Homo sapiens OX=9606 GN=POLR2A PE=1 SV=2 | 2 | 7 | 3 | 1.83E+06 | 1.39E+06 | 2.05E+06 | 2.48E+06 | 2.34E+06 | 1.59E+06 | | 1.85E+06 | 1.86E+06 | 1.63E+06 | 1.77E+06 | 1.34E+06 | 1.95E+06 | 1.69E+06 | 1.15 | 0.20 | 0.234559605 |
| Q7L5N1 | COP9 signalosome complex subunit 6 OS=Homo sapiens OX=9606 GN=COPS6 PE=1 SV=1 | 17 | 23 | 3 | 6.92E+06 | 8.18E+06 | 7.36E+06 | 5.36E+06 | 7.01E+06 | 4.40E+06 | 7.10E+06 | 4.36E+06 | 4.23E+06 | 6.48E+06 | 4.76E+06 | 5.25E+06 | 6.54E+06 | 5.36E+06 | 1.22 | 0.29 | 0.146007342 |
| O95372 | Acyl-protein thioesterase 2 OS=Homo sapiens OX=9606 GN=LYPLA2 PE=1 SV=1 | 18 | 13 | 3 | 2.36E+06 | 4.70E+06 | 5.16E+06 | 3.54E+06 | 4.05E+06 | 5.29E+06 | 6.12E+06 | 5.87E+06 | 6.71E+06 | 3.72E+06 | 5.30E+06 | 4.22E+06 | 4.18E+06 | 5.32E+06 | 0.79 | -0.35 | 0.112929322 |
| P19784 | Casein kinase II subunit alpha' OS=Homo sapiens OX=9606 GN=CSNK2A2 PE=1 SV=1 | 12 | 15 | 3 | 6.34E+07 | 1.48E+08 | 6.95E+07 | 4.96E+07 | 6.84E+07 | 1.19E+08 | 8.41E+07 | 1.94E+08 | 3.99E+06 | 4.54E+07 | 1.43E+08 | 1.00E+08 | 8.63E+07 | 9.50E+07 | 0.91 | -0.14 | 0.790388108 |
| Q05519 | Serine/arginine-rich splicing factor 11 OS=Homo sapiens OX=9606 GN=SRSF11 PE=1 SV=1 | 8 | 24 | 3 | 8.74E+06 | 7.06E+06 | 6.56E+06 | 5.45E+06 | 4.80E+06 | 5.22E+06 | 3.81E+06 | 4.28E+06 | 4.63E+06 | 2.55E+06 | 3.90E+06 | 4.07E+06 | 6.31E+06 | 3.87E+06 | 1.63 | 0.70 | 0.00765212 |
| P08123 | Collagen alpha-2(I) chain OS=Homo sapiens OX=9606 GN=COL1A2 PE=1 SV=7 | 3 | 12 | 3 | 1.69E+06 | 1.65E+06 | 2.03E+06 | 1.81E+06 | 1.68E+06 | 1.77E+06 | 3.25E+06 | 2.74E+06 | 3.76E+06 | 2.08E+06 | 2.86E+06 | 8.95E+06 | 1.77E+06 | 3.94E+06 | 0.45 | -1.15 | 0.088473523 |
| Q9Y5X3 | Sorting nexin-5 OS=Homo sapiens OX=9606 GN=SNX5 PE=1 SV=1 | 10 | 40 | 3 | 6.43E+06 | 6.32E+06 | 3.75E+06 | 3.91E+06 | 4.07E+06 | 4.64E+06 | 6.16E+06 | 6.38E+06 | 4.31E+06 | 5.23E+06 | 4.82E+06 | 4.29E+06 | 4.85E+06 | 5.20E+06 | 0.93 | -0.10 | 0.591266506 |
| P35625 | Metalloproteinase inhibitor 3 OS=Homo sapiens OX=9606 GN=TIMP3 PE=1 SV=2 | 16 | 40 | 3 | 5.91E+06 | 6.52E+06 | 5.85E+06 | 6.06E+06 | 6.59E+06 | 5.77E+06 | 7.90E+06 | 8.52E+06 | 1.32E+07 | 7.76E+06 | 7.55E+06 | 1.19E+07 | 6.12E+06 | 9.46E+06 | 0.65 | -0.63 | 0.019232046 |
| P37837 | Transaldolase OS=Homo sapiens OX=9606 GN=TALDO1 PE=1 SV=2 | 12 | 42 | 3 | 9.87E+06 | 1.04E+07 | 9.93E+06 | 9.17E+06 | 9.26E+06 | 8.91E+06 | 7.17E+06 | 6.61E+06 | 6.74E+06 | 4.16E+06 | 6.50E+06 | 5.65E+06 | 9.60E+06 | 6.14E+06 | 1.56 | 0.64 | 0.000160045 |
| P84098 | 60S ribosomal protein L19 OS=Homo sapiens OX=9606 GN=RPL19 PE=1 SV=1 | 17 | 32 | 3 | 1.57E+07 | 1.58E+07 | 1.50E+07 | 1.76E+07 | 1.49E+07 | 1.40E+07 | 9.77E+06 | 9.29E+06 | 1.15E+07 | 1.11E+07 | 1.00E+07 | 9.07E+06 | 1.55E+07 | 1.01E+07 | 1.53 | 0.61 | 9.46844E-06 |
| Q96KP1 | Exocyst complex component 2 OS=Homo sapiens OX=9606 GN=EXOC2 PE=1 SV=1 | 6 | 9 | 3 | 2.91E+06 | 1.13E+06 | 1.94E+06 | 5.53E+05 | 1.90E+06 | 2.82E+06 | 1.68E+06 | 7.22E+05 | 1.37E+06 | 2.71E+05 | 7.30E+05 | 2.27E+06 | 1.88E+06 | 1.17E+06 | 1.60 | 0.68 | 0.177895094 |
| P02647 | Apolipoprotein A-I OS=Homo sapiens OX=9606 GN=APOA1 PE=1 SV=1 | 15 | 6 | 3 | 3.16E+06 | 3.59E+06 | 2.33E+06 | 3.00E+06 | 4.17E+06 | 4.06E+06 | 8.49E+06 | 8.66E+06 | 6.93E+06 | 3.96E+06 | 7.00E+06 | 7.10E+06 | 3.39E+06 | 7.02E+06 | 0.48 | -1.05 | 0.002048351 |
| P09382 | Galectin-1 OS=Homo sapiens OX=9606 GN=LGALS1 PE=1 SV=2 | 26 | 90 | 3 | 3.17E+07 | 3.24E+07 | 3.37E+07 | 3.36E+07 | 3.64E+07 | 3.07E+07 | 2.79E+07 | 2.72E+07 | 2.82E+07 | 2.89E+07 | 2.58E+07 | 2.30E+07 | 3.31E+07 | 2.68E+07 | 1.23 | 0.30 | 0.000399451 |
| Q8TAT6 | Nuclear protein localization protein 4 homolog OS=Homo sapiens OX=9606 GN=NPLOC4 PE=1 SV=3 | 7 | 24 | 3 | 3.37E+06 | 2.39E+06 | 2.33E+06 | 3.68E+06 | 2.01E+06 | 3.09E+06 | 1.68E+06 | 1.79E+06 | 2.81E+06 | 3.12E+06 | 1.48E+06 | 1.26E+06 | 2.81E+06 | 2.03E+06 | 1.39 | 0.47 | 0.084756209 |
| Q92466 | DNA damage-binding protein 2 OS=Homo sapiens OX=9606 GN=DDB2 PE=1 SV=1 | 10 | 14 | 3 | 5.29E+06 | 5.72E+06 | 5.36E+06 | 4.12E+06 | 4.69E+06 | 3.89E+06 | 4.01E+06 | 3.87E+06 | 4.11E+06 | 4.43E+06 | 4.19E+06 | 3.43E+06 | 4.85E+06 | 4.01E+06 | 1.21 | 0.27 | 0.038324733 |
| Q14624 | Inter-alpha-trypsin inhibitor heavy chain H4 OS=Homo sapiens OX=9606 GN=ITIH4 PE=1 SV=4 | 3 | 167 | 3 | 1.55E+08 | 1.34E+08 | 1.11E+08 | 1.73E+08 | 1.38E+08 | 1.09E+08 | 1.59E+08 | 1.43E+08 | 2.16E+08 | 2.36E+08 | 1.44E+08 | 1.52E+08 | 1.37E+08 | 1.75E+08 | 0.78 | -0.35 | 0.082923911 |
| Q13557 | Calcium/calmodulin-dependent protein kinase type II subunit delta OS=Homo sapiens OX=9606 GN=CAMK2D PE=1 SV=3 | 10 | 6 | 3 | 1.84E+06 | 1.06E+06 | 2.07E+06 | 2.32E+06 | 1.56E+06 | 2.02E+06 | 1.72E+06 | 1.07E+06 | 1.12E+06 | 1.58E+06 | 1.43E+06 | 1.37E+06 | 1.81E+06 | 1.38E+06 | 1.31 | 0.39 | 0.074753917 |
| O95084 | Serine protease 23 OS=Homo sapiens OX=9606 GN=PRSS23 PE=1 SV=1 | 9 | 38 | 3 | 5.55E+06 | 5.15E+06 | 4.81E+06 | 5.70E+06 | 4.80E+06 | 5.01E+06 | 1.08E+07 | 9.62E+06 | 9.50E+06 | 1.05E+07 | 9.60E+06 | 9.04E+06 | 5.17E+06 | 9.85E+06 | 0.52 | -0.93 | 5.63232E-07 |
| P27658 | Collagen alpha-1(VIII) chain OS=Homo sapiens OX=9606 GN=COL8A1 PE=1 SV=2 | 7 | 64 | 3 | 1.24E+07 | 1.27E+07 | 7.58E+06 | 8.73E+06 | 8.28E+06 | 1.15E+07 | 1.46E+07 | 1.20E+07 | 8.10E+06 | 8.23E+06 | 1.22E+07 | 1.27E+07 | 1.02E+07 | 1.13E+07 | 0.90 | -0.15 | 0.447102809 |
| P49773 | Histidine triad nucleotide-binding protein 1 OS=Homo sapiens OX=9606 GN=HINT1 PE=1 SV=2 | 24 | 33 | 3 | 1.20E+07 | 1.53E+07 | 1.42E+07 | 1.56E+07 | 1.32E+07 | 1.39E+07 | 1.08E+07 | 1.32E+07 | 9.12E+06 | 1.24E+07 | 8.93E+06 | 1.04E+07 | 1.40E+07 | 1.08E+07 | 1.30 | 0.38 | 0.005186774 |
| O75400 | Pre-mRNA-processing factor 40 homolog A OS=Homo sapiens OX=9606 GN=PRPF40A PE=1 SV=2 | 4 | 44 | 3 | 5.02E+06 | 5.99E+06 | 4.91E+06 | 4.01E+06 | 6.58E+06 | 4.94E+06 | 5.25E+06 | 5.88E+06 | 4.48E+06 | 3.63E+06 | 4.34E+06 | 3.47E+06 | 5.24E+06 | 4.51E+06 | 1.16 | 0.22 | 0.195267676 |
| O00391 | Sulfhydryl oxidase 1 OS=Homo sapiens OX=9606 GN=QSOX1 PE=1 SV=3 | 5 | 46 | 3 | 8.67E+06 | 9.00E+06 | 6.76E+06 | 8.63E+06 | 8.76E+06 | 7.09E+06 | 1.30E+07 | 1.26E+07 | 1.09E+07 | 1.07E+07 | 1.08E+07 | 1.10E+07 | 8.15E+06 | 1.15E+07 | 0.71 | -0.50 | 0.000170821 |
| Q9NR19 | Acetyl-coenzyme A synthetase, cytoplasmic OS=Homo sapiens OX=9606 GN=ACSS2 PE=1 SV=1 | 6 | 19 | 3 | 2.32E+06 | 3.03E+06 | 3.02E+06 | 3.51E+06 | 2.68E+06 | 2.44E+06 | 3.77E+06 | 3.69E+06 | 3.14E+06 | 3.89E+06 | 3.54E+06 | 2.79E+06 | 2.83E+06 | 3.47E+06 | 0.82 | -0.29 | 0.028903981 |
| Q14008 | Cytoskeleton-associated protein 5 OS=Homo sapiens OX=9606 GN=CKAP5 PE=1 SV=3 | 2 | 10 | 3 | 6.38E+06 | 5.81E+06 | 5.81E+06 | 6.09E+06 | 5.81E+06 | 5.33E+06 | 4.33E+06 | 1.47E+06 | 3.03E+06 | 1.66E+06 | 3.24E+06 | 2.55E+06 | 5.87E+06 | 2.71E+06 | 2.16 | 1.11 | 0.000436591 |
| Q96TA1 | Protein Niban 2 OS=Homo sapiens OX=9606 GN=NIBAN2 PE=1 SV=3 | 6 | 11 | 3 | 2.19E+06 | 1.15E+06 | 9.30E+05 | 3.84E+05 | 2.18E+06 | 2.08E+06 | 3.23E+06 | 1.98E+06 | 1.56E+06 | 1.66E+06 | 1.55E+06 | 1.67E+06 | 1.48E+06 | 1.94E+06 | 0.76 | -0.39 | 0.292581822 |
| Q92882 | Osteoclast-stimulating factor 1 OS=Homo sapiens OX=9606 GN=OSTF1 PE=1 SV=2 | 24 | 16 | 3 | 2.87E+06 | 3.20E+06 | 2.53E+06 | 3.28E+06 | 1.83E+06 | 2.02E+06 | 3.66E+06 | 3.69E+06 | 1.79E+06 | 1.86E+06 | 1.72E+06 | 2.75E+06 | 2.62E+06 | 2.58E+06 | 1.02 | 0.02 | 0.926527936 |
| Q92905 | COP9 signalosome complex subunit 5 OS=Homo sapiens OX=9606 GN=COPS5 PE=1 SV=4 | 14 | 12 | 3 | 3.85E+06 | 3.72E+06 | 3.08E+06 | 4.02E+06 | 4.06E+06 | 3.34E+06 | 3.76E+06 | 2.35E+06 | 7.62E+05 | 2.26E+06 | 3.33E+06 | 9.73E+05 | 3.68E+06 | 2.24E+06 | 1.64 | 0.72 | 0.031820873 |
| O15145 | Actin-related protein 2/3 complex subunit 3 OS=Homo sapiens OX=9606 GN=ARPC3 PE=1 SV=3 | 13 | 52 | 3 | 1.27E+07 | 1.21E+07 | 1.23E+07 | 1.18E+07 | 1.03E+07 | 1.03E+07 | 1.17E+07 | 1.10E+07 | 1.12E+07 | 1.05E+07 | 1.03E+07 | 9.65E+06 | 1.16E+07 | 1.07E+07 | 1.09 | 0.12 | 0.107525469 |
| Q02413 | Desmoglein-1 OS=Homo sapiens OX=9606 GN=DSG1 PE=1 SV=2 | 5 | 10 | 3 | 2.01E+06 | 2.05E+06 | 2.32E+05 | 1.34E+06 | 9.91E+05 | 1.73E+06 | 3.47E+05 | 1.83E+06 | 3.49E+05 | 1.58E+06 | 1.35E+06 | 3.62E+05 | 1.39E+06 | 9.69E+05 | 1.44 | 0.52 | 0.3164794 |
| P53992 | Protein transport protein Sec24C OS=Homo sapiens OX=9606 GN=SEC24C PE=1 SV=3 | 4 | 10 | 3 | 1.39E+06 | 4.83E+05 | 1.28E+06 | 1.28E+05 | 1.44E+06 | 1.08E+06 | 1.21E+06 | | 3.26E+05 | 6.46E+05 | 6.90E+05 | 6.11E+05 | 9.66E+05 | 6.96E+05 | 1.39 | 0.47 | 0.330388582 |
| P01009 | Alpha-1-antitrypsin OS=Homo sapiens OX=9606 GN=SERPINA1 PE=1 SV=3 | 8 | 6 | 3 | | | | | | | 6.57E+05 | | | | | 1.06E+07 | | 5.60E+06 | | | |
| P35052 | Glypican-1 OS=Homo sapiens OX=9606 GN=GPC1 PE=1 SV=2 | 8 | 29 | 3 | 6.39E+06 | 6.25E+06 | 5.66E+06 | 6.03E+06 | 4.05E+06 | 5.66E+06 | 6.10E+06 | 6.12E+06 | 4.05E+06 | 5.19E+06 | 3.73E+06 | 2.66E+06 | 5.67E+06 | 4.64E+06 | 1.22 | 0.29 | 0.15898109 |
| Q86TI2 | Dipeptidyl peptidase 9 OS=Homo sapiens OX=9606 GN=DPP9 PE=1 SV=3 | 4 | 27 | 3 | 5.30E+06 | 5.15E+06 | 4.77E+06 | 4.33E+06 | 4.25E+06 | 4.50E+06 | 4.52E+06 | 3.94E+06 | 3.64E+06 | 4.55E+06 | 4.49E+06 | 3.70E+06 | 4.72E+06 | 4.14E+06 | 1.14 | 0.19 | 0.0435091 |
| Q76LX8 | A disintegrin and metalloproteinase with thrombospondin motifs 13 OS=Homo sapiens OX=9606 GN=ADAMTS13 PE=1 SV=1 | 2 | 84 | 3 | 9.85E+06 | 1.37E+07 | 1.44E+07 | 1.52E+07 | 1.52E+07 | 1.46E+07 | 2.32E+07 | 2.10E+07 | 2.41E+07 | 1.78E+07 | 1.86E+07 | 1.88E+07 | 1.38E+07 | 2.06E+07 | 0.67 | -0.57 | 0.000639763 |

### Supplementary Table 4. List of all proteins identified by label free proteomics analysis.

| Accession | Description | Coverage [%] | # PSMs | # Unique Peptides | Amitriptyline 1 | Amitriptyline 2 | Amitriptyline 3 | Amitriptyline 4 | Amitriptyline 5 | Amitriptyline 6 | Vehicle 1 | Vehicle 2 | Vehicle 3 | Vehicle 4 | Vehicle 5 | Vehicle 6 | Average amitriptyline | Average vehicle | Ratio amitriptyline/ vehicle | Log 2 fold change (amitriptyline /vehicle) | P-value |
|---|---|---|---|---|---|---|---|---|---|---|---|---|---|---|---|---|---|---|---|---|---|
| Q14117 | Dihydropyrimidinase OS=Homo sapiens OX=9606 GN=DPYS PE=1 SV=1 | 6 | 40 | 3 | 3.43E+06 | 4.63E+06 | 5.54E+06 | 6.17E+06 | 5.23E+06 | 4.08E+06 | 6.61E+06 | 6.97E+06 | 8.02E+06 | 8.69E+06 | 7.28E+06 | 5.87E+06 | 4.85E+06 | 7.24E+06 | 0.67 | -0.58 | 0.002029804 |
| Q9NY12 | H/ACA ribonucleoprotein complex subunit 1 OS=Homo sapiens OX=9606 GN=GAR1 PE=1 SV=1 | 16 | 30 | 3 | 4.55E+06 | 2.57E+06 | 4.22E+06 | 3.93E+06 | 3.79E+06 | 4.76E+06 | 2.37E+06 | 3.31E+06 | 2.52E+06 | 3.14E+06 | 2.82E+06 | 2.05E+06 | 3.97E+06 | 2.70E+06 | 1.47 | 0.56 | 0.008748343 |
| P40926 | Malate dehydrogenase, mitochondrial OS=Homo sapiens OX=9606 GN=MDH2 PE=1 SV=3 | 11 | 25 | 3 | 2.25E+06 | 9.30E+05 | 1.23E+06 | 1.56E+06 | 1.31E+06 | 1.33E+06 |  | 3.56E+06 | 7.83E+05 | 1.54E+06 | 1.15E+06 | 1.55E+06 | 1.44E+06 | 1.72E+06 | 0.84 | -0.26 | 0.609708468 |
| P78344 | Eukaryotic translation initiation factor 4 gamma 2 OS=Homo sapiens OX=9606 GN=EIF4G2 PE=1 SV=1 | 3 | 8 | 3 | 1.11E+06 | 8.26E+05 | 1.50E+06 | 7.72E+05 | 9.79E+05 | 1.68E+06 | 6.40E+05 | 1.89E+06 | 2.91E+05 |  | 1.18E+06 | 6.40E+05 | 1.14E+06 | 9.28E+05 | 1.23 | 0.30 | 0.51971241 |
| P20042 | Eukaryotic translation initiation factor 2 subunit 2 OS=Homo sapiens OX=9606 GN=EIF2S2 PE=1 SV=2 | 9 | 22 | 3 | 5.03E+06 | 4.08E+06 | 5.94E+06 | 7.06E+06 | 6.62E+06 | 5.32E+06 | 3.77E+06 | 3.81E+06 | 4.53E+06 | 3.86E+06 | 3.74E+06 | 3.78E+06 | 5.68E+06 | 3.91E+06 | 1.45 | 0.54 | 0.009558287 |
| O75832 | 26S proteasome non-ATPase regulatory subunit 10 OS=Homo sapiens OX=9606 GN=PSMD10 PE=1 SV=1 | 19 | 16 | 3 | 4.33E+06 | 5.20E+06 | 4.15E+06 | 4.56E+06 | 6.28E+06 | 5.32E+06 | 4.12E+06 | 4.53E+06 | 4.87E+06 | 4.10E+06 | 3.58E+06 | 2.36E+06 | 4.97E+06 | 3.93E+06 | 1.27 | 0.34 | 0.056710384 |
| P26196 | Probable ATP-dependent RNA helicase DDX6 OS=Homo sapiens OX=9606 GN=DDX6 PE=1 SV=2 | 8 | 15 | 3 | 3.29E+06 | 3.14E+06 | 2.29E+06 | 3.33E+06 | 1.83E+06 | 2.45E+06 | 2.21E+06 | 1.09E+06 | 6.47E+05 | 1.18E+06 | 1.60E+06 | 1.90E+06 | 2.72E+06 | 1.44E+06 | 1.89 | 0.92 | 0.003985184 |
| Q9HC38 | Glyoxalase domain-containing protein 4 OS=Homo sapiens OX=9606 GN=GLOD4 PE=1 SV=1 | 10 | 47 | 3 | 9.93E+06 | 1.02E+07 | 1.11E+07 | 9.23E+06 | 9.46E+06 | 1.08E+07 | 1.32E+07 | 1.28E+07 | 1.33E+07 | 1.19E+07 | 1.37E+07 | 1.05E+07 | 1.01E+07 | 1.26E+07 | 0.80 | -0.32 | 0.002170711 |
| P30044 | Peroxiredoxin-5, mitochondrial OS=Homo sapiens OX=9606 GN=PRDX5 PE=1 SV=4 | 30 | 6 | 3 | 1.33E+06 | 6.28E+05 | 3.22E+06 | 2.47E+06 | 9.15E+05 | 3.91E+06 |  |  |  |  | 1.03E+06 | 2.48E+05 | 2.08E+06 | 6.37E+05 | 3.26 | 1.71 | 0.08373926 |
| Q13045 | Protein flightless-1 homolog OS=Homo sapiens OX=9606 GN=FLII PE=1 SV=2 | 3 | 26 | 3 | 5.45E+06 | 3.77E+06 | 4.98E+06 | 5.44E+06 | 4.84E+06 | 4.89E+06 | 4.21E+06 | 3.45E+06 | 2.83E+06 | 4.22E+06 | 3.43E+06 | 2.74E+06 | 4.89E+06 | 3.48E+06 | 1.41 | 0.49 | 0.003001446 |
| P48444 | Coatomer subunit delta OS=Homo sapiens OX=9606 GN=ARCN1 PE=1 SV=1 | 7 | 10 | 3 | 3.91E+06 | 3.70E+06 | 3.39E+06 | 3.58E+06 | 3.03E+06 | 2.16E+06 | 2.04E+06 | 1.78E+06 | 2.15E+06 | 2.59E+06 | 1.30E+06 | 8.04E+05 | 3.29E+06 | 1.78E+06 | 1.85 | 0.89 | 0.002042845 |
| P83731 | 60S ribosomal protein L24 OS=Homo sapiens OX=9606 GN=RPL24 PE=1 SV=1 | 16 | 52 | 3 | 2.29E+07 | 2.14E+07 | 2.16E+07 | 2.21E+07 | 2.48E+07 | 2.06E+07 | 1.48E+07 | 1.55E+07 | 1.28E+07 | 1.39E+07 | 1.25E+07 | 1.08E+07 | 2.22E+07 | 1.34E+07 | 1.66 | 0.73 | 2.56297E-06 |
| O00339 | Matrilin-2 OS=Homo sapiens OX=9606 GN=MATN2 PE=1 SV=4 | 5 | 13 | 3 |  | 3.03E+06 |  |  | 1.05E+06 |  |  | 4.13E+06 | 1.56E+06 |  | 3.02E+06 | 1.19E+06 | 2.04E+06 | 2.48E+06 | 0.82 | -0.28 | 0.751689528 |
| Q9NQW7 | Xaa-Pro aminopeptidase 1 OS=Homo sapiens OX=9606 GN=XPNPEP1 PE=1 SV=3 | 5 | 32 | 3 | 5.13E+06 | 5.81E+06 | 3.12E+06 | 3.92E+06 | 3.59E+06 | 3.14E+06 | 7.20E+06 | 8.32E+06 | 7.01E+06 | 6.54E+06 | 6.08E+06 | 5.48E+06 | 4.12E+06 | 6.77E+06 | 0.61 | -0.72 | 0.001427047 |
| P62851 | 40S ribosomal protein S25 OS=Homo sapiens OX=9606 GN=RPS25 PE=1 SV=1 | 22 | 93 | 3 | 5.64E+07 | 5.37E+07 | 6.71E+07 | 5.58E+07 | 7.96E+07 | 7.19E+07 | 4.00E+07 | 3.70E+07 | 4.91E+07 | 5.63E+07 | 4.21E+07 | 4.10E+07 | 6.41E+07 | 4.43E+07 | 1.45 | 0.53 | 0.004167885 |
| Q14847 | LIM and SH3 domain protein 1 OS=Homo sapiens OX=9606 GN=LASP1 PE=1 SV=2 | 13 | 42 | 3 | 5.68E+06 | 6.65E+06 | 6.00E+06 | 5.56E+06 | 5.79E+06 | 5.46E+06 | 3.43E+06 | 3.40E+06 | 4.39E+06 | 5.69E+06 | 3.31E+06 | 2.42E+06 | 5.86E+06 | 3.77E+06 | 1.55 | 0.63 | 0.004726468 |
| Q96RP9 | Elongation factor G, mitochondrial OS=Homo sapiens OX=9606 GN=GFM1 PE=1 SV=2 | 4 | 10 | 3 | 3.10E+06 | 2.99E+06 | 2.11E+06 | 2.58E+06 | 2.25E+06 | 2.17E+06 | 1.77E+06 | 8.56E+05 | 1.32E+06 | 1.15E+06 | 1.46E+06 | 7.14E+05 | 2.53E+06 | 1.21E+06 | 2.09 | 1.06 | 0.000243346 |
| Q9Y371 | Endophilin-B1 OS=Homo sapiens OX=9606 GN=SH3GLB1 PE=1 SV=1 | 8 | 18 | 3 | 1.53E+06 | 4.09E+06 | 3.57E+06 | 3.87E+06 | 3.40E+06 | 3.63E+06 | 3.24E+06 | 3.89E+06 | 3.64E+06 | 3.52E+06 | 3.00E+06 | 2.84E+06 | 3.35E+06 | 3.35E+06 | 1.00 | 0.00 | 0.985668439 |
| P49902 | Cytosolic purine 5'-nucleotidase OS=Homo sapiens OX=9606 GN=NT5C2 PE=1 SV=1 | 8 | 5 | 3 | 1.33E+06 | 5.91E+05 | 8.37E+05 | 6.10E+05 | 9.24E+05 | 6.89E+05 |  |  | 1.08E+06 |  | 8.37E+05 | 6.84E+05 | 8.30E+05 | 8.68E+05 | 0.96 | -0.07 | 0.82136733 |
| P16885 | 1-phosphatidylinositol 4,5-bisphosphate phosphodiesterase gamma-2 OS=Homo sapiens OX=9606 GN=PLCG2 PE=1 SV=4 | 3 | 14 | 3 | 2.60E+06 | 2.35E+06 | 2.25E+06 | 1.75E+06 | 1.82E+06 | 2.63E+06 | 3.10E+06 | 3.39E+06 | 4.39E+06 | 3.84E+06 | 2.75E+06 | 2.64E+06 | 2.23E+06 | 3.36E+06 | 0.67 | -0.59 | 0.00746715 |
| P05198 | Eukaryotic translation initiation factor 2 subunit 1 OS=Homo sapiens OX=9606 GN=EIF2S1 PE=1 SV=3 | 9 | 62 | 3 | 1.57E+07 | 1.64E+07 | 1.48E+07 | 1.58E+07 | 1.60E+07 | 1.46E+07 | 1.44E+07 | 1.58E+07 | 1.49E+07 | 1.55E+07 | 1.41E+07 | 1.16E+07 | 1.56E+07 | 1.44E+07 | 1.08 | 0.11 | 0.134856856 |
| P27701 | CD82 antigen OS=Homo sapiens OX=9606 GN=CD82 PE=1 SV=1 | 15 | 18 | 3 | 3.48E+06 | 2.75E+06 | 5.80E+06 | 9.13E+06 | 4.64E+06 | 3.87E+06 | 2.62E+06 | 6.00E+06 | 8.98E+06 | 6.31E+06 | 4.27E+06 | 4.88E+06 | 4.95E+06 | 5.51E+06 | 0.90 | -0.16 | 0.670783977 |
| P49959 | Double-strand break repair protein MRE11 OS=Homo sapiens OX=9606 GN=MRE11 PE=1 SV=3 | 6 | 6 | 3 | 1.83E+06 | 1.15E+06 | 4.95E+05 | 8.97E+05 | 1.09E+06 | 1.10E+06 | 1.04E+06 | 9.11E+05 |  |  | 2.38E+05 | 6.12E+05 | 1.09E+06 | 7.00E+05 | 1.56 | 0.64 | 0.159461392 |
| P14324 | Farnesyl pyrophosphate synthase OS=Homo sapiens OX=9606 GN=FDPS PE=1 SV=4 | 11 | 10 | 3 | 5.64E+05 | 8.04E+05 | 1.61E+06 | 1.70E+06 | 1.44E+06 | 2.21E+06 | 1.48E+06 |  | 1.65E+06 | 7.02E+05 | 1.30E+06 | 1.06E+06 | 1.39E+06 | 1.24E+06 | 1.12 | 0.17 | 0.627853302 |
| Q6NZI2 | Caveolae-associated protein 1 OS=Homo sapiens OX=9606 GN=CAVIN1 PE=1 SV=1 | 8 | 26 | 3 | 5.89E+06 | 6.94E+06 | 8.46E+06 | 8.54E+06 | 8.49E+06 | 6.84E+06 | 3.24E+06 | 2.50E+06 | 2.18E+06 | 2.57E+06 | 2.15E+06 | 1.95E+06 | 7.53E+06 | 2.43E+06 | 3.09 | 1.63 | 2.60723E-05 |
| O60264 | SWI/SNF-related matrix-associated actin-dependent regulator of chromatin subfamily A member 5 OS=Homo sapiens OX=9606 GN=SMARCA5 PE=1 SV=1 | 3 | 17 | 3 | 4.11E+06 | 4.77E+06 | 3.77E+06 | 4.28E+06 | 3.75E+06 | 3.72E+06 | 1.78E+06 | 2.12E+06 | 1.49E+06 | 2.06E+06 | 2.28E+06 | 2.20E+06 | 4.07E+06 | 1.99E+06 | 2.05 | 1.03 | 3.27802E-06 |
| O75116 | Rho-associated protein kinase 2 OS=Homo sapiens OX=9606 GN=ROCK2 PE=1 SV=4 | 3 | 7 | 3 | 7.20E+05 | 9.83E+05 | 1.19E+06 | 9.49E+05 | 4.24E+05 | 1.92E+06 | 2.33E+06 | 1.43E+06 | 1.82E+06 | 1.74E+06 | 1.54E+06 | 1.57E+06 | 1.03E+06 | 1.74E+06 | 0.59 | -0.75 | 0.018961965 |
| Q15057 | Arf-GAP with coiled-coil, ANK repeat and PH domain-containing protein 2 OS=Homo sapiens OX=9606 GN=ACAP2 PE=1 SV=3 | 6 | 23 | 3 | 3.98E+06 | 7.92E+06 | 1.13E+07 | 1.85E+07 | 1.19E+07 | 9.56E+06 | 9.70E+06 | 8.77E+06 | 7.41E+06 | 3.37E+06 | 3.39E+06 | 5.91E+06 | 1.05E+07 | 6.42E+06 | 1.64 | 0.71 | 0.108334526 |
| P23381 | Tryptophan--tRNA ligase, cytoplasmic OS=Homo sapiens OX=9606 GN=WARS PE=1 SV=2 | 9 | 9 | 3 | 1.93E+06 | 1.76E+06 | 1.93E+06 | 1.43E+06 | 5.95E+05 | 1.37E+06 | 5.69E+05 | 3.57E+06 |  | 1.06E+06 | 6.82E+05 | 4.95E+05 | 1.50E+06 | 1.28E+06 | 1.18 | 0.24 | 0.729472475 |
| O95445 | Apolipoprotein M OS=Homo sapiens OX=9606 GN=APOM PE=1 SV=2 | 18 | 58 | 3 | 4.67E+07 | 5.28E+07 | 4.75E+07 | 4.04E+07 | 4.82E+07 | 3.66E+07 | 6.34E+07 | 6.73E+07 | 5.99E+07 | 4.70E+07 | 5.17E+07 | 4.94E+07 | 4.54E+07 | 5.64E+07 | 0.80 | -0.32 | 0.024820283 |
| P10643 | Complement component C7 OS=Homo sapiens OX=9606 GN=C7 PE=1 SV=2 | 2 | 66 | 3 | 2.86E+07 | 2.55E+07 | 1.75E+07 | 2.02E+07 | 1.89E+07 | 2.26E+07 | 3.47E+07 | 4.28E+07 | 2.34E+07 | 2.49E+07 | 2.54E+07 | 2.71E+07 | 2.22E+07 | 2.97E+07 | 0.75 | -0.42 | 0.066148483 |
| O14744 | Protein arginine N-methyltransferase 5 OS=Homo sapiens OX=9606 GN=PRMT5 PE=1 SV=4 | 5 | 20 | 3 | 9.49E+06 | 1.01E+07 | 8.38E+06 | 8.59E+06 | 8.66E+06 | 2.17E+06 | 9.44E+06 | 2.00E+06 | 1.51E+06 | 9.03E+06 | 7.46E+06 | 1.82E+06 | 7.89E+06 | 5.21E+06 | 1.52 | 0.60 | 0.201352765 |
| P06132 | Uroporphyrinogen decarboxylase OS=Homo sapiens OX=9606 GN=UROD PE=1 SV=2 | 9 | 34 | 3 | 7.46E+06 | 6.25E+06 | 5.11E+06 | 3.64E+06 | 6.43E+06 | 5.66E+06 | 7.21E+06 | 7.85E+06 | 3.98E+06 | 7.70E+06 | 6.87E+06 | 7.15E+06 | 5.76E+06 | 6.79E+06 | 0.85 | -0.24 | 0.218458482 |
| P23229 | Integrin alpha-6 OS=Homo sapiens OX=9606 GN=ITGA6 PE=1 SV=5 | 3 | 12 | 3 |  |  | 6.07E+05 | 6.50E+05 | 7.27E+05 |  | 4.19E+05 |  | 6.23E+05 | 7.26E+05 | 5.38E+05 | 8.45E+05 | 6.61E+05 | 6.30E+05 | 1.05 | 0.07 | 0.7171384 |
| P08514 | Integrin alpha-IIb OS=Homo sapiens OX=9606 GN=ITGA2B PE=1 SV=3 | 4 | 6 | 3 | 3.92E+05 |  | 5.04E+05 | 2.18E+05 | 1.42E+06 | 1.11E+06 | 1.21E+06 | 7.58E+05 |  | 9.91E+05 | 5.52E+05 | 4.89E+06 | 7.29E+05 | 1.68E+06 | 0.43 | -1.20 | 0.313874082 |
| Q969P0 | Immunoglobulin superfamily member 8 OS=Homo sapiens OX=9606 GN=IGSF8 PE=1 SV=1 | 5 | 23 | 3 | 3.35E+06 | 2.72E+06 | 3.64E+06 | 4.16E+06 | 4.29E+06 | 3.81E+06 | 2.21E+06 | 2.14E+06 | 2.03E+06 | 2.13E+06 | 2.16E+06 | 1.83E+06 | 3.66E+06 | 2.08E+06 | 1.76 | 0.82 | 0.000820177 |
| Q00059 | Transcription factor A, mitochondrial OS=Homo sapiens OX=9606 GN=TFAM PE=1 SV=1 | 12 | 30 | 3 | 7.65E+06 | 8.65E+06 | 8.05E+06 | 9.73E+06 | 8.79E+06 | 7.88E+06 | 3.69E+06 | 7.24E+06 | 7.11E+06 | 6.92E+06 | 3.76E+06 | 6.09E+06 | 8.46E+06 | 5.80E+06 | 1.46 | 0.54 | 0.00907494 |
| P18077 | 60S ribosomal protein L35a OS=Homo sapiens OX=9606 GN=RPL35A PE=1 SV=2 | 22 | 73 | 3 | 1.62E+07 | 1.57E+07 | 1.60E+07 | 1.64E+07 | 1.83E+07 | 1.59E+07 | 1.46E+07 | 1.51E+07 | 1.59E+07 | 1.43E+07 | 1.35E+07 | 1.14E+07 | 1.64E+07 | 1.41E+07 | 1.16 | 0.21 | 0.015389441 |
| Q99798 | Aconitate hydratase, mitochondrial OS=Homo sapiens OX=9606 GN=ACO2 PE=1 SV=2 | 4 | 14 | 3 | 1.55E+06 | 1.89E+06 | 2.20E+06 | 1.45E+06 | 1.65E+06 | 2.29E+06 | 1.48E+06 | 1.54E+06 | 2.02E+06 | 1.62E+06 | 1.89E+06 | 1.19E+06 | 1.84E+06 | 1.62E+06 | 1.13 | 0.18 | 0.274144395 |
| P07738 | Bisphosphoglycerate mutase OS=Homo sapiens OX=9606 GN=BPGM PE=1 SV=2 | 12 | 30 | 3 | 4.99E+06 | 5.04E+06 | 5.21E+06 | 4.67E+06 | 5.19E+06 | 5.16E+06 | 6.92E+06 | 6.11E+06 | 6.09E+06 | 5.80E+06 | 5.91E+06 | 5.98E+06 | 5.04E+06 | 6.14E+06 | 0.82 | -0.28 | 0.000489287 |
| O60218 | Aldo-keto reductase family 1 member B10 OS=Homo sapiens OX=9606 GN=AKR1B10 PE=1 SV=2 | 5 | 47 | 3 | 5.05E+07 | 5.07E+07 | 4.66E+07 | 5.32E+07 | 5.42E+07 | 3.95E+07 | 6.54E+07 | 6.43E+07 | 7.26E+07 | 7.95E+07 | 5.64E+07 | 5.72E+07 | 4.91E+07 | 6.59E+07 | 0.74 | -0.42 | 0.004037189 |
| Q9UNS2 | COP9 signalosome complex subunit 3 OS=Homo sapiens OX=9606 GN=COPS3 PE=1 SV=3 | 10 | 9 | 3 |  | 3.81E+06 | 3.27E+06 | 9.85E+05 | 1.75E+06 | 3.46E+06 | 3.46E+06 | 2.78E+06 | 3.34E+06 | 3.87E+06 | 8.15E+05 | 4.12E+06 | 2.66E+06 | 3.07E+06 | 0.87 | -0.21 | 0.589485633 |
| P46063 | ATP-dependent DNA helicase Q1 OS=Homo sapiens OX=9606 GN=RECQL PE=1 SV=3 | 5 | 26 | 3 | 2.27E+07 | 2.61E+07 | 2.46E+07 | 2.72E+07 | 2.37E+07 | 2.32E+07 | 2.39E+07 | 2.68E+07 | 2.48E+07 | 2.77E+07 | 2.39E+07 | 2.26E+07 | 2.46E+07 | 2.49E+07 | 0.99 | -0.02 | 0.75712383 |
| P62899 | 60S ribosomal protein L31 OS=Homo sapiens OX=9606 GN=RPL31 PE=1 SV=1 | 33 | 8 | 3 | 1.63E+06 | 5.84E+06 | 4.79E+06 | 5.55E+06 | 5.95E+06 | 5.41E+06 | 4.36E+06 | 4.54E+06 | 4.33E+06 | 5.21E+06 | 3.90E+06 | 3.01E+06 | 4.86E+06 | 4.22E+06 | 1.15 | 0.20 | 0.411989117 |
| P00374 | Dihydrofolate reductase OS=Homo sapiens OX=9606 GN=DHFR PE=1 SV=2 | 26 | 32 | 3 | 1.38E+07 | 1.53E+07 | 1.49E+07 | 1.19E+07 | 1.49E+07 | 1.45E+07 | 1.52E+07 | 1.54E+07 | 1.53E+07 | 1.68E+07 | 1.43E+07 | 1.11E+07 | 1.42E+07 | 1.47E+07 | 0.97 | -0.05 | 0.638692333 |
| P04003 | C4b-binding protein alpha chain OS=Homo sapiens OX=9606 GN=C4BPA PE=1 SV=2 | 5 | 42 | 3 | 8.99E+06 | 1.13E+07 | 4.05E+07 | 2.42E+07 | 1.91E+07 | 2.33E+07 | 1.37E+07 | 7.87E+06 | 1.03E+07 | 1.05E+07 | 1.08E+07 | 1.89E+07 | 2.12E+07 | 1.20E+07 | 1.77 | 0.82 | 0.106291964 |
| Q13144 | Translation initiation factor eIF-2B subunit epsilon OS=Homo sapiens OX=9606 GN=EIF2B5 PE=1 SV=3 | 5 | 5 | 3 | 3.86E+06 | 1.36E+06 |  | 2.81E+06 | 2.85E+06 | 4.81E+06 | 4.70E+06 |  | 6.72E+05 | 9.80E+05 | 3.30E+06 | 3.78E+06 | 3.14E+06 | 2.69E+06 | 1.17 | 0.22 | 0.659111765 |
| Q9UDY2 | Tight junction protein ZO-2 OS=Homo sapiens OX=9606 GN=TJP2 PE=1 SV=2 | 2 | 15 | 3 | 2.39E+06 | 4.13E+06 | 4.34E+06 | 4.84E+06 | 5.06E+06 | 3.55E+06 | 3.92E+06 | 1.84E+06 | 4.41E+06 | 2.37E+06 | 3.85E+06 | 2.84E+06 | 4.05E+06 | 3.21E+06 | 1.26 | 0.34 | 0.170537399 |
| P54727 | UV excision repair protein RAD23 homolog B OS=Homo sapiens OX=9606 GN=RAD23B PE=1 SV=1 | 7 | 18 | 3 | 9.79E+06 | 1.15E+07 | 1.33E+07 | 1.52E+07 | 1.39E+07 | 9.91E+06 | 1.32E+07 | 1.32E+07 | 1.19E+07 | 1.38E+07 | 1.28E+07 | 1.17E+07 | 1.23E+07 | 1.28E+07 | 0.96 | -0.06 | 0.606198214 |
| P29144 | Tripeptidyl-peptidase 2 OS=Homo sapiens OX=9606 GN=TPP2 PE=1 SV=4 | 2 | 6 | 3 | 3.27E+06 | 1.50E+06 | 3.85E+06 | 3.66E+06 | 3.30E+06 | 3.69E+06 | 2.75E+06 | 3.52E+05 | 3.78E+06 | 2.91E+06 | 4.28E+05 | 1.95E+06 | 3.21E+06 | 2.03E+06 | 1.58 | 0.66 | 0.11428255 |
| P15151 | Poliovirus receptor OS=Homo sapiens OX=9606 GN=PVR PE=1 SV=2 | 6 | 10 | 3 | 5.23E+06 | 5.87E+06 | 6.04E+06 | 6.52E+06 | 6.14E+06 | 5.69E+06 | 3.50E+06 | 4.12E+06 | 2.72E+06 | 2.79E+06 | 2.35E+06 | 2.70E+06 | 5.91E+06 | 3.03E+06 | 1.95 | 0.96 | 1.07461E-05 |
| Q13509 | Tubulin beta-3 chain OS=Homo sapiens OX=9606 GN=TUBB3 PE=1 SV=2 | 47 | 3130 | 4 | 3.59E+06 | 3.48E+06 | 2.09E+06 | 2.39E+06 | 1.48E+06 | 2.46E+06 | 3.29E+06 | 2.25E+06 | 1.45E+06 | 1.38E+06 | 1.06E+06 | 2.48E+06 | 2.58E+06 | 1.98E+06 | 1.30 | 0.38 | 0.2376746 |
| Q5QNW6 | Histone H2B type 2-F OS=Homo sapiens OX=9606 GN=HIST2H2BF PE=1 SV=3 | 56 | 6410 | 4 | 7.85E+09 | 6.59E+09 | 9.43E+09 | 7.36E+09 | 7.96E+09 | 7.69E+09 | 8.01E+09 | 7.00E+09 | 8.29E+09 | 7.38E+09 | 7.73E+09 | 7.48E+09 | 7.81E+09 | 7.65E+09 | 1.02 | 0.03 | 0.708002684 |
| P13929 | Beta-enolase OS=Homo sapiens OX=9606 GN=ENO3 PE=1 SV=5 | 31 | 397 | 4 | 1.55E+07 | 1.45E+07 | 2.07E+07 | 2.24E+07 | 2.13E+07 | 1.90E+07 | 2.69E+07 | 2.14E+07 | 2.25E+07 | 1.85E+07 | 2.29E+07 | 2.26E+07 | 1.89E+07 | 2.25E+07 | 0.84 | -0.25 | 0.067197207 |
| Q05639 | Elongation factor 1-alpha 2 OS=Homo sapiens OX=9606 GN=EEF1A2 PE=1 SV=1 | 42 | 693 | 4 | 6.80E+06 | 5.47E+06 | 8.77E+06 | 5.51E+06 | 7.23E+06 | 5.20E+06 | 8.79E+06 | 5.14E+06 | 3.74E+06 | 3.13E+06 | 6.12E+06 | 3.98E+06 | 6.50E+06 | 5.15E+06 | 1.26 | 0.34 | 0.219976107 |
| P04899 | Guanine nucleotide-binding protein G(i) subunit alpha-2 OS=Homo sapiens OX=9606 GN=GNAI2 PE=1 SV=3 | 30 | 148 | 4 | 5.93E+07 | 6.23E+07 | 7.74E+07 | 6.77E+07 | 8.36E+07 | 5.95E+07 | 6.07E+07 | 5.52E+07 | 6.86E+07 | 6.24E+07 | 7.33E+07 | 6.46E+07 | 6.83E+07 | 6.41E+07 | 1.07 | 0.09 | 0.415255847 |
| P31943 | Heterogeneous nuclear ribonucleoprotein H OS=Homo sapiens OX=9606 GN=HNRNPH1 PE=1 SV=4 | 20 | 150 | 4 | 2.52E+07 | 2.34E+07 | 1.93E+07 | 2.04E+07 | 1.99E+07 | 2.16E+07 | 1.28E+07 | 1.12E+07 | 1.03E+07 | 9.23E+06 | 8.21E+06 | 9.97E+06 | 2.17E+07 | 1.03E+07 | 2.11 | 1.08 | 3.79739E-06 |
| P02774 | Vitamin D-binding protein OS=Homo sapiens OX=9606 GN=GC PE=1 SV=2 | 9 | 94 | 4 | 3.51E+07 | 2.37E+07 | 2.12E+07 | 2.90E+07 | 2.67E+07 | 3.18E+07 | 4.14E+07 | 3.57E+07 | 3.07E+07 | 3.41E+07 | 3.08E+07 | 2.93E+07 | 2.79E+07 | 3.37E+07 | 0.83 | -0.27 | 0.0658469 |
| P35241 | Radixin OS=Homo sapiens OX=9606 GN=RDX PE=1 SV=1 | 19 | 221 | 4 | 4.70E+06 | 1.92E+06 |  |  | 1.21E+06 | 9.43E+05 | 1.39E+06 | 9.99E+05 |  | 7.45E+05 | 8.19E+05 | 1.32E+06 | 2.19E+06 | 1.06E+06 | 2.08 | 1.06 | 0.278437355 |
| P62314 | Small nuclear ribonucleoprotein Sm D1 OS=Homo sapiens OX=9606 GN=SNRPD1 PE=1 SV=1 | 55 | 183 | 4 | 5.98E+07 | 6.77E+07 | 7.79E+07 | 6.34E+07 | 7.18E+07 | 8.12E+07 | 4.09E+07 | 5.27E+07 | 5.14E+07 | 4.71E+07 | 5.61E+07 | 3.49E+07 | 7.03E+07 | 4.72E+07 | 1.49 | 0.57 | 0.000601081 |
| P60033 | CD81 antigen OS=Homo sapiens OX=9606 GN=CD81 PE=1 SV=1 | 28 | 74 | 4 | 3.14E+07 | 2.77E+07 | 1.89E+07 | 2.33E+07 | 2.74E+07 | 2.85E+07 | 2.30E+07 | 1.88E+07 | 1.30E+07 | 1.59E+07 | 1.70E+07 | 1.72E+07 | 2.62E+07 | 1.75E+07 | 1.50 | 0.58 | 0.003674527 |
| Q00266 | S-adenosylmethionine synthase isoform type-1 OS=Homo sapiens OX=9606 GN=MAT1A PE=1 SV=2 | 19 | 144 | 4 | 3.53E+07 | 3.40E+07 | 3.62E+07 | 3.55E+07 | 3.22E+07 | 2.47E+07 | 4.37E+07 | 4.43E+07 | 3.01E+07 | 4.05E+07 | 2.92E+07 | 3.67E+07 | 3.30E+07 | 3.74E+07 | 0.88 | -0.18 | 0.201708997 |
| P51148 | Ras-related protein Rab-5C OS=Homo sapiens OX=9606 GN=RAB5C PE=1 SV=2 | 37 | 124 | 4 | 3.82E+07 | 4.11E+07 | 3.34E+07 | 3.22E+07 | 4.14E+07 | 3.60E+07 | 3.01E+07 | 2.99E+07 | 2.58E+07 | 2.63E+07 | 2.66E+07 | 2.19E+07 | 3.70E+07 | 2.68E+07 | 1.38 | 0.47 | 0.000536151 |
| Q16576 | Histone-binding protein RBBP7 OS=Homo sapiens OX=9606 GN=RBBP7 PE=1 SV=1 | 28 | 68 | 4 | 1.66E+07 | 1.78E+07 | 1.29E+07 | 1.50E+07 | 1.30E+07 | 1.56E+07 | 1.12E+07 | 1.31E+07 | 7.86E+06 | 9.68E+06 | 9.00E+06 | 8.76E+06 | 1.52E+07 | 9.93E+06 | 1.53 | 0.61 | 0.000839949 |
| Q15436 | Protein transport protein Sec23A OS=Homo sapiens OX=9606 GN=SEC23A PE=1 SV=2 | 12 | 66 | 4 | 2.59E+07 | 2.99E+07 | 2.97E+07 | 3.08E+07 | 2.46E+07 | 2.60E+07 | 2.29E+07 | 1.65E+07 | 1.39E+07 | 1.38E+07 | 1.87E+07 | 1.66E+07 | 2.78E+07 | 1.70E+07 | 1.63 | 0.71 | 0.00014234 |
| Q15233 | Non-POU domain-containing octamer-binding protein OS=Homo sapiens OX=9606 GN=NONO PE=1 SV=4 | 12 | 107 | 4 | 1.83E+07 | 1.90E+07 | 2.26E+07 | 2.61E+07 | 2.08E+07 | 1.80E+07 | 1.17E+07 | 1.33E+07 | 1.25E+07 | 1.34E+07 | 1.31E+07 | 8.48E+06 | 2.08E+07 | 1.21E+07 | 1.72 | 0.78 | 0.000329338 |
| O15144 | Actin-related protein 2/3 complex subunit 2 OS=Homo sapiens OX=9606 GN=ARPC2 PE=1 SV=1 | 23 | 70 | 4 | 1.19E+07 | 1.13E+07 | 1.53E+07 | 1.21E+07 | 1.54E+07 | 1.20E+07 | 1.08E+07 | 1.22E+07 | 1.38E+07 | 6.46E+06 | 1.40E+07 | 1.03E+07 | 1.30E+07 | 1.13E+07 | 1.15 | 0.21 | 0.235976115 |
| P61769 | Beta-2-microglobulin OS=Homo sapiens OX=9606 GN=B2M PE=1 SV=1 | 38 | 122 | 4 | 8.57E+07 | 8.34E+07 | 8.73E+07 | 7.20E+07 | 6.87E+07 | 9.52E+07 | 4.37E+07 | 4.30E+07 | 2.96E+07 | 3.39E+07 | 3.59E+07 | 2.66E+07 | 8.20E+07 | 3.55E+07 | 2.31 | 1.21 | 6.24067E-06 |
| P51991 | Heterogeneous nuclear ribonucleoprotein A3 OS=Homo sapiens OX=9606 GN=HNRNPA3 PE=1 SV=2 | 22 | 53 | 4 | 1.06E+07 | 1.42E+07 | 5.58E+06 | 1.18E+07 | 8.52E+06 | 9.57E+06 | 3.34E+06 | 3.31E+06 | 4.09E+06 | 5.76E+06 | 1.42E+06 | 1.59E+06 | 1.00E+07 | 3.25E+06 | 3.09 | 1.63 | 0.001201704 |
| P61106 | Ras-related protein Rab-14 OS=Homo sapiens OX=9606 GN=RAB14 PE=1 SV=4 | 33 | 50 | 4 | 2.16E+07 | 1.61E+07 | 1.87E+07 | 1.71E+07 | 1.90E+07 | 1.78E+07 | 1.19E+07 | 1.11E+07 | 1.59E+07 | 1.58E+07 | 1.27E+07 | 9.22E+06 | 1.84E+07 | 1.28E+07 | 1.44 | 0.53 | 0.002222368 |
| O15143 | Actin-related protein 2/3 complex subunit 1B OS=Homo sapiens OX=9606 GN=ARPC1B PE=1 SV=3 | 19 | 92 | 4 | 1.58E+07 | 2.03E+07 | 1.94E+07 | 1.67E+07 | 1.58E+07 | 1.83E+07 | 1.51E+07 | 1.78E+07 | 1.57E+07 | 1.13E+07 | 1.16E+07 | 1.44E+07 | 1.77E+07 | 1.43E+07 | 1.24 | 0.31 | 0.025025205 |
| P06744 | Glucose-6-phosphate isomerase OS=Homo sapiens OX=9606 GN=GPI PE=1 SV=4 | 12 | 64 | 4 | 1.02E+07 | 6.92E+06 | 1.14E+07 | 1.81E+07 | 1.21E+07 | 8.26E+06 | 1.35E+07 | 1.11E+07 | 1.46E+07 | 1.01E+07 | 1.23E+07 | 1.27E+07 | 1.12E+07 | 1.24E+07 | 0.90 | -0.15 | 0.508463254 |
| P50914 | 60S ribosomal protein L14 OS=Homo sapiens OX=9606 GN=RPL14 PE=1 SV=4 | 22 | 166 | 4 | 6.57E+07 | 8.24E+07 | 7.31E+07 | 7.94E+07 | 7.33E+07 | 6.91E+07 | 4.28E+07 | 5.18E+07 | 4.04E+07 | 4.62E+07 | 3.82E+07 | 3.66E+07 | 7.38E+07 | 4.27E+07 | 1.73 | 0.79 | 4.06751E-06 |
| Q99829 | Copine-1 OS=Homo sapiens OX=9606 GN=CPNE1 PE=1 SV=1 | 10 | 62 | 4 | 1.28E+07 | 8.10E+06 | 1.00E+07 | 5.82E+06 | 6.55E+06 | 8.61E+06 | 7.79E+06 | 6.38E+06 | 4.78E+06 | 4.20E+06 | 6.42E+06 | 5.05E+06 | 8.64E+06 | 5.77E+06 | 1.50 | 0.58 | 0.039803457 |
| P05546 | Heparin cofactor 2 OS=Homo sapiens OX=9606 GN=SERPIND1 PE=1 SV=3 | 7 | 187 | 4 | 6.27E+07 | 7.02E+07 | 5.85E+07 | 7.13E+07 | 7.12E+07 | 6.78E+07 | 7.65E+07 | 7.93E+07 | 8.66E+07 | 8.90E+07 | 8.80E+07 | 8.06E+07 | 6.69E+07 | 8.33E+07 | 0.80 | -0.32 | 0.000284045 |
| Q04917 | 14-3-3 protein eta OS=Homo sapiens OX=9606 GN=YWHAH PE=1 SV=4 | 24 | 61 | 4 | 1.17E+07 | 1.25E+07 | 6.08E+06 | 8.65E+06 | 8.37E+06 | 7.92E+06 | 1.14E+07 | 9.22E+06 | 8.40E+06 | 1.05E+07 | 5.59E+06 | 6.22E+06 | 9.20E+06 | 8.55E+06 | 1.08 | 0.11 | 0.644696961 |
| P12268 | Inosine-5'-monophosphate dehydrogenase 2 OS=Homo sapiens OX=9606 GN=IMPDH2 PE=1 SV=2 | 14 | 70 | 4 | 1.79E+07 | 1.85E+07 | 1.46E+07 | 1.71E+07 | 1.61E+07 | 1.52E+07 | 1.28E+07 | 1.46E+07 | 1.05E+07 | 1.20E+07 | 1.07E+07 | 8.56E+06 | 1.66E+07 | 1.15E+07 | 1.44 | 0.52 | 0.000948165 |
| Q05682 | Caldesmon OS=Homo sapiens OX=9606 GN=CALD1 PE=1 SV=3 | 8 | 22 | 4 | 3.14E+06 | 2.45E+06 | 4.74E+06 | 4.89E+06 | 3.72E+06 | 2.84E+06 | 1.51E+06 | 1.70E+06 | 3.04E+06 | 1.12E+06 | 1.50E+06 | 1.02E+06 | 3.63E+06 | 1.65E+06 | 2.20 | 1.14 | 0.003523827 |
| P35221 | Catenin alpha-1 OS=Homo sapiens OX=9606 GN=CTNNA1 PE=1 SV=1 | 8 | 24 | 4 | 1.73E+07 | 1.91E+07 | 1.82E+07 | 1.91E+07 | 1.79E+07 | 1.73E+07 | 1.94E+07 | 2.04E+07 | 1.87E+07 | 2.31E+07 | 2.07E+07 | 1.83E+07 | 1.81E+07 | 2.00E+07 | 0.91 | -0.14 | 0.04429624 |
| P40261 | Nicotinamide N-methyltransferase OS=Homo sapiens OX=9606 GN=NNMT PE=1 SV=1 | 20 | 75 | 4 | 2.53E+07 | 2.90E+07 | 2.90E+07 | 2.78E+07 | 2.63E+07 | 2.82E+07 | 1.56E+07 | 1.46E+07 | 1.55E+07 | 1.51E+07 | 1.93E+07 | 1.51E+07 | 2.76E+07 | 1.55E+07 | 1.78 | 0.83 | 4.99702E-07 |
| P49720 | Proteasome subunit beta type-3 OS=Homo sapiens OX=9606 GN=PSMB3 PE=1 SV=2 | 29 | 50 | 4 | 1.07E+07 | 1.23E+07 | 9.89E+06 | 7.73E+06 | 8.75E+06 | 9.77E+06 | 1.07E+07 | 1.13E+07 | 9.39E+06 | 7.30E+06 | 1.08E+07 | 1.05E+07 | 9.87E+06 | 1.00E+07 | 0.99 | -0.02 | 0.878413799 |
| Q5JWF2 | Guanine nucleotide-binding protein G(s) subunit alpha isoforms XLas OS=Homo sapiens OX=9606 GN=GNAS PE=1 SV=2 | 6 | 69 | 4 | 9.22E+06 | 1.06E+07 | 8.95E+06 | 8.88E+06 | 8.47E+06 | 9.13E+06 | 8.37E+06 | 7.69E+06 | 6.63E+06 | 6.84E+06 | 6.43E+06 | 6.78E+06 | 9.21E+06 | 7.12E+06 | 1.29 | 0.37 | 0.000638556 |
| P20339 | Ras-related protein Rab-5A OS=Homo sapiens OX=9606 GN=RAB5A PE=1 SV=2 | 37 | 87 | 4 | 7.82E+06 | 8.49E+06 | 8.06E+06 | 8.14E+06 | 7.48E+06 | 8.52E+06 | 7.36E+06 | 6.87E+06 | 7.79E+06 | 5.80E+06 | 6.49E+06 | 6.94E+06 | 8.09E+06 | 6.87E+06 | 1.18 | 0.23 | 0.005963896 |
| O14974 | Protein phosphatase 1 regulatory subunit 12A OS=Homo sapiens OX=9606 GN=PPP1R12A PE=1 SV=1 | 4 | 20 | 4 | 7.81E+06 | 3.32E+06 | 5.41E+06 | 3.06E+06 | 2.63E+06 | 1.84E+06 | 5.00E+06 | 1.72E+06 | 9.14E+05 | 3.86E+06 | 7.49E+05 | 2.46E+06 | 4.01E+06 | 2.45E+06 | 1.64 | 0.71 | 0.200776314 |
| P02747 | Complement C1q subcomponent subunit C OS=Homo sapiens OX=9606 GN=C1QC PE=1 SV=3 | 21 | 36 | 4 | 1.20E+06 |  |  | 5.59E+05 | 8.09E+05 | 4.88E+05 | 1.88E+07 | 3.58E+06 | 3.10E+06 | 1.49E+06 | 2.53E+06 | 2.00E+08 | 7.64E+05 | 3.82E+07 | 0.02 | -5.64 | 0.300396701 |
| Q96A65 | Exocyst complex component 4 OS=Homo sapiens OX=9606 GN=EXOC4 PE=1 SV=1 | 7 | 14 | 4 | 3.90E+06 | 2.48E+06 | 2.69E+06 | 3.31E+06 | 3.06E+06 | 2.36E+06 | 1.73E+06 | 5.05E+05 | 3.82E+05 | 7.55E+05 | 3.17E+05 | 6.99E+05 | 2.97E+06 | 7.32E+05 | 4.06 | 2.02 | 3.84562E-05 |
| P00751 | Complement factor B OS=Homo sapiens OX=9606 GN=CFB PE=1 SV=2 | 9 | 84 | 4 | 1.61E+07 | 1.75E+07 | 1.46E+07 | 1.38E+07 | 1.40E+07 | 1.56E+07 | 1.82E+07 | 2.32E+07 | 2.41E+07 | 1.59E+07 | 2.44E+07 | 1.76E+07 | 1.53E+07 | 2.06E+07 | 0.74 | -0.43 | 0.015932008 |
| Q96QD8 | Sodium-coupled neutral amino acid transporter 2 OS=Homo sapiens OX=9606 GN=SLC38A2 PE=1 SV=2 | 12 | 42 | 4 | 1.44E+07 | 1.74E+07 | 1.34E+07 | 1.21E+07 | 1.25E+07 | 1.45E+07 | 8.73E+06 | 7.75E+06 | 6.00E+06 | 6.47E+06 | 4.88E+06 | 6.23E+06 | 1.41E+07 | 6.67E+06 | 2.11 | 1.07 | 2.88716E-05 |
| P15153 | Ras-related C3 botulinum toxin substrate 2 OS=Homo sapiens OX=9606 GN=RAC2 PE=1 SV=1 | 35 | 116 | 4 | 5.76E+07 | 5.51E+07 | 5.48E+07 | 6.74E+07 | 5.78E+07 | 5.91E+07 | 5.93E+07 | 5.96E+07 | 5.96E+07 | 7.03E+07 | 5.59E+07 | 5.52E+07 | 5.86E+07 | 6.00E+07 | 0.98 | -0.03 | 0.660346795 |
| P50552 | Vasodilator-stimulated phosphoprotein OS=Homo sapiens OX=9606 GN=VASP PE=1 SV=3 | 14 | 116 | 4 | 2.16E+07 | 2.12E+07 | 1.62E+07 | 1.55E+07 | 1.60E+07 | 1.82E+07 | 1.83E+07 | 1.77E+07 | 1.38E+07 | 1.78E+07 | 1.34E+07 | 1.67E+07 | 1.81E+07 | 1.63E+07 | 1.11 | 0.15 | 0.223982082 |
| P04424 | Argininosuccinate lyase OS=Homo sapiens OX=9606 GN=ASL PE=1 SV=4 | 11 | 52 | 4 | 1.16E+07 | 9.85E+06 | 1.17E+07 | 9.74E+06 | 1.01E+07 | 1.10E+07 | 1.82E+07 | 1.58E+07 | 1.62E+07 | 1.24E+07 | 1.43E+07 | 1.40E+07 | 1.07E+07 | 1.51E+07 | 0.70 | -0.51 | 0.001821252 |
| P12110 | Collagen alpha-2(VI) chain OS=Homo sapiens OX=9606 GN=COL6A2 PE=1 SV=4 | 5 | 88 | 4 | 1.06E+07 | 1.65E+07 | 9.88E+06 | 9.65E+06 | 1.20E+07 | 1.29E+07 | 2.75E+07 | 2.68E+07 | 1.86E+07 | 1.50E+07 | 2.42E+07 | 1.33E+07 | 1.19E+07 | 2.09E+07 | 0.57 | -0.81 | 0.013728395 |
| P05026 | Sodium/potassium-transporting ATPase subunit beta-1 OS=Homo sapiens OX=9606 GN=ATP1B1 PE=1 SV=1 | 21 | 39 | 4 | 8.26E+06 | 6.25E+06 | 1.06E+07 | 8.44E+06 | 8.98E+06 | 7.77E+06 | 6.14E+06 | 4.68E+06 | 4.16E+06 | 4.03E+06 | 2.38E+06 | 3.82E+06 | 8.38E+06 | 4.20E+06 | 2.00 | 1.00 | 0.000305952 |
| Q9Y5L0 | Transportin-3 OS=Homo sapiens OX=9606 GN=TNPO3 PE=1 SV=3 | 8 | 16 | 4 | 2.00E+06 | 2.10E+06 | 2.59E+06 | 8.89E+05 | 2.86E+06 | 2.66E+06 | 4.95E+05 |  | 1.49E+06 | 6.06E+05 | 6.88E+05 | 3.36E+05 | 2.18E+06 | 7.22E+05 | 3.02 | 1.60 | 0.002960487 |
| O15162 | Phospholipid scramblase 1 OS=Homo sapiens OX=9606 GN=PLSCR1 PE=1 SV=1 | 12 | 73 | 4 | 3.81E+07 | 3.25E+07 | 2.88E+07 | 2.88E+07 | 2.37E+07 | 3.09E+07 | 2.13E+07 | 1.74E+07 | 2.00E+07 | 1.29E+07 | 2.29E+07 | 1.58E+07 | 3.05E+07 | 1.84E+07 | 1.66 | 0.73 | 0.00074725 |
| P51114 | Fragile X mental retardation syndrome-related protein 1 OS=Homo sapiens OX=9606 GN=FXR1 PE=1 SV=3 | 11 | 78 | 4 | 1.36E+07 | 1.65E+07 | 1.55E+07 | 1.77E+07 | 1.65E+07 | 1.44E+07 | 7.18E+06 | 7.78E+06 | 7.57E+06 | 7.94E+06 | 7.71E+06 | 6.41E+06 | 1.57E+07 | 7.43E+06 | 2.11 | 1.08 | 1.06546E-05 |
| P62854 | 40S ribosomal protein S26 OS=Homo sapiens OX=9606 GN=RPS26 PE=1 SV=3 | 44 | 123 | 4 | 5.79E+07 | 5.22E+07 | 4.75E+07 | 5.47E+07 | 5.00E+07 | 4.84E+07 | 3.36E+07 | 3.61E+07 | 2.95E+07 | 3.09E+07 | 3.01E+07 | 2.54E+07 | 5.18E+07 | 3.09E+07 | 1.67 | 0.74 | 2.71958E-06 |
| P67870 | Casein kinase II subunit beta OS=Homo sapiens OX=9606 GN=CSNK2B PE=1 SV=1 | 23 | 67 | 4 | 2.70E+07 | 3.08E+07 | 2.76E+07 | 3.10E+07 | 3.02E+07 | 2.74E+07 | 1.73E+07 | 1.80E+07 | 1.96E+07 | 1.83E+07 | 1.64E+07 | 1.37E+07 | 2.90E+07 | 1.72E+07 | 1.69 | 0.75 | 1.10671E-06 |
| P48740 | Mannan-binding lectin serine protease 1 OS=Homo sapiens OX=9606 GN=MASP1 PE=1 SV=3 | 9 | 102 | 4 | 3.53E+07 | 3.67E+07 | 3.73E+07 | 4.34E+07 | 3.84E+07 | 4.64E+07 | 4.52E+07 | 4.49E+07 | 4.72E+07 | 4.49E+07 | 4.97E+07 | 3.90E+07 | 3.96E+07 | 4.51E+07 | 0.88 | -0.19 | 0.037053961 |

### Supplementary Table 4. List of all proteins identified by label free proteomics analysis.

| Accession | Description | Coverage [%] | # PSMs | # Unique Peptides | Amitriptyline 1 | Amitriptyline 2 | Amitriptyline 3 | Amitriptyline 4 | Amitriptyline 5 | Amitriptyline 6 | Vehicle 1 | Vehicle 2 | Vehicle 3 | Vehicle 4 | Vehicle 5 | Vehicle 6 | Average amitriptyline | Average vehicle | Ratio amitriptyline/ vehicle | Log 2 fold change (amitriptyline /vehicle) | P-value |
|---|---|---|---|---|---|---|---|---|---|---|---|---|---|---|---|---|---|---|---|---|---|
| Q9BXJ4 | Complement C1q tumor necrosis factor-related protein 3 OS=Homo sapiens OX=9606 GN=C1QTNF3 PE=1 SV=1 | 13 | 74 | 4 | 2.71E+07 | 2.37E+07 | 2.01E+07 | 2.30E+07 | 2.01E+07 | 1.75E+07 | 3.27E+07 | 2.61E+07 | 2.44E+07 | 2.18E+07 | 1.86E+07 | 1.76E+07 | 2.19E+07 | 2.35E+07 | 0.93 | -0.10 | 0.557221439 |
| Q92542 | Nicastrin OS=Homo sapiens OX=9606 GN=NCSTN PE=1 SV=2 | 10 | 10 | 4 | 6.13E+06 | 4.68E+06 | 4.07E+06 | 4.39E+06 | 5.85E+06 | 5.45E+06 | 1.24E+06 | 1.39E+06 | 1.33E+06 | 2.15E+06 | 1.36E+06 | 1.84E+06 | 5.09E+06 | 1.55E+06 | 3.28 | 1.72 | 3.50612E-05 |
| P00738 | Haptoglobin OS=Homo sapiens OX=9606 GN=HP PE=1 SV=1 | 10 | 91 | 4 | 3.01E+07 | 2.98E+07 | 2.57E+07 | 2.76E+07 | 2.37E+07 | 1.92E+07 | 3.19E+07 | 2.94E+07 | 2.56E+07 | 3.06E+07 | 2.07E+07 | 1.82E+07 | 2.60E+07 | 2.61E+07 | 1.00 | 0.00 | 0.989369582 |
| Q13409 | Cytoplasmic dynein 1 intermediate chain 2 OS=Homo sapiens OX=9606 GN=DYNC1I2 PE=1 SV=3 | 12 | 12 | 4 | 3.06E+06 | 5.11E+06 | 3.45E+06 | 2.03E+06 | 2.03E+06 | 4.08E+06 | | 1.41E+06 | 1.86E+06 | 1.60E+06 | 3.79E+06 | 2.48E+06 | 3.29E+06 | 2.23E+06 | 1.48 | 0.56 | 0.136684388 |
| P16070 | CD44 antigen OS=Homo sapiens OX=9606 GN=CD44 PE=1 SV=3 | 6 | 142 | 4 | 4.09E+07 | 4.02E+07 | 3.89E+07 | 4.50E+07 | 4.18E+07 | 4.64E+07 | 2.12E+07 | 2.21E+07 | 2.06E+07 | 2.05E+07 | 1.83E+07 | 1.65E+07 | 4.22E+07 | 1.99E+07 | 2.12 | 1.09 | 8.68511E-08 |
| O96019 | Actin-like protein 6A OS=Homo sapiens OX=9606 GN=ACTL6A PE=1 SV=1 | 16 | 54 | 4 | 1.66E+07 | 1.95E+07 | 1.16E+07 | 8.42E+06 | 1.24E+07 | 1.45E+07 | 1.12E+07 | 1.43E+07 | 9.68E+06 | 8.90E+06 | 9.96E+06 | 9.43E+06 | 1.38E+07 | 1.06E+07 | 1.31 | 0.39 | 0.109866641 |
| P26022 | Pentraxin-related protein PTX3 OS=Homo sapiens OX=9606 GN=PTX3 PE=1 SV=1 | 14 | 28 | 4 | 1.67E+06 | 1.19E+06 | 4.85E+05 | 1.35E+06 | 1.65E+06 | 3.47E+06 | 9.31E+06 | 1.07E+07 | 5.79E+06 | 7.26E+06 | 7.28E+06 | 7.29E+06 | 1.64E+06 | 7.94E+06 | 0.21 | -2.28 | 6.25913E-05 |
| Q99805 | Transmembrane 9 superfamily member 2 OS=Homo sapiens OX=9606 GN=TM9SF2 PE=1 SV=1 | 9 | 48 | 4 | 8.56E+06 | 7.88E+06 | 8.28E+06 | 8.04E+06 | 7.52E+06 | 8.58E+06 | 6.05E+06 | 6.04E+06 | 4.48E+06 | 5.49E+06 | 5.60E+06 | 4.17E+06 | 8.14E+06 | 5.30E+06 | 1.54 | 0.62 | 7.88823E-05 |
| Q00577 | Transcriptional activator protein Pur-alpha OS=Homo sapiens OX=9606 GN=PURA PE=1 SV=2 | 25 | 22 | 4 | 1.05E+07 | 1.09E+07 | 5.85E+06 | 5.95E+06 | 6.16E+06 | 7.87E+06 | 9.49E+06 | 8.84E+06 | 5.97E+06 | 9.55E+06 | 6.46E+06 | 5.28E+06 | 7.87E+06 | 7.60E+06 | 1.04 | 0.05 | 0.829657978 |
| Q92522 | Histone H1x OS=Homo sapiens OX=9606 GN=H1FX PE=1 SV=1 | 16 | 114 | 4 | 5.32E+07 | 5.80E+07 | 5.83E+07 | 4.88E+07 | 4.63E+07 | 4.95E+07 | 4.97E+07 | 5.09E+07 | 4.75E+07 | 4.23E+07 | 4.02E+07 | 3.82E+07 | 5.23E+07 | 4.48E+07 | 1.17 | 0.22 | 0.029697859 |
| Q13085 | Acetyl-CoA carboxylase 1 OS=Homo sapiens OX=9606 GN=ACACA PE=1 SV=2 | 3 | 23 | 4 | 4.93E+06 | 5.51E+06 | 6.90E+06 | 5.10E+06 | 6.78E+06 | 4.51E+06 | 5.26E+06 | 5.87E+06 | 5.36E+06 | 6.02E+06 | 6.61E+06 | 5.10E+06 | 5.62E+06 | 5.70E+06 | 0.99 | -0.02 | 0.867454914 |
| P61289 | Proteasome activator complex subunit 3 OS=Homo sapiens OX=9606 GN=PSME3 PE=1 SV=1 | 17 | 73 | 4 | 1.15E+07 | 1.43E+07 | 9.00E+06 | 1.23E+07 | 1.16E+07 | 9.75E+06 | 9.43E+06 | 8.79E+06 | 8.27E+06 | 6.82E+06 | 5.64E+06 | 5.68E+06 | 1.14E+07 | 7.44E+06 | 1.53 | 0.62 | 0.003058654 |
| P04632 | Calpain small subunit 1 OS=Homo sapiens OX=9606 GN=CAPNS1 PE=1 SV=1 | 20 | 32 | 4 | 8.09E+06 | 6.43E+06 | 7.89E+06 | 7.54E+06 | 7.83E+06 | 7.12E+06 | 5.44E+06 | 4.11E+06 | 4.42E+06 | 5.24E+06 | 5.50E+06 | 4.33E+06 | 7.48E+06 | 4.84E+06 | 1.55 | 0.63 | 2.26971E-05 |
| Q7Z2W4 | Zinc finger CCCH-type antiviral protein 1 OS=Homo sapiens OX=9606 GN=ZC3HAV1 PE=1 SV=3 | 6 | 42 | 4 | 7.69E+06 | 8.72E+06 | 7.96E+06 | 9.03E+06 | 7.56E+06 | 8.36E+06 | 3.36E+06 | 3.87E+06 | 4.57E+06 | 4.65E+06 | 4.58E+06 | 4.44E+06 | 8.22E+06 | 4.24E+06 | 1.94 | 0.95 | 2.32232E-07 |
| Q8NE71 | ATP-binding cassette sub-family F member 1 OS=Homo sapiens OX=9606 GN=ABCF1 PE=1 SV=2 | 8 | 20 | 4 | 4.76E+06 | 3.93E+06 | 5.03E+06 | 6.96E+06 | 5.55E+06 | 5.82E+06 | 3.85E+06 | 1.87E+06 | 3.64E+06 | 3.15E+06 | 4.97E+05 | 1.70E+06 | 5.34E+06 | 2.45E+06 | 2.18 | 1.12 | 0.001916005 |
| P62888 | 60S ribosomal protein L30 OS=Homo sapiens OX=9606 GN=RPL30 PE=1 SV=2 | 51 | 62 | 4 | 1.83E+07 | 1.70E+07 | 1.40E+07 | 1.46E+07 | 1.48E+07 | 1.72E+07 | 1.37E+07 | 1.35E+07 | 8.98E+06 | 1.18E+07 | 1.13E+07 | 9.87E+06 | 1.60E+07 | 1.15E+07 | 1.39 | 0.47 | 0.001729187 |
| P40763 | Signal transducer and activator of transcription 3 OS=Homo sapiens OX=9606 GN=STAT3 PE=1 SV=2 | 9 | 34 | 4 | 4.89E+06 | 6.78E+06 | 5.04E+06 | 5.30E+06 | 5.42E+06 | 4.39E+06 | 5.88E+06 | 7.17E+06 | 5.89E+06 | 6.80E+06 | 5.99E+06 | 5.98E+06 | 5.30E+06 | 6.28E+06 | 0.84 | -0.24 | 0.037770284 |
| Q96FW1 | Ubiquitin thioesterase OTUB1 OS=Homo sapiens OX=9606 GN=OTUB1 PE=1 SV=2 | 19 | 26 | 4 | 7.77E+06 | 7.05E+06 | 8.09E+06 | 1.01E+07 | 7.81E+06 | 7.74E+06 | 8.51E+06 | 8.36E+06 | 8.38E+06 | 7.84E+06 | 7.69E+06 | 7.06E+06 | 8.10E+06 | 7.97E+06 | 1.02 | 0.02 | 0.802514473 |
| Q12905 | Interleukin enhancer-binding factor 2 OS=Homo sapiens OX=9606 GN=ILF2 PE=1 SV=2 | 15 | 42 | 4 | 9.74E+06 | 9.56E+06 | 1.30E+07 | 1.07E+07 | 7.77E+06 | 8.11E+06 | 1.26E+07 | 9.84E+06 | 9.95E+06 | 5.79E+06 | 1.09E+07 | 8.63E+06 | 9.81E+06 | 9.62E+06 | 1.02 | 0.03 | 0.874965993 |
| O75340 | Programmed cell death protein 6 OS=Homo sapiens OX=9606 GN=PDCD6 PE=1 SV=1 | 31 | 84 | 4 | 2.20E+07 | 2.25E+07 | 2.36E+07 | 1.97E+07 | 1.96E+07 | 1.77E+07 | 1.48E+07 | 1.55E+07 | 1.21E+07 | 1.40E+07 | 1.22E+07 | 2.05E+07 | 2.08E+07 | 1.48E+07 | 1.40 | 0.49 | 0.003738271 |
| O95834 | Echinoderm microtubule-associated protein-like 2 OS=Homo sapiens OX=9606 GN=EML2 PE=1 SV=1 | 9 | 36 | 4 | 7.87E+06 | 8.61E+06 | 7.93E+06 | 6.34E+06 | 5.89E+06 | 6.77E+06 | 9.98E+06 | 1.04E+07 | 7.77E+06 | 7.49E+06 | 7.60E+06 | 7.94E+06 | 7.24E+06 | 8.53E+06 | 0.85 | -0.24 | 0.090156155 |
| P62913 | 60S ribosomal protein L11 OS=Homo sapiens OX=9606 GN=RPL11 PE=1 SV=2 | 31 | 82 | 4 | 1.72E+07 | 1.78E+07 | 1.51E+07 | 1.35E+07 | 1.66E+07 | 1.33E+07 | 1.57E+07 | 1.50E+07 | 1.37E+07 | 1.35E+07 | 1.75E+07 | 1.87E+07 | 1.56E+07 | 1.57E+07 | 0.99 | -0.01 | 0.912240929 |
| Q9Y5P6 | Mannose-1-phosphate guanyltransferase beta OS=Homo sapiens OX=9606 GN=GMPPB PE=1 SV=2 | 17 | 30 | 4 | 3.88E+06 | 4.75E+06 | 4.91E+06 | 3.60E+06 | 3.26E+06 | 3.76E+06 | 5.02E+06 | 5.78E+06 | 4.11E+06 | 6.92E+06 | 4.21E+06 | 3.37E+06 | 4.03E+06 | 4.90E+06 | 0.82 | -0.28 | 0.179544866 |
| Q9NR31 | GTP-binding protein SAR1a OS=Homo sapiens OX=9606 GN=SAR1A PE=1 SV=1 | 31 | 64 | 4 | 1.40E+07 | 1.02E+07 | 9.49E+06 | 8.31E+06 | 9.59E+06 | 1.14E+07 | 9.35E+06 | 8.75E+06 | 6.72E+06 | 5.95E+06 | 9.36E+06 | 8.26E+06 | 1.05E+07 | 8.06E+06 | 1.30 | 0.38 | 0.038075372 |
| O14950 | Myosin regulatory light chain 12B OS=Homo sapiens OX=9606 GN=MYL12B PE=1 SV=2 | 29 | 65 | 4 | 2.18E+07 | 2.16E+07 | 2.18E+07 | 2.51E+07 | 2.14E+07 | 2.17E+07 | 2.15E+07 | 2.20E+07 | 2.21E+07 | 1.94E+07 | 1.84E+07 | 1.67E+07 | 2.22E+07 | 2.00E+07 | 1.11 | 0.15 | 0.068965072 |
| P28072 | Proteasome subunit beta type-6 OS=Homo sapiens OX=9606 GN=PSMB6 PE=1 SV=4 | 29 | 76 | 4 | 2.32E+07 | 1.50E+07 | 1.33E+07 | 1.20E+07 | 1.37E+07 | 1.75E+07 | 2.40E+07 | 2.29E+07 | 2.46E+07 | 2.00E+07 | 1.76E+07 | 1.29E+07 | 1.58E+07 | 2.04E+07 | 0.77 | -0.37 | 0.093320352 |
| Q07666 | KH domain-containing, RNA-binding, signal transduction-associated protein 1 OS=Homo sapiens OX=9606 GN=KHDRBS1 PE=1 SV=1 | 12 | 30 | 4 | 1.23E+07 | 1.23E+07 | 1.18E+07 | 1.27E+07 | 1.02E+07 | 1.16E+07 | 8.84E+06 | 8.75E+06 | 8.21E+06 | 8.68E+06 | 7.06E+06 | 7.06E+06 | 1.18E+07 | 8.10E+06 | 1.46 | 0.54 | 2.24987E-05 |
| P10809 | 60 kDa heat shock protein, mitochondrial OS=Homo sapiens OX=9606 GN=HSPD1 PE=1 SV=2 | 11 | 45 | 4 | 5.21E+06 | 1.06E+07 | 5.19E+06 | 4.57E+06 | 4.18E+06 | 3.87E+06 | 9.21E+06 | 6.96E+06 | 7.72E+06 | 9.40E+06 | 6.51E+06 | 9.72E+06 | 5.61E+06 | 8.25E+06 | 0.68 | -0.56 | 0.05428114 |
| P02746 | Complement C1q subcomponent subunit B OS=Homo sapiens OX=9606 GN=C1QB PE=1 SV=3 | 23 | 24 | 4 | 7.51E+05 | 9.34E+05 | 6.46E+05 | 6.95E+05 | 9.15E+05 | 9.43E+05 | 8.47E+06 | 1.79E+06 | 9.93E+05 | 1.73E+06 | 2.27E+06 | 4.98E+07 | 8.14E+05 | 1.08E+07 | 0.08 | -3.73 | 0.258666568 |
| Q13740 | CD166 antigen OS=Homo sapiens OX=9606 GN=ALCAM PE=1 SV=2 | 13 | 16 | 4 | 3.15E+06 | 4.11E+06 | 2.82E+06 | 2.98E+06 | 2.82E+06 | 2.11E+06 | 1.52E+06 | 2.02E+06 | 1.02E+06 | 1.28E+06 | 1.16E+06 | 8.32E+05 | 3.00E+06 | 1.30E+06 | 2.30 | 1.20 | 0.000537756 |
| Q9Y285 | Phenylalanine--tRNA ligase alpha subunit OS=Homo sapiens OX=9606 GN=FARSA PE=1 SV=3 | 11 | 39 | 4 | 1.39E+07 | 1.68E+07 | 1.53E+07 | 1.57E+07 | 1.61E+07 | 1.53E+07 | 1.18E+07 | 1.20E+07 | 8.52E+06 | 1.25E+07 | 1.05E+07 | 1.06E+07 | 1.55E+07 | 1.10E+07 | 1.41 | 0.50 | 0.000155366 |
| O15427 | Monocarboxylate transporter 4 OS=Homo sapiens OX=9606 GN=SLC16A3 PE=1 SV=1 | 10 | 63 | 4 | 1.25E+07 | 1.30E+07 | 1.28E+07 | 1.58E+07 | 1.56E+07 | 1.05E+07 | 9.39E+06 | 7.77E+06 | 1.10E+07 | 1.04E+07 | 8.47E+06 | 5.79E+06 | 1.34E+07 | 8.81E+06 | 1.52 | 0.60 | 0.002350571 |
| P22570 | NADPH:adrenodoxin oxidoreductase, mitochondrial OS=Homo sapiens OX=9606 GN=FDXR PE=1 SV=3 | 12 | 50 | 4 | 1.21E+07 | 7.90E+06 | 1.19E+07 | 1.19E+07 | 1.21E+07 | 6.34E+06 | 1.07E+07 | 1.04E+07 | 1.08E+07 | 8.30E+06 | 8.77E+06 | 4.87E+06 | 1.04E+07 | 8.98E+06 | 1.16 | 0.21 | 0.339028786 |
| P62244 | 40S ribosomal protein S15a OS=Homo sapiens OX=9606 GN=RPS15A PE=1 SV=2 | 31 | 146 | 4 | 5.17E+07 | 5.71E+07 | 7.44E+07 | 7.29E+07 | 7.60E+07 | 6.92E+07 | 4.27E+07 | 4.70E+07 | 5.73E+07 | 5.67E+07 | 5.65E+07 | 4.01E+07 | 6.69E+07 | 5.01E+07 | 1.34 | 0.42 | 0.009614808 |
| Q14566 | DNA replication licensing factor MCM6 OS=Homo sapiens OX=9606 GN=MCM6 PE=1 SV=1 | 6 | 20 | 4 | 3.66E+06 | 3.09E+06 | 3.45E+06 | 3.92E+06 | 3.34E+06 | 3.12E+06 | 9.27E+05 | 1.93E+06 | 9.75E+05 | | 1.21E+06 | 2.12E+06 | 3.43E+06 | 1.43E+06 | 2.39 | 1.26 | 0.000356056 |
| Q9HD45 | Transmembrane 9 superfamily member 3 OS=Homo sapiens OX=9606 GN=TM9SF3 PE=1 SV=2 | 9 | 46 | 4 | 5.28E+06 | 5.08E+06 | 6.56E+06 | 5.20E+06 | 6.47E+06 | 7.45E+06 | 4.77E+06 | 3.69E+06 | 4.83E+06 | 6.96E+06 | 5.14E+06 | 3.90E+06 | 6.01E+06 | 4.88E+06 | 1.23 | 0.30 | 0.099303025 |
| P63151 | Serine/threonine-protein phosphatase 2A 55 kDa regulatory subunit B alpha isoform OS=Homo sapiens OX=9606 GN=PPP2R2A PE=1 SV=1 | 13 | 34 | 4 | 4.89E+06 | 4.59E+06 | 7.21E+06 | 7.90E+06 | 6.68E+06 | 8.53E+06 | 5.94E+06 | 4.81E+06 | 7.54E+06 | 6.18E+06 | 5.85E+06 | 4.81E+06 | 6.63E+06 | 5.86E+06 | 1.13 | 0.18 | 0.343088155 |
| P29401 | Transketolase OS=Homo sapiens OX=9606 GN=TKT PE=1 SV=3 | 9 | 52 | 4 | 5.87E+06 | 5.77E+06 | 7.08E+06 | 5.03E+06 | 5.63E+06 | 4.09E+06 | 6.79E+06 | 5.49E+06 | 6.14E+06 | 6.42E+06 | 6.29E+06 | 6.94E+05 | 5.58E+06 | 5.30E+06 | 1.05 | 0.07 | 0.796319818 |
| Q15691 | Microtubule-associated protein RP/EB family member 1 OS=Homo sapiens OX=9606 GN=MAPRE1 PE=1 SV=3 | 26 | 40 | 4 | 9.57E+06 | 8.08E+06 | 9.56E+06 | 9.00E+06 | 9.62E+06 | 9.56E+06 | 5.66E+06 | 5.38E+06 | 7.12E+06 | 5.99E+06 | 4.59E+06 | 4.12E+06 | 9.23E+06 | 5.48E+06 | 1.69 | 0.75 | 6.85236E-05 |
| P50502 | Hsc70-interacting protein OS=Homo sapiens OX=9606 GN=ST13 PE=1 SV=2 | 9 | 81 | 4 | 1.68E+07 | 2.02E+07 | 1.98E+07 | 2.09E+07 | 1.59E+07 | 1.61E+07 | 2.00E+07 | 1.97E+07 | 9.00E+06 | 2.11E+07 | 1.71E+07 | 1.94E+07 | 1.83E+07 | 1.77E+07 | 1.03 | 0.05 | 0.784924673 |
| P00352 | Retinal dehydrogenase 1 OS=Homo sapiens OX=9606 GN=ALDH1A1 PE=1 SV=2 | 9 | 24 | 4 | 1.10E+07 | 1.22E+07 | 1.25E+07 | 1.39E+07 | 1.16E+07 | 1.12E+07 | 1.85E+07 | 1.59E+07 | 1.91E+07 | 1.78E+07 | 1.63E+07 | 1.45E+07 | 1.21E+07 | 1.70E+07 | 0.71 | -0.50 | 0.000290323 |
| O15031 | Plexin-B2 OS=Homo sapiens OX=9606 GN=PLXNB2 PE=1 SV=3 | 4 | 48 | 4 | 8.91E+06 | 7.33E+06 | 6.55E+06 | 8.93E+06 | 9.15E+06 | 7.05E+06 | 9.00E+06 | 1.05E+07 | 1.21E+07 | 1.14E+07 | 9.17E+06 | 9.01E+06 | 7.99E+06 | 1.02E+07 | 0.78 | -0.35 | 0.012249348 |
| P14678 | Small nuclear ribonucleoprotein-associated proteins B and B' OS=Homo sapiens OX=9606 GN=SNRPB PE=1 SV=2 | 15 | 139 | 4 | 2.24E+07 | 2.32E+07 | 2.44E+07 | 2.19E+07 | 2.61E+07 | 2.50E+07 | 1.87E+07 | 1.65E+07 | 1.69E+07 | 1.75E+07 | 2.36E+07 | 1.87E+07 | 2.38E+07 | 1.86E+07 | 1.28 | 0.36 | 0.002776556 |
| P18621 | 60S ribosomal protein L17 OS=Homo sapiens OX=9606 GN=RPL17 PE=1 SV=3 | 27 | 22 | 4 | 1.17E+07 | 1.70E+07 | 4.19E+06 | 9.24E+06 | 9.04E+06 | 1.48E+07 | 2.19E+06 | 1.57E+07 | 5.12E+06 | 4.71E+06 | 7.71E+06 | 7.75E+06 | 1.10E+07 | 7.20E+06 | 1.53 | 0.61 | 0.186243026 |
| Q15113 | Procollagen C-endopeptidase enhancer 1 OS=Homo sapiens OX=9606 GN=PCOLCE PE=1 SV=2 | 14 | 46 | 4 | 4.40E+06 | 4.91E+06 | 2.93E+06 | 3.58E+06 | 2.77E+06 | 6.22E+06 | 6.21E+06 | 6.35E+06 | 3.06E+06 | 4.11E+06 | 4.09E+06 | 7.38E+06 | 4.14E+06 | 5.20E+06 | 0.79 | -0.33 | 0.249950727 |
| P61604 | 10 kDa heat shock protein, mitochondrial OS=Homo sapiens OX=9606 GN=HSPE1 PE=1 SV=2 | 33 | 64 | 4 | 1.29E+07 | 1.38E+07 | 1.49E+07 | 1.72E+07 | 1.52E+07 | 2.10E+07 | 1.11E+07 | 1.06E+07 | 1.02E+07 | 9.13E+06 | 8.29E+06 | 7.45E+06 | 1.58E+07 | 9.46E+06 | 1.67 | 0.74 | 0.001754542 |
| Q9NRN5 | Olfactomedin-like protein 3 OS=Homo sapiens OX=9606 GN=OLFML3 PE=2 SV=1 | 12 | 43 | 4 | 2.80E+07 | 2.63E+07 | 2.12E+07 | 2.22E+07 | 2.27E+07 | 2.10E+07 | 2.37E+07 | 1.96E+07 | 2.34E+07 | 2.15E+07 | 2.13E+07 | 9.08E+06 | 2.36E+07 | 1.98E+07 | 1.19 | 0.26 | 0.169956284 |
| Q10471 | Polypeptide N-acetylgalactosaminyltransferase 2 OS=Homo sapiens OX=9606 GN=GALNT2 PE=1 SV=1 | 10 | 10 | 4 | 4.49E+06 | 5.16E+06 | 3.97E+06 | 2.30E+06 | 4.45E+06 | 4.03E+06 | 4.06E+06 | 3.02E+06 | 3.67E+06 | 3.68E+06 | 1.92E+06 | 4.25E+06 | 4.07E+06 | 3.43E+06 | 1.18 | 0.24 | 0.255727416 |
| O43172 | U4/U6 small nuclear ribonucleoprotein Prp4 OS=Homo sapiens OX=9606 GN=PRPF4 PE=1 SV=2 | 10 | 16 | 4 | 3.91E+06 | 1.76E+06 | 2.82E+06 | 3.95E+06 | 3.41E+06 | 3.17E+06 | 1.37E+06 | 8.87E+05 | 2.36E+06 | 2.43E+06 | 1.17E+06 | 1.74E+06 | 3.17E+06 | 1.66E+06 | 1.91 | 0.94 | 0.005380444 |
| P63241 | Eukaryotic translation initiation factor 5A-1 OS=Homo sapiens OX=9606 GN=EIF5A PE=1 SV=2 | 36 | 65 | 4 | 1.29E+07 | 1.75E+07 | 1.41E+07 | 1.58E+07 | 1.64E+07 | 1.54E+07 | 1.13E+07 | 1.47E+07 | 1.35E+07 | 1.67E+07 | 1.11E+07 | 1.29E+07 | 1.54E+07 | 1.34E+07 | 1.15 | 0.20 | 0.100193863 |
| Q99961 | Endophilin-A2 OS=Homo sapiens OX=9606 GN=SH3GL1 PE=1 SV=1 | 12 | 24 | 4 | 4.10E+06 | 3.92E+06 | 3.92E+06 | 3.66E+06 | 3.99E+06 | 3.90E+06 | 2.22E+06 | 2.25E+06 | 2.24E+06 | 2.23E+06 | 1.59E+06 | 2.18E+06 | 3.92E+06 | 2.12E+06 | 1.85 | 0.89 | 5.87087E-07 |
| P45974 | Ubiquitin carboxyl-terminal hydrolase 5 OS=Homo sapiens OX=9606 GN=USP5 PE=1 SV=2 | 5 | 42 | 4 | 5.82E+06 | 4.42E+06 | 5.58E+06 | 5.72E+06 | 5.73E+06 | 4.24E+06 | 7.36E+06 | 7.47E+06 | 7.64E+06 | 5.86E+06 | 6.61E+06 | 5.56E+06 | 5.25E+06 | 6.75E+06 | 0.78 | -0.36 | 0.009644704 |
| P53004 | Biliverdin reductase A OS=Homo sapiens OX=9606 GN=BLVRA PE=1 SV=2 | 18 | 50 | 4 | 4.20E+06 | 7.30E+06 | 4.04E+06 | 4.78E+06 | 6.60E+06 | 6.61E+06 | 5.90E+06 | 6.29E+06 | 5.99E+06 | 5.71E+06 | 5.53E+06 | 4.76E+06 | 5.59E+06 | 5.69E+06 | 0.98 | -0.03 | 0.867922569 |
| P46779 | 60S ribosomal protein L28 OS=Homo sapiens OX=9606 GN=RPL28 PE=1 SV=3 | 28 | 52 | 4 | 1.67E+07 | 1.58E+07 | 1.92E+07 | 2.10E+07 | 1.93E+07 | 1.45E+07 | 1.38E+07 | 1.34E+07 | 1.60E+07 | 1.47E+07 | 1.25E+07 | 1.13E+06 | 1.77E+07 | 1.22E+07 | 1.45 | 0.54 | 0.034339462 |
| O43747 | AP-1 complex subunit gamma-1 OS=Homo sapiens OX=9606 GN=AP1G1 PE=1 SV=5 | 6 | 22 | 4 | 1.90E+06 | 4.62E+06 | 7.73E+06 | 7.88E+06 | 6.92E+06 | 6.17E+06 | 2.42E+06 | 2.18E+06 | 3.07E+06 | 3.11E+06 | 4.21E+06 | 4.44E+06 | 5.87E+06 | 3.24E+06 | 1.81 | 0.86 | 0.036294548 |
| Q9H0B6 | Kinesin light chain 2 OS=Homo sapiens OX=9606 GN=KLC2 PE=1 SV=1 | 10 | 24 | 4 | 4.20E+06 | 5.44E+06 | 6.52E+06 | 6.42E+06 | 4.24E+06 | 6.07E+06 | 3.00E+06 | 5.02E+06 | 2.95E+06 | | 4.86E+06 | 2.91E+06 | 5.48E+06 | 3.75E+06 | 1.46 | 0.55 | 0.026893434 |
| Q8IZP0 | Abl interactor 1 OS=Homo sapiens OX=9606 GN=ABI1 PE=1 SV=4 | 10 | 14 | 4 | 5.98E+06 | 6.32E+06 | 6.96E+06 | 6.53E+06 | 6.35E+06 | 4.84E+06 | 4.59E+06 | 4.31E+06 | 3.88E+06 | 3.94E+06 | 3.24E+06 | 4.05E+06 | 6.16E+06 | 4.00E+06 | 1.54 | 0.62 | 0.000210215 |
| P07358 | Complement component C8 beta chain OS=Homo sapiens OX=9606 GN=C8B PE=1 SV=3 | 11 | 21 | 4 | 3.19E+06 | 2.70E+06 | 4.11E+06 | 4.15E+06 | 3.58E+06 | 4.54E+06 | 5.86E+06 | 5.75E+06 | 5.26E+06 | 6.23E+06 | 5.02E+06 | 4.42E+06 | 3.72E+06 | 5.42E+06 | 0.69 | -0.54 | 0.001212776 |
| P11717 | Cation-independent mannose-6-phosphate receptor OS=Homo sapiens OX=9606 GN=IGF2R PE=1 SV=3 | 2 | 54 | 4 | 1.45E+07 | 1.28E+07 | 1.14E+07 | 1.28E+07 | 1.25E+07 | 1.12E+07 | 2.01E+07 | 1.86E+07 | 1.72E+07 | 2.01E+07 | 1.33E+07 | 1.42E+07 | 1.25E+07 | 1.73E+07 | 0.73 | -0.46 | 0.008936043 |
| Q8WVM8 | Sec1 family domain-containing protein 1 OS=Homo sapiens OX=9606 GN=SCFD1 PE=1 SV=4 | 12 | 8 | 4 | 1.45E+06 | 1.85E+06 | 1.41E+06 | 2.05E+06 | 1.86E+06 | 1.80E+06 | 1.40E+06 | 3.64E+06 | 1.05E+06 | 1.25E+06 | 1.35E+06 | 2.05E+06 | 1.74E+06 | 1.79E+06 | 0.97 | -0.04 | 0.901008575 |
| Q15493 | Regucalcin OS=Homo sapiens OX=9606 GN=RGN PE=1 SV=1 | 15 | 64 | 4 | 2.54E+07 | 2.72E+07 | 2.20E+07 | 3.09E+07 | 2.39E+07 | 2.45E+07 | 3.19E+07 | 3.60E+07 | 3.61E+07 | 3.89E+07 | 2.73E+07 | 2.92E+07 | 2.57E+07 | 3.32E+07 | 0.77 | -0.37 | 0.007920422 |
| P54619 | 5'-AMP-activated protein kinase subunit gamma-1 OS=Homo sapiens OX=9606 GN=PRKAG1 PE=1 SV=1 | 19 | 12 | 4 | 5.40E+05 | 3.46E+05 | 6.47E+05 | 8.88E+05 | 2.40E+05 | 3.20E+05 | | 4.89E+05 | 4.50E+05 | 3.98E+05 | 1.16E+05 | 8.25E+05 | 4.97E+05 | 4.56E+05 | 1.09 | 0.13 | 0.790746493 |
| O43324 | Eukaryotic translation elongation factor 1 epsilon-1 OS=Homo sapiens OX=9606 GN=EEF1E1 PE=1 SV=1 | 32 | 14 | 4 | 1.13E+07 | 1.26E+07 | 1.06E+07 | 1.11E+07 | 1.01E+07 | 9.03E+06 | 6.76E+06 | 7.27E+06 | 4.93E+06 | 5.77E+06 | 4.82E+06 | 5.37E+06 | 1.08E+07 | 5.82E+06 | 1.85 | 0.89 | 2.07018E-05 |
| Q9UQE7 | Structural maintenance of chromosomes protein 3 OS=Homo sapiens OX=9606 GN=SMC3 PE=1 SV=2 | 4 | 20 | 4 | 2.31E+06 | 7.16E+06 | 4.86E+06 | 4.67E+06 | 4.71E+06 | 4.53E+06 | 1.27E+06 | 7.65E+05 | 9.60E+05 | 9.11E+05 | 1.35E+06 | 2.26E+06 | 4.71E+06 | 1.25E+06 | 3.76 | 1.91 | 0.001810189 |
| P62910 | 60S ribosomal protein L32 OS=Homo sapiens OX=9606 GN=RPL32 PE=1 SV=2 | 27 | 61 | 4 | 2.65E+07 | 2.74E+07 | 2.62E+07 | 2.96E+07 | 2.63E+07 | 2.69E+07 | 1.82E+07 | 1.48E+07 | 1.37E+07 | 1.52E+07 | 1.61E+07 | 1.56E+07 | 2.71E+07 | 1.56E+07 | 1.74 | 0.80 | 7.65129E-08 |
| O75165 | DnaJ homolog subfamily C member 13 OS=Homo sapiens OX=9606 GN=DNAJC13 PE=1 SV=5 | 2 | 8 | 4 | 1.72E+06 | 1.26E+06 | 1.90E+06 | 1.67E+06 | 2.75E+06 | 2.14E+06 | 2.55E+06 | 1.58E+06 | 1.47E+06 | 1.04E+06 | 1.92E+06 | 1.74E+06 | 1.91E+06 | 1.72E+06 | 1.11 | 0.15 | 0.535117128 |
| P28070 | Proteasome subunit beta type-4 OS=Homo sapiens OX=9606 GN=PSMB4 PE=1 SV=4 | 18 | 48 | 4 | 1.84E+07 | 2.05E+07 | 2.05E+07 | 2.37E+07 | 2.22E+07 | 1.97E+07 | 2.12E+07 | 2.01E+07 | 2.10E+07 | 2.20E+07 | 2.02E+07 | 1.66E+07 | 2.08E+07 | 2.02E+07 | 1.03 | 0.05 | 0.550972287 |
| P25788 | Proteasome subunit alpha type-3 OS=Homo sapiens OX=9606 GN=PSMA3 PE=1 SV=2 | 23 | 52 | 4 | 1.75E+07 | 2.01E+07 | 1.82E+07 | 1.65E+07 | 1.78E+07 | 1.63E+07 | 1.78E+07 | 1.77E+07 | 1.50E+07 | 1.65E+07 | 1.40E+07 | 1.38E+07 | 1.77E+07 | 1.58E+07 | 1.12 | 0.17 | 0.058585997 |
| A0AVT1 | Ubiquitin-like modifier-activating enzyme 6 OS=Homo sapiens OX=9606 GN=UBA6 PE=1 SV=1 | 4 | 29 | 4 | 4.95E+06 | 5.60E+06 | 5.70E+06 | 5.16E+06 | 4.92E+06 | 5.57E+06 | 5.18E+06 | 3.93E+06 | 4.13E+06 | 1.80E+06 | 4.11E+06 | 3.56E+06 | 5.32E+06 | 3.79E+06 | 1.40 | 0.49 | 0.018250787 |
| P26373 | 60S ribosomal protein L13 OS=Homo sapiens OX=9606 GN=RPL13 PE=1 SV=4 | 19 | 66 | 4 | 2.49E+07 | 2.44E+07 | 2.45E+07 | 2.54E+07 | 2.25E+07 | 2.31E+07 | 1.72E+07 | 1.58E+07 | 1.30E+07 | 1.45E+07 | 1.36E+07 | 1.01E+07 | 2.41E+07 | 1.40E+07 | 1.72 | 0.78 | 3.62621E-05 |
| P25789 | Proteasome subunit alpha type-4 OS=Homo sapiens OX=9606 GN=PSMA4 PE=1 SV=1 | 16 | 50 | 4 | 1.80E+07 | 2.19E+07 | 1.84E+07 | 2.11E+07 | 2.09E+07 | 1.80E+07 | 1.94E+07 | 2.00E+07 | 1.94E+07 | 1.63E+07 | 1.90E+07 | 1.79E+07 | 1.97E+07 | 1.87E+07 | 1.06 | 0.08 | 0.286361005 |
| O60716 | Catenin delta-1 OS=Homo sapiens OX=9606 GN=CTNND1 PE=1 SV=1 | 7 | 7 | 4 | 6.04E+06 | 5.20E+06 | 5.02E+06 | 5.11E+06 | 4.87E+06 | 4.80E+06 | 5.08E+06 | 4.85E+06 | 6.14E+05 | 4.77E+06 | 4.10E+06 | 3.78E+06 | 5.17E+06 | 3.87E+06 | 1.34 | 0.42 | 0.11552467 |
| P35555 | Fibrillin-1 OS=Homo sapiens OX=9606 GN=FBN1 PE=1 SV=4 | 1 | 8 | 4 | | | | | | | 8.60E+05 | 2.40E+06 | 9.23E+05 | 6.22E+05 | 6.64E+05 | 3.14E+06 | | 1.43E+06 | | | |
| P51812 | Ribosomal protein S6 kinase alpha-3 OS=Homo sapiens OX=9606 GN=RPS6KA3 PE=1 SV=1 | 6 | 39 | 4 | 3.26E+06 | 4.99E+06 | 4.70E+06 | 5.95E+06 | 5.52E+06 | 4.91E+06 | 5.38E+06 | 5.91E+06 | 3.65E+06 | 3.56E+06 | 3.99E+06 | 5.82E+06 | 4.89E+06 | 4.72E+06 | 1.04 | 0.05 | 0.778021883 |
| P35659 | Protein DEK OS=Homo sapiens OX=9606 GN=DEK PE=1 SV=1 | 12 | 54 | 4 | 8.53E+06 | 7.44E+06 | 9.49E+06 | 1.05E+07 | 9.94E+06 | 9.08E+06 | 5.63E+06 | 7.31E+06 | 9.18E+06 | 8.01E+06 | 7.53E+06 | 5.41E+06 | 9.16E+06 | 7.18E+06 | 1.28 | 0.35 | 0.023811245 |
| Q15185 | Prostaglandin E synthase 3 OS=Homo sapiens OX=9606 GN=PTGES3 PE=1 SV=1 | 19 | 39 | 4 | 4.99E+06 | 3.90E+06 | 4.96E+06 | 4.76E+06 | 1.17E+07 | 1.38E+07 | 4.64E+06 | 3.64E+06 | 4.08E+06 | 9.07E+06 | 4.33E+06 | 7.46E+06 | 7.34E+06 | 5.54E+06 | 1.33 | 0.41 | 0.383040309 |
| Q99816 | Tumor susceptibility gene 101 protein OS=Homo sapiens OX=9606 GN=TSG101 PE=1 SV=2 | 12 | 26 | 4 | 5.48E+06 | 5.24E+06 | 4.16E+06 | 6.52E+06 | 2.52E+06 | 4.43E+06 | 6.29E+06 | 5.27E+06 | 7.75E+06 | 6.52E+06 | 7.26E+06 | 5.47E+06 | 4.73E+06 | 6.43E+06 | 0.74 | -0.44 | 0.034824239 |
| Q13616 | Cullin-1 OS=Homo sapiens OX=9606 GN=CUL1 PE=1 SV=2 | 5 | 19 | 4 | 3.37E+06 | 3.60E+06 | 1.93E+06 | 3.56E+06 | 2.88E+06 | 1.54E+06 | 3.81E+06 | 1.37E+06 | 4.52E+06 | 4.21E+06 | 4.86E+05 | | 2.81E+06 | 2.88E+06 | 0.98 | -0.03 | 0.944076077 |
| P05543 | Thyroxine-binding globulin OS=Homo sapiens OX=9606 GN=SERPINA7 PE=1 SV=2 | 9 | 166 | 4 | 1.71E+07 | 1.60E+07 | 1.45E+07 | 1.41E+07 | 2.19E+07 | 1.78E+07 | 1.99E+07 | 1.77E+07 | 1.69E+07 | 1.50E+07 | 1.57E+07 | 1.56E+07 | 1.69E+07 | 1.71E+07 | 0.99 | -0.02 | 0.876206932 |
| P48556 | 26S proteasome non-ATPase regulatory subunit 8 OS=Homo sapiens OX=9606 GN=PSMD8 PE=1 SV=2 | 9 | 39 | 4 | 9.68E+06 | 1.07E+07 | 9.22E+06 | 1.08E+07 | 7.02E+06 | 6.80E+06 | 9.04E+06 | 8.47E+06 | 8.72E+06 | 6.37E+06 | 9.01E+06 | 6.81E+06 | 8.95E+06 | 8.07E+06 | 1.11 | 0.15 | 0.317642821 |
| Q16777 | Histone H2A type 2-C OS=Homo sapiens OX=9606 GN=HIST2H2AC PE=1 SV=4 | 64 | 3161 | 5 | 1.04E+10 | 9.14E+09 | 9.08E+09 | 1.01E+10 | 9.55E+09 | 9.77E+09 | 1.15E+10 | 1.07E+10 | 9.90E+09 | 1.10E+10 | 9.78E+09 | 8.59E+09 | 9.67E+09 | 1.03E+10 | 0.94 | -0.08 | 0.265918412 |
| P02538 | Keratin, type II cytoskeletal 6A OS=Homo sapiens OX=9606 GN=KRT6A PE=1 SV=3 | 37 | 547 | 5 | 1.58E+08 | 7.58E+07 | 4.61E+07 | 4.63E+07 | 4.73E+07 | 9.47E+07 | 3.48E+07 | 1.45E+08 | 9.48E+07 | 8.55E+07 | 6.01E+07 | 3.77E+07 | 7.80E+07 | 7.63E+07 | 1.02 | 0.03 | 0.948056734 |
| Q71UI9 | Histone H2A.V OS=Homo sapiens OX=9606 GN=H2AFV PE=1 SV=3 | 54 | 1139 | 5 | 6.84E+08 | 6.50E+08 | 6.56E+08 | 6.61E+08 | 7.38E+08 | 5.87E+08 | 4.20E+08 | 4.02E+08 | 3.30E+08 | 3.61E+08 | 2.88E+08 | 2.90E+08 | 6.63E+08 | 3.48E+08 | 1.90 | 0.93 | 1.30272E-06 |
| P62140 | Serine/threonine-protein phosphatase PP1-beta catalytic subunit OS=Homo sapiens OX=9606 GN=PPP1CB PE=1 SV=3 | 44 | 194 | 5 | 1.66E+07 | 1.59E+07 | 1.66E+07 | 1.78E+07 | 1.71E+07 | 1.69E+07 | 1.45E+07 | 1.14E+07 | 1.29E+07 | 1.36E+07 | 1.30E+07 | 1.32E+07 | 1.68E+07 | 1.31E+07 | 1.28 | 0.36 | 4.79012E-05 |
| P23528 | Cofilin-1 OS=Homo sapiens OX=9606 GN=CFL1 PE=1 SV=3 | 59 | 182 | 5 | 6.83E+07 | 7.79E+07 | 6.22E+07 | 6.70E+07 | 5.91E+07 | 7.31E+07 | 7.53E+07 | 7.33E+07 | 4.97E+07 | 4.22E+07 | 5.67E+07 | 5.19E+07 | 6.79E+07 | 5.82E+07 | 1.17 | 0.22 | 0.153270067 |
| P36955 | Pigment epithelium-derived factor OS=Homo sapiens OX=9606 GN=SERPINF1 PE=1 SV=4 | 13 | 616 | 5 | 3.92E+08 | 4.05E+08 | 2.86E+08 | 3.18E+08 | 3.08E+08 | 3.17E+08 | 4.62E+08 | 4.93E+08 | 3.80E+08 | 3.71E+08 | 3.53E+08 | 3.54E+08 | 3.38E+08 | 4.02E+08 | 0.84 | -0.25 | 0.069712677 |
| Q10567 | AP-1 complex subunit beta-1 OS=Homo sapiens OX=9606 GN=AP1B1 PE=1 SV=2 | 14 | 194 | 5 | 1.70E+07 | 1.78E+07 | 1.51E+07 | 1.54E+07 | 1.49E+07 | 1.79E+07 | 1.35E+07 | 1.19E+07 | 1.11E+07 | 1.16E+07 | 1.13E+07 | 8.29E+06 | 1.64E+07 | 1.13E+07 | 1.45 | 0.54 | 0.000233445 |
| P31946 | 14-3-3 protein beta/alpha OS=Homo sapiens OX=9606 GN=YWHAB PE=1 SV=3 | 39 | 89 | 5 | 6.81E+06 | 7.73E+06 | 1.28E+07 | 1.64E+07 | 9.24E+06 | 8.02E+06 | 7.86E+06 | 9.06E+06 | 8.70E+06 | 7.75E+06 | 8.55E+06 | 6.05E+06 | 1.02E+07 | 8.00E+06 | 1.27 | 0.35 | 0.219056139 |
| P36507 | Dual specificity mitogen-activated protein kinase kinase 2 OS=Homo sapiens OX=9606 GN=MAP2K2 PE=1 SV=1 | 36 | 106 | 5 | 1.36E+07 | 1.43E+07 | 1.31E+07 | 1.49E+07 | 1.33E+07 | 1.39E+07 | 9.83E+06 | 9.03E+06 | 8.01E+06 | 7.59E+06 | 6.90E+06 | 8.62E+06 | 1.39E+07 | 8.33E+06 | 1.67 | 0.74 | 2.69903E-06 |
| P62258 | 14-3-3 protein epsilon OS=Homo sapiens OX=9606 GN=YWHAE PE=1 SV=1 | 25 | 83 | 5 | 1.60E+07 | 2.04E+07 | 2.12E+07 | 2.91E+07 | 2.71E+07 | 2.69E+07 | 1.40E+07 | 1.81E+07 | 1.98E+07 | 2.17E+07 | 1.76E+07 | 1.42E+07 | 2.35E+07 | 1.76E+07 | 1.34 | 0.42 | 0.039169243 |
| P61981 | 14-3-3 protein gamma OS=Homo sapiens OX=9606 GN=YWHAG PE=1 SV=2 | 33 | 99 | 5 | 8.54E+06 | 9.68E+06 | 1.14E+07 | 1.39E+07 | 1.14E+07 | 1.15E+07 | 8.15E+06 | 1.17E+07 | 9.38E+06 | 1.24E+07 | 1.05E+07 | 8.65E+06 | 1.10E+07 | 1.01E+07 | 1.09 | 0.13 | 0.382799466 |
| P34896 | Serine hydroxymethyltransferase, cytosolic OS=Homo sapiens OX=9606 GN=SHMT1 PE=1 SV=1 | 16 | 151 | 5 | 1.78E+07 | 1.83E+07 | 2.28E+07 | 2.12E+07 | 2.38E+07 | 1.74E+07 | 2.33E+07 | 1.57E+07 | 2.12E+07 | 2.29E+07 | 2.05E+07 | 1.80E+07 | 2.02E+07 | 2.02E+07 | 1.00 | 0.00 | 0.987347147 |
| P80723 | Brain acid soluble protein 1 OS=Homo sapiens OX=9606 GN=BASP1 PE=1 SV=2 | 48 | 82 | 5 | 1.48E+07 | 1.32E+07 | 2.12E+07 | 2.34E+07 | 1.59E+07 | 1.90E+07 | 2.00E+07 | 1.84E+07 | 2.32E+07 | 2.13E+07 | 1.45E+07 | 1.44E+07 | 1.79E+07 | 1.86E+07 | 0.96 | -0.06 | 0.754653108 |
| O14773 | Tripeptidyl-peptidase 1 OS=Homo sapiens OX=9606 GN=TPP1 PE=1 SV=2 | 15 | 104 | 5 | 3.14E+07 | 3.05E+07 | 3.17E+07 | 2.75E+07 | 2.89E+07 | 3.15E+07 | 2.19E+07 | 2.13E+07 | 2.09E+07 | 1.30E+07 | 2.14E+07 | 1.54E+07 | 3.02E+07 | 1.90E+07 | 1.59 | 0.67 | 0.000313032 |
| O00303 | Eukaryotic translation initiation factor 3 subunit F OS=Homo sapiens OX=9606 GN=EIF3F PE=1 SV=1 | 21 | 100 | 5 | 1.58E+07 | 2.47E+07 | 2.95E+07 | 3.44E+07 | 3.77E+07 | 2.31E+07 | 1.19E+07 | 1.52E+07 | 1.69E+07 | 1.30E+07 | 1.47E+07 | 1.38E+07 | 2.75E+07 | 1.42E+07 | 1.93 | 0.95 | 0.00888529 |
| P06732 | Creatine kinase M-type OS=Homo sapiens OX=9606 GN=CKM PE=1 SV=2 | 24 | 66 | 5 | 1.24E+07 | 1.40E+07 | 1.19E+07 | 1.45E+07 | 1.33E+07 | 1.46E+07 | 2.11E+07 | 2.11E+07 | 1.49E+07 | 1.32E+07 | 1.54E+07 | 1.75E+07 | 1.34E+07 | 1.72E+07 | 0.78 | -0.35 | 0.038577006 |
| P53985 | Monocarboxylate transporter 1 OS=Homo sapiens OX=9606 GN=SLC16A1 PE=1 SV=3 | 12 | 48 | 5 | 1.44E+07 | 1.43E+07 | 1.49E+07 | 1.97E+07 | 1.79E+07 | 1.59E+07 | 8.07E+06 | 6.27E+06 | 8.68E+06 | 7.09E+06 | 8.55E+06 | 5.91E+06 | 1.62E+07 | 7.43E+06 | 2.18 | 1.12 | 2.84234E-05 |
| P05161 | Ubiquitin-like protein ISG15 OS=Homo sapiens OX=9606 GN=ISG15 PE=1 SV=5 | 31 | 116 | 5 | 3.86E+07 | 4.26E+07 | 3.29E+07 | 3.77E+07 | 3.94E+07 | 3.35E+07 | 2.51E+07 | 2.44E+07 | 2.49E+07 | 2.64E+07 | 2.23E+07 | 1.70E+07 | 3.74E+07 | 2.33E+07 | 1.60 | 0.68 | 1.4367E-05 |
| P49411 | Elongation factor Tu, mitochondrial OS=Homo sapiens OX=9606 GN=TUFM PE=1 SV=2 | 16 | 48 | 5 | 2.07E+07 | 2.02E+07 | 2.26E+07 | 2.30E+07 | 1.83E+07 | 1.96E+07 | 1.24E+07 | 1.14E+07 | 8.12E+06 | 1.10E+07 | 1.02E+07 | 8.00E+06 | 2.07E+07 | 1.02E+07 | 2.03 | 1.02 | 1.42903E-06 |

### Supplementary Table 4. List of all proteins identified by label free proteomics analysis.

| Accession | Description | Coverage [%] | # PSMs | # Unique Peptides | Amitriptyline 1 | Amitriptyline 2 | Amitriptyline 3 | Amitriptyline 4 | Amitriptyline 5 | Amitriptyline 6 | Vehicle 1 | Vehicle 2 | Vehicle 3 | Vehicle 4 | Vehicle 5 | Vehicle 6 | Average amitriptyline | Average vehicle | Ratio amitriptyline/ vehicle | Log 2 fold change (amitriptyline /vehicle) | P-value |
|---|---|---|---|---|---|---|---|---|---|---|---|---|---|---|---|---|---|---|---|---|---|
| P01834 | Immunoglobulin kappa constant OS=Homo sapiens OX=9606 GN=IGKC PE=1 SV=2 | 79 | 15 | 5 | 1.02E+06 | | | | | | 4.28E+06 | 2.56E+05 | | | 3.65E+05 | 1.98E+07 | 1.02E+06 | 6.18E+06 | 0.16 | -2.60 | |
| Q9Y240 | C-type lectin domain family 11 member A OS=Homo sapiens OX=9606 GN=CLEC11A PE=1 SV=1 | 15 | 261 | 5 | 9.65E+07 | 8.48E+07 | 8.38E+07 | 8.63E+07 | 7.80E+07 | 7.39E+07 | 1.22E+08 | 1.10E+08 | 9.88E+07 | 1.02E+08 | 9.51E+07 | 8.28E+07 | 8.39E+07 | 1.02E+08 | 0.82 | -0.28 | 0.021382525 |
| Q02750 | Dual specificity mitogen-activated protein kinase kinase 1 OS=Homo sapiens OX=9606 GN=MAP2K1 PE=1 SV=2 | 34 | 66 | 5 | 1.61E+07 | 1.69E+07 | 1.90E+07 | 2.26E+07 | 1.75E+07 | 1.75E+07 | 1.37E+07 | 1.63E+07 | 1.32E+07 | 1.86E+07 | 1.39E+07 | 1.52E+07 | 1.83E+07 | 1.52E+07 | 1.20 | 0.27 | 0.034602911 |
| P13804 | Electron transfer flavoprotein subunit alpha, mitochondrial OS=Homo sapiens OX=9606 GN=ETFA PE=1 SV=1 | 27 | 42 | 5 | 7.63E+06 | 7.46E+06 | 1.04E+07 | 1.03E+07 | 7.46E+06 | 1.14E+07 | 6.06E+06 | 3.16E+06 | 5.17E+06 | 2.99E+06 | 3.78E+06 | 3.32E+06 | 9.11E+06 | 4.08E+06 | 2.23 | 1.16 | 0.000318344 |
| P30613 | Pyruvate kinase PKLR OS=Homo sapiens OX=9606 GN=PKLR PE=1 SV=2 | 13 | 196 | 5 | 1.42E+07 | 1.59E+07 | 1.16E+07 | 1.16E+07 | 1.07E+07 | 1.16E+07 | 1.78E+07 | 1.98E+07 | 1.49E+07 | 1.55E+07 | 1.35E+07 | 1.17E+07 | 1.26E+07 | 1.55E+07 | 0.81 | -0.30 | 0.073344756 |
| P52907 | F-actin-capping protein subunit alpha-1 OS=Homo sapiens OX=9606 GN=CAPZA1 PE=1 SV=3 | 43 | 96 | 5 | 2.30E+07 | 2.20E+07 | 1.84E+07 | 2.44E+07 | 1.80E+07 | 2.46E+07 | 1.82E+07 | 1.56E+07 | 1.55E+07 | 1.55E+07 | 1.59E+07 | 1.44E+07 | 2.17E+07 | 1.59E+07 | 1.37 | 0.45 | 0.002975255 |
| P16278 | Beta-galactosidase OS=Homo sapiens OX=9606 GN=GLB1 PE=1 SV=2 | 13 | 36 | 5 | 1.31E+07 | 1.24E+07 | 1.06E+07 | 8.56E+06 | 6.56E+06 | 6.15E+06 | 3.46E+06 | 3.49E+06 | 1.27E+06 | 3.84E+06 | 3.66E+06 | 2.16E+06 | 9.57E+06 | 2.98E+06 | 3.21 | 1.68 | 0.001890192 |
| P17844 | Probable ATP-dependent RNA helicase DDX5 OS=Homo sapiens OX=9606 GN=DDX5 PE=1 SV=1 | 20 | 105 | 5 | 9.90E+06 | 1.16E+07 | 1.05E+07 | 1.02E+07 | 1.08E+07 | 8.26E+06 | 5.33E+06 | 6.22E+06 | 4.79E+06 | 5.40E+06 | 4.52E+06 | 3.79E+06 | 1.02E+07 | 5.01E+06 | 2.04 | 1.03 | 6.16793E-06 |
| O00487 | 26S proteasome non-ATPase regulatory subunit 14 OS=Homo sapiens OX=9606 GN=PSMD14 PE=1 SV=1 | 36 | 63 | 5 | 1.92E+07 | 2.30E+07 | 2.31E+07 | 1.72E+07 | 1.77E+07 | 1.42E+07 | 1.16E+07 | 1.92E+07 | 9.12E+06 | 7.35E+06 | 8.91E+06 | 9.83E+06 | 1.91E+07 | 1.10E+07 | 1.74 | 0.80 | 0.004985815 |
| P06858 | Lipoprotein lipase OS=Homo sapiens OX=9606 GN=LPL PE=1 SV=1 | 19 | 52 | 5 | 1.08E+07 | 9.72E+06 | 6.17E+06 | 1.01E+07 | 1.10E+07 | 9.79E+06 | 1.57E+07 | 1.35E+07 | 1.41E+07 | 1.34E+07 | 1.18E+07 | 1.25E+07 | 9.60E+06 | 1.35E+07 | 0.71 | -0.49 | 0.001792388 |
| O43175 | D-3-phosphoglycerate dehydrogenase OS=Homo sapiens OX=9606 GN=PHGDH PE=1 SV=4 | 15 | 112 | 5 | 4.05E+07 | 4.23E+07 | 3.45E+07 | 4.36E+07 | 3.18E+07 | 3.62E+07 | 5.10E+07 | 5.20E+07 | 4.43E+07 | 4.18E+07 | 4.34E+07 | 3.86E+07 | 3.82E+07 | 4.52E+07 | 0.84 | -0.24 | 0.034862253 |
| Q9Y696 | Chloride intracellular channel protein 4 OS=Homo sapiens OX=9606 GN=CLIC4 PE=1 SV=4 | 35 | 92 | 5 | 2.15E+07 | 1.79E+07 | 1.91E+07 | 1.93E+07 | 1.95E+07 | 1.93E+07 | 2.08E+07 | 1.79E+07 | 1.64E+07 | 2.04E+07 | 1.45E+07 | 1.52E+07 | 1.94E+07 | 1.75E+07 | 1.11 | 0.15 | 0.14832385 |
| P30050 | 60S ribosomal protein L12 OS=Homo sapiens OX=9606 GN=RPL12 PE=1 SV=1 | 45 | 142 | 5 | 3.88E+07 | 4.15E+07 | 3.07E+07 | 2.72E+07 | 3.13E+07 | 3.34E+07 | 3.38E+07 | 4.05E+07 | 2.90E+07 | 3.08E+07 | 2.72E+07 | 2.65E+07 | 3.38E+07 | 3.13E+07 | 1.08 | 0.11 | 0.430112375 |
| P21926 | CD9 antigen OS=Homo sapiens OX=9606 GN=CD9 PE=1 SV=4 | 21 | 179 | 5 | 1.01E+08 | 9.41E+07 | 9.44E+07 | 1.09E+08 | 9.57E+07 | 9.34E+07 | 7.68E+07 | 7.32E+07 | 7.22E+07 | 6.42E+07 | 6.12E+07 | 6.42E+07 | 9.79E+07 | 6.86E+07 | 1.43 | 0.51 | 8.4871E-06 |
| Q9Y4K0 | Lysyl oxidase homolog 2 OS=Homo sapiens OX=9606 GN=LOXL2 PE=1 SV=1 | 11 | 40 | 5 | 5.42E+06 | 6.29E+06 | 5.30E+06 | 5.92E+06 | 5.19E+06 | 4.71E+06 | 1.15E+07 | 1.16E+07 | 1.16E+07 | 9.89E+06 | 9.13E+06 | 8.83E+06 | 5.47E+06 | 1.04E+07 | 0.52 | -0.93 | 6.99966E-05 |
| P10644 | cAMP-dependent protein kinase type I-alpha regulatory subunit OS=Homo sapiens OX=9606 GN=PRKAR1A PE=1 SV=1 | 22 | 82 | 5 | 1.23E+07 | 1.12E+07 | 9.75E+06 | 1.27E+07 | 1.40E+07 | 1.13E+07 | 1.28E+07 | 1.24E+07 | 7.55E+06 | 7.83E+06 | 1.24E+07 | 1.08E+07 | 1.19E+07 | 1.06E+07 | 1.12 | 0.16 | 0.297722065 |
| O14786 | Neuropilin-1 OS=Homo sapiens OX=9606 GN=NRP1 PE=1 SV=3 | 11 | 52 | 5 | 1.34E+07 | 1.46E+07 | 1.25E+07 | 1.41E+07 | 1.16E+07 | 1.04E+07 | 1.05E+07 | 9.38E+06 | 7.56E+06 | 9.75E+06 | 7.60E+06 | 7.86E+06 | 1.28E+07 | 8.78E+06 | 1.45 | 0.54 | 0.000812827 |
| P22234 | Multifunctional protein ADE2 OS=Homo sapiens OX=9606 GN=PAICS PE=1 SV=3 | 14 | 45 | 5 | 1.19E+07 | 1.36E+07 | 6.33E+06 | 8.07E+06 | 6.12E+06 | 1.26E+07 | 5.78E+06 | 6.46E+06 | 3.37E+06 | 3.42E+06 | 5.30E+06 | 5.72E+06 | 9.77E+06 | 5.01E+06 | 1.95 | 0.97 | 0.015273477 |
| O00151 | PDZ and LIM domain protein 1 OS=Homo sapiens OX=9606 GN=PDLIM1 PE=1 SV=4 | 21 | 24 | 5 | 1.27E+07 | 1.09E+07 | 7.60E+06 | 9.94E+06 | 1.03E+07 | 1.01E+07 | 6.99E+06 | 7.05E+06 | 4.52E+06 | 5.86E+06 | 5.37E+06 | 5.54E+06 | 1.03E+07 | 5.89E+06 | 1.74 | 0.80 | 0.000500038 |
| P20618 | Proteasome subunit beta type-1 OS=Homo sapiens OX=9606 GN=PSMB1 PE=1 SV=2 | 32 | 76 | 5 | 2.01E+07 | 2.03E+07 | 2.04E+07 | 2.10E+07 | 2.19E+07 | 2.31E+07 | 2.31E+07 | 2.23E+07 | 2.30E+07 | 2.37E+07 | 2.03E+07 | 1.79E+07 | 2.11E+07 | 2.17E+07 | 0.97 | -0.04 | 0.598565877 |
| P27361 | Mitogen-activated protein kinase 3 OS=Homo sapiens OX=9606 GN=MAPK3 PE=1 SV=4 | 25 | 76 | 5 | 2.81E+07 | 2.60E+07 | 1.81E+07 | 2.44E+07 | 2.19E+07 | 2.22E+07 | 1.88E+07 | 2.18E+07 | 1.80E+07 | 2.00E+07 | 1.77E+07 | 1.60E+07 | 2.34E+07 | 1.87E+07 | 1.25 | 0.33 | 0.020872159 |
| P11279 | Lysosome-associated membrane glycoprotein 1 OS=Homo sapiens OX=9606 GN=LAMP1 PE=1 SV=3 | 12 | 126 | 5 | 6.23E+07 | 5.77E+07 | 5.88E+07 | 5.94E+07 | 5.52E+07 | 5.30E+07 | 2.05E+07 | 3.26E+07 | 3.07E+07 | 1.82E+07 | 1.81E+07 | 2.63E+07 | 5.77E+07 | 2.44E+07 | 2.37 | 1.24 | 5.57772E-06 |
| P61019 | Ras-related protein Rab-2A OS=Homo sapiens OX=9606 GN=RAB2A PE=1 SV=1 | 32 | 80 | 5 | 1.40E+07 | 1.21E+07 | 1.52E+07 | 1.34E+07 | 1.47E+07 | 1.53E+07 | 1.27E+07 | 1.06E+07 | 1.12E+07 | 9.47E+06 | 1.09E+07 | 8.89E+06 | 1.41E+07 | 1.06E+07 | 1.32 | 0.41 | 0.000985454 |
| P52292 | Importin subunit alpha-1 OS=Homo sapiens OX=9606 GN=KPNA2 PE=1 SV=1 | 19 | 26 | 5 | 9.66E+06 | 6.44E+06 | 4.96E+06 | 6.12E+06 | 3.96E+06 | 7.21E+06 | 4.46E+06 | 2.14E+06 | 1.96E+06 | 8.19E+05 | 2.80E+06 | 2.43E+06 | 6.39E+06 | 2.44E+06 | 2.62 | 1.39 | 0.002783473 |
| Q6V0I7 | Protocadherin Fat 4 OS=Homo sapiens OX=9606 GN=FAT4 PE=1 SV=2 | 2 | 26 | 5 | 4.67E+06 | 3.73E+06 | 4.64E+06 | 3.10E+06 | 4.70E+06 | 3.50E+06 | 7.24E+06 | 7.46E+06 | 6.02E+06 | 5.83E+06 | 6.19E+06 | 7.14E+06 | 4.06E+06 | 6.65E+06 | 0.61 | -0.71 | 8.19768E-05 |
| Q15435 | Protein phosphatase 1 regulatory subunit 7 OS=Homo sapiens OX=9606 GN=PPP1R7 PE=1 SV=1 | 19 | 90 | 5 | 1.10E+07 | 1.48E+07 | 1.20E+07 | 8.35E+06 | 1.47E+07 | 1.27E+07 | 1.37E+07 | 8.95E+06 | 1.34E+07 | 1.67E+07 | 1.18E+07 | 1.20E+07 | 1.23E+07 | 1.28E+07 | 0.96 | -0.06 | 0.734981671 |
| P35268 | 60S ribosomal protein L22 OS=Homo sapiens OX=9606 GN=RPL22 PE=1 SV=2 | 49 | 64 | 5 | 5.15E+07 | 4.91E+07 | 4.93E+07 | 5.14E+07 | 4.93E+07 | 4.71E+07 | 2.43E+07 | 2.47E+07 | 1.91E+07 | 2.28E+07 | 2.20E+07 | 1.92E+07 | 4.96E+07 | 2.20E+07 | 2.25 | 1.17 | 3.8406E-09 |
| P60900 | Proteasome subunit alpha type-6 OS=Homo sapiens OX=9606 GN=PSMA6 PE=1 SV=1 | 25 | 115 | 5 | 4.05E+07 | 4.58E+07 | 4.04E+07 | 3.98E+07 | 4.14E+07 | 3.38E+07 | 4.53E+07 | 4.77E+07 | 4.93E+07 | 5.82E+07 | 4.23E+07 | 3.27E+07 | 4.03E+07 | 4.59E+07 | 0.88 | -0.19 | 0.179858842 |
| Q9UBQ7 | Glyoxylate reductase/hydroxypyruvate reductase OS=Homo sapiens OX=9606 GN=GRHPR PE=1 SV=1 | 26 | 66 | 5 | 8.73E+06 | 9.20E+06 | 9.21E+06 | 1.15E+07 | 8.29E+06 | 1.16E+07 | 1.00E+07 | 9.92E+06 | 9.63E+06 | 1.08E+07 | 1.09E+07 | 9.65E+06 | 9.76E+06 | 1.01E+07 | 0.96 | -0.06 | 0.562018192 |
| P61225 | Ras-related protein Rap-2b OS=Homo sapiens OX=9606 GN=RAP2B PE=1 SV=1 | 33 | 66 | 5 | 1.85E+07 | 1.83E+07 | 1.70E+07 | 1.88E+07 | 2.06E+07 | 2.30E+07 | 1.78E+07 | 1.66E+07 | 1.59E+07 | 1.81E+07 | 1.44E+07 | 1.61E+07 | 1.94E+07 | 1.65E+07 | 1.18 | 0.23 | 0.021366338 |
| Q09666 | Neuroblast differentiation-associated protein AHNAK OS=Homo sapiens OX=9606 GN=AHNAK PE=1 SV=2 | 3 | 27 | 5 | 3.84E+06 | 5.61E+06 | 8.92E+06 | 9.47E+06 | 6.91E+06 | 1.13E+07 | 7.25E+06 | 8.82E+06 | 8.62E+06 | 7.20E+06 | 8.20E+06 | 5.66E+06 | 7.68E+06 | 7.63E+06 | 1.01 | 0.01 | 0.969113912 |
| Q9BR76 | Coronin-1B OS=Homo sapiens OX=9606 GN=CORO1B PE=1 SV=1 | 14 | 66 | 5 | 2.03E+07 | 1.75E+07 | 1.75E+07 | 1.75E+07 | 1.79E+07 | 1.41E+07 | 1.97E+07 | 1.82E+07 | 1.83E+07 | 1.59E+07 | 1.78E+07 | 1.53E+07 | 1.75E+07 | 1.75E+07 | 1.00 | 0.00 | 0.970067923 |
| P47756 | F-actin-capping protein subunit beta OS=Homo sapiens OX=9606 GN=CAPZB PE=1 SV=4 | 26 | 50 | 5 | 1.51E+07 | 1.50E+07 | 1.34E+07 | 1.26E+07 | 1.19E+07 | 1.39E+07 | 9.57E+06 | 1.16E+07 | 1.03E+07 | 9.71E+06 | 7.71E+06 | 7.58E+06 | 1.36E+07 | 9.41E+06 | 1.45 | 0.53 | 0.000481615 |
| Q9BXS5 | AP-1 complex subunit mu-1 OS=Homo sapiens OX=9606 GN=AP1M1 PE=1 SV=3 | 18 | 61 | 5 | 1.28E+07 | 1.29E+07 | 1.50E+07 | 1.40E+07 | 1.33E+07 | 1.23E+07 | 7.88E+06 | 7.54E+06 | 5.41E+06 | 6.78E+06 | 5.27E+06 | 5.69E+06 | 1.34E+07 | 6.43E+06 | 2.08 | 1.06 | 5.59084E-07 |
| Q96IJ6 | Mannose-1-phosphate guanyltransferase alpha OS=Homo sapiens OX=9606 GN=GMPPA PE=1 SV=1 | 15 | 60 | 5 | 6.63E+06 | 7.10E+06 | 8.22E+06 | 8.71E+06 | 7.61E+06 | 6.13E+06 | 9.09E+06 | 8.77E+06 | 7.53E+06 | 8.97E+06 | 7.95E+06 | 6.29E+06 | 7.40E+06 | 8.10E+06 | 0.91 | -0.13 | 0.265446138 |
| P12004 | Proliferating cell nuclear antigen OS=Homo sapiens OX=9606 GN=PCNA PE=1 SV=1 | 25 | 40 | 5 | 6.63E+06 | 1.22E+07 | 1.52E+07 | 1.40E+07 | 1.50E+07 | 1.31E+07 | 5.89E+06 | 1.09E+07 | 6.54E+06 | 1.22E+07 | 9.93E+06 | 1.08E+07 | 1.27E+07 | 9.38E+06 | 1.35 | 0.44 | 0.077290312 |
| P07686 | Beta-hexosaminidase subunit beta OS=Homo sapiens OX=9606 GN=HEXB PE=1 SV=3 | 15 | 17 | 5 | 5.18E+06 | 3.34E+06 | 4.16E+05 | 2.34E+06 | 1.69E+06 | 1.77E+06 | 6.27E+05 | 1.76E+06 | 6.21E+05 | 1.22E+06 | 8.90E+05 | 1.85E+06 | 2.46E+06 | 1.16E+06 | 2.11 | 1.08 | 0.115498476 |
| Q96KP4 | Cytosolic non-specific dipeptidase OS=Homo sapiens OX=9606 GN=CNDP2 PE=1 SV=2 | 19 | 91 | 5 | 1.56E+07 | 2.93E+07 | 2.37E+07 | 1.67E+07 | 1.51E+07 | 1.82E+07 | 1.73E+07 | 2.43E+07 | 2.42E+07 | 1.88E+07 | 2.53E+07 | 1.68E+07 | 1.98E+07 | 2.11E+07 | 0.94 | -0.09 | 0.643617171 |
| P04062 | Lysosomal acid glucosylceramidase OS=Homo sapiens OX=9606 GN=GBA PE=1 SV=3 | 15 | 42 | 5 | 7.62E+06 | 6.78E+06 | 7.85E+06 | 5.88E+06 | 6.87E+06 | 6.11E+06 | 5.90E+06 | 5.59E+06 | 4.51E+06 | 4.12E+06 | 5.15E+06 | 4.50E+06 | 6.85E+06 | 4.96E+06 | 1.38 | 0.47 | 0.001357749 |
| P27797 | Calreticulin OS=Homo sapiens OX=9606 GN=CALR PE=1 SV=1 | 17 | 82 | 5 | 2.02E+07 | 2.07E+07 | 2.18E+07 | 1.84E+07 | 2.20E+07 | 1.90E+07 | 1.98E+07 | 2.07E+07 | 1.78E+07 | 1.58E+07 | 1.66E+07 | 1.69E+07 | 2.03E+07 | 1.79E+07 | 1.13 | 0.18 | 0.035794604 |
| P23368 | NAD-dependent malic enzyme, mitochondrial OS=Homo sapiens OX=9606 GN=ME2 PE=1 SV=1 | 18 | 25 | 5 | 6.11E+06 | 3.75E+06 | 3.51E+06 | 5.36E+06 | 4.53E+06 | 5.87E+06 | 5.98E+06 | 4.01E+06 | 4.29E+06 | 5.41E+06 | 3.07E+06 | 3.32E+06 | 4.85E+06 | 4.35E+06 | 1.12 | 0.16 | 0.453001888 |
| P36776 | Lon protease homolog, mitochondrial OS=Homo sapiens OX=9606 GN=LONP1 PE=1 SV=2 | 8 | 17 | 5 | 2.53E+06 | 2.53E+06 | 1.78E+06 | 2.27E+06 | 2.17E+06 | 2.32E+06 | 2.19E+06 | 1.85E+06 | 1.83E+06 | 1.68E+06 | 1.66E+06 | 1.42E+06 | 2.27E+06 | 1.77E+06 | 1.28 | 0.36 | 0.009350397 |
| P02792 | Ferritin light chain OS=Homo sapiens OX=9606 GN=FTL PE=1 SV=2 | 40 | 53 | 5 | 1.13E+07 | 1.90E+07 | 1.28E+07 | 1.49E+07 | 1.22E+07 | 1.19E+07 | 2.15E+07 | 1.72E+07 | 2.19E+07 | 3.44E+07 | 2.06E+07 | 1.32E+07 | 1.37E+07 | 2.15E+07 | 0.64 | -0.65 | 0.043533764 |
| Q5VYK3 | Proteasome adapter and scaffold protein ECM29 OS=Homo sapiens OX=9606 GN=ECPAS PE=1 SV=2 | 5 | 25 | 5 | 6.62E+06 | 4.18E+06 | 4.80E+06 | 4.68E+06 | 4.78E+06 | 5.28E+06 | 3.96E+06 | 2.77E+06 | 2.68E+06 | 2.96E+06 | 3.16E+06 | 2.99E+06 | 5.06E+06 | 3.09E+06 | 1.64 | 0.71 | 0.001122316 |
| O75351 | Vacuolar protein sorting-associated protein 4B OS=Homo sapiens OX=9606 GN=VPS4B PE=1 SV=2 | 20 | 47 | 5 | 1.08E+07 | 8.59E+06 | 8.40E+06 | 6.57E+06 | 7.51E+06 | 1.23E+07 | 6.46E+06 | 7.44E+06 | 6.18E+06 | 3.77E+06 | 5.48E+06 | 4.84E+06 | 9.03E+06 | 5.70E+06 | 1.58 | 0.66 | 0.010976714 |
| P13861 | cAMP-dependent protein kinase type II-alpha regulatory subunit OS=Homo sapiens OX=9606 GN=PRKAR2A PE=1 SV=2 | 15 | 85 | 5 | 1.74E+07 | 1.89E+07 | 1.75E+07 | 1.86E+07 | 1.74E+07 | 1.78E+07 | 2.09E+07 | 2.29E+07 | 2.00E+07 | 2.09E+07 | 1.69E+07 | 1.56E+07 | 1.79E+07 | 1.95E+07 | 0.92 | -0.12 | 0.21276132 |
| P43243 | Matrin-3 OS=Homo sapiens OX=9606 GN=MATR3 PE=1 SV=2 | 12 | 30 | 5 | 9.61E+06 | 7.61E+06 | 8.51E+06 | 1.00E+07 | 6.20E+06 | 7.35E+06 | 2.46E+06 | 2.57E+06 | 2.07E+06 | 2.57E+06 | 3.04E+06 | 2.14E+06 | 8.22E+06 | 2.47E+06 | 3.32 | 1.73 | 0.000121366 |
| O60506 | Heterogeneous nuclear ribonucleoprotein Q OS=Homo sapiens OX=9606 GN=SYNCRIP PE=1 SV=2 | 11 | 56 | 5 | 1.62E+07 | 1.76E+07 | 1.73E+07 | 1.58E+07 | 1.71E+07 | 1.56E+07 | 1.30E+07 | 1.05E+07 | 9.27E+06 | 9.31E+06 | 1.01E+07 | 7.55E+06 | 1.66E+07 | 9.96E+06 | 1.67 | 0.74 | 7.59348E-05 |
| P02771 | Alpha-fetoprotein OS=Homo sapiens OX=9606 GN=AFP PE=1 SV=1 | 11 | 144 | 5 | 3.98E+07 | 3.97E+07 | 4.10E+07 | 4.88E+07 | 4.16E+07 | 3.78E+07 | 6.95E+07 | 6.76E+07 | 6.59E+07 | 6.52E+07 | 6.56E+07 | 5.71E+07 | 4.15E+07 | 6.52E+07 | 0.64 | -0.65 | 1.56342E-06 |
| P33991 | DNA replication licensing factor MCM4 OS=Homo sapiens OX=9606 GN=MCM4 PE=1 SV=5 | 11 | 35 | 5 | 6.63E+06 | 8.08E+06 | 6.24E+06 | 5.87E+06 | 5.05E+06 | 8.19E+06 | 3.68E+06 | 5.04E+06 | 2.43E+06 | 2.28E+06 | 3.84E+06 | 2.99E+06 | 6.68E+06 | 3.38E+06 | 1.98 | 0.98 | 0.000596737 |
| Q9UQ80 | Proliferation-associated protein 2G4 OS=Homo sapiens OX=9606 GN=PA2G4 PE=1 SV=3 | 17 | 47 | 5 | 1.01E+07 | 1.24E+07 | 1.20E+07 | 1.23E+07 | 1.27E+07 | 1.18E+07 | 1.03E+07 | 1.03E+07 | 1.32E+07 | 1.24E+07 | 9.78E+06 | 9.46E+06 | 1.19E+07 | 1.09E+07 | 1.09 | 0.12 | 0.226877842 |
| Q15365 | Poly(rC)-binding protein 1 OS=Homo sapiens OX=9606 GN=PCBP1 PE=1 SV=2 | 22 | 110 | 5 | 3.54E+07 | 3.38E+07 | 2.77E+07 | 3.10E+07 | 3.06E+07 | 3.01E+07 | 2.63E+07 | 2.65E+07 | 2.27E+07 | 2.24E+07 | 2.06E+07 | 1.97E+07 | 3.15E+07 | 2.30E+07 | 1.37 | 0.45 | 0.000383193 |
| Q9BTW9 | Tubulin-specific chaperone D OS=Homo sapiens OX=9606 GN=TBCD PE=1 SV=2 | 8 | 32 | 5 | 6.15E+06 | 6.30E+06 | 6.13E+06 | 6.49E+06 | 6.58E+06 | 5.51E+06 | 3.11E+06 | 3.83E+06 | 4.12E+06 | 2.06E+06 | 4.03E+06 | 2.25E+06 | 6.19E+06 | 3.23E+06 | 1.92 | 0.94 | 0.000191785 |
| P25205 | DNA replication licensing factor MCM3 OS=Homo sapiens OX=9606 GN=MCM3 PE=1 SV=3 | 10 | 44 | 5 | 1.04E+07 | 8.82E+06 | 8.87E+06 | 8.62E+06 | 8.63E+06 | 8.66E+06 | 6.62E+06 | 6.13E+06 | 5.33E+06 | 4.92E+06 | 5.51E+06 | 5.11E+06 | 9.01E+06 | 5.60E+06 | 1.61 | 0.69 | 6.13952E-06 |
| Q9Y6M1 | Insulin-like growth factor 2 mRNA-binding protein 2 OS=Homo sapiens OX=9606 GN=IGF2BP2 PE=1 SV=2 | 11 | 24 | 5 | 3.93E+06 | 4.20E+06 | 3.68E+06 | 3.80E+06 | 3.99E+06 | 3.32E+06 | 1.71E+06 | 1.90E+06 | 1.87E+06 | 1.78E+06 | 1.16E+06 | 9.68E+05 | 3.82E+06 | 1.57E+06 | 2.44 | 1.29 | 1.20321E-06 |
| P62316 | Small nuclear ribonucleoprotein Sm D2 OS=Homo sapiens OX=9606 GN=SNRPD2 PE=1 SV=1 | 47 | 68 | 5 | 3.45E+07 | 2.64E+07 | 2.82E+07 | 2.77E+07 | 2.63E+07 | 3.42E+07 | 2.25E+07 | 2.42E+07 | 2.04E+07 | 7.11E+06 | 1.86E+07 | 2.12E+07 | 2.96E+07 | 1.90E+07 | 1.56 | 0.64 | 0.006621797 |
| P27816 | Microtubule-associated protein 4 OS=Homo sapiens OX=9606 GN=MAP4 PE=1 SV=3 | 6 | 56 | 5 | 1.39E+07 | 1.52E+07 | 1.48E+07 | 1.61E+07 | 1.45E+07 | 1.31E+07 | 6.64E+06 | 6.15E+06 | 6.32E+06 | 5.74E+06 | 6.10E+06 | 4.06E+06 | 1.46E+07 | 5.83E+06 | 2.50 | 1.32 | 2.52748E-08 |
| P34932 | Heat shock 70 kDa protein 4 OS=Homo sapiens OX=9606 GN=HSPA4 PE=1 SV=4 | 8 | 82 | 5 | 1.57E+07 | 1.51E+07 | 1.52E+07 | 1.55E+07 | 1.65E+07 | 1.76E+07 | 1.94E+07 | 1.84E+07 | 1.64E+07 | 1.84E+07 | 1.40E+07 | 1.48E+07 | 1.59E+07 | 1.69E+07 | 0.94 | -0.09 | 0.345171426 |
| Q00341 | Vigilin OS=Homo sapiens OX=9606 GN=HDLBP PE=1 SV=2 | 6 | 40 | 5 | 9.11E+06 | 1.08E+07 | 7.95E+06 | 8.51E+06 | 7.19E+06 | 8.08E+06 | 5.16E+06 | 4.72E+06 | 4.06E+06 | 3.95E+06 | 5.15E+06 | 4.62E+06 | 8.60E+06 | 4.61E+06 | 1.87 | 0.90 | 0.000201907 |
| Q9Y3F4 | Serine-threonine kinase receptor-associated protein OS=Homo sapiens OX=9606 GN=STRAP PE=1 SV=1 | 24 | 34 | 5 | 1.05E+07 | 1.18E+07 | 1.01E+07 | 1.12E+07 | 1.19E+07 | 1.16E+07 | 1.14E+07 | 9.83E+06 | 1.05E+07 | 9.85E+06 | 6.47E+06 | 7.24E+06 | 1.12E+07 | 9.21E+06 | 1.21 | 0.28 | 0.054658688 |
| P54709 | Sodium/potassium-transporting ATPase subunit beta-3 OS=Homo sapiens OX=9606 GN=ATP1B3 PE=1 SV=1 | 21 | 62 | 5 | 2.02E+07 | 2.12E+07 | 1.59E+07 | 1.94E+07 | 1.76E+07 | 1.36E+07 | 1.44E+07 | 1.43E+07 | 1.33E+07 | 1.45E+07 | 1.09E+07 | 1.13E+07 | 1.80E+07 | 1.31E+07 | 1.37 | 0.45 | 0.006781486 |
| Q99497 | Protein/nucleic acid deglycase DJ-1 OS=Homo sapiens OX=9606 GN=PARK7 PE=1 SV=2 | 39 | 53 | 5 | 9.49E+06 | 8.05E+06 | 1.21E+07 | 1.06E+07 | 1.15E+07 | 1.01E+07 | 9.31E+06 | 7.26E+06 | 8.94E+06 | 8.18E+06 | 7.39E+06 | 7.27E+06 | 1.03E+07 | 8.06E+06 | 1.28 | 0.35 | 0.011315917 |
| Q14166 | Tubulin--tyrosine ligase-like protein 12 OS=Homo sapiens OX=9606 GN=TTLL12 PE=1 SV=2 | 11 | 34 | 5 | 8.17E+06 | 6.43E+06 | 5.84E+06 | 8.56E+06 | 5.17E+06 | 6.21E+06 | 6.63E+06 | 5.30E+06 | 6.24E+06 | 7.87E+06 | 6.99E+06 | 4.26E+06 | 6.73E+06 | 6.21E+06 | 1.08 | 0.11 | 0.512186462 |
| O15118 | NPC intracellular cholesterol transporter 1 OS=Homo sapiens OX=9606 GN=NPC1 PE=1 SV=2 | 4 | 42 | 5 | 1.56E+07 | 1.32E+07 | 1.37E+07 | 1.60E+07 | 1.51E+07 | 1.21E+07 | 1.32E+07 | 7.41E+06 | 1.02E+07 | 1.16E+07 | 1.11E+07 | 7.45E+06 | 1.43E+07 | 1.02E+07 | 1.40 | 0.49 | 0.006053696 |
| P20073 | Annexin A7 OS=Homo sapiens OX=9606 GN=ANXA7 PE=1 SV=3 | 13 | 108 | 5 | 1.99E+07 | 1.87E+07 | 1.36E+07 | 1.49E+07 | 1.36E+07 | 1.49E+07 | 1.67E+07 | 1.79E+07 | 1.25E+07 | 1.22E+07 | 1.20E+07 | 1.28E+07 | 1.59E+07 | 1.40E+07 | 1.14 | 0.18 | 0.238708273 |
| Q9BY44 | Eukaryotic translation initiation factor 2A OS=Homo sapiens OX=9606 GN=EIF2A PE=1 SV=3 | 11 | 26 | 5 | 8.84E+06 | 3.67E+06 | 6.74E+06 | 7.36E+06 | 7.52E+06 | 6.59E+06 | 2.47E+06 | 2.77E+06 | 3.13E+06 | 2.33E+06 | 3.50E+06 | 3.21E+06 | 6.79E+06 | 2.90E+06 | 2.34 | 1.23 | 0.002071084 |
| P61254 | 60S ribosomal protein L26 OS=Homo sapiens OX=9606 GN=RPL26 PE=1 SV=1 | 32 | 91 | 5 | 2.08E+07 | 1.85E+07 | 2.54E+07 | 2.60E+07 | 2.09E+07 | 1.88E+07 | 1.58E+07 | 1.49E+07 | 1.75E+07 | 2.33E+07 | 1.14E+07 | 1.32E+07 | 2.17E+07 | 1.60E+07 | 1.35 | 0.44 | 0.025386429 |
| Q13228 | Methanethiol oxidase OS=Homo sapiens OX=9606 GN=SELENBP1 PE=1 SV=2 | 16 | 39 | 5 | 6.16E+06 | 6.36E+06 | 5.34E+06 | 3.99E+06 | 4.85E+06 | 6.32E+06 | 9.24E+06 | 7.76E+06 | 6.86E+06 | 4.56E+06 | 7.35E+06 | 8.62E+06 | 5.50E+06 | 7.40E+06 | 0.74 | -0.43 | 0.039496506 |
| Q9Y617 | Phosphoserine aminotransferase OS=Homo sapiens OX=9606 GN=PSAT1 PE=1 SV=2 | 16 | 76 | 5 | 1.51E+07 | 1.64E+07 | 1.50E+07 | 1.42E+07 | 1.20E+07 | 1.34E+07 | 1.83E+07 | 1.80E+07 | 1.71E+07 | 1.42E+07 | 1.79E+07 | 1.26E+07 | 1.43E+07 | 1.63E+07 | 0.88 | -0.19 | 0.117640796 |
| Q15181 | Inorganic pyrophosphatase OS=Homo sapiens OX=9606 GN=PPA1 PE=1 SV=2 | 26 | 32 | 5 | 1.06E+07 | 8.62E+06 | 8.62E+06 | 9.53E+06 | 8.18E+06 | 1.18E+07 | 8.94E+06 | 1.33E+07 | 1.19E+07 | 6.12E+06 | 5.81E+06 | 5.69E+06 | 9.55E+06 | 8.64E+06 | 1.11 | 0.15 | 0.557071521 |
| Q9UBQ0 | Vacuolar protein sorting-associated protein 29 OS=Homo sapiens OX=9606 GN=VPS29 PE=1 SV=1 | 34 | 59 | 5 | 1.73E+07 | 1.75E+07 | 1.73E+07 | 1.87E+07 | 1.90E+07 | 1.72E+07 | 1.56E+07 | 1.59E+07 | 1.57E+07 | 1.67E+07 | 1.55E+07 | 1.57E+07 | 1.78E+07 | 1.58E+07 | 1.13 | 0.17 | 0.000669906 |
| P04181 | Ornithine aminotransferase, mitochondrial OS=Homo sapiens OX=9606 GN=OAT PE=1 SV=1 | 15 | 29 | 5 | 7.33E+06 | 7.43E+06 | 6.50E+06 | 6.20E+06 | 6.96E+06 | 7.40E+06 | 4.81E+06 | 4.63E+06 | 4.32E+06 | 4.15E+06 | 3.46E+06 | 3.33E+06 | 6.97E+06 | 4.12E+06 | 1.69 | 0.76 | 5.95811E-06 |
| P40429 | 60S ribosomal protein L13a OS=Homo sapiens OX=9606 GN=RPL13A PE=1 SV=2 | 24 | 70 | 5 | 3.95E+07 | 4.99E+07 | 4.55E+07 | 4.79E+07 | 4.38E+07 | 4.67E+07 | 2.89E+07 | 2.84E+07 | 2.82E+07 | 3.15E+07 | 2.93E+07 | 2.79E+07 | 4.56E+07 | 2.90E+07 | 1.57 | 0.65 | 3.23595E-05 |
| P07339 | Cathepsin D OS=Homo sapiens OX=9606 GN=CTSD PE=1 SV=1 | 15 | 86 | 5 | 2.61E+07 | 2.50E+07 | 2.32E+07 | 2.19E+07 | 3.05E+07 | 2.33E+07 | 1.36E+07 | 1.40E+07 | 1.08E+07 | 1.13E+07 | 1.17E+07 | 1.01E+07 | 2.50E+07 | 1.19E+07 | 2.10 | 1.07 | 2.45181E-05 |
| Q8TD19 | Serine/threonine-protein kinase Nek9 OS=Homo sapiens OX=9606 GN=NEK9 PE=1 SV=2 | 6 | 45 | 5 | 3.38E+06 | 4.54E+06 | 3.16E+06 | 3.07E+06 | 3.42E+06 | 3.07E+06 | 3.09E+06 | 3.23E+06 | 2.07E+06 | 1.72E+06 | 2.75E+06 | 3.07E+06 | 3.44E+06 | 2.66E+06 | 1.30 | 0.37 | 0.044037434 |
| P46087 | Probable 28S rRNA (cytosine(4447)-C(5))-methyltransferase OS=Homo sapiens OX=9606 GN=NOP2 PE=1 SV=2 | 9 | 14 | 5 | 7.31E+06 | 7.03E+06 | 5.33E+06 | 5.02E+06 | 3.93E+06 | 7.82E+06 | 7.97E+06 | 7.34E+06 | 3.93E+06 | 3.44E+06 | 4.11E+06 | 6.41E+06 | 6.07E+06 | 5.54E+06 | 1.10 | 0.13 | 0.607165816 |
| P04040 | Catalase OS=Homo sapiens OX=9606 GN=CAT PE=1 SV=3 | 14 | 18 | 5 | 5.16E+06 | 4.34E+06 | 4.65E+06 | 4.93E+06 | 4.55E+06 | 5.64E+06 | 6.97E+06 | 6.87E+06 | 1.00E+07 | 6.22E+06 | 7.38E+06 | 7.33E+06 | 4.88E+06 | 7.47E+06 | 0.65 | -0.61 | 0.003562322 |
| Q9NTJ3 | Structural maintenance of chromosomes protein 4 OS=Homo sapiens OX=9606 GN=SMC4 PE=1 SV=2 | 4 | 14 | 5 | 4.84E+05 | 1.32E+06 | 3.60E+06 | 1.79E+06 | 1.07E+06 | 1.57E+06 | | | 7.95E+05 | 7.29E+05 | 7.80E+05 | 6.66E+05 | 1.64E+06 | 7.43E+05 | 2.21 | 1.14 | 0.093052534 |
| Q04837 | Single-stranded DNA-binding protein, mitochondrial OS=Homo sapiens OX=9606 GN=SSBP1 PE=1 SV=1 | 36 | 93 | 5 | 3.42E+07 | 3.68E+07 | 3.23E+07 | 3.45E+07 | 2.97E+07 | 3.24E+07 | 2.38E+07 | 2.66E+07 | 2.26E+07 | 2.28E+07 | 2.17E+07 | 2.06E+07 | 3.33E+07 | 2.30E+07 | 1.45 | 0.54 | 1.34645E-05 |
| P05106 | Integrin beta-3 OS=Homo sapiens OX=9606 GN=ITGB3 PE=1 SV=2 | 7 | 57 | 5 | 2.14E+07 | 1.09E+07 | 9.50E+06 | 1.16E+07 | 8.20E+06 | 2.04E+07 | 2.88E+07 | 1.35E+07 | 1.25E+07 | 1.17E+07 | 1.21E+07 | 2.48E+07 | 1.37E+07 | 1.72E+07 | 0.79 | -0.33 | 0.381975866 |
| P00749 | Urokinase-type plasminogen activator OS=Homo sapiens OX=9606 GN=PLAU PE=1 SV=2 | 15 | 14 | 5 | 8.93E+06 | 8.22E+06 | 6.44E+06 | 7.28E+06 | 7.20E+06 | 7.57E+06 | 1.02E+07 | 1.11E+07 | 8.64E+06 | 9.66E+06 | 9.51E+06 | 8.39E+06 | 7.61E+06 | 9.59E+06 | 0.79 | -0.33 | 0.004750868 |
| P62829 | 60S ribosomal protein L23 OS=Homo sapiens OX=9606 GN=RPL23 PE=1 SV=1 | 39 | 60 | 5 | 2.23E+07 | 2.44E+07 | 2.13E+07 | 2.18E+07 | 1.96E+07 | 2.11E+07 | 1.50E+07 | 1.50E+07 | 1.24E+07 | 1.23E+07 | 8.00E+06 | 1.01E+07 | 2.18E+07 | 1.21E+07 | 1.79 | 0.84 | 7.8162E-05 |
| Q86X55 | Histone-arginine methyltransferase CARM1 OS=Homo sapiens OX=9606 GN=CARM1 PE=1 SV=3 | 11 | 147 | 5 | 3.95E+08 | 2.49E+08 | 3.85E+08 | 2.89E+08 | 2.64E+08 | 2.73E+08 | 4.10E+08 | 3.37E+08 | 2.33E+08 | 2.26E+08 | 2.19E+08 | 1.87E+08 | 3.09E+08 | 2.69E+08 | 1.15 | 0.20 | 0.381481246 |
| O76021 | Ribosomal L1 domain-containing protein 1 OS=Homo sapiens OX=9606 GN=RSL1D1 PE=1 SV=3 | 13 | 44 | 5 | 6.59E+06 | 6.39E+06 | 6.31E+06 | 6.33E+06 | 7.31E+06 | 1.55E+07 | 5.28E+06 | 4.69E+06 | 4.02E+06 | 4.83E+06 | 5.69E+06 | 5.76E+06 | 8.08E+06 | 5.04E+06 | 1.60 | 0.68 | 0.099416429 |
| P78417 | Glutathione S-transferase omega-1 OS=Homo sapiens OX=9606 GN=GSTO1 PE=1 SV=2 | 21 | 54 | 5 | 1.50E+07 | 1.73E+07 | 1.59E+07 | 1.69E+07 | 1.59E+07 | 1.54E+07 | 1.53E+07 | 1.63E+07 | 1.44E+07 | 1.36E+07 | 1.30E+07 | 1.29E+07 | 1.61E+07 | 1.42E+07 | 1.13 | 0.17 | 0.022482459 |
| Q6ZNJ1 | Neurobeachin-like protein 2 OS=Homo sapiens OX=9606 GN=NBEAL2 PE=1 SV=2 | 3 | 9 | 5 | 2.84E+06 | 2.33E+06 | 8.23E+06 | 9.92E+06 | 1.06E+07 | 2.30E+06 | 4.91E+06 | 1.72E+07 | 1.07E+07 | 8.89E+06 | 1.05E+07 | 8.69E+06 | 6.04E+06 | 1.01E+07 | 0.60 | -0.75 | 0.106168347 |
| O94874 | E3 UFM1-protein ligase 1 OS=Homo sapiens OX=9606 GN=UFL1 PE=1 SV=2 | 8 | 27 | 5 | 3.37E+06 | 4.62E+06 | 3.90E+06 | 4.11E+06 | 3.47E+06 | 3.84E+06 | 1.30E+06 | 1.98E+06 | 1.95E+06 | 2.84E+06 | 1.74E+06 | 1.26E+06 | 3.88E+06 | 1.84E+06 | 2.11 | 1.08 | 6.06823E-05 |
| Q14683 | Structural maintenance of chromosomes protein 1A OS=Homo sapiens OX=9606 GN=SMC1A PE=1 SV=2 | 6 | 23 | 5 | 1.81E+08 | 1.64E+08 | 6.37E+07 | 6.73E+07 | 7.03E+07 | 8.31E+07 | 9.49E+07 | 1.13E+08 | 3.29E+07 | 8.94E+07 | 6.14E+07 | 7.39E+07 | 1.05E+08 | 7.75E+07 | 1.35 | 0.44 | 0.297287302 |
| Q9Y2J2 | Band 4.1-like protein 3 OS=Homo sapiens OX=9606 GN=EPB41L3 PE=1 SV=2 | 5 | 19 | 5 | 1.56E+07 | 1.61E+07 | 1.48E+07 | 1.76E+07 | 2.25E+07 | 2.24E+07 | 2.09E+07 | 2.01E+07 | 2.13E+07 | 1.87E+07 | 2.16E+07 | 2.36E+07 | 1.82E+07 | 2.10E+07 | 0.86 | -0.21 | 0.109082343 |
| P07437 | Tubulin beta chain OS=Homo sapiens OX=9606 GN=TUBB PE=1 SV=2 | 81 | 5407 | 6 | 3.48E+08 | 3.53E+08 | 3.01E+08 | 2.74E+08 | 2.69E+08 | 3.84E+08 | 2.53E+08 | 2.71E+08 | 2.04E+08 | 2.24E+08 | 2.43E+08 | 2.67E+08 | 3.21E+08 | 2.44E+08 | 1.32 | 0.40 | 0.007631842 |
| P68032 | Actin, alpha cardiac muscle 1 OS=Homo sapiens OX=9606 GN=ACTC1 PE=1 SV=1 | 58 | 2921 | 6 | 9.37E+07 | 8.82E+07 | 2.05E+08 | 1.16E+08 | 1.56E+08 | 9.49E+07 | 3.79E+08 | 1.01E+08 | 1.36E+08 | 1.03E+08 | 1.26E+08 | 1.04E+08 | 1.26E+08 | 1.58E+08 | 0.79 | -0.33 | 0.52182273 |
| Q9NZN3 | EH domain-containing protein 3 OS=Homo sapiens OX=9606 GN=EHD3 PE=1 SV=2 | 42 | 307 | 6 | 1.79E+07 | 1.54E+07 | 1.23E+07 | 1.57E+07 | 1.55E+07 | 1.67E+07 | 2.47E+07 | 2.46E+07 | 1.68E+07 | 1.84E+07 | 1.92E+07 | 1.81E+07 | 1.56E+07 | 2.03E+07 | 0.77 | -0.38 | 0.019071455 |
| P11217 | Glycogen phosphorylase, muscle form OS=Homo sapiens OX=9606 GN=PYGM PE=1 SV=6 | 27 | 374 | 6 | 8.94E+06 | 1.01E+07 | 7.51E+06 | 7.45E+06 | 8.65E+06 | 7.27E+06 | 9.64E+06 | 1.08E+07 | 1.09E+07 | 1.05E+07 | 9.59E+06 | 1.06E+07 | 8.32E+06 | 1.04E+07 | 0.80 | -0.32 | 0.004743457 |
| Q9H4B7 | Tubulin beta-1 chain OS=Homo sapiens OX=9606 GN=TUBB1 PE=1 SV=1 | 25 | 1622 | 6 | 6.58E+07 | 7.48E+07 | 7.20E+07 | 5.71E+07 | 5.93E+07 | 8.79E+07 | 6.44E+07 | 1.08E+08 | 8.59E+07 | 8.66E+07 | 9.03E+07 | 8.90E+07 | 6.95E+07 | 8.74E+07 | 0.79 | -0.33 | 0.035736857 |
| P52597 | Heterogeneous nuclear ribonucleoprotein F OS=Homo sapiens OX=9606 GN=HNRNPF PE=1 SV=3 | 28 | 134 | 6 | 4.94E+07 | 4.04E+07 | 4.15E+07 | 4.41E+07 | 4.16E+07 | 4.26E+07 | 3.03E+07 | 2.67E+07 | 2.25E+07 | 2.59E+07 | 2.31E+07 | 2.20E+07 | 4.33E+07 | 2.51E+07 | 1.72 | 0.78 | 2.02455E-06 |
| P17096 | High mobility group protein HMG-I/HMG-Y OS=Homo sapiens OX=9606 GN=HMGA1 PE=1 SV=3 | 41 | 293 | 6 | 2.09E+08 | 2.12E+08 | 1.89E+08 | 2.16E+08 | 1.77E+08 | 2.00E+08 | 1.43E+08 | 1.32E+08 | 1.21E+08 | 1.42E+08 | 1.08E+08 | 1.09E+08 | 2.00E+08 | 1.26E+08 | 1.59 | 0.67 | 7.659E-06 |
| P46782 | 40S ribosomal protein S5 OS=Homo sapiens OX=9606 GN=RPS5 PE=1 SV=4 | 43 | 85 | 6 | 2.11E+07 | 1.79E+07 | 1.80E+07 | 1.60E+07 | 1.94E+07 | 2.24E+07 | 1.71E+07 | 2.08E+07 | 1.37E+07 | 1.28E+07 | 1.59E+07 | 1.41E+07 | 1.91E+07 | 1.57E+07 | 1.22 | 0.28 | 0.053631251 |
| P38919 | Eukaryotic initiation factor 4A-III OS=Homo sapiens OX=9606 GN=EIF4A3 PE=1 SV=4 | 23 | 178 | 6 | 2.45E+07 | 2.80E+07 | 2.74E+07 | 2.92E+07 | 2.38E+07 | 2.37E+07 | 1.88E+07 | 2.30E+07 | 1.80E+07 | 1.95E+07 | 1.81E+07 | 2.04E+07 | 2.61E+07 | 1.96E+07 | 1.33 | 0.41 | 0.000477859 |
| O95782 | AP-2 complex subunit alpha-1 OS=Homo sapiens OX=9606 GN=AP2A1 PE=1 SV=3 | 18 | 81 | 6 | 1.08E+07 | 1.18E+07 | 1.09E+07 | 1.16E+07 | 1.23E+07 | 8.36E+06 | 8.85E+06 | 8.23E+06 | 8.22E+06 | 8.51E+06 | 8.29E+06 | 8.34E+06 | 1.09E+07 | 8.41E+06 | 1.30 | 0.38 | 0.005996316 |

### Supplementary Table 4. List of all proteins identified by label free proteomics analysis.

| Accession | Description | Coverage [%] | # PSMs | # Unique Peptides | Amitriptyline 1 | Amitriptyline 2 | Amitriptyline 3 | Amitriptyline 4 | Amitriptyline 5 | Amitriptyline 6 | Vehicle 1 | Vehicle 2 | Vehicle 3 | Vehicle 4 | Vehicle 5 | Vehicle 6 | Average amitriptyline | Average vehicle | Ratio amitriptyline/ vehicle | Log 2 fold change (amitriptyline /vehicle) | P-value |
|---|---|---|---|---|---|---|---|---|---|---|---|---|---|---|---|---|---|---|---|---|---|
| P09651 | Heterogeneous nuclear ribonucleoprotein A1 OS=Homo sapiens OX=9606 GN=HNRNPA1 PE=1 SV=5 | 23 | 98 | 6 | 1.76E+07 | 2.55E+07 | 1.97E+07 | 2.18E+07 | 1.83E+07 | 2.23E+07 | 1.45E+07 | 1.45E+07 | 1.12E+07 | 1.13E+07 | 1.17E+07 | 9.34E+06 | 2.09E+07 | 1.21E+07 | 1.73 | 0.79 | 0.000206591 |
| P07858 | Cathepsin B OS=Homo sapiens OX=9606 GN=CTSB PE=1 SV=3 | 24 | 92 | 6 | 2.81E+07 | 2.12E+07 | 2.62E+07 | 2.42E+07 | 2.77E+07 | 2.60E+07 | 1.45E+07 | 1.37E+07 | 1.13E+07 | 1.15E+07 | 1.21E+07 | 1.06E+07 | 2.56E+07 | 1.23E+07 | 2.08 | 1.06 | 4.00638E-06 |
| Q12904 | Aminoacyl tRNA synthase complex-interacting multifunctional protein 1 OS=Homo sapiens OX=9606 GN=AIMP1 PE=1 SV=2 | 25 | 72 | 6 | 2.07E+07 | 1.86E+07 | 1.56E+07 | 1.89E+07 | 1.63E+07 | 1.60E+07 | 9.11E+06 | 8.70E+06 | 8.62E+06 | 9.56E+06 | 1.06E+07 | 7.70E+06 | 1.77E+07 | 9.05E+06 | 1.96 | 0.97 | 2.65149E-05 |
| Q15555 | Microtubule-associated protein RP/EB family member 2 OS=Homo sapiens OX=9606 GN=MAPRE2 PE=1 SV=1 | 25 | 123 | 6 | 1.68E+07 | 1.75E+07 | 1.89E+07 | 1.87E+07 | 1.58E+07 | 1.95E+07 | 2.30E+07 | 2.80E+07 | 2.61E+07 | 1.98E+07 | 1.81E+07 | 2.10E+07 | 1.79E+07 | 2.27E+07 | 0.79 | -0.34 | 0.0257944 |
| Q99714 | 3-hydroxyacyl-CoA dehydrogenase type-2 OS=Homo sapiens OX=9606 GN=HSD17B10 PE=1 SV=3 | 44 | 66 | 6 | 1.76E+07 | 1.70E+07 | 1.98E+07 | 1.21E+07 | 1.21E+07 | 1.57E+07 | 1.41E+07 | 1.19E+07 | 1.03E+07 | 8.00E+06 | 1.03E+07 | 1.01E+07 | 1.57E+07 | 1.08E+07 | 1.46 | 0.54 | 0.011127501 |
| Q9Y2X3 | Nucleolar protein 58 OS=Homo sapiens OX=9606 GN=NOP58 PE=1 SV=1 | 16 | 96 | 6 | 1.32E+07 | 1.45E+07 | 1.91E+07 | 1.66E+07 | 2.05E+07 | 1.67E+07 | 9.61E+06 | 8.58E+06 | 1.05E+07 | 9.78E+06 | 1.15E+07 | 8.40E+06 | 1.68E+07 | 9.72E+06 | 1.73 | 0.79 | 0.000710381 |
| P30419 | Glycylpeptide N-tetradecanoyltransferase 1 OS=Homo sapiens OX=9606 GN=NMT1 PE=1 SV=2 | 23 | 92 | 6 | 3.09E+07 | 2.94E+07 | 2.74E+07 | 2.53E+07 | 2.73E+07 | 2.67E+07 | 2.32E+07 | 1.97E+07 | 2.32E+07 | 2.30E+07 | 1.94E+07 | 1.73E+07 | 2.78E+07 | 2.10E+07 | 1.33 | 0.41 | 0.00045387 |
| P01857 | Immunoglobulin heavy constant gamma 1 OS=Homo sapiens OX=9606 GN=IGHG1 PE=1 SV=1 | 38 | 33 | 6 | 1.12E+07 | 1.57E+07 | 2.12E+07 | 2.34E+07 | 2.25E+07 | 2.19E+07 | 3.45E+07 | 1.78E+07 | 2.08E+07 | 1.98E+07 | 2.03E+07 | 1.07E+08 | 1.93E+07 | 3.68E+07 | 0.53 | -0.93 | 0.280131754 |
| P18669 | Phosphoglycerate mutase 1 OS=Homo sapiens OX=9606 GN=PGAM1 PE=1 SV=2 | 37 | 125 | 6 | 4.05E+07 | 3.85E+07 | 2.86E+07 | 3.69E+07 | 3.00E+07 | 3.15E+07 | 3.95E+07 | 3.70E+07 | 2.89E+07 | 3.37E+07 | 3.36E+07 | 3.18E+07 | 3.43E+07 | 3.41E+07 | 1.01 | 0.01 | 0.92093334 |
| P04745 | Alpha-amylase 1 OS=Homo sapiens OX=9606 GN=AMY1A PE=1 SV=2 | 12 | 304 | 6 | 1.38E+08 | 1.43E+08 | 1.24E+08 | 1.37E+08 | 1.25E+08 | 1.18E+08 | 1.64E+08 | 1.49E+08 | 1.39E+08 | 1.58E+08 | 1.24E+08 | 1.33E+08 | 1.31E+08 | 1.44E+08 | 0.91 | -0.14 | 0.100332006 |
| P31146 | Coronin-1A OS=Homo sapiens OX=9606 GN=CORO1A PE=1 SV=4 | 21 | 135 | 6 | 2.56E+07 | 2.32E+07 | 2.21E+07 | 2.74E+07 | 2.44E+07 | 2.36E+07 | 3.64E+07 | 3.98E+07 | 2.85E+07 | 3.05E+07 | 2.62E+07 | 3.04E+07 | 2.44E+07 | 3.20E+07 | 0.76 | -0.39 | 0.013136093 |
| P35613 | Basigin OS=Homo sapiens OX=9606 GN=BSG PE=1 SV=2 | 21 | 131 | 6 | 5.11E+07 | 5.84E+07 | 5.56E+07 | 5.55E+07 | 5.05E+07 | 5.36E+07 | 2.78E+07 | 3.20E+07 | 3.03E+07 | 3.34E+07 | 2.75E+07 | 2.47E+07 | 5.41E+07 | 2.93E+07 | 1.85 | 0.88 | 8.32458E-08 |
| P08243 | Asparagine synthetase [glutamine-hydrolyzing] OS=Homo sapiens OX=9606 GN=ASNS PE=1 SV=4 | 18 | 52 | 6 | 9.54E+06 | 7.80E+06 | 9.53E+06 | 1.03E+07 | 6.90E+06 | 7.59E+06 | 6.53E+06 | 7.28E+06 | 6.34E+06 | 7.06E+06 | 6.78E+06 | 6.90E+06 | 8.61E+06 | 6.81E+06 | 1.26 | 0.34 | 0.02148764 |
| O60749 | Sorting nexin-2 OS=Homo sapiens OX=9606 GN=SNX2 PE=1 SV=2 | 20 | 131 | 6 | 2.13E+07 | 2.37E+07 | 2.27E+07 | 2.30E+07 | 2.12E+07 | 1.97E+07 | 2.27E+07 | 2.25E+07 | 2.10E+07 | 2.05E+07 | 2.18E+07 | 1.57E+07 | 2.19E+07 | 2.07E+07 | 1.06 | 0.08 | 0.34619375 |
| Q93099 | Homogentisate 1,2-dioxygenase OS=Homo sapiens OX=9606 GN=HGD PE=1 SV=2 | 19 | 158 | 6 | 1.75E+07 | 1.93E+07 | 2.31E+07 | 1.62E+07 | 2.53E+07 | 1.65E+07 | 2.50E+07 | 2.58E+07 | 2.19E+07 | 2.40E+07 | 2.30E+07 | 1.98E+07 | 1.97E+07 | 2.32E+07 | 0.85 | -0.24 | 0.077537453 |
| P27635 | 60S ribosomal protein L10 OS=Homo sapiens OX=9606 GN=RPL10 PE=1 SV=4 | 31 | 101 | 6 | 5.33E+07 | 4.82E+07 | 4.88E+07 | 5.54E+07 | 4.78E+07 | 4.54E+07 | 2.62E+07 | 2.29E+07 | 2.77E+07 | 3.15E+07 | 2.17E+07 | 1.91E+07 | 4.98E+07 | 2.48E+07 | 2.01 | 1.00 | 1.34317E-06 |
| P84103 | Serine/arginine-rich splicing factor 3 OS=Homo sapiens OX=9606 GN=SRSF3 PE=1 SV=1 | 41 | 152 | 6 | 4.95E+07 | 4.40E+07 | 3.66E+07 | 4.11E+07 | 4.37E+07 | 3.71E+07 | 3.21E+07 | 2.95E+07 | 2.70E+07 | 3.03E+07 | 2.61E+07 | 2.32E+07 | 4.20E+07 | 2.80E+07 | 1.50 | 0.58 | 0.000270987 |
| Q7L2H7 | Eukaryotic translation initiation factor 3 subunit M OS=Homo sapiens OX=9606 GN=EIF3M PE=1 SV=1 | 24 | 28 | 6 | 9.52E+06 | 8.40E+06 | 5.87E+06 | 5.96E+06 | 3.78E+06 | 5.44E+06 | 3.09E+06 | 3.76E+06 | 2.18E+05 | 1.39E+06 | 2.54E+06 | 1.33E+06 | 6.50E+06 | 2.05E+06 | 3.16 | 1.66 | 0.002022277 |
| P31689 | DnaJ homolog subfamily A member 1 OS=Homo sapiens OX=9606 GN=DNAJA1 PE=1 SV=2 | 23 | 55 | 6 | 1.21E+07 | 1.75E+07 | 1.47E+07 | 1.60E+07 | 1.73E+07 | 1.65E+07 | 6.92E+06 | 7.20E+06 | 7.66E+06 | 8.71E+06 | 6.53E+06 | 7.15E+06 | 1.57E+07 | 7.36E+06 | 2.13 | 1.09 | 5.72054E-05 |
| Q16394 | Exostosin-1 OS=Homo sapiens OX=9606 GN=EXT1 PE=1 SV=2 | 11 | 116 | 6 | 1.95E+07 | 1.89E+07 | 1.44E+07 | 1.38E+07 | 1.18E+07 | 1.75E+07 | 2.58E+07 | 2.82E+07 | 1.96E+07 | 1.61E+07 | 2.11E+07 | 2.37E+07 | 1.60E+07 | 2.24E+07 | 0.71 | -0.49 | 0.016203075 |
| P49721 | Proteasome subunit beta type-2 OS=Homo sapiens OX=9606 GN=PSMB2 PE=1 SV=1 | 46 | 75 | 6 | 2.10E+07 | 2.27E+07 | 1.97E+07 | 1.81E+07 | 1.79E+07 | 2.01E+07 | 2.06E+07 | 2.28E+07 | 1.84E+07 | 1.55E+07 | 2.10E+07 | 1.58E+07 | 1.99E+07 | 1.90E+07 | 1.05 | 0.07 | 0.54208023 |
| Q9UL46 | Proteasome activator complex subunit 2 OS=Homo sapiens OX=9606 GN=PSME2 PE=1 SV=4 | 35 | 28 | 6 | 7.37E+06 | 4.87E+06 | 4.62E+06 | 7.52E+06 | 6.49E+06 | 5.52E+06 | 4.52E+06 | 3.71E+06 | 2.78E+06 | 3.65E+06 | 4.86E+06 | 3.40E+06 | 6.06E+06 | 3.82E+06 | 1.59 | 0.67 | 0.005264919 |
| P52209 | 6-phosphogluconate dehydrogenase, decarboxylating OS=Homo sapiens OX=9606 GN=PGD PE=1 SV=3 | 20 | 120 | 6 | 4.99E+07 | 5.13E+07 | 5.08E+07 | 5.75E+07 | 5.54E+07 | 4.96E+07 | 6.58E+07 | 6.37E+07 | 7.60E+07 | 8.16E+07 | 5.83E+07 | 5.44E+07 | 5.24E+07 | 6.66E+07 | 0.79 | -0.35 | 0.018681226 |
| P30566 | Adenylosuccinate lyase OS=Homo sapiens OX=9606 GN=ADSL PE=1 SV=2 | 23 | 71 | 6 | 2.29E+07 | 2.59E+07 | 2.34E+07 | 2.17E+07 | 2.08E+07 | 2.37E+07 | 2.74E+07 | 2.63E+07 | 2.20E+07 | 2.36E+07 | 2.41E+07 | 1.69E+07 | 2.31E+07 | 2.34E+07 | 0.99 | -0.02 | 0.845010626 |
| Q15459 | Splicing factor 3A subunit 1 OS=Homo sapiens OX=9606 GN=SF3A1 PE=1 SV=1 | 14 | 35 | 6 | 8.78E+06 | 5.91E+06 | 7.94E+06 | 6.98E+06 | 6.81E+06 | 7.93E+06 | 4.34E+06 | 4.82E+06 | 6.06E+06 | 3.75E+06 | 4.32E+06 | 1.34E+06 | 7.39E+06 | 4.10E+06 | 1.80 | 0.85 | 0.002164005 |
| P15559 | NAD(P)H dehydrogenase [quinone] 1 OS=Homo sapiens OX=9606 GN=NQO1 PE=1 SV=1 | 31 | 57 | 6 | 2.66E+07 | 2.97E+07 | 3.93E+07 | 4.00E+07 | 3.68E+07 | 3.27E+07 | 2.74E+07 | 2.49E+07 | 3.00E+07 | 1.98E+07 | 1.69E+07 | 1.64E+07 | 3.42E+07 | 2.26E+07 | 1.51 | 0.60 | 0.004792335 |
| Q9UHD8 | Septin-9 OS=Homo sapiens OX=9606 GN=SEPTIN9 PE=1 SV=2 | 13 | 62 | 6 | 1.83E+07 | 2.21E+07 | 1.99E+07 | 2.03E+07 | 1.79E+07 | 1.80E+07 | 2.03E+07 | 1.54E+07 | 1.66E+07 | 1.70E+07 | 1.53E+07 | 1.27E+07 | 1.94E+07 | 1.62E+07 | 1.20 | 0.26 | 0.027412474 |
| Q99536 | Synaptic vesicle membrane protein VAT-1 homolog OS=Homo sapiens OX=9606 GN=VAT1 PE=1 SV=2 | 30 | 65 | 6 | 1.42E+07 | 1.51E+07 | 1.92E+07 | 1.47E+07 | 1.36E+07 | 1.73E+07 | 1.53E+07 | 1.49E+07 | 9.02E+06 | 1.78E+07 | 1.68E+07 | 1.40E+07 | 1.57E+07 | 1.46E+07 | 1.07 | 0.10 | 0.503314891 |
| Q92499 | ATP-dependent RNA helicase DDX1 OS=Homo sapiens OX=9606 GN=DDX1 PE=1 SV=2 | 10 | 43 | 6 | 7.75E+06 | 7.82E+06 | 9.85E+06 | 8.70E+06 | 9.69E+06 | 7.80E+06 | 6.82E+06 | 5.74E+06 | 6.85E+06 | 7.58E+06 | 6.76E+06 | 7.25E+06 | 8.60E+06 | 6.84E+06 | 1.26 | 0.33 | 0.005145748 |
| P08174 | Complement decay-accelerating factor OS=Homo sapiens OX=9606 GN=CD55 PE=1 SV=4 | 20 | 76 | 6 | 2.20E+07 | 1.70E+07 | 1.96E+07 | 1.78E+07 | 2.24E+07 | 1.69E+07 | 8.18E+06 | 7.82E+06 | 8.13E+06 | 6.92E+06 | 7.10E+06 | 6.14E+06 | 1.93E+07 | 7.38E+06 | 2.61 | 1.38 | 2.79711E-05 |
| Q1KMD3 | Heterogeneous nuclear ribonucleoprotein U-like protein 2 OS=Homo sapiens OX=9606 GN=HNRNPUL2 PE=1 SV=1 | 11 | 66 | 6 | 2.15E+07 | 2.07E+07 | 2.43E+07 | 1.80E+07 | 2.26E+07 | 2.01E+07 | 1.82E+07 | 2.04E+07 | 1.71E+07 | 2.28E+07 | 2.01E+07 | 1.56E+07 | 2.12E+07 | 1.90E+07 | 1.11 | 0.16 | 0.145696718 |
| P46776 | 60S ribosomal protein L27a OS=Homo sapiens OX=9606 GN=RPL27A PE=1 SV=2 | 31 | 117 | 6 | 4.46E+07 | 4.34E+07 | 4.90E+07 | 4.00E+07 | 3.82E+07 | 3.96E+07 | 3.00E+07 | 2.88E+07 | 2.53E+07 | 2.32E+07 | 2.60E+07 | 2.12E+07 | 4.25E+07 | 2.57E+07 | 1.65 | 0.72 | 1.6396E-05 |
| Q92743 | Serine protease HTRA1 OS=Homo sapiens OX=9606 GN=HTRA1 PE=1 SV=1 | 15 | 66 | 6 | 5.77E+06 | 1.22E+07 | 9.57E+06 | 1.27E+07 | 1.01E+07 | 1.15E+07 | 1.78E+07 | 2.12E+07 | 1.54E+07 | 1.47E+07 | 1.74E+07 | 1.85E+07 | 1.03E+07 | 1.75E+07 | 0.59 | -0.76 | 0.000459262 |
| P25786 | Proteasome subunit alpha type-1 OS=Homo sapiens OX=9606 GN=PSMA1 PE=1 SV=1 | 22 | 127 | 6 | 4.76E+07 | 4.74E+07 | 4.43E+07 | 3.88E+07 | 4.41E+07 | 3.70E+07 | 3.91E+07 | 4.90E+07 | 4.45E+07 | 4.22E+07 | 3.83E+07 | 3.15E+07 | 4.32E+07 | 4.08E+07 | 1.06 | 0.08 | 0.442285012 |
| P49419 | Alpha-aminoadipic semialdehyde dehydrogenase OS=Homo sapiens OX=9606 GN=ALDH7A1 PE=1 SV=5 | 17 | 72 | 6 | 1.41E+07 | 1.78E+07 | 1.34E+07 | 1.23E+07 | 1.25E+07 | 1.30E+07 | 1.59E+07 | 1.43E+07 | 1.12E+07 | 1.15E+07 | 1.10E+07 | 8.97E+06 | 1.38E+07 | 1.21E+07 | 1.14 | 0.19 | 0.227588154 |
| Q92598 | Heat shock protein 105 kDa OS=Homo sapiens OX=9606 GN=HSPH1 PE=1 SV=1 | 8 | 50 | 6 | 1.54E+07 | 1.46E+07 | 1.24E+07 | 1.23E+07 | 1.27E+07 | 1.18E+07 | 1.24E+07 | 9.98E+06 | 9.11E+06 | 1.10E+07 | 7.68E+06 | 6.71E+06 | 1.32E+07 | 9.48E+06 | 1.39 | 0.48 | 0.0062372 |
| O43684 | Mitotic checkpoint protein BUB3 OS=Homo sapiens OX=9606 GN=BUB3 PE=1 SV=1 | 21 | 87 | 6 | 1.86E+07 | 2.10E+07 | 2.21E+07 | 2.12E+07 | 2.19E+07 | 1.88E+07 | 1.61E+07 | 1.75E+07 | 1.66E+07 | 1.65E+07 | 1.53E+07 | 1.36E+07 | 2.06E+07 | 1.59E+07 | 1.29 | 0.37 | 0.000247396 |
| P49736 | DNA replication licensing factor MCM2 OS=Homo sapiens OX=9606 GN=MCM2 PE=1 SV=4 | 9 | 56 | 6 | 1.31E+07 | 1.43E+07 | 1.69E+07 | 1.33E+07 | 1.60E+07 | 1.38E+07 | 9.81E+06 | 9.82E+06 | 8.69E+06 | 8.93E+06 | 9.03E+06 | 7.76E+06 | 1.46E+07 | 9.01E+06 | 1.62 | 0.69 | 7.07947E-05 |
| Q9H4A4 | Aminopeptidase B OS=Homo sapiens OX=9606 GN=RNPEP PE=1 SV=2 | 12 | 40 | 6 | 1.39E+07 | 1.29E+07 | 1.26E+07 | 1.36E+07 | 1.25E+07 | 1.35E+07 | 8.93E+06 | 1.15E+07 | 4.36E+06 | 3.67E+06 | 4.94E+06 | 6.92E+06 | 1.32E+07 | 6.71E+06 | 1.96 | 0.97 | 0.002910677 |
| Q9NVA2 | Septin-11 OS=Homo sapiens OX=9606 GN=SEPTIN11 PE=1 SV=3 | 21 | 129 | 6 | 3.29E+07 | 3.33E+07 | 2.63E+07 | 2.76E+07 | 2.24E+07 | 2.42E+07 | 2.91E+07 | 2.87E+07 | 1.98E+07 | 1.91E+07 | 1.94E+07 | 2.20E+07 | 2.78E+07 | 2.30E+07 | 1.21 | 0.27 | 0.099519283 |
| P23246 | Splicing factor, proline- and glutamine-rich OS=Homo sapiens OX=9606 GN=SFPQ PE=1 SV=2 | 12 | 81 | 6 | 2.68E+07 | 2.37E+07 | 3.41E+07 | 2.87E+07 | 3.32E+07 | 2.77E+07 | 2.03E+07 | 2.21E+07 | 1.95E+07 | 1.94E+07 | 1.84E+07 | 1.63E+07 | 2.91E+07 | 1.93E+07 | 1.50 | 0.59 | 0.000880256 |
| Q12874 | Splicing factor 3A subunit 3 OS=Homo sapiens OX=9606 GN=SF3A3 PE=1 SV=1 | 17 | 44 | 6 | 1.28E+07 | 1.25E+07 | 1.14E+07 | 1.23E+07 | 1.26E+07 | 1.15E+07 | 8.11E+06 | 7.72E+06 | 7.08E+06 | 9.39E+06 | 7.56E+06 | 6.80E+06 | 1.22E+07 | 7.77E+06 | 1.57 | 0.65 | 5.90732E-06 |
| P13674 | Prolyl 4-hydroxylase subunit alpha-1 OS=Homo sapiens OX=9606 GN=P4HA1 PE=1 SV=2 | 17 | 60 | 6 | 7.72E+06 | 8.28E+06 | 6.85E+06 | 4.89E+06 | 4.89E+06 | 5.87E+06 | 8.82E+06 | 7.62E+06 | 6.69E+06 | 7.83E+06 | 7.50E+06 | 6.00E+06 | 6.42E+06 | 7.41E+06 | 0.87 | -0.21 | 0.195447961 |
| P09871 | Complement C1s subcomponent OS=Homo sapiens OX=9606 GN=C1S PE=1 SV=1 | 11 | 13 | 6 | 5.76E+05 | | | | | | 8.86E+05 | | | | | 1.88E+07 | 5.76E+05 | 9.86E+06 | 0.06 | -4.10 | #DIV/0! |
| Q13435 | Splicing factor 3B subunit 2 OS=Homo sapiens OX=9606 GN=SF3B2 PE=1 SV=2 | 7 | 44 | 6 | 1.58E+07 | 1.47E+07 | 1.68E+07 | 1.44E+07 | 2.00E+07 | 1.64E+07 | 7.08E+06 | 9.87E+06 | 1.29E+07 | 2.82E+07 | 7.94E+06 | 6.02E+06 | 1.63E+07 | 1.20E+07 | 1.36 | 0.45 | 0.262811855 |
| O15372 | Eukaryotic translation initiation factor 3 subunit H OS=Homo sapiens OX=9606 GN=EIF3H PE=1 SV=1 | 24 | 73 | 6 | 2.12E+07 | 1.88E+07 | 2.90E+07 | 2.79E+07 | 2.95E+07 | 2.37E+07 | 1.66E+07 | 1.75E+07 | 1.88E+07 | 1.53E+07 | 1.78E+07 | 1.32E+07 | 2.50E+07 | 1.65E+07 | 1.51 | 0.60 | 0.003848281 |
| P62263 | 40S ribosomal protein S14 OS=Homo sapiens OX=9606 GN=RPS14 PE=1 SV=3 | 39 | 72 | 6 | 3.80E+07 | 2.58E+07 | 2.70E+07 | 2.61E+07 | 2.73E+07 | 2.49E+07 | 1.62E+07 | 1.65E+07 | 2.15E+07 | 1.78E+07 | 1.57E+07 | 1.24E+07 | 2.82E+07 | 1.67E+07 | 1.69 | 0.76 | 0.001061454 |
| Q14517 | Protocadherin Fat 1 OS=Homo sapiens OX=9606 GN=FAT1 PE=1 SV=2 | 2 | 46 | 6 | 1.31E+07 | 1.45E+07 | 1.54E+07 | 1.25E+07 | 1.14E+07 | 1.40E+07 | 1.48E+07 | 7.13E+06 | 7.92E+06 | 1.27E+07 | 1.40E+07 | 1.14E+07 | 1.35E+07 | 1.13E+07 | 1.19 | 0.25 | 0.167339673 |
| Q9UBG0 | C-type mannose receptor 2 OS=Homo sapiens OX=9606 GN=MRC2 PE=1 SV=2 | 6 | 61 | 6 | 1.47E+07 | 1.32E+07 | 1.19E+07 | 1.39E+07 | 1.37E+07 | 1.37E+07 | 1.71E+07 | 1.67E+07 | 1.16E+07 | 1.48E+07 | 1.33E+07 | 1.23E+07 | 1.35E+07 | 1.43E+07 | 0.94 | -0.08 | 0.46326587 |
| Q99733 | Nucleosome assembly protein 1-like 4 OS=Homo sapiens OX=9606 GN=NAP1L4 PE=1 SV=1 | 24 | 51 | 6 | 1.84E+07 | 1.84E+07 | 1.49E+07 | 1.42E+07 | 1.32E+07 | 1.54E+07 | 1.10E+07 | 7.33E+06 | 8.50E+06 | 6.57E+06 | 8.42E+06 | 7.27E+06 | 1.57E+07 | 8.18E+06 | 1.92 | 0.94 | 7.26046E-05 |
| Q16851 | UTP--glucose-1-phosphate uridylyltransferase OS=Homo sapiens OX=9606 GN=UGP2 PE=1 SV=5 | 16 | 36 | 6 | 8.19E+06 | 6.86E+06 | 6.14E+06 | 6.28E+06 | 4.99E+06 | 6.00E+06 | 1.00E+07 | 9.29E+06 | 8.24E+06 | 7.16E+06 | 7.37E+06 | 6.66E+06 | 6.41E+06 | 8.12E+06 | 0.79 | -0.34 | 0.033310502 |
| Q13683 | Integrin alpha-7 OS=Homo sapiens OX=9606 GN=ITGA7 PE=1 SV=3 | 7 | 23 | 6 | 7.75E+06 | 8.52E+06 | 7.25E+06 | 8.76E+06 | 9.57E+06 | 6.80E+06 | 4.34E+06 | 5.19E+06 | 5.29E+06 | 4.20E+06 | 3.87E+06 | 2.95E+06 | 8.11E+06 | 4.31E+06 | 1.88 | 0.91 | 4.79915E-05 |
| P11166 | Solute carrier family 2, facilitated glucose transporter member 1 OS=Homo sapiens OX=9606 GN=SLC2A1 PE=1 SV=2 | 12 | 187 | 6 | 8.92E+07 | 6.32E+07 | 8.55E+07 | 6.35E+07 | 8.25E+07 | 7.13E+07 | 6.94E+07 | 6.55E+07 | 8.53E+07 | 5.97E+07 | 8.30E+07 | 6.68E+07 | 7.59E+07 | 7.16E+07 | 1.06 | 0.08 | 0.507956287 |
| Q16222 | UDP-N-acetylhexosamine pyrophosphorylase OS=Homo sapiens OX=9606 GN=UAP1 PE=1 SV=3 | 17 | 20 | 6 | 6.54E+06 | 6.43E+06 | 6.16E+06 | 4.99E+06 | 5.56E+06 | 5.04E+06 | 7.44E+06 | 6.76E+06 | 5.90E+06 | 3.17E+06 | 4.53E+06 | 4.35E+06 | 5.79E+06 | 5.36E+06 | 1.08 | 0.11 | 0.568264035 |
| Q7Z6Z7 | E3 ubiquitin-protein ligase HUWE1 OS=Homo sapiens OX=9606 GN=HUWE1 PE=1 SV=3 | 2 | 29 | 6 | 5.56E+06 | 4.88E+06 | 6.19E+06 | 5.88E+06 | 7.25E+06 | 5.81E+06 | 2.73E+06 | 4.26E+06 | 5.58E+06 | 4.87E+06 | 4.21E+06 | 1.09E+07 | 5.93E+06 | 5.43E+06 | 1.09 | 0.13 | 0.696891237 |
| Q93034 | Cullin-5 OS=Homo sapiens OX=9606 GN=CUL5 PE=1 SV=4 | 9 | 86 | 6 | 2.76E+08 | 2.68E+08 | 3.76E+08 | 4.26E+08 | 4.57E+08 | 4.00E+08 | 4.00E+08 | 4.29E+08 | 6.89E+08 | 8.41E+08 | 8.59E+08 | 6.44E+08 | 3.67E+08 | 6.44E+08 | 0.57 | -0.81 | 0.016371436 |
| Q8IWA5 | Choline transporter-like protein 2 OS=Homo sapiens OX=9606 GN=SLC44A2 PE=1 SV=3 | 9 | 42 | 6 | 8.62E+06 | 8.41E+06 | 8.19E+06 | 1.51E+07 | 8.46E+06 | 7.89E+06 | 5.70E+06 | 5.34E+06 | 9.07E+06 | 8.18E+06 | 7.09E+06 | 5.02E+06 | 9.44E+06 | 6.73E+06 | 1.40 | 0.49 | 0.073208123 |
| Q9BUF5 | Tubulin beta-6 chain OS=Homo sapiens OX=9606 GN=TUBB6 PE=1 SV=1 | 60 | 2411 | 7 | 6.44E+07 | 4.91E+07 | 4.98E+07 | 4.22E+07 | 4.95E+07 | 4.27E+07 | 4.96E+07 | 4.88E+07 | 3.95E+07 | 3.66E+07 | 4.34E+07 | 4.43E+07 | 4.96E+07 | 4.37E+07 | 1.14 | 0.18 | 0.162762495 |
| P61204 | ADP-ribosylation factor 3 OS=Homo sapiens OX=9606 GN=ARF3 PE=1 SV=2 | 60 | 426 | 7 | 6.60E+07 | 8.32E+07 | 8.24E+07 | 8.45E+07 | 6.34E+07 | 8.96E+07 | 9.20E+07 | 1.04E+08 | 7.19E+07 | 6.10E+07 | 9.23E+07 | 7.52E+07 | 7.82E+07 | 8.27E+07 | 0.95 | -0.08 | 0.581869184 |
| P61163 | Alpha-centractin OS=Homo sapiens OX=9606 GN=ACTR1A PE=1 SV=1 | 52 | 238 | 7 | 9.21E+07 | 8.38E+07 | 8.41E+07 | 8.55E+07 | 7.50E+07 | 8.56E+07 | 7.73E+07 | 7.98E+07 | 5.94E+07 | 6.45E+07 | 6.61E+07 | 6.40E+07 | 8.44E+07 | 6.85E+07 | 1.23 | 0.30 | 0.003470943 |
| P15311 | Ezrin OS=Homo sapiens OX=9606 GN=EZR PE=1 SV=4 | 26 | 288 | 7 | 2.53E+07 | 1.48E+07 | 2.03E+07 | 1.44E+07 | 1.65E+07 | 1.67E+07 | 2.35E+07 | 1.17E+07 | 1.76E+07 | 1.60E+07 | 1.40E+07 | 1.62E+07 | 1.80E+07 | 1.65E+07 | 1.09 | 0.13 | 0.530375089 |
| P69905 | Hemoglobin subunit alpha OS=Homo sapiens OX=9606 GN=HBA1 PE=1 SV=2 | 38 | 3552 | 7 | 4.25E+09 | 4.41E+09 | 5.57E+09 | 5.43E+09 | 5.89E+09 | 4.83E+09 | 5.95E+09 | 5.61E+09 | 7.16E+09 | 6.91E+09 | 6.52E+09 | 6.70E+09 | 5.06E+09 | 6.47E+09 | 0.78 | -0.36 | 0.003064574 |
| P67809 | Y-box-binding protein 1 OS=Homo sapiens OX=9606 GN=YBX1 PE=1 SV=3 | 34 | 118 | 7 | 2.10E+07 | 2.50E+07 | 2.39E+07 | 2.50E+07 | 2.24E+07 | 2.62E+07 | 2.41E+07 | 2.70E+07 | 2.33E+07 | 2.64E+07 | 2.46E+07 | 1.96E+07 | 2.39E+07 | 2.42E+07 | 0.99 | -0.01 | 0.863935241 |
| P55209 | Nucleosome assembly protein 1-like 1 OS=Homo sapiens OX=9606 GN=NAP1L1 PE=1 SV=1 | 36 | 142 | 7 | 3.19E+07 | 3.15E+07 | 3.25E+07 | 2.79E+07 | 2.60E+07 | 2.92E+07 | 2.32E+07 | 1.95E+07 | 1.65E+07 | 1.69E+07 | 1.80E+07 | 1.85E+07 | 2.98E+07 | 1.88E+07 | 1.59 | 0.67 | 1.72108E-05 |
| P51884 | Lumican OS=Homo sapiens OX=9606 GN=LUM PE=1 SV=2 | 14 | 923 | 7 | 5.41E+08 | 5.37E+08 | 7.21E+08 | 5.90E+08 | 7.00E+08 | 6.26E+08 | 5.77E+08 | 6.52E+08 | 8.30E+08 | 7.90E+08 | 9.92E+08 | 7.55E+08 | 6.19E+08 | 7.66E+08 | 0.81 | -0.31 | 0.061402872 |
| P29992 | Guanine nucleotide-binding protein subunit alpha-11 OS=Homo sapiens OX=9606 GN=GNA11 PE=1 SV=2 | 33 | 141 | 7 | 4.22E+07 | 3.71E+07 | 3.72E+07 | 3.75E+07 | 3.46E+07 | 3.91E+07 | 3.01E+07 | 2.91E+07 | 2.51E+07 | 2.71E+07 | 2.35E+07 | 2.52E+07 | 3.79E+07 | 2.67E+07 | 1.42 | 0.51 | 1.79168E-05 |
| P62979 | Ubiquitin-40S ribosomal protein S27a OS=Homo sapiens OX=9606 GN=RPS27A PE=1 SV=2 | 44 | 732 | 7 | 3.76E+08 | 5.17E+08 | 4.74E+08 | 4.74E+08 | 4.48E+08 | 4.33E+08 | 2.69E+08 | 2.93E+08 | 3.61E+08 | 3.33E+08 | 3.24E+08 | 2.65E+08 | 4.54E+08 | 3.08E+08 | 1.47 | 0.56 | 0.000186257 |
| P60953 | Cell division control protein 42 homolog OS=Homo sapiens OX=9606 GN=CDC42 PE=1 SV=2 | 47 | 118 | 7 | 3.86E+07 | 3.29E+07 | 3.40E+07 | 2.59E+07 | 2.64E+07 | 3.38E+07 | 3.29E+07 | 3.44E+07 | 2.51E+07 | 2.44E+07 | 2.82E+07 | 2.73E+07 | 3.19E+07 | 2.87E+07 | 1.11 | 0.15 | 0.245962254 |
| Q06033 | Inter-alpha-trypsin inhibitor heavy chain H3 OS=Homo sapiens OX=9606 GN=ITIH3 PE=1 SV=2 | 6 | 320 | 7 | 8.87E+07 | 1.34E+08 | 8.37E+07 | 9.72E+07 | 8.58E+07 | 1.04E+08 | 1.13E+08 | 1.82E+08 | 1.13E+08 | 1.04E+08 | 1.09E+08 | 1.25E+08 | 9.89E+07 | 1.24E+08 | 0.80 | -0.33 | 0.107814671 |
| Q96P70 | Importin-9 OS=Homo sapiens OX=9606 GN=IPO9 PE=1 SV=3 | 12 | 72 | 7 | 1.52E+08 | 1.10E+08 | 1.34E+08 | 1.50E+08 | 1.37E+08 | 1.59E+07 | 1.58E+08 | 1.12E+08 | 1.06E+08 | 8.80E+07 | 1.36E+08 | 1.41E+08 | 1.17E+08 | 1.24E+08 | 0.94 | -0.08 | 0.774235564 |
| Q15758 | Neutral amino acid transporter B(0) OS=Homo sapiens OX=9606 GN=SLC1A5 PE=1 SV=2 | 22 | 148 | 7 | 8.06E+07 | 6.87E+07 | 7.93E+07 | 6.85E+07 | 6.46E+07 | 9.39E+07 | 5.88E+07 | 4.92E+07 | 4.55E+07 | 4.17E+07 | 5.06E+07 | 4.38E+07 | 7.59E+07 | 4.83E+07 | 1.57 | 0.65 | 0.000657092 |
| P07305 | Histone H1.0 OS=Homo sapiens OX=9606 GN=H1-0 PE=1 SV=3 | 24 | 907 | 7 | 2.67E+08 | 3.26E+08 | 2.72E+08 | 3.12E+08 | 2.62E+08 | 2.96E+08 | 2.60E+08 | 3.13E+08 | 2.60E+08 | 2.73E+08 | 2.59E+08 | 2.51E+08 | 2.89E+08 | 2.69E+08 | 1.07 | 0.10 | 0.195341041 |
| P12109 | Collagen alpha-1(VI) chain OS=Homo sapiens OX=9606 GN=COL6A1 PE=1 SV=3 | 9 | 244 | 7 | 1.13E+08 | 1.23E+08 | 9.71E+07 | 1.10E+08 | 1.10E+08 | 9.07E+07 | 1.54E+08 | 1.61E+08 | 1.53E+08 | 1.44E+08 | 1.42E+08 | 1.43E+08 | 1.07E+08 | 1.50E+08 | 0.72 | -0.48 | 4.92381E-05 |
| P51665 | 26S proteasome non-ATPase regulatory subunit 7 OS=Homo sapiens OX=9606 GN=PSMD7 PE=1 SV=2 | 28 | 80 | 7 | 2.32E+07 | 2.99E+07 | 1.96E+07 | 2.09E+07 | 1.95E+07 | 2.42E+07 | 1.29E+07 | 1.94E+07 | 9.98E+06 | 1.06E+07 | 1.62E+07 | 1.30E+07 | 2.29E+07 | 1.37E+07 | 1.67 | 0.74 | 0.001720333 |
| P04004 | Vitronectin OS=Homo sapiens OX=9606 GN=VTN PE=1 SV=1 | 14 | 362 | 7 | 4.73E+08 | 3.08E+08 | 2.89E+08 | 3.47E+08 | 2.78E+08 | 3.32E+08 | 3.78E+08 | 2.76E+08 | 2.12E+08 | 2.46E+08 | 2.51E+08 | 2.83E+08 | 3.38E+08 | 2.74E+08 | 1.23 | 0.30 | 0.119987945 |
| Q13501 | Sequestosome-1 OS=Homo sapiens OX=9606 GN=SQSTM1 PE=1 SV=1 | 30 | 52 | 7 | 2.63E+07 | 2.28E+07 | 2.24E+07 | 2.59E+07 | 2.27E+07 | 2.87E+07 | 1.89E+06 | 2.80E+06 | 2.35E+06 | | 5.79E+05 | | 2.48E+07 | 1.91E+06 | 13.01 | 3.70 | 2.64697E-07 |
| P02794 | Ferritin heavy chain OS=Homo sapiens OX=9606 GN=FTH1 PE=1 SV=2 | 56 | 124 | 7 | 3.11E+07 | 3.84E+07 | 4.63E+07 | 4.85E+07 | 4.42E+07 | 4.17E+07 | 5.34E+07 | 5.38E+07 | 6.87E+07 | 6.62E+07 | 5.73E+07 | 5.33E+07 | 4.17E+07 | 5.88E+07 | 0.71 | -0.50 | 0.00119783 |
| Q9HBI1 | Beta-parvin OS=Homo sapiens OX=9606 GN=PARVB PE=1 SV=1 | 22 | 127 | 7 | 2.56E+07 | 2.37E+07 | 2.59E+07 | 2.53E+07 | 2.58E+07 | 2.44E+07 | 3.55E+07 | 3.27E+07 | 3.31E+07 | 2.85E+07 | 3.04E+07 | 2.72E+07 | 2.51E+07 | 3.12E+07 | 0.80 | -0.31 | 0.003782095 |
| Q9NR45 | Sialic acid synthase OS=Homo sapiens OX=9606 GN=NANS PE=1 SV=2 | 35 | 134 | 7 | 4.37E+07 | 4.82E+07 | 3.69E+07 | 3.73E+07 | 3.26E+07 | 3.82E+07 | 3.31E+07 | 4.37E+07 | 3.20E+07 | 2.79E+07 | 3.34E+07 | 2.59E+07 | 3.95E+07 | 3.27E+07 | 1.21 | 0.27 | 0.072020388 |
| Q13813 | Spectrin alpha chain, non-erythrocytic 1 OS=Homo sapiens OX=9606 GN=SPTAN1 PE=1 SV=3 | 4 | 63 | 7 | 7.78E+06 | 1.14E+07 | 9.48E+06 | 1.11E+07 | 8.82E+06 | 1.00E+07 | 6.43E+06 | 6.21E+06 | 6.19E+06 | 5.88E+06 | 6.22E+06 | 4.38E+06 | 9.77E+06 | 5.88E+06 | 1.66 | 0.73 | 0.000340235 |
| P49747 | Cartilage oligomeric matrix protein OS=Homo sapiens OX=9606 GN=COMP PE=1 SV=2 | 18 | 211 | 7 | 8.29E+07 | 8.67E+07 | 8.78E+07 | 9.13E+07 | 8.82E+07 | 8.13E+07 | 1.16E+08 | 1.28E+08 | 1.07E+08 | 1.11E+08 | 1.06E+08 | 9.83E+07 | 8.63E+07 | 1.11E+08 | 0.78 | -0.36 | 0.001160575 |
| O43505 | Beta-1,4-glucuronyltransferase 1 OS=Homo sapiens OX=9606 GN=B4GAT1 PE=1 SV=1 | 25 | 124 | 7 | 3.48E+07 | 3.31E+07 | 2.68E+07 | 2.90E+07 | 2.98E+07 | 4.24E+07 | 5.31E+07 | 5.40E+07 | 5.23E+07 | 3.13E+07 | 3.89E+07 | 3.61E+07 | 3.27E+07 | 4.43E+07 | 0.74 | -0.44 | 0.037966268 |
| P62195 | 26S proteasome regulatory subunit 8 OS=Homo sapiens OX=9606 GN=PSMC5 PE=1 SV=1 | 27 | 133 | 7 | 2.47E+07 | 2.74E+07 | 2.06E+07 | 1.98E+07 | 2.09E+07 | 2.76E+07 | 1.83E+07 | 1.74E+07 | 1.50E+07 | 1.32E+07 | 1.69E+07 | 1.30E+07 | 2.35E+07 | 1.56E+07 | 1.50 | 0.59 | 0.001466562 |
| Q13347 | Eukaryotic translation initiation factor 3 subunit I OS=Homo sapiens OX=9606 GN=EIF3I PE=1 SV=1 | 37 | 92 | 7 | 2.65E+07 | 2.62E+07 | 2.40E+07 | 2.58E+07 | 2.26E+07 | 2.15E+07 | 1.74E+07 | 1.59E+07 | 1.50E+07 | 1.63E+07 | 1.50E+07 | 1.40E+07 | 2.44E+07 | 1.56E+07 | 1.56 | 0.65 | 1.8692E-05 |
| Q14108 | Lysosome membrane protein 2 OS=Homo sapiens OX=9606 GN=SCARB2 PE=1 SV=2 | 24 | 74 | 7 | 2.33E+07 | 2.90E+07 | 2.48E+07 | 2.64E+07 | 2.25E+07 | 2.31E+07 | 7.18E+06 | 8.13E+06 | 9.35E+06 | 1.03E+07 | 8.65E+06 | 9.93E+06 | 2.49E+07 | 8.93E+06 | 2.78 | 1.48 | 1.73165E-06 |
| P62266 | 40S ribosomal protein S23 OS=Homo sapiens OX=9606 GN=RPS23 PE=1 SV=3 | 49 | 127 | 7 | 8.56E+07 | 7.89E+07 | 5.66E+07 | 7.36E+07 | 6.22E+07 | 5.39E+07 | 4.96E+07 | 5.10E+07 | 3.87E+07 | 5.01E+07 | 3.43E+07 | 3.34E+07 | 6.85E+07 | 4.29E+07 | 1.60 | 0.68 | 0.002918802 |
| P14550 | Aldo-keto reductase family 1 member A1 OS=Homo sapiens OX=9606 GN=AKR1A1 PE=1 SV=3 | 28 | 293 | 7 | 1.61E+08 | 1.78E+08 | 1.56E+08 | 2.13E+08 | 1.65E+08 | 1.94E+08 | 2.25E+08 | 2.42E+08 | 2.79E+08 | 2.99E+08 | 2.58E+08 | 2.03E+08 | 1.78E+08 | 2.51E+08 | 0.71 | -0.50 | 0.002271934 |
| P29692 | Elongation factor 1-delta OS=Homo sapiens OX=9606 GN=EEF1D PE=1 SV=5 | 30 | 183 | 7 | 4.45E+07 | 5.20E+07 | 4.26E+07 | 4.71E+07 | 5.52E+07 | 4.31E+07 | 3.27E+07 | 4.11E+07 | 4.09E+07 | 3.40E+07 | 3.40E+07 | 3.28E+07 | 4.74E+07 | 3.59E+07 | 1.32 | 0.40 | 0.00172015 |
| Q13155 | Aminoacyl tRNA synthase complex-interacting multifunctional protein 2 OS=Homo sapiens OX=9606 GN=AIMP2 PE=1 SV=2 | 32 | 58 | 7 | 2.61E+07 | 2.24E+07 | 1.98E+07 | 1.87E+07 | 2.14E+07 | 2.38E+07 | 1.19E+07 | 1.07E+07 | 1.03E+07 | 1.05E+07 | 9.29E+06 | 9.44E+06 | 2.20E+07 | 1.04E+07 | 2.13 | 1.09 | 4.55228E-05 |
| O75131 | Copine-3 OS=Homo sapiens OX=9606 GN=CPNE3 PE=1 SV=1 | 19 | 132 | 7 | 3.23E+07 | 2.94E+07 | 2.88E+07 | 2.83E+07 | 3.04E+07 | 2.75E+07 | 1.87E+07 | 1.47E+07 | 1.40E+07 | 1.63E+07 | 1.83E+07 | 1.41E+07 | 2.94E+07 | 1.60E+07 | 1.84 | 0.88 | 3.93441E-07 |
| P28074 | Proteasome subunit beta type-5 OS=Homo sapiens OX=9606 GN=PSMB5 PE=1 SV=3 | 27 | 84 | 7 | 1.84E+07 | 2.00E+07 | 1.95E+07 | 2.05E+07 | 1.93E+07 | 1.57E+07 | 1.98E+07 | 1.88E+07 | 2.02E+07 | 1.87E+07 | 1.69E+07 | 1.49E+07 | 1.89E+07 | 1.82E+07 | 1.04 | 0.05 | 0.536343382 |
| P16152 | Carbonyl reductase [NADPH] 1 OS=Homo sapiens OX=9606 GN=CBR1 PE=1 SV=3 | 38 | 85 | 7 | 2.02E+07 | 2.17E+07 | 1.56E+07 | 1.73E+07 | 1.68E+07 | 1.65E+07 | 2.20E+07 | 2.08E+07 | 1.51E+07 | 1.74E+07 | 1.72E+07 | 1.65E+07 | 1.80E+07 | 1.82E+07 | 0.99 | -0.01 | 0.918345136 |
| P25787 | Proteasome subunit alpha type-2 OS=Homo sapiens OX=9606 GN=PSMA2 PE=1 SV=2 | 35 | 106 | 7 | 3.09E+07 | 2.36E+07 | 2.35E+07 | 2.90E+07 | 2.29E+07 | 2.87E+07 | 3.31E+07 | 2.49E+07 | 2.23E+07 | 2.09E+07 | 2.40E+07 | 2.04E+07 | 2.64E+07 | 2.43E+07 | 1.09 | 0.12 | 0.386872424 |
| P02649 | Apolipoprotein E OS=Homo sapiens OX=9606 GN=APOE PE=1 SV=1 | 25 | 348 | 7 | 2.17E+08 | 2.16E+08 | 1.98E+08 | 2.01E+08 | 1.89E+08 | 1.87E+08 | 2.27E+08 | 2.32E+08 | 2.00E+08 | 2.19E+08 | 2.07E+08 | 2.15E+08 | 2.01E+08 | 2.17E+08 | 0.93 | -0.11 | 0.055619113 |
| P07910 | Heterogeneous nuclear ribonucleoproteins C1/C2 OS=Homo sapiens OX=9606 GN=HNRNPC PE=1 SV=4 | 26 | 255 | 7 | 8.48E+07 | 9.15E+07 | 7.37E+07 | 9.42E+07 | 7.60E+07 | 7.29E+07 | 5.74E+07 | 4.89E+07 | 5.63E+07 | 5.68E+07 | 3.94E+07 | 4.69E+07 | 8.22E+07 | 5.10E+07 | 1.61 | 0.69 | 9.39164E-05 |
| P32119 | Peroxiredoxin-2 OS=Homo sapiens OX=9606 GN=PRDX2 PE=1 SV=5 | 34 | 134 | 7 | 4.30E+07 | 3.34E+07 | 3.70E+07 | 3.63E+07 | 2.72E+07 | 3.13E+07 | 3.00E+07 | 2.52E+07 | 2.24E+07 | 1.94E+07 | 1.87E+07 | 1.91E+07 | 3.47E+07 | 2.25E+07 | 1.54 | 0.63 | 0.001765888 |
| P50454 | Serpin H1 OS=Homo sapiens OX=9606 GN=SERPINH1 PE=1 SV=2 | 23 | 116 | 7 | 2.30E+07 | 2.66E+07 | 1.73E+07 | 2.30E+07 | 2.07E+07 | 2.16E+07 | 3.03E+07 | 3.04E+07 | 2.20E+07 | 2.28E+07 | 2.65E+07 | 1.77E+07 | 2.20E+07 | 2.50E+07 | 0.88 | -0.18 | 0.25874207 |
| O00154 | Cytosolic acyl coenzyme A thioester hydrolase OS=Homo sapiens OX=9606 GN=ACOT7 PE=1 SV=3 | 24 | 66 | 7 | 2.76E+07 | 3.13E+07 | 1.84E+07 | 2.40E+07 | 2.50E+07 | 2.45E+07 | 2.40E+07 | 2.46E+07 | 1.61E+07 | 1.55E+07 | 1.63E+07 | 1.70E+07 | 2.51E+07 | 1.89E+07 | 1.33 | 0.41 | 0.029559103 |
| P37802 | Transgelin-2 OS=Homo sapiens OX=9606 GN=TAGLN2 PE=1 SV=3 | 50 | 114 | 7 | 3.21E+07 | 3.31E+07 | 3.11E+07 | 3.05E+07 | 2.34E+07 | 2.66E+07 | 3.05E+07 | 2.42E+07 | 2.14E+07 | 1.83E+07 | 2.01E+07 | 2.35E+07 | 2.95E+07 | 2.30E+07 | 1.28 | 0.36 | 0.019334721 |

### Supplementary Table 4. List of all proteins identified by label free proteomics analysis.

| Accession | Description | Coverage [%] | # PSMs | # Unique Peptides | Amitriptyline 1 | Amitriptyline 2 | Amitriptyline 3 | Amitriptyline 4 | Amitriptyline 5 | Amitriptyline 6 | Vehicle 1 | Vehicle 2 | Vehicle 3 | Vehicle 4 | Vehicle 5 | Vehicle 6 | Average amitriptyline | Average vehicle | Ratio amitriptyline/ vehicle | Log 2 fold change (amitriptyline /vehicle) | P-value |
|---|---|---|---|---|---|---|---|---|---|---|---|---|---|---|---|---|---|---|---|---|---|
| P00736 | Complement C1r subcomponent OS=Homo sapiens OX=9606 GN=C1R PE=1 SV=2 | 14 | 43 | 7 | 5.75E+06 | 3.35E+06 | 3.62E+06 | 6.35E+06 | 5.03E+06 | 5.67E+06 | 8.38E+06 | 7.63E+06 | 9.83E+06 | 7.21E+06 | 6.76E+06 | 3.00E+07 | 4.96E+06 | 1.16E+07 | 0.43 | -1.23 | 0.131891979 |
| Q9NR30 | Nucleolar RNA helicase 2 OS=Homo sapiens OX=9606 GN=DDX21 PE=1 SV=5 | 14 | 64 | 7 | 8.58E+06 | 9.13E+06 | 8.47E+06 | 7.28E+06 | 6.59E+06 | 7.86E+06 | 7.70E+06 | 6.80E+06 | 4.86E+06 | 3.92E+06 | 5.67E+06 | 5.53E+06 | 7.98E+06 | 5.75E+06 | 1.39 | 0.47 | 0.008790562 |
| P32969 | 60S ribosomal protein L9 OS=Homo sapiens OX=9606 GN=RPL9 PE=1 SV=1 | 59 | 59 | 7 | 1.53E+07 | 2.64E+07 | 1.97E+07 | 1.74E+07 | 1.70E+07 | 2.45E+07 | 1.12E+07 | 2.33E+07 | 1.45E+07 | 1.46E+07 | 2.19E+07 | 1.63E+07 | 2.00E+07 | 1.70E+07 | 1.18 | 0.24 | 0.269518433 |
| Q06830 | Peroxiredoxin-1 OS=Homo sapiens OX=9606 GN=PRDX1 PE=1 SV=1 | 34 | 142 | 7 | 5.53E+07 | 5.20E+07 | 5.12E+07 | 4.76E+07 | 4.75E+07 | 4.87E+07 | 2.41E+07 | 2.74E+07 | 2.49E+07 | 2.41E+07 | 2.85E+07 | 1.68E+07 | 5.04E+07 | 2.43E+07 | 2.07 | 1.05 | 4.18809E-07 |
| O15230 | Laminin subunit alpha-5 OS=Homo sapiens OX=9606 GN=LAMA5 PE=1 SV=8 | 3 | 46 | 7 | 1.13E+07 | 5.86E+06 | 8.21E+06 | 8.33E+06 | 1.06E+07 | 8.31E+06 | 1.20E+07 | 9.05E+06 | 7.92E+06 | 9.33E+06 | 8.67E+06 | 6.30E+06 | 8.76E+06 | 8.87E+06 | 0.99 | -0.02 | 0.920905658 |
| O60488 | Long-chain-fatty-acid--CoA ligase 4 OS=Homo sapiens OX=9606 GN=ACSL4 PE=1 SV=2 | 16 | 58 | 7 | 2.51E+07 | 1.92E+07 | 1.17E+07 | 1.24E+07 | 1.43E+07 | 1.38E+07 | 1.22E+07 | 1.29E+07 | 8.17E+06 | 1.08E+07 | 1.00E+07 | 1.00E+07 | 1.61E+07 | 1.07E+07 | 1.50 | 0.59 | 0.050392678 |
| P14543 | Nidogen-1 OS=Homo sapiens OX=9606 GN=NID1 PE=1 SV=3 | 8 | 82 | 7 | 1.56E+07 | 1.78E+07 | 1.53E+07 | 1.50E+07 | 1.39E+07 | 1.35E+07 | 2.08E+07 | 2.22E+07 | 1.69E+07 | 1.96E+07 | 1.61E+07 | 1.46E+07 | 1.52E+07 | 1.84E+07 | 0.83 | -0.27 | 0.049726486 |
| P61313 | 60S ribosomal protein L15 OS=Homo sapiens OX=9606 GN=RPL15 PE=1 SV=2 | 31 | 137 | 7 | 6.73E+07 | 7.21E+07 | 7.68E+07 | 7.45E+07 | 6.81E+07 | 6.44E+07 | 4.58E+07 | 4.50E+07 | 4.35E+07 | 4.63E+07 | 3.94E+07 | 3.56E+07 | 7.05E+07 | 4.26E+07 | 1.66 | 0.73 | 8.53724E-07 |
| Q16666 | Gamma-interferon-inducible protein 16 OS=Homo sapiens OX=9606 GN=IFI16 PE=1 SV=3 | 10 | 73 | 7 | 1.92E+07 | 1.85E+07 | 1.95E+07 | 2.25E+07 | 2.14E+07 | 2.00E+07 | 9.15E+06 | 7.06E+06 | 8.27E+06 | 8.06E+06 | 8.27E+06 | 5.68E+06 | 2.02E+07 | 7.75E+06 | 2.60 | 1.38 | 3.29988E-08 |
| O75475 | PC4 and SFRS1-interacting protein OS=Homo sapiens OX=9606 GN=PSIP1 PE=1 SV=1 | 17 | 65 | 7 | 1.43E+07 | 1.48E+07 | 1.44E+07 | 1.50E+07 | 1.34E+07 | 1.39E+07 | 5.63E+06 | 5.20E+06 | 4.91E+06 | 4.04E+06 | 3.67E+06 | 5.50E+06 | 1.43E+07 | 4.82E+06 | 2.96 | 1.57 | 2.05195E-09 |
| P30101 | Protein disulfide-isomerase A3 OS=Homo sapiens OX=9606 GN=PDIA3 PE=1 SV=4 | 14 | 124 | 7 | 2.60E+07 | 2.43E+07 | 2.41E+07 | 2.47E+07 | 2.78E+07 | 2.38E+07 | 2.85E+07 | 2.99E+07 | 2.83E+07 | 2.65E+07 | 2.52E+07 | 2.38E+07 | 2.51E+07 | 2.71E+07 | 0.93 | -0.11 | 0.118676508 |
| P30041 | Peroxiredoxin-6 OS=Homo sapiens OX=9606 GN=PRDX6 PE=1 SV=3 | 38 | 71 | 7 | 1.63E+07 | 1.64E+07 | 1.75E+07 | 1.69E+07 | 1.76E+07 | 1.89E+07 | 1.59E+07 | 1.59E+07 | 1.45E+07 | 1.50E+07 | 1.45E+07 | 1.62E+07 | 1.72E+07 | 1.53E+07 | 1.13 | 0.17 | 0.003671198 |
| P39748 | Flap endonuclease 1 OS=Homo sapiens OX=9606 GN=FEN1 PE=1 SV=1 | 17 | 94 | 7 | 9.95E+07 | 1.00E+08 | 9.81E+07 | 1.04E+08 | 8.75E+07 | 9.39E+07 | 8.25E+07 | 8.12E+07 | 8.73E+07 | 8.85E+07 | 7.82E+07 | 6.97E+07 | 9.72E+07 | 8.12E+07 | 1.20 | 0.26 | 0.001486215 |
| O94776 | Metastasis-associated protein MTA2 OS=Homo sapiens OX=9606 GN=MTA2 PE=1 SV=1 | 12 | 28 | 7 | 1.29E+07 | 1.39E+07 | 1.22E+07 | 1.42E+07 | 1.08E+07 | 1.16E+07 | 9.80E+06 | 6.46E+06 | 8.68E+06 | 7.37E+06 | 5.13E+06 | 5.52E+06 | 1.26E+07 | 7.16E+06 | 1.76 | 0.82 | 0.00020902 |
| Q9NZN4 | EH domain-containing protein 2 OS=Homo sapiens OX=9606 GN=EHD2 PE=1 SV=2 | 18 | 132 | 7 | 1.48E+07 | 1.28E+07 | 1.26E+07 | 1.39E+07 | 1.23E+07 | 1.43E+07 | 1.46E+07 | 1.23E+07 | 1.21E+07 | 1.24E+07 | 1.17E+07 | 1.05E+07 | 1.34E+07 | 1.23E+07 | 1.10 | 0.13 | 0.119221272 |
| Q9Y223 | Bifunctional UDP-N-acetylglucosamine 2-epimerase/N-acetylmannosamine kinase OS=Homo sapiens OX=9606 GN=GNE PE=1 SV=1 | 12 | 68 | 7 | 1.28E+07 | 1.41E+07 | 1.11E+07 | 1.18E+07 | 1.22E+07 | 1.03E+07 | 1.41E+07 | 1.44E+07 | 1.24E+07 | 1.38E+07 | 1.17E+07 | 9.75E+06 | 1.21E+07 | 1.27E+07 | 0.95 | -0.07 | 0.503888253 |
| P27708 | CAD protein OS=Homo sapiens OX=9606 GN=CAD PE=1 SV=3 | 4 | 36 | 7 | 7.57E+06 | 9.99E+06 | 1.19E+07 | 1.31E+07 | 1.09E+07 | 1.13E+07 | 7.77E+06 | 8.09E+06 | 8.37E+06 | 6.71E+06 | 8.07E+06 | 7.63E+06 | 1.08E+07 | 7.77E+06 | 1.39 | 0.48 | 0.00991683 |
| O43143 | Pre-mRNA-splicing factor ATP-dependent RNA helicase DHX15 OS=Homo sapiens OX=9606 GN=DHX15 PE=1 SV=2 | 10 | 48 | 7 | 1.08E+07 | 1.34E+07 | 1.35E+07 | 1.71E+07 | 1.24E+07 | 1.30E+07 | 1.20E+07 | 1.27E+07 | 1.30E+07 | 1.19E+07 | 1.10E+07 | 7.52E+06 | 1.34E+07 | 1.13E+07 | 1.18 | 0.24 | 0.11746924 |
| Q9Y2A7 | Nck-associated protein 1 OS=Homo sapiens OX=9606 GN=NCKAP1 PE=1 SV=1 | 6 | 83 | 7 | 1.05E+07 | 1.16E+07 | 7.48E+06 | 1.18E+07 | 1.02E+07 | 8.36E+06 | 8.68E+06 | 7.82E+06 | 1.04E+07 | 9.92E+06 | 6.88E+06 | 6.63E+06 | 9.99E+06 | 8.39E+06 | 1.19 | 0.25 | 0.124727346 |
| P38117 | Electron transfer flavoprotein subunit beta OS=Homo sapiens OX=9606 GN=ETFB PE=1 SV=3 | 35 | 40 | 7 | 1.80E+07 | 1.83E+07 | 1.52E+07 | 1.77E+07 | 1.66E+07 | 1.85E+07 | 9.78E+06 | 9.01E+06 | 1.17E+07 | 1.12E+07 | 8.02E+06 | 6.54E+06 | 1.74E+07 | 9.38E+06 | 1.85 | 0.89 | 1.90347E-05 |
| P62081 | 40S ribosomal protein S7 OS=Homo sapiens OX=9606 GN=RPS7 PE=1 SV=1 | 46 | 60 | 7 | 2.19E+07 | 1.64E+07 | 1.60E+07 | 1.38E+07 | 1.41E+07 | 1.62E+07 | 1.08E+07 | 1.12E+07 | 8.22E+06 | 8.43E+06 | 9.74E+06 | 8.86E+06 | 1.64E+07 | 9.55E+06 | 1.72 | 0.78 | 0.00123988 |
| P43034 | Platelet-activating factor acetylhydrolase IB subunit alpha OS=Homo sapiens OX=9606 GN=PAFAH1B1 PE=1 SV=2 | 21 | 46 | 7 | 2.44E+07 | 2.48E+07 | 2.75E+07 | 2.65E+07 | 3.13E+07 | 2.50E+07 | 1.79E+07 | 1.93E+07 | 1.91E+07 | 1.68E+07 | 1.66E+07 | 1.69E+07 | 2.66E+07 | 1.77E+07 | 1.50 | 0.58 | 0.000117847 |
| P60709 | Actin, cytoplasmic 1 OS=Homo sapiens OX=9606 GN=ACTB PE=1 SV=1 | 83 | 4529 | 8 | 2.68E+09 | 2.75E+09 | 2.76E+09 | 2.73E+09 | 2.74E+09 | 2.87E+09 | 2.36E+09 | 2.40E+09 | 2.21E+09 | 2.45E+09 | 2.43E+09 | 2.28E+09 | 2.75E+09 | 2.35E+09 | 1.17 | 0.23 | 1.23776E-05 |
| P02533 | Keratin, type I cytoskeletal 14 OS=Homo sapiens OX=9606 GN=KRT14 PE=1 SV=4 | 45 | 534 | 8 | 5.76E+07 | 4.54E+07 | 1.69E+07 | 2.94E+07 | 2.68E+07 | 6.84E+07 | 8.94E+06 | 6.32E+07 | 3.50E+07 | 3.79E+07 | 3.44E+07 | 1.94E+07 | 4.08E+07 | 3.31E+07 | 1.23 | 0.30 | 0.505745948 |
| P27348 | 14-3-3 protein theta OS=Homo sapiens OX=9606 GN=YWHAQ PE=1 SV=1 | 42 | 159 | 8 | 4.20E+07 | 4.31E+07 | 5.30E+07 | 4.45E+07 | 4.81E+07 | 4.45E+07 | 4.50E+07 | 4.42E+07 | 4.62E+07 | 4.66E+07 | 4.69E+07 | 3.25E+07 | 4.59E+07 | 4.36E+07 | 1.05 | 0.07 | 0.428267914 |
| P01023 | Alpha-2-macroglobulin OS=Homo sapiens OX=9606 GN=A2M PE=1 SV=3 | 9 | 2864 | 8 | 9.99E+09 | 9.91E+09 | 1.19E+10 | 1.07E+10 | 1.12E+10 | 1.06E+10 | 9.80E+09 | 8.98E+09 | 1.09E+10 | 1.00E+10 | 1.13E+10 | 9.11E+09 | 1.07E+10 | 1.00E+10 | 1.07 | 0.10 | 0.180898824 |
| O94973 | AP-2 complex subunit alpha-2 OS=Homo sapiens OX=9606 GN=AP2A2 PE=1 SV=2 | 25 | 100 | 8 | 2.86E+07 | 3.25E+07 | 3.30E+07 | 3.31E+07 | 3.49E+07 | 3.88E+07 | 1.85E+07 | 2.06E+07 | 2.26E+07 | 1.49E+07 | 3.09E+07 | 1.96E+07 | 3.35E+07 | 2.12E+07 | 1.58 | 0.66 | 0.001271791 |
| P62937 | Peptidyl-prolyl cis-trans isomerase A OS=Homo sapiens OX=9606 GN=PPIA PE=1 SV=2 | 67 | 305 | 8 | 1.42E+08 | 1.43E+08 | 9.47E+07 | 9.63E+07 | 8.57E+07 | 1.16E+08 | 1.91E+08 | 1.44E+08 | 9.83E+07 | 1.02E+08 | 1.18E+08 | 1.20E+08 | 1.13E+08 | 1.29E+08 | 0.88 | -0.19 | 0.377781509 |
| Q15019 | Septin-2 OS=Homo sapiens OX=9606 GN=SEPTIN2 PE=1 SV=1 | 36 | 190 | 8 | 4.72E+07 | 4.46E+07 | 4.02E+07 | 3.99E+07 | 3.86E+07 | 3.69E+07 | 4.29E+07 | 3.89E+07 | 2.66E+07 | 2.86E+07 | 3.24E+07 | 3.18E+07 | 4.12E+07 | 3.35E+07 | 1.23 | 0.30 | 0.031995356 |
| Q9NQC3 | Reticulon-4 OS=Homo sapiens OX=9606 GN=RTN4 PE=1 SV=2 | 13 | 131 | 8 | 4.46E+07 | 4.44E+07 | 4.48E+07 | 4.97E+07 | 4.65E+07 | 4.10E+07 | 4.26E+07 | 4.08E+07 | 4.33E+07 | 5.24E+07 | 3.73E+07 | 3.51E+07 | 4.52E+07 | 4.19E+07 | 1.08 | 0.11 | 0.27020323 |
| P60174 | Triosephosphate isomerase OS=Homo sapiens OX=9606 GN=TPI1 PE=1 SV=3 | 40 | 213 | 8 | 6.58E+07 | 4.64E+07 | 4.64E+07 | 4.85E+07 | 4.24E+07 | 4.63E+07 | 6.22E+07 | 4.86E+07 | 4.31E+07 | 4.26E+07 | 4.30E+07 | 3.95E+07 | 4.93E+07 | 4.65E+07 | 1.06 | 0.08 | 0.567813307 |
| Q16401 | 26S proteasome non-ATPase regulatory subunit 5 OS=Homo sapiens OX=9606 GN=PSMD5 PE=1 SV=3 | 21 | 104 | 8 | 1.96E+07 | 1.58E+07 | 1.48E+07 | 1.93E+07 | 1.32E+07 | 1.60E+07 | 1.49E+07 | 1.26E+07 | 9.54E+06 | 7.63E+06 | 9.71E+06 | 1.13E+07 | 1.64E+07 | 1.09E+07 | 1.50 | 0.59 | 0.00393216 |
| P22626 | Heterogeneous nuclear ribonucleoproteins A2/B1 OS=Homo sapiens OX=9606 GN=HNRNPA2B1 PE=1 SV=2 | 27 | 122 | 8 | 3.88E+07 | 3.97E+07 | 3.34E+07 | 3.71E+07 | 3.07E+07 | 3.67E+07 | 2.36E+07 | 2.36E+07 | 2.04E+07 | 2.11E+07 | 2.04E+07 | 1.85E+07 | 3.61E+07 | 2.13E+07 | 1.70 | 0.76 | 1.46727E-05 |
| P67775 | Serine/threonine-protein phosphatase 2A catalytic subunit alpha isoform OS=Homo sapiens OX=9606 GN=PPP2CA PE=1 SV=1 | 44 | 140 | 8 | 5.45E+07 | 5.15E+07 | 4.01E+07 | 3.96E+07 | 4.75E+07 | 5.46E+07 | 5.02E+07 | 4.70E+07 | 3.50E+07 | 2.99E+07 | 3.59E+07 | 3.58E+07 | 4.80E+07 | 3.90E+07 | 1.23 | 0.30 | 0.060717614 |
| Q9NY33 | Dipeptidyl peptidase 3 OS=Homo sapiens OX=9606 GN=DPP3 PE=1 SV=2 | 17 | 117 | 8 | 1.95E+07 | 1.60E+07 | 1.80E+07 | 2.01E+07 | 1.67E+07 | 1.71E+07 | 2.43E+07 | 2.39E+07 | 2.49E+07 | 2.26E+07 | 2.22E+07 | 1.96E+07 | 1.79E+07 | 2.29E+07 | 0.78 | -0.36 | 0.000668799 |
| Q93063 | Exostosin-2 OS=Homo sapiens OX=9606 GN=EXT2 PE=1 SV=1 | 17 | 106 | 8 | 1.48E+07 | 1.64E+07 | 1.52E+07 | 1.61E+07 | 1.31E+07 | 1.67E+07 | 2.85E+07 | 2.63E+07 | 2.37E+07 | 2.21E+07 | 2.86E+07 | 2.55E+07 | 1.54E+07 | 2.58E+07 | 0.60 | -0.74 | 3.60702E-05 |
| P19823 | Inter-alpha-trypsin inhibitor heavy chain H2 OS=Homo sapiens OX=9606 GN=ITIH2 PE=1 SV=2 | 9 | 416 | 8 | 2.90E+08 | 3.31E+08 | 3.31E+08 | 3.32E+08 | 3.85E+08 | 2.87E+08 | 4.20E+08 | 4.60E+08 | 5.65E+08 | 4.51E+08 | 5.07E+08 | 3.40E+08 | 3.26E+08 | 4.57E+08 | 0.71 | -0.49 | 0.006559864 |
| P08582 | Melanotransferrin OS=Homo sapiens OX=9606 GN=MELTF PE=1 SV=2 | 18 | 115 | 8 | 2.63E+07 | 2.31E+07 | 2.64E+07 | 2.30E+07 | 2.08E+07 | 2.10E+07 | 1.47E+07 | 1.13E+07 | 1.31E+07 | 1.27E+07 | 1.18E+07 | 8.00E+06 | 2.34E+07 | 1.19E+07 | 1.97 | 0.98 | 7.58315E-06 |
| P17980 | 26S proteasome regulatory subunit 6A OS=Homo sapiens OX=9606 GN=PSMC3 PE=1 SV=3 | 22 | 141 | 8 | 3.58E+07 | 3.22E+07 | 3.17E+07 | 2.99E+07 | 2.39E+07 | 3.26E+07 | 2.60E+07 | 2.33E+07 | 1.82E+07 | 1.36E+07 | 1.86E+07 | 1.85E+07 | 3.10E+07 | 1.97E+07 | 1.57 | 0.65 | 0.00088522 |
| P01008 | Antithrombin-III OS=Homo sapiens OX=9606 GN=SERPINC1 PE=1 SV=1 | 21 | 736 | 8 | 5.84E+08 | 6.21E+08 | 7.02E+08 | 6.40E+08 | 6.61E+08 | 5.92E+08 | 7.60E+08 | 7.41E+08 | 8.97E+08 | 7.94E+08 | 8.18E+08 | 6.46E+08 | 6.33E+08 | 7.76E+08 | 0.82 | -0.29 | 0.006791831 |
| P01031 | Complement C5 OS=Homo sapiens OX=9606 GN=C5 PE=1 SV=4 | 5 | 223 | 8 | 1.34E+08 | 1.42E+08 | 1.28E+08 | 1.39E+08 | 1.40E+08 | 1.25E+08 | 1.87E+08 | 1.87E+08 | 1.88E+08 | 1.88E+08 | 1.76E+08 | 1.64E+08 | 1.35E+08 | 1.81E+08 | 0.74 | -0.43 | 5.50429E-06 |
| P50570 | Dynamin-2 OS=Homo sapiens OX=9606 GN=DNM2 PE=1 SV=2 | 15 | 110 | 8 | 2.89E+07 | 3.21E+07 | 2.77E+07 | 2.75E+07 | 2.56E+07 | 3.04E+07 | 2.34E+07 | 2.36E+07 | 2.28E+07 | 1.57E+07 | 2.06E+07 | 1.83E+07 | 2.87E+07 | 2.07E+07 | 1.38 | 0.47 | 0.000773146 |
| P08621 | U1 small nuclear ribonucleoprotein 70 kDa OS=Homo sapiens OX=9606 GN=SNRNP70 PE=1 SV=2 | 20 | 193 | 8 | 4.25E+07 | 4.27E+07 | 4.38E+07 | 4.73E+07 | 4.07E+07 | 4.33E+07 | 3.46E+07 | 3.38E+07 | 3.31E+07 | 3.42E+07 | 3.05E+07 | 2.70E+07 | 4.34E+07 | 3.22E+07 | 1.35 | 0.43 | 3.25549E-05 |
| Q15907 | Ras-related protein Rab-11B OS=Homo sapiens OX=9606 GN=RAB11B PE=1 SV=4 | 42 | 130 | 8 | 6.31E+07 | 6.65E+07 | 4.77E+07 | 5.90E+07 | 5.91E+07 | 5.52E+07 | 5.24E+07 | 5.69E+07 | 4.94E+07 | 5.08E+07 | 4.93E+07 | 4.88E+07 | 5.84E+07 | 5.12E+07 | 1.14 | 0.19 | 0.044612654 |
| P61353 | 60S ribosomal protein L27 OS=Homo sapiens OX=9606 GN=RPL27 PE=1 SV=2 | 49 | 241 | 8 | 1.17E+08 | 1.19E+08 | 1.41E+08 | 1.33E+08 | 1.26E+08 | 1.38E+08 | 8.89E+07 | 9.29E+07 | 9.13E+07 | 9.59E+07 | 9.12E+07 | 7.79E+07 | 1.29E+08 | 8.97E+07 | 1.44 | 0.53 | 2.64042E-05 |
| P06756 | Integrin alpha-V OS=Homo sapiens OX=9606 GN=ITGAV PE=1 SV=2 | 12 | 64 | 8 | 1.48E+07 | 1.58E+07 | 1.41E+07 | 1.35E+07 | 1.47E+07 | 1.48E+07 | 9.80E+06 | 6.18E+06 | 9.05E+06 | 7.78E+06 | 8.31E+06 | 9.03E+06 | 1.46E+07 | 8.36E+06 | 1.75 | 0.81 | 5.26082E-06 |
| P11047 | Laminin subunit gamma-1 OS=Homo sapiens OX=9606 GN=LAMC1 PE=1 SV=3 | 7 | 107 | 8 | 1.60E+07 | 9.65E+06 | 1.17E+07 | 1.32E+07 | 1.12E+07 | 1.21E+07 | 1.95E+07 | 1.80E+07 | 1.57E+07 | 1.80E+07 | 1.56E+07 | 1.55E+07 | 1.23E+07 | 1.71E+07 | 0.72 | -0.47 | 0.001789035 |
| P38646 | Stress-70 protein, mitochondrial OS=Homo sapiens OX=9606 GN=HSPA9 PE=1 SV=2 | 15 | 56 | 8 | 1.91E+07 | 1.80E+07 | 1.11E+07 | 1.15E+07 | 1.23E+07 | 1.80E+07 | 8.14E+06 | 1.13E+07 | 8.76E+06 | 6.02E+06 | 7.88E+06 | 7.42E+06 | 1.50E+07 | 8.25E+06 | 1.82 | 0.86 | 0.004856812 |
| Q92900 | Regulator of nonsense transcripts 1 OS=Homo sapiens OX=9606 GN=UPF1 PE=1 SV=2 | 11 | 49 | 8 | 1.85E+07 | 1.64E+07 | 2.09E+07 | 1.73E+07 | 1.77E+07 | 1.46E+07 | 1.21E+07 | 1.23E+07 | 1.35E+07 | 1.00E+07 | 1.23E+07 | 1.60E+07 | 1.75E+07 | 1.27E+07 | 1.38 | 0.47 | 0.002070798 |
| Q92945 | Far upstream element-binding protein 2 OS=Homo sapiens OX=9606 GN=KHSRP PE=1 SV=4 | 18 | 102 | 8 | 2.83E+07 | 2.78E+07 | 2.63E+07 | 3.12E+07 | 2.36E+07 | 2.51E+07 | 1.56E+07 | 1.84E+07 | 1.62E+07 | 1.42E+07 | 1.48E+07 | 1.47E+07 | 2.71E+07 | 1.56E+07 | 1.73 | 0.79 | 1.74522E-05 |
| P17301 | Integrin alpha-2 OS=Homo sapiens OX=9606 GN=ITGA2 PE=1 SV=1 | 9 | 64 | 8 | 1.90E+07 | 1.61E+07 | 1.58E+07 | 1.72E+07 | 1.43E+07 | 1.47E+07 | 8.75E+06 | 1.09E+07 | 9.42E+06 | 8.93E+06 | 1.06E+07 | 7.97E+06 | 1.62E+07 | 9.42E+06 | 1.72 | 0.78 | 2.91955E-05 |
| P11766 | Alcohol dehydrogenase class-3 OS=Homo sapiens OX=9606 GN=ADH5 PE=1 SV=4 | 24 | 187 | 8 | 3.74E+07 | 3.81E+07 | 3.78E+07 | 4.07E+07 | 3.21E+07 | 3.87E+07 | 5.01E+07 | 5.70E+07 | 3.78E+07 | 4.01E+07 | 4.94E+07 | 3.49E+07 | 3.75E+07 | 4.49E+07 | 0.83 | -0.26 | 0.089698419 |
| P62917 | 60S ribosomal protein L8 OS=Homo sapiens OX=9606 GN=RPL8 PE=1 SV=2 | 23 | 154 | 8 | 4.87E+07 | 4.96E+07 | 5.11E+07 | 5.71E+07 | 5.16E+07 | 4.62E+07 | 3.38E+07 | 3.29E+07 | 3.32E+07 | 3.65E+07 | 2.81E+07 | 2.50E+07 | 5.07E+07 | 3.16E+07 | 1.61 | 0.68 | 8.62458E-06 |
| Q13263 | Transcription intermediary factor 1-beta OS=Homo sapiens OX=9606 GN=TRIM28 PE=1 SV=5 | 19 | 43 | 8 | 1.53E+07 | 1.62E+07 | 1.23E+07 | 1.24E+07 | 1.73E+07 | 1.43E+07 | 6.57E+06 | 5.53E+06 | 5.32E+06 | 8.10E+06 | 6.01E+06 | 4.43E+06 | 1.46E+07 | 5.99E+06 | 2.44 | 1.29 | 1.59513E-05 |
| Q13838 | Spliceosome RNA helicase DDX39B OS=Homo sapiens OX=9606 GN=DDX39B PE=1 SV=1 | 21 | 157 | 8 | 1.97E+07 | 2.60E+07 | 2.28E+07 | 2.15E+07 | 2.24E+07 | 2.26E+07 | 1.85E+07 | 2.16E+07 | 1.89E+07 | 1.64E+07 | 1.54E+07 | 1.24E+07 | 2.25E+07 | 1.72E+07 | 1.31 | 0.39 | 0.007938427 |
| P61221 | ATP-binding cassette sub-family E member 1 OS=Homo sapiens OX=9606 GN=ABCE1 PE=1 SV=1 | 23 | 84 | 8 | 1.78E+07 | 2.09E+07 | 2.38E+07 | 2.07E+07 | 1.96E+07 | 1.73E+07 | 1.47E+07 | 1.20E+07 | 1.63E+07 | 1.51E+07 | 1.58E+07 | 1.37E+07 | 2.00E+07 | 1.46E+07 | 1.37 | 0.45 | 0.001276361 |
| P21281 | V-type proton ATPase subunit B, brain isoform OS=Homo sapiens OX=9606 GN=ATP6V1B2 PE=1 SV=3 | 21 | 54 | 8 | 1.42E+07 | 1.28E+07 | 1.34E+07 | 1.25E+07 | 1.28E+07 | 1.31E+07 | 1.26E+07 | 1.37E+07 | 1.05E+07 | 9.22E+06 | 9.38E+06 | 1.03E+07 | 1.31E+07 | 1.09E+07 | 1.20 | 0.26 | 0.029433453 |
| P15144 | Aminopeptidase N OS=Homo sapiens OX=9606 GN=ANPEP PE=1 SV=4 | 12 | 40 | 8 | 8.50E+06 | 1.12E+07 | 1.14E+07 | 8.86E+06 | 9.72E+06 | 9.20E+06 | 3.79E+06 | 4.35E+06 | 4.98E+06 | 3.39E+06 | 4.84E+06 | 6.03E+06 | 9.80E+06 | 4.56E+06 | 2.15 | 1.10 | 1.11135E-05 |
| Q02809 | Procollagen-lysine,2-oxoglutarate 5-dioxygenase 1 OS=Homo sapiens OX=9606 GN=PLOD1 PE=1 SV=2 | 13 | 67 | 8 | 1.61E+07 | 1.18E+07 | 1.21E+07 | 1.15E+07 | 1.10E+07 | 1.09E+07 | 1.40E+07 | 1.53E+07 | 1.80E+07 | 1.55E+07 | 1.22E+07 | 1.23E+07 | 1.22E+07 | 1.46E+07 | 0.84 | -0.25 | 0.08155761 |
| P33992 | DNA replication licensing factor MCM5 OS=Homo sapiens OX=9606 GN=MCM5 PE=1 SV=5 | 14 | 76 | 8 | 2.52E+07 | 2.60E+07 | 2.12E+07 | 2.32E+07 | 2.15E+07 | 2.08E+07 | 1.51E+07 | 1.57E+07 | 1.37E+07 | 1.44E+07 | 1.21E+07 | 1.12E+07 | 2.30E+07 | 1.37E+07 | 1.68 | 0.75 | 1.3824E-05 |
| Q12860 | Contactin-1 OS=Homo sapiens OX=9606 GN=CNTN1 PE=1 SV=1 | 10 | 62 | 8 | 1.96E+07 | 1.30E+07 | 1.38E+07 | 1.95E+07 | 1.11E+07 | 1.44E+07 | 1.62E+07 | 1.62E+07 | 1.40E+07 | 1.75E+07 | 1.80E+07 | 1.19E+07 | 1.52E+07 | 1.56E+07 | 0.97 | -0.04 | 0.813448145 |
| Q01082 | Spectrin beta chain, non-erythrocytic 1 OS=Homo sapiens OX=9606 GN=SPTBN1 PE=1 SV=2 | 5 | 43 | 8 | 1.10E+07 | 1.06E+07 | 1.03E+07 | 1.11E+07 | 1.14E+07 | 1.20E+07 | 7.27E+06 | 7.06E+06 | 8.15E+06 | 6.19E+06 | 6.53E+06 | 6.95E+06 | 1.11E+07 | 7.02E+06 | 1.58 | 0.66 | 6.96185E-07 |
| Q92973 | Transportin-1 OS=Homo sapiens OX=9606 GN=TNPO1 PE=1 SV=2 | 13 | 57 | 8 | 1.85E+07 | 1.73E+07 | 1.80E+07 | 1.85E+07 | 1.64E+07 | 1.74E+07 | 1.25E+07 | 1.20E+07 | 1.13E+07 | 1.10E+07 | 1.07E+07 | 8.06E+06 | 1.77E+07 | 1.09E+07 | 1.62 | 0.69 | 1.83323E-05 |
| P54577 | Tyrosine--tRNA ligase, cytoplasmic OS=Homo sapiens OX=9606 GN=YARS PE=1 SV=4 | 18 | 52 | 8 | 1.21E+07 | 1.23E+07 | 1.35E+07 | 1.43E+07 | 1.49E+07 | 1.32E+07 | 8.28E+06 | 9.78E+06 | 9.88E+06 | 7.45E+06 | 6.91E+06 | 1.61E+07 | 1.34E+07 | 9.73E+06 | 1.38 | 0.46 | 0.042814489 |
| O00203 | AP-3 complex subunit beta-1 OS=Homo sapiens OX=9606 GN=AP3B1 PE=1 SV=3 | 10 | 47 | 8 | 1.11E+08 | 1.18E+08 | 5.69E+07 | 4.55E+07 | 8.81E+07 | 5.21E+07 | 4.64E+07 | 4.90E+07 | 2.88E+07 | 5.36E+07 | 4.81E+07 | 4.17E+07 | 7.85E+07 | 4.46E+07 | 1.76 | 0.81 | 0.045268859 |
| Q93008 | Probable ubiquitin carboxyl-terminal hydrolase FAF-X OS=Homo sapiens OX=9606 GN=USP9X PE=1 SV=3 | 3 | 26 | 8 | 8.62E+06 | 9.68E+06 | 1.05E+07 | 1.13E+07 | 9.99E+06 | 8.78E+06 | 7.69E+06 | 8.59E+06 | 8.72E+06 | 9.61E+06 | 7.29E+06 | 6.43E+06 | 9.81E+06 | 8.06E+06 | 1.22 | 0.28 | 0.019114085 |
| P46459 | Vesicle-fusing ATPase OS=Homo sapiens OX=9606 GN=NSF PE=1 SV=3 | 13 | 55 | 8 | 1.56E+07 | 1.59E+07 | 1.58E+07 | 1.50E+07 | 1.34E+07 | 1.30E+07 | 1.23E+07 | 1.08E+07 | 1.24E+07 | 1.17E+07 | 1.12E+07 | 1.00E+07 | 1.48E+07 | 1.14E+07 | 1.29 | 0.37 | 0.000463665 |
| P00734 | Prothrombin OS=Homo sapiens OX=9606 GN=F2 PE=1 SV=2 | 11 | 3066 | 9 | 8.01E+08 | 7.08E+08 | 8.31E+08 | 8.31E+08 | 9.22E+08 | 7.84E+08 | 1.02E+09 | 1.03E+09 | 1.12E+09 | 1.21E+09 | 1.16E+09 | 1.01E+09 | 8.13E+08 | 1.09E+09 | 0.74 | -0.43 | 0.000100157 |
| P50395 | Rab GDP dissociation inhibitor beta OS=Homo sapiens OX=9606 GN=GDI2 PE=1 SV=2 | 40 | 386 | 9 | 1.20E+08 | 8.70E+07 | 9.20E+07 | 9.64E+07 | 9.06E+07 | 1.00E+08 | 1.47E+08 | 1.20E+08 | 1.10E+08 | 8.74E+07 | 1.29E+08 | 1.11E+08 | 9.77E+07 | 1.17E+08 | 0.83 | -0.26 | 0.073244611 |
| O00560 | Syntenin-1 OS=Homo sapiens OX=9606 GN=SDCBP PE=1 SV=1 | 59 | 216 | 9 | 7.15E+07 | 6.29E+07 | 5.89E+07 | 5.89E+07 | 6.12E+07 | 5.68E+07 | 7.19E+07 | 6.80E+07 | 6.23E+07 | 5.64E+07 | 6.03E+07 | 5.38E+07 | 6.17E+07 | 6.21E+07 | 0.99 | -0.01 | 0.907255649 |
| P13647 | Keratin, type II cytoskeletal 5 OS=Homo sapiens OX=9606 GN=KRT5 PE=1 SV=3 | 28 | 498 | 9 | 6.50E+07 | 7.00E+07 | 2.60E+07 | 4.25E+07 | 4.26E+07 | 5.87E+07 | 1.93E+07 | 5.90E+07 | 3.99E+07 | 3.70E+07 | 4.14E+07 | 2.92E+07 | 5.08E+07 | 3.76E+07 | 1.35 | 0.43 | 0.162039929 |
| P11940 | Polyadenylate-binding protein 1 OS=Homo sapiens OX=9606 GN=PABPC1 PE=1 SV=2 | 24 | 228 | 9 | 7.64E+07 | 7.53E+07 | 6.74E+07 | 6.82E+07 | 7.07E+07 | 7.41E+07 | 5.39E+07 | 5.54E+07 | 4.58E+07 | 4.45E+07 | 4.53E+07 | 4.09E+07 | 7.20E+07 | 4.76E+07 | 1.51 | 0.60 | 1.42994E-05 |
| P61160 | Actin-related protein 2 OS=Homo sapiens OX=9606 GN=ACTR2 PE=1 SV=1 | 35 | 78 | 9 | 2.17E+07 | 2.46E+07 | 2.43E+07 | 2.52E+07 | 2.43E+07 | 2.45E+07 | 2.18E+07 | 2.68E+07 | 1.81E+07 | 2.21E+07 | 2.05E+07 | 1.91E+07 | 2.41E+07 | 2.14E+07 | 1.13 | 0.17 | 0.085349431 |
| P05388 | 60S acidic ribosomal protein P0 OS=Homo sapiens OX=9606 GN=RPLP0 PE=1 SV=1 | 42 | 160 | 9 | 4.09E+07 | 4.27E+07 | 2.96E+07 | 3.50E+07 | 3.67E+07 | 3.80E+07 | 4.53E+07 | 4.47E+07 | 3.39E+07 | 3.27E+07 | 3.67E+07 | 3.21E+07 | 3.72E+07 | 3.76E+07 | 0.99 | -0.02 | 0.891558759 |
| O00232 | 26S proteasome non-ATPase regulatory subunit 12 OS=Homo sapiens OX=9606 GN=PSMD12 PE=1 SV=3 | 22 | 136 | 9 | 3.36E+07 | 3.44E+07 | 4.02E+07 | 4.22E+07 | 4.25E+07 | 3.70E+07 | 2.79E+07 | 2.53E+07 | 2.65E+07 | 2.78E+07 | 2.91E+07 | 2.37E+07 | 3.83E+07 | 2.67E+07 | 1.43 | 0.52 | 0.00026546 |
| P09211 | Glutathione S-transferase P OS=Homo sapiens OX=9606 GN=GSTP1 PE=1 SV=2 | 57 | 171 | 9 | 5.80E+07 | 6.38E+07 | 5.36E+07 | 5.53E+07 | 5.47E+07 | 5.06E+07 | 6.44E+07 | 6.33E+07 | 5.61E+07 | 6.47E+07 | 5.87E+07 | 6.16E+07 | 5.60E+07 | 6.14E+07 | 0.91 | -0.13 | 0.041210727 |
| Q99988 | Growth/differentiation factor 15 OS=Homo sapiens OX=9606 GN=GDF15 PE=1 SV=3 | 40 | 198 | 9 | 5.54E+07 | 5.52E+07 | 6.27E+07 | 5.68E+07 | 6.13E+07 | 5.23E+07 | 6.05E+07 | 6.11E+07 | 4.94E+07 | 5.86E+07 | 4.89E+07 | 4.73E+07 | 5.73E+07 | 5.43E+07 | 1.05 | 0.08 | 0.360980918 |
| P62191 | 26S proteasome regulatory subunit 4 OS=Homo sapiens OX=9606 GN=PSMC1 PE=1 SV=1 | 26 | 163 | 9 | 7.57E+07 | 7.45E+07 | 7.12E+07 | 6.68E+07 | 7.14E+07 | 6.96E+07 | 4.97E+07 | 5.61E+07 | 4.22E+07 | 4.90E+07 | 4.20E+07 | 3.87E+07 | 7.16E+07 | 4.63E+07 | 1.55 | 0.63 | 4.23889E-05 |
| P10768 | S-formylglutathione hydrolase OS=Homo sapiens OX=9606 GN=ESD PE=1 SV=2 | 50 | 155 | 9 | 5.12E+07 | 4.97E+07 | 4.80E+07 | 4.15E+07 | 4.51E+07 | 4.50E+07 | 4.28E+07 | 3.80E+07 | 4.14E+07 | 4.88E+07 | 4.04E+07 | 4.06E+07 | 4.68E+07 | 4.20E+07 | 1.11 | 0.15 | 0.046050263 |
| O14818 | Proteasome subunit alpha type-7 OS=Homo sapiens OX=9606 GN=PSMA7 PE=1 SV=1 | 40 | 174 | 9 | 5.27E+07 | 6.95E+07 | 1.15E+08 | 4.62E+07 | 4.56E+07 | 5.54E+07 | 5.14E+07 | 6.65E+07 | 4.57E+07 | 4.49E+07 | 3.49E+07 | 2.01E+08 | 6.41E+07 | 7.40E+07 | 0.87 | -0.21 | 0.73465334 |
| P09543 | 2',3'-cyclic-nucleotide 3'-phosphodiesterase OS=Homo sapiens OX=9606 GN=CNP PE=1 SV=2 | 29 | 81 | 9 | 1.56E+07 | 1.50E+07 | 1.73E+07 | 1.55E+07 | 1.57E+07 | 1.56E+07 | 8.02E+06 | 1.11E+07 | 8.51E+06 | 1.13E+07 | 1.04E+07 | 6.98E+06 | 1.58E+07 | 9.38E+06 | 1.68 | 0.75 | 9.51072E-05 |
| P07737 | Profilin-1 OS=Homo sapiens OX=9606 GN=PFN1 PE=1 SV=2 | 71 | 775 | 9 | 1.12E+08 | 1.04E+08 | 1.92E+08 | 1.32E+08 | 1.78E+08 | 1.20E+08 | 9.25E+07 | 7.79E+07 | 1.96E+08 | 1.28E+08 | 2.05E+08 | 1.16E+08 | 1.40E+08 | 1.36E+08 | 1.03 | 0.04 | 0.891070543 |
| Q13418 | Integrin-linked protein kinase OS=Homo sapiens OX=9606 GN=ILK PE=1 SV=2 | 28 | 188 | 9 | 3.53E+07 | 3.83E+07 | 3.05E+07 | 3.66E+07 | 3.10E+07 | 3.63E+07 | 4.58E+07 | 4.37E+07 | 3.67E+07 | 3.91E+07 | 3.37E+07 | 3.54E+07 | 3.47E+07 | 3.91E+07 | 0.89 | -0.17 | 0.094881845 |
| O15067 | Phosphoribosylformylglycinamidine synthase OS=Homo sapiens OX=9606 GN=PFAS PE=1 SV=4 | 9 | 102 | 9 | 6.49E+07 | 6.28E+07 | 4.35E+07 | 5.14E+07 | 4.44E+07 | 5.66E+07 | 6.76E+07 | 8.71E+07 | 6.03E+07 | 5.62E+07 | 6.16E+07 | 6.54E+07 | 5.39E+07 | 6.64E+07 | 0.81 | -0.30 | 0.058764056 |
| O76094 | Signal recognition particle subunit SRP72 OS=Homo sapiens OX=9606 GN=SRP72 PE=1 SV=3 | 20 | 92 | 9 | 2.78E+07 | 2.75E+07 | 2.25E+07 | 2.85E+07 | 2.54E+07 | 2.73E+07 | 1.40E+07 | 1.18E+07 | 1.23E+07 | 1.33E+07 | 1.13E+07 | 1.18E+07 | 2.65E+07 | 1.24E+07 | 2.14 | 1.09 | 2.15844E-06 |
| P17655 | Calpain-2 catalytic subunit OS=Homo sapiens OX=9606 GN=CAPN2 PE=1 SV=6 | 22 | 56 | 9 | 1.37E+07 | 1.52E+07 | 1.36E+07 | 9.32E+06 | 1.69E+07 | 1.14E+07 | 1.65E+07 | 1.08E+07 | 6.80E+06 | 1.30E+07 | 1.13E+07 | 7.92E+06 | 1.34E+07 | 1.11E+07 | 1.21 | 0.27 | 0.234843222 |
| P02452 | Collagen alpha-1(I) chain OS=Homo sapiens OX=9606 GN=COL1A1 PE=1 SV=5 | 9 | 26 | 9 | 1.15E+06 | 8.39E+05 | 6.36E+05 | 1.57E+06 | 7.73E+05 | 1.21E+06 | 5.38E+06 | 3.71E+06 | 2.86E+06 | 2.34E+06 | 2.32E+06 | 3.00E+07 | 1.03E+06 | 7.77E+06 | 0.13 | -2.92 | 0.192156695 |
| Q07020 | 60S ribosomal protein L18 OS=Homo sapiens OX=9606 GN=RPL18 PE=1 SV=2 | 39 | 347 | 9 | 1.49E+08 | 1.57E+08 | 1.34E+08 | 1.46E+08 | 1.44E+08 | 1.46E+08 | 1.10E+08 | 1.13E+08 | 9.49E+07 | 1.08E+08 | 9.83E+07 | 9.17E+07 | 1.46E+08 | 1.03E+08 | 1.42 | 0.51 | 4.32067E-06 |
| P49189 | 4-trimethylaminobutyraldehyde dehydrogenase OS=Homo sapiens OX=9606 GN=ALDH9A1 PE=1 SV=3 | 23 | 337 | 9 | 1.34E+08 | 1.46E+08 | 1.29E+08 | 1.70E+08 | 1.62E+08 | 1.60E+08 | 1.80E+08 | 1.97E+08 | 2.11E+08 | 2.14E+08 | 1.86E+08 | 1.62E+08 | 1.50E+08 | 1.92E+08 | 0.78 | -0.35 | 0.002838842 |
| P02768 | Serum albumin OS=Homo sapiens OX=9606 GN=ALB PE=1 SV=2 | 13 | 1326 | 9 | 1.76E+09 | 1.81E+09 | 1.82E+09 | 1.90E+09 | 2.12E+09 | 2.05E+09 | 3.15E+09 | 3.12E+09 | 3.84E+09 | 3.51E+09 | 3.44E+09 | 2.80E+09 | 1.91E+09 | 3.31E+09 | 0.58 | -0.79 | 7.63559E-05 |
| Q16658 | Fascin OS=Homo sapiens OX=9606 GN=FSCN1 PE=1 SV=3 | 27 | 96 | 9 | 1.95E+07 | 1.80E+07 | 1.85E+07 | 1.63E+07 | 1.52E+07 | 1.82E+07 | 1.70E+07 | 1.88E+07 | 1.68E+07 | 1.71E+07 | 1.50E+07 | 1.77E+07 | 1.76E+07 | 1.71E+07 | 1.03 | 0.04 | 0.531155237 |
| P51149 | Ras-related protein Rab-7a OS=Homo sapiens OX=9606 GN=RAB7A PE=1 SV=1 | 52 | 225 | 9 | 6.92E+07 | 6.53E+07 | 7.49E+07 | 5.76E+07 | 6.23E+07 | 6.36E+07 | 6.34E+07 | 6.12E+07 | 5.95E+07 | 5.41E+07 | 5.62E+07 | 4.97E+07 | 6.55E+07 | 5.73E+07 | 1.14 | 0.19 | 0.029785659 |
| O43795 | Unconventional myosin-Ib OS=Homo sapiens OX=9606 GN=MYO1B PE=1 SV=3 | 10 | 75 | 9 | 1.34E+07 | 1.30E+07 | 1.55E+07 | 1.61E+07 | 1.35E+07 | 1.31E+07 | 9.80E+06 | 8.86E+06 | 8.47E+06 | 1.03E+07 | 6.56E+06 | 5.13E+06 | 1.41E+07 | 8.19E+06 | 1.72 | 0.78 | 0.000209614 |
| Q7KZF4 | Staphylococcal nuclease domain-containing protein 1 OS=Homo sapiens OX=9606 GN=SND1 PE=1 SV=1 | 15 | 90 | 9 | 2.32E+07 | 2.47E+07 | 2.12E+07 | 2.23E+07 | 2.16E+07 | 2.33E+07 | 1.54E+07 | 1.59E+07 | 1.62E+07 | 1.59E+07 | 1.45E+07 | 1.13E+07 | 2.27E+07 | 1.49E+07 | 1.53 | 0.61 | 1.44295E-05 |
| P07741 | Adenine phosphoribosyltransferase OS=Homo sapiens OX=9606 GN=APRT PE=1 SV=2 | 54 | 158 | 9 | 4.91E+07 | 4.39E+07 | 3.86E+07 | 3.15E+07 | 3.25E+07 | 3.61E+07 | 3.46E+07 | 3.38E+07 | 3.16E+07 | 2.59E+07 | 2.90E+07 | 2.81E+07 | 3.86E+07 | 3.05E+07 | 1.27 | 0.34 | 0.033032681 |
| P17812 | CTP synthase 1 OS=Homo sapiens OX=9606 GN=CTPS1 PE=1 SV=2 | 19 | 62 | 9 | 1.62E+07 | 1.66E+07 | 1.96E+07 | 1.31E+07 | 1.98E+07 | 1.34E+07 | 1.50E+07 | 1.19E+07 | 8.01E+06 | 9.70E+06 | 1.13E+07 | 1.25E+07 | 1.65E+07 | 1.14E+07 | 1.44 | 0.53 | 0.008538244 |
| P62277 | 40S ribosomal protein S13 OS=Homo sapiens OX=9606 GN=RPS13 PE=1 SV=2 | 44 | 178 | 9 | 6.04E+07 | 5.87E+07 | 5.66E+07 | 5.97E+07 | 5.51E+07 | 5.56E+07 | 3.66E+07 | 4.07E+07 | 3.28E+07 | 3.30E+07 | 3.67E+07 | 4.37E+07 | 5.77E+07 | 3.73E+07 | 1.55 | 0.63 | 1.07631E-05 |
| O60832 | H/ACA ribonucleoprotein complex subunit DKC1 OS=Homo sapiens OX=9606 GN=DKC1 PE=1 SV=3 | 29 | 68 | 9 | 3.95E+07 | 3.06E+07 | 2.10E+07 | 1.87E+07 | 2.39E+07 | 2.60E+07 | 1.74E+07 | 2.41E+07 | 1.07E+07 | 1.10E+07 | 9.26E+06 | 1.50E+07 | 2.66E+07 | 1.46E+07 | 1.82 | 0.87 | 0.011629865 |
| P08648 | Integrin alpha-5 OS=Homo sapiens OX=9606 GN=ITGA5 PE=1 SV=2 | 12 | 79 | 9 | 3.38E+07 | 3.04E+07 | 2.66E+07 | 2.68E+07 | 3.43E+07 | 2.31E+07 | 1.25E+07 | 9.50E+06 | 1.13E+07 | 1.26E+07 | 1.10E+07 | 1.04E+07 | 2.92E+07 | 1.12E+07 | 2.60 | 1.38 | 9.5166E-05 |

### Supplementary Table 4. List of all proteins identified by label free proteomics analysis.

| Accession | Description | Coverage [%] | # PSMs | # Unique Peptides | Amitriptyline 1 | Amitriptyline 2 | Amitriptyline 3 | Amitriptyline 4 | Amitriptyline 5 | Amitriptyline 6 | Vehicle 1 | Vehicle 2 | Vehicle 3 | Vehicle 4 | Vehicle 5 | Vehicle 6 | Average amitriptyline | Average vehicle | Ratio amitriptyline/ vehicle | Log 2 fold change (amitriptyline /vehicle) | P-value |
|---|---|---|---|---|---|---|---|---|---|---|---|---|---|---|---|---|---|---|---|---|---|
| Q9NSD9 | Phenylalanine--tRNA ligase beta subunit OS=Homo sapiens OX=9606 GN=FARSB PE=1 SV=3 | 18 | 72 | 9 | 1.66E+07 | 1.58E+07 | 1.80E+07 | 2.25E+07 | 1.52E+07 | 2.04E+07 | 1.42E+07 | 1.35E+07 | 1.15E+07 | 1.41E+07 | 1.32E+07 | 1.37E+07 | 1.81E+07 | 1.34E+07 | 1.35 | 0.44 | 0.008214467 |
| Q04637 | Eukaryotic translation initiation factor 4 gamma 1 OS=Homo sapiens OX=9606 GN=EIF4G1 PE=1 SV=4 | 8 | 76 | 9 | 2.43E+07 | 2.43E+07 | 2.31E+07 | 2.50E+07 | 2.23E+07 | 2.11E+07 | 1.08E+07 | 1.20E+07 | 1.09E+07 | 1.32E+07 | 1.07E+07 | 9.81E+06 | 2.33E+07 | 1.13E+07 | 2.07 | 1.05 | 4.12192E-08 |
| Q07955 | Serine/arginine-rich splicing factor 1 OS=Homo sapiens OX=9606 GN=SRSF1 PE=1 SV=2 | 35 | 68 | 9 | 1.62E+07 | 1.55E+07 | 1.62E+07 | 1.71E+07 | 1.48E+07 | 1.57E+07 | 1.52E+07 | 1.58E+07 | 1.45E+07 | 1.40E+07 | 1.29E+07 | 1.32E+07 | 1.59E+07 | 1.43E+07 | 1.12 | 0.16 | 0.016194833 |
| Q9Y6G9 | Cytoplasmic dynein 1 light intermediate chain 1 OS=Homo sapiens OX=9606 GN=DYNC1LI1 PE=1 SV=3 | 22 | 47 | 9 | 8.72E+07 | 8.26E+07 | 8.82E+07 | 8.81E+07 | 9.80E+07 | 8.93E+07 | 9.47E+07 | 1.11E+08 | 9.29E+07 | 1.12E+08 | 9.19E+07 | 7.83E+07 | 8.89E+07 | 9.68E+07 | 0.92 | -0.12 | 0.205676263 |
| P04439 | HLA class I histocompatibility antigen, A alpha chain OS=Homo sapiens OX=9606 GN=HLA-A PE=1 SV=2 | 52 | 729 | 10 | 3.40E+08 | 3.67E+08 | 3.35E+08 | 3.67E+08 | 3.49E+08 | 3.52E+08 | 2.03E+08 | 2.21E+08 | 1.95E+08 | 2.14E+08 | 1.88E+08 | 1.74E+08 | 3.52E+08 | 1.99E+08 | 1.77 | 0.82 | 2.07833E-08 |
| P63104 | 14-3-3 protein zeta/delta OS=Homo sapiens OX=9606 GN=YWHAZ PE=1 SV=1 | 53 | 210 | 10 | 4.59E+07 | 3.89E+07 | 4.21E+07 | 5.23E+07 | 4.67E+07 | 4.03E+07 | 4.65E+07 | 4.39E+07 | 4.42E+07 | 4.25E+07 | 3.57E+07 | 3.66E+07 | 4.43E+07 | 4.16E+07 | 1.07 | 0.09 | 0.326383081 |
| O43707 | Alpha-actinin-4 OS=Homo sapiens OX=9606 GN=ACTN4 PE=1 SV=2 | 23 | 198 | 10 | 2.65E+07 | 2.46E+07 | 1.98E+07 | 2.32E+07 | 2.10E+07 | 2.21E+07 | 2.46E+07 | 2.07E+07 | 1.94E+07 | 1.82E+07 | 1.75E+07 | 1.69E+07 | 2.29E+07 | 1.95E+07 | 1.17 | 0.23 | 0.053517293 |
| Q16181 | Septin-7 OS=Homo sapiens OX=9606 GN=SEPTIN7 PE=1 SV=2 | 29 | 139 | 10 | 3.77E+07 | 2.80E+07 | 3.42E+07 | 3.22E+07 | 3.11E+07 | 3.62E+07 | 3.98E+07 | 3.01E+07 | 3.23E+07 | 2.72E+07 | 2.72E+07 | 2.99E+07 | 3.32E+07 | 3.11E+07 | 1.07 | 0.10 | 0.398713546 |
| O75533 | Splicing factor 3B subunit 1 OS=Homo sapiens OX=9606 GN=SF3B1 PE=1 SV=3 | 14 | 121 | 10 | 2.24E+07 | 2.11E+07 | 2.33E+07 | 2.28E+07 | 2.54E+07 | 2.29E+07 | 1.99E+07 | 1.88E+07 | 1.24E+07 | 1.35E+07 | 1.72E+07 | 1.36E+07 | 2.30E+07 | 1.59E+07 | 1.44 | 0.53 | 0.001566099 |
| O15371 | Eukaryotic translation initiation factor 3 subunit D OS=Homo sapiens OX=9606 GN=EIF3D PE=1 SV=1 | 28 | 105 | 10 | 2.09E+07 | 1.66E+07 | 2.64E+07 | 1.69E+07 | 2.56E+07 | 2.85E+07 | 9.59E+06 | 9.56E+06 | 1.41E+07 | 7.46E+06 | 1.49E+07 | 1.42E+07 | 2.25E+07 | 1.16E+07 | 1.93 | 0.95 | 0.001958966 |
| P41091 | Eukaryotic translation initiation factor 2 subunit 3 OS=Homo sapiens OX=9606 GN=EIF2S3 PE=1 SV=3 | 37 | 128 | 10 | 3.39E+07 | 3.87E+07 | 3.39E+07 | 3.80E+07 | 3.54E+07 | 3.72E+07 | 2.79E+07 | 2.86E+07 | 2.38E+07 | 2.64E+07 | 2.73E+07 | 2.59E+07 | 3.62E+07 | 2.66E+07 | 1.36 | 0.44 | 7.93979E-06 |
| P43121 | Cell surface glycoprotein MUC18 OS=Homo sapiens OX=9606 GN=MCAM PE=1 SV=2 | 24 | 106 | 10 | 3.71E+07 | 3.03E+07 | 3.18E+07 | 2.89E+07 | 2.73E+07 | 3.22E+07 | 1.54E+07 | 1.51E+07 | 1.24E+07 | 1.19E+07 | 1.26E+07 | 1.40E+07 | 3.13E+07 | 1.36E+07 | 2.30 | 1.20 | 9.26782E-06 |
| Q9BUJ2 | Heterogeneous nuclear ribonucleoprotein U-like protein 1 OS=Homo sapiens OX=9606 GN=HNRNPUL1 PE=1 SV=2 | 22 | 73 | 10 | 2.09E+07 | 1.55E+07 | 1.95E+07 | 2.32E+07 | 2.42E+07 | 2.04E+07 | 1.75E+07 | 1.09E+07 | 1.04E+07 | 1.24E+07 | 9.57E+06 | 8.70E+06 | 2.06E+07 | 1.16E+07 | 1.78 | 0.83 | 0.000525162 |
| Q02878 | 60S ribosomal protein L6 OS=Homo sapiens OX=9606 GN=RPL6 PE=1 SV=3 | 42 | 185 | 10 | 9.32E+07 | 9.06E+07 | 7.39E+07 | 6.28E+07 | 7.31E+07 | 6.57E+07 | 5.74E+07 | 5.36E+07 | 4.93E+07 | 4.22E+07 | 4.92E+07 | 3.67E+07 | 7.66E+07 | 4.81E+07 | 1.59 | 0.67 | 0.001378615 |
| P26599 | Polypyrimidine tract-binding protein 1 OS=Homo sapiens OX=9606 GN=PTBP1 PE=1 SV=1 | 28 | 126 | 10 | 3.28E+07 | 3.04E+07 | 2.67E+07 | 3.01E+07 | 2.80E+07 | 3.16E+07 | 2.33E+07 | 2.15E+07 | 1.77E+07 | 1.83E+07 | 1.80E+07 | 1.38E+07 | 2.99E+07 | 1.88E+07 | 1.60 | 0.67 | 8.70672E-05 |
| P61978 | Heterogeneous nuclear ribonucleoprotein K OS=Homo sapiens OX=9606 GN=HNRNPK PE=1 SV=1 | 28 | 161 | 10 | 7.59E+07 | 8.44E+07 | 7.77E+07 | 7.31E+07 | 7.04E+07 | 7.33E+07 | 4.64E+07 | 5.21E+07 | 4.07E+07 | 4.36E+07 | 4.11E+07 | 3.97E+07 | 7.58E+07 | 4.40E+07 | 1.72 | 0.79 | 4.35594E-07 |
| Q16531 | DNA damage-binding protein 1 OS=Homo sapiens OX=9606 GN=DDB1 PE=1 SV=1 | 14 | 105 | 10 | 3.41E+07 | 3.71E+07 | 2.40E+07 | 2.48E+07 | 2.52E+07 | 2.33E+07 | 2.37E+07 | 2.19E+07 | 1.60E+07 | 1.73E+07 | 1.78E+07 | 1.74E+07 | 2.81E+07 | 1.90E+07 | 1.48 | 0.56 | 0.0113477 |
| P14866 | Heterogeneous nuclear ribonucleoprotein L OS=Homo sapiens OX=9606 GN=HNRNPL PE=1 SV=2 | 26 | 78 | 10 | 2.48E+07 | 2.29E+07 | 2.19E+07 | 1.98E+07 | 1.99E+07 | 2.39E+07 | 1.19E+07 | 1.23E+07 | 1.21E+07 | 1.21E+07 | 1.14E+07 | 6.79E+06 | 2.22E+07 | 1.11E+07 | 2.00 | 1.00 | 3.65452E-06 |
| O00571 | ATP-dependent RNA helicase DDX3X OS=Homo sapiens OX=9606 GN=DDX3X PE=1 SV=3 | 20 | 96 | 10 | 2.64E+07 | 2.56E+07 | 2.61E+07 | 2.49E+07 | 1.99E+07 | 2.56E+07 | 1.81E+07 | 1.92E+07 | 1.55E+07 | 1.26E+07 | 1.63E+07 | 1.14E+07 | 2.48E+07 | 1.55E+07 | 1.59 | 0.67 | 0.000212078 |
| P22087 | rRNA 2'-O-methyltransferase fibrillarin OS=Homo sapiens OX=9606 GN=FBL PE=1 SV=2 | 41 | 127 | 10 | 3.48E+07 | 3.73E+07 | 3.76E+07 | 3.30E+07 | 3.77E+07 | 3.88E+07 | 2.55E+07 | 2.71E+07 | 2.09E+07 | 2.20E+07 | 1.90E+07 | 2.09E+07 | 3.65E+07 | 2.26E+07 | 1.62 | 0.69 | 8.52681E-06 |
| P53618 | Coatomer subunit beta OS=Homo sapiens OX=9606 GN=COPB1 PE=1 SV=3 | 17 | 90 | 10 | 2.82E+07 | 3.15E+07 | 3.62E+07 | 2.66E+07 | 4.54E+07 | 3.74E+07 | 1.96E+07 | 2.07E+07 | 2.41E+07 | 1.85E+07 | 1.99E+07 | 1.61E+07 | 3.42E+07 | 1.98E+07 | 1.73 | 0.79 | 0.002640345 |
| Q15582 | Transforming growth factor-beta-induced protein ig-h3 OS=Homo sapiens OX=9606 GN=TGFBI PE=1 SV=1 | 19 | 94 | 10 | 1.63E+07 | 1.66E+07 | 2.40E+07 | 2.19E+07 | 2.22E+07 | 2.39E+07 | 3.79E+07 | 2.42E+07 | 3.34E+07 | 3.04E+07 | 3.88E+07 | 3.80E+07 | 2.08E+07 | 3.38E+07 | 0.62 | -0.70 | 0.0013146 |
| Q9NTK5 | Obg-like ATPase 1 OS=Homo sapiens OX=9606 GN=OLA1 PE=1 SV=2 | 28 | 148 | 10 | 3.21E+07 | 3.01E+07 | 2.74E+07 | 2.57E+07 | 2.97E+07 | 2.58E+07 | 3.69E+07 | 3.37E+07 | 3.18E+07 | 2.56E+07 | 3.38E+07 | 1.96E+07 | 2.85E+07 | 3.02E+07 | 0.94 | -0.09 | 0.548411122 |
| Q08945 | FACT complex subunit SSRP1 OS=Homo sapiens OX=9606 GN=SSRP1 PE=1 SV=1 | 19 | 93 | 10 | 3.07E+07 | 2.70E+07 | 3.08E+07 | 3.08E+07 | 2.70E+07 | 2.86E+07 | 2.07E+07 | 1.59E+07 | 1.74E+07 | 1.56E+07 | 1.28E+07 | 1.03E+07 | 2.92E+07 | 1.55E+07 | 1.89 | 0.92 | 5.03591E-05 |
| Q92616 | eIF-2-alpha kinase activator GCN1 OS=Homo sapiens OX=9606 GN=GCN1 PE=1 SV=6 | 6 | 45 | 10 | 3.90E+07 | 3.54E+07 | 3.28E+07 | 2.19E+07 | 3.39E+07 | 3.02E+07 | 3.42E+07 | 3.87E+07 | 3.93E+07 | 2.11E+07 | 3.84E+07 | 1.86E+07 | 3.22E+07 | 3.17E+07 | 1.01 | 0.02 | 0.919279436 |
| Q16610 | Extracellular matrix protein 1 OS=Homo sapiens OX=9606 GN=ECM1 PE=1 SV=2 | 28 | 28 | 10 | 4.92E+06 | 4.40E+06 | 1.29E+06 | 3.02E+06 | 1.36E+06 | 3.21E+06 | 1.45E+07 | 4.69E+06 | 6.20E+06 | 1.45E+07 | 5.83E+06 | 5.18E+07 | 3.03E+06 | 1.62E+07 | 0.19 | -2.42 | 0.131775697 |
| P27694 | Replication protein A 70 kDa DNA-binding subunit OS=Homo sapiens OX=9606 GN=RPA1 PE=1 SV=2 | 24 | 94 | 10 | 3.02E+07 | 3.32E+07 | 2.30E+07 | 2.59E+07 | 2.36E+07 | 3.01E+07 | 2.36E+07 | 2.38E+07 | 1.84E+07 | 1.73E+07 | 1.66E+07 | 1.52E+07 | 2.77E+07 | 1.91E+07 | 1.44 | 0.53 | 0.003686854 |
| P62249 | 40S ribosomal protein S16 OS=Homo sapiens OX=9606 GN=RPS16 PE=1 SV=2 | 49 | 420 | 10 | 1.68E+08 | 1.76E+08 | 1.57E+08 | 1.58E+08 | 1.52E+08 | 1.47E+08 | 1.22E+08 | 1.27E+08 | 1.02E+08 | 1.06E+08 | 1.14E+08 | 1.03E+08 | 1.60E+08 | 1.12E+08 | 1.42 | 0.50 | 1.45355E-05 |
| P43490 | Nicotinamide phosphoribosyltransferase OS=Homo sapiens OX=9606 GN=NAMPT PE=1 SV=1 | 30 | 77 | 10 | 2.78E+07 | 2.95E+07 | 2.70E+07 | 2.76E+07 | 2.73E+07 | 2.24E+07 | 2.44E+07 | 2.02E+07 | 1.80E+07 | 2.02E+07 | 2.24E+07 | 1.59E+07 | 2.69E+07 | 2.02E+07 | 1.33 | 0.42 | 0.001806899 |
| P35573 | Glycogen debranching enzyme OS=Homo sapiens OX=9606 GN=AGL PE=1 SV=3 | 9 | 130 | 10 | 2.65E+07 | 2.66E+07 | 2.20E+07 | 1.90E+07 | 1.97E+07 | 2.48E+07 | 3.69E+07 | 3.59E+07 | 2.96E+07 | 2.56E+07 | 3.36E+07 | 2.33E+07 | 2.31E+07 | 3.08E+07 | 0.75 | -0.42 | 0.01975363 |
| O60763 | General vesicular transport factor p115 OS=Homo sapiens OX=9606 GN=USO1 PE=1 SV=2 | 11 | 94 | 10 | 8.87E+07 | 9.47E+07 | 1.34E+08 | 1.54E+08 | 1.55E+08 | 1.21E+08 | 1.33E+08 | 1.22E+08 | 1.88E+08 | 1.68E+08 | 1.38E+08 | 1.19E+08 | 1.25E+08 | 1.45E+08 | 0.86 | -0.21 | 0.244413418 |
| P22102 | Trifunctional purine biosynthetic protein adenosine-3 OS=Homo sapiens OX=9606 GN=GART PE=1 SV=1 | 16 | 60 | 10 | 2.28E+07 | 2.67E+07 | 2.14E+07 | 2.88E+07 | 2.37E+07 | 2.06E+07 | 1.44E+07 | 1.62E+07 | 1.36E+07 | 1.55E+07 | 1.19E+07 | 1.07E+07 | 2.40E+07 | 1.37E+07 | 1.75 | 0.80 | 0.000120978 |
| O00429 | Dynamin-1-like protein OS=Homo sapiens OX=9606 GN=DNM1L PE=1 SV=2 | 19 | 81 | 10 | 1.56E+07 | 2.01E+07 | 1.72E+07 | 1.49E+07 | 1.72E+07 | 1.56E+07 | 1.48E+07 | 1.45E+07 | 1.78E+07 | 1.41E+07 | 1.88E+07 | 1.30E+07 | 1.68E+07 | 1.55E+07 | 1.08 | 0.11 | 0.316199479 |
| P20774 | Mimecan OS=Homo sapiens OX=9606 GN=OGN PE=1 SV=1 | 27 | 295 | 10 | 1.03E+08 | 9.22E+07 | 9.30E+07 | 1.25E+08 | 1.11E+08 | 9.77E+07 | 1.46E+08 | 1.28E+08 | 1.27E+08 | 1.42E+08 | 1.29E+08 | 1.17E+08 | 1.04E+08 | 1.32E+08 | 0.79 | -0.35 | 0.001925452 |
| P18206 | Vinculin OS=Homo sapiens OX=9606 GN=VCL PE=1 SV=4 | 10 | 94 | 10 | 2.48E+07 | 2.66E+07 | 3.00E+07 | 3.05E+07 | 2.24E+07 | 2.70E+07 | 3.10E+07 | 2.94E+07 | 3.32E+07 | 3.17E+07 | 3.14E+07 | 3.15E+07 | 2.69E+07 | 3.14E+07 | 0.86 | -0.22 | 0.013918794 |
| Q9UNH7 | Sorting nexin-6 OS=Homo sapiens OX=9606 GN=SNX6 PE=1 SV=1 | 22 | 64 | 10 | 1.76E+07 | 2.24E+07 | 2.11E+07 | 2.02E+07 | 1.99E+07 | 1.82E+07 | 1.57E+07 | 1.89E+07 | 1.78E+07 | 1.45E+07 | 1.78E+07 | 1.78E+07 | 1.99E+07 | 1.71E+07 | 1.16 | 0.22 | 0.018710378 |
| P08779 | Keratin, type I cytoskeletal 16 OS=Homo sapiens OX=9606 GN=KRT16 PE=1 SV=4 | 53 | 480 | 11 | 1.66E+08 | 7.54E+07 | 4.25E+07 | 5.44E+07 | 4.59E+07 | 1.39E+08 | 2.00E+07 | 1.66E+08 | 8.20E+07 | 9.51E+07 | 5.68E+07 | 3.56E+07 | 8.72E+07 | 7.59E+07 | 1.15 | 0.20 | 0.715868387 |
| P06748 | Nucleophosmin OS=Homo sapiens OX=9606 GN=NPM1 PE=1 SV=2 | 38 | 225 | 11 | 1.21E+08 | 1.30E+08 | 1.53E+08 | 1.42E+08 | 1.36E+08 | 1.55E+08 | 9.02E+07 | 8.84E+07 | 9.02E+07 | 6.97E+07 | 7.51E+07 | 7.11E+07 | 1.39E+08 | 8.08E+07 | 1.73 | 0.79 | 8.36565E-06 |
| P16401 | Histone H1.5 OS=Homo sapiens OX=9606 GN=H1-5 PE=1 SV=3 | 36 | 739 | 11 | 8.66E+08 | 8.83E+08 | 8.33E+08 | 8.93E+08 | 8.87E+08 | 8.12E+08 | 7.64E+08 | 7.75E+08 | 8.18E+08 | 8.20E+08 | 7.09E+08 | 6.48E+08 | 8.62E+08 | 7.56E+08 | 1.14 | 0.19 | 0.009254022 |
| P12814 | Alpha-actinin-1 OS=Homo sapiens OX=9606 GN=ACTN1 PE=1 SV=2 | 25 | 185 | 11 | 4.80E+07 | 4.40E+07 | 4.02E+07 | 3.82E+07 | 3.56E+07 | 4.15E+07 | 3.72E+07 | 3.15E+07 | 2.38E+07 | 2.43E+07 | 2.76E+07 | 2.64E+07 | 4.13E+07 | 2.85E+07 | 1.45 | 0.54 | 0.000939916 |
| P00338 | L-lactate dehydrogenase A chain OS=Homo sapiens OX=9606 GN=LDHA PE=1 SV=2 | 51 | 387 | 11 | 1.40E+08 | 1.53E+08 | 1.33E+08 | 1.43E+08 | 1.43E+08 | 1.29E+08 | 9.49E+07 | 9.04E+07 | 9.97E+07 | 1.01E+08 | 8.53E+07 | 8.32E+07 | 1.40E+08 | 9.25E+07 | 1.52 | 0.60 | 1.18618E-06 |
| P23142 | Fibulin-1 OS=Homo sapiens OX=9606 GN=FBLN1 PE=1 SV=4 | 16 | 591 | 11 | 2.25E+08 | 2.13E+08 | 1.96E+08 | 2.24E+08 | 1.89E+08 | 1.91E+08 | 3.24E+08 | 3.25E+08 | 2.76E+08 | 3.48E+08 | 2.69E+08 | 2.54E+08 | 2.06E+08 | 2.99E+08 | 0.69 | -0.54 | 0.000983748 |
| P07195 | L-lactate dehydrogenase B chain OS=Homo sapiens OX=9606 GN=LDHB PE=1 SV=2 | 50 | 544 | 11 | 2.07E+08 | 1.91E+08 | 1.82E+08 | 1.88E+08 | 1.86E+08 | 1.63E+08 | 1.68E+08 | 1.60E+08 | 1.68E+08 | 1.77E+08 | 1.70E+08 | 1.56E+08 | 1.86E+08 | 1.67E+08 | 1.12 | 0.16 | 0.018377262 |
| P00966 | Argininosuccinate synthase OS=Homo sapiens OX=9606 GN=ASS1 PE=1 SV=2 | 41 | 372 | 11 | 1.21E+08 | 1.16E+08 | 1.30E+08 | 1.25E+08 | 1.27E+08 | 1.35E+08 | 1.98E+08 | 1.81E+08 | 1.68E+08 | 1.67E+08 | 1.75E+08 | 1.47E+08 | 1.26E+08 | 1.73E+08 | 0.73 | -0.46 | 0.00050468 |
| Q9ULV4 | Coronin-1C OS=Homo sapiens OX=9606 GN=CORO1C PE=1 SV=1 | 30 | 244 | 11 | 6.23E+07 | 6.02E+07 | 5.89E+07 | 5.74E+07 | 5.43E+07 | 4.94E+07 | 6.24E+07 | 5.12E+07 | 5.33E+07 | 4.33E+07 | 4.56E+07 | 4.28E+07 | 5.71E+07 | 4.98E+07 | 1.15 | 0.20 | 0.074563415 |
| Q9Y265 | RuvB-like 1 OS=Homo sapiens OX=9606 GN=RUVBL1 PE=1 SV=1 | 32 | 216 | 11 | 5.63E+07 | 5.52E+07 | 4.22E+07 | 4.58E+07 | 4.16E+07 | 4.78E+07 | 3.88E+07 | 4.20E+07 | 2.87E+07 | 3.48E+07 | 3.03E+07 | 2.88E+07 | 4.81E+07 | 3.39E+07 | 1.42 | 0.50 | 0.002162321 |
| Q92841 | Probable ATP-dependent RNA helicase DDX17 OS=Homo sapiens OX=9606 GN=DDX17 PE=1 SV=2 | 26 | 154 | 11 | 6.10E+07 | 6.65E+07 | 5.71E+07 | 6.29E+07 | 6.32E+07 | 5.89E+07 | 4.85E+07 | 4.42E+07 | 3.78E+07 | 4.24E+07 | 3.36E+07 | 3.26E+07 | 6.16E+07 | 3.99E+07 | 1.55 | 0.63 | 8.79285E-05 |
| P46821 | Microtubule-associated protein 1B OS=Homo sapiens OX=9606 GN=MAP1B PE=1 SV=2 | 8 | 69 | 11 | 2.06E+07 | 2.04E+07 | 1.70E+07 | 1.81E+07 | 1.61E+07 | 1.90E+07 | 1.21E+07 | 1.36E+07 | 9.90E+06 | 1.02E+07 | 9.57E+06 | 1.07E+07 | 1.86E+07 | 1.10E+07 | 1.68 | 0.75 | 1.75459E-05 |
| P55263 | Adenosine kinase OS=Homo sapiens OX=9606 GN=ADK PE=1 SV=2 | 35 | 173 | 11 | 5.46E+07 | 4.62E+07 | 4.31E+07 | 6.32E+07 | 5.59E+07 | 5.94E+07 | 7.04E+07 | 7.28E+07 | 6.43E+07 | 6.00E+07 | 6.26E+07 | 5.23E+07 | 5.37E+07 | 6.37E+07 | 0.84 | -0.25 | 0.045136293 |
| P01871 | Immunoglobulin heavy constant mu OS=Homo sapiens OX=9606 GN=IGHM PE=1 SV=4 | 31 | 44 | 11 | 9.35E+05 | 8.21E+05 |  | 4.69E+05 |  | 3.15E+05 | 8.75E+06 | 1.57E+06 | 4.66E+05 |  |  | 7.40E+07 | 6.35E+05 | 2.12E+07 | 0.03 | -5.06 | 0.329295506 |
| P62333 | 26S proteasome regulatory subunit 10B OS=Homo sapiens OX=9606 GN=PSMC6 PE=1 SV=1 | 34 | 140 | 11 | 5.19E+07 | 4.97E+07 | 4.26E+07 | 4.42E+07 | 4.56E+07 | 4.76E+07 | 3.50E+07 | 3.60E+07 | 3.47E+07 | 3.29E+07 | 3.12E+07 | 2.81E+07 | 4.69E+07 | 3.30E+07 | 1.42 | 0.51 | 2.3661E-05 |
| P33176 | Kinesin-1 heavy chain OS=Homo sapiens OX=9606 GN=KIF5B PE=1 SV=1 | 18 | 94 | 11 | 2.13E+07 | 1.83E+07 | 2.05E+07 | 2.04E+07 | 2.01E+07 | 2.24E+07 | 1.08E+07 | 8.95E+06 | 8.30E+06 | 8.43E+06 | 8.53E+06 | 8.81E+06 | 2.05E+07 | 8.96E+06 | 2.29 | 1.19 | 5.34554E-08 |
| P33993 | DNA replication licensing factor MCM7 OS=Homo sapiens OX=9606 GN=MCM7 PE=1 SV=4 | 22 | 108 | 11 | 2.79E+07 | 2.39E+07 | 2.49E+07 | 2.60E+07 | 2.32E+07 | 2.40E+07 | 1.27E+07 | 1.43E+07 | 1.29E+07 | 1.34E+07 | 1.25E+07 | 1.11E+07 | 2.50E+07 | 1.28E+07 | 1.95 | 0.96 | 3.55614E-07 |
| Q99873 | Protein arginine N-methyltransferase 1 OS=Homo sapiens OX=9606 GN=PRMT1 PE=1 SV=3 | 30 | 165 | 11 | 5.12E+07 | 5.65E+07 | 5.33E+07 | 5.57E+07 | 5.20E+07 | 4.87E+07 | 4.43E+07 | 4.30E+07 | 3.91E+07 | 4.16E+07 | 3.71E+07 | 3.65E+07 | 5.29E+07 | 4.03E+07 | 1.31 | 0.39 | 3.3042E-05 |
| Q14203 | Dynactin subunit 1 OS=Homo sapiens OX=9606 GN=DCTN1 PE=1 SV=3 | 12 | 91 | 11 | 2.45E+07 | 2.60E+07 | 2.64E+07 | 2.68E+07 | 2.70E+07 | 2.61E+07 | 1.31E+07 | 1.44E+07 | 1.63E+07 | 1.48E+07 | 1.42E+07 | 1.32E+07 | 2.61E+07 | 1.43E+07 | 1.83 | 0.87 | 7.37935E-09 |
| Q7L576 | Cytoplasmic FMR1-interacting protein 1 OS=Homo sapiens OX=9606 GN=CYFIP1 PE=1 SV=1 | 10 | 141 | 11 | 3.54E+07 | 3.15E+07 | 2.30E+07 | 2.78E+07 | 2.78E+07 | 2.41E+07 | 2.41E+07 | 2.09E+07 | 2.03E+07 | 2.21E+07 | 1.94E+07 | 1.81E+07 | 2.83E+07 | 2.08E+07 | 1.36 | 0.44 | 0.00889316 |
| O94979 | Protein transport protein Sec31A OS=Homo sapiens OX=9606 GN=SEC31A PE=1 SV=3 | 11 | 122 | 11 | 3.68E+07 | 3.97E+07 | 3.11E+07 | 3.55E+07 | 3.64E+07 | 3.76E+07 | 3.10E+07 | 2.71E+07 | 2.65E+07 | 2.48E+07 | 2.48E+07 | 2.38E+07 | 3.62E+07 | 2.63E+07 | 1.37 | 0.46 | 9.75311E-05 |
| Q02543 | 60S ribosomal protein L18a OS=Homo sapiens OX=9606 GN=RPL18A PE=1 SV=2 | 44 | 214 | 11 | 9.26E+07 | 8.26E+07 | 7.36E+07 | 8.74E+07 | 8.43E+07 | 8.05E+07 | 5.81E+07 | 5.09E+07 | 4.45E+07 | 5.21E+07 | 4.48E+07 | 4.56E+07 | 8.35E+07 | 4.93E+07 | 1.69 | 0.76 | 2.02155E-06 |
| Q9P2R3 | Rabankyrin-5 OS=Homo sapiens OX=9606 GN=ANKFY1 PE=1 SV=2 | 13 | 62 | 11 | 1.95E+07 | 1.84E+07 | 1.68E+07 | 1.54E+07 | 1.49E+07 | 1.75E+07 | 1.71E+07 | 1.45E+07 | 1.70E+07 | 1.29E+07 | 1.47E+07 | 1.47E+07 | 1.71E+07 | 1.52E+07 | 1.13 | 0.17 | 0.078339289 |
| P49591 | Serine--tRNA ligase, cytoplasmic OS=Homo sapiens OX=9606 GN=SARS PE=1 SV=3 | 31 | 147 | 11 | 2.75E+07 | 2.76E+07 | 2.58E+07 | 2.40E+07 | 2.64E+07 | 3.08E+07 | 1.85E+07 | 1.74E+07 | 1.30E+07 | 1.25E+07 | 1.50E+07 | 1.34E+07 | 2.70E+07 | 1.50E+07 | 1.81 | 0.85 | 5.81886E-06 |
| P47897 | Glutamine--tRNA ligase OS=Homo sapiens OX=9606 GN=QARS PE=1 SV=1 | 18 | 115 | 11 | 2.85E+07 | 2.53E+07 | 2.30E+07 | 3.19E+07 | 2.51E+07 | 2.34E+07 | 1.47E+07 | 1.33E+07 | 1.98E+07 | 1.72E+07 | 1.52E+07 | 1.38E+07 | 2.62E+07 | 1.57E+07 | 1.67 | 0.74 | 0.000166012 |
| P46781 | 40S ribosomal protein S9 OS=Homo sapiens OX=9606 GN=RPS9 PE=1 SV=3 | 38 | 376 | 11 | 1.46E+08 | 1.61E+08 | 1.80E+08 | 1.68E+08 | 1.80E+08 | 1.43E+08 | 1.18E+08 | 1.23E+08 | 1.16E+08 | 1.20E+08 | 1.01E+08 | 8.01E+07 | 1.63E+08 | 1.10E+08 | 1.49 | 0.57 | 0.000204504 |
| Q9BXJ9 | N-alpha-acetyltransferase 15, NatA auxiliary subunit OS=Homo sapiens OX=9606 GN=NAA15 PE=1 SV=1 | 13 | 76 | 11 | 2.13E+07 | 1.90E+07 | 1.90E+07 | 1.97E+07 | 1.83E+07 | 2.14E+07 | 1.44E+07 | 1.39E+07 | 1.05E+07 | 1.22E+07 | 1.27E+07 | 1.11E+07 | 1.98E+07 | 1.25E+07 | 1.59 | 0.66 | 5.6599E-06 |
| P68366 | Tubulin alpha-4A chain OS=Homo sapiens OX=9606 GN=TUBA4A PE=1 SV=1 | 75 | 3579 | 12 | 3.82E+08 | 3.64E+08 | 2.90E+08 | 2.84E+08 | 2.80E+08 | 3.51E+08 | 3.82E+08 | 4.03E+08 | 3.42E+08 | 2.93E+08 | 3.91E+08 | 3.20E+08 | 3.25E+08 | 3.55E+08 | 0.92 | -0.13 | 0.271398581 |
| P68104 | Elongation factor 1-alpha 1 OS=Homo sapiens OX=9606 GN=EEF1A1 PE=1 SV=1 | 53 | 1737 | 12 | 1.45E+09 | 1.56E+09 | 1.49E+09 | 1.58E+09 | 1.60E+09 | 1.62E+09 | 1.23E+09 | 1.33E+09 | 1.43E+09 | 1.36E+09 | 1.43E+09 | 1.11E+09 | 1.55E+09 | 1.32E+09 | 1.18 | 0.23 | 0.004161394 |
| P06576 | ATP synthase subunit beta, mitochondrial OS=Homo sapiens OX=9606 GN=ATP5F1B PE=1 SV=3 | 43 | 227 | 12 | 4.97E+07 | 5.47E+07 | 5.59E+07 | 4.16E+07 | 5.42E+07 | 4.97E+07 | 5.14E+07 | 5.91E+07 | 4.67E+07 | 4.52E+07 | 5.16E+07 | 4.65E+07 | 5.10E+07 | 5.01E+07 | 1.02 | 0.03 | 0.769530694 |
| P08865 | 40S ribosomal protein SA OS=Homo sapiens OX=9606 GN=RPSA PE=1 SV=4 | 44 | 427 | 12 | 2.68E+08 | 2.89E+08 | 2.13E+08 | 2.48E+08 | 2.18E+08 | 2.37E+08 | 2.00E+08 | 2.27E+08 | 1.84E+08 | 2.00E+08 | 1.87E+08 | 1.82E+08 | 2.45E+08 | 1.97E+08 | 1.25 | 0.32 | 0.007669479 |
| P09525 | Annexin A4 OS=Homo sapiens OX=9606 GN=ANXA4 PE=1 SV=4 | 50 | 226 | 12 | 5.51E+07 | 4.82E+07 | 4.95E+07 | 5.84E+07 | 4.55E+07 | 5.70E+07 | 5.13E+07 | 4.61E+07 | 4.60E+07 | 3.68E+07 | 3.96E+07 | 3.66E+07 | 5.23E+07 | 4.27E+07 | 1.22 | 0.29 | 0.015035297 |
| P61224 | Ras-related protein Rap-1b OS=Homo sapiens OX=9606 GN=RAP1B PE=1 SV=1 | 73 | 360 | 12 | 1.86E+08 | 1.83E+08 | 1.73E+08 | 1.91E+08 | 1.71E+08 | 1.83E+08 | 2.09E+08 | 1.98E+08 | 1.96E+08 | 1.91E+08 | 2.07E+08 | 1.88E+08 | 1.81E+08 | 1.98E+08 | 0.91 | -0.13 | 0.004417743 |
| O75390 | Citrate synthase, mitochondrial OS=Homo sapiens OX=9606 GN=CS PE=1 SV=2 | 34 | 274 | 12 | 8.63E+07 | 9.06E+07 | 7.76E+07 | 9.45E+07 | 9.24E+07 | 8.71E+07 | 7.71E+07 | 7.70E+07 | 6.26E+07 | 6.73E+07 | 6.63E+07 | 6.43E+07 | 8.81E+07 | 6.91E+07 | 1.27 | 0.35 | 0.000347916 |
| P06396 | Gelsolin OS=Homo sapiens OX=9606 GN=GSN PE=1 SV=1 | 20 | 523 | 12 | 1.95E+08 | 1.96E+08 | 1.56E+08 | 1.86E+08 | 1.62E+08 | 1.69E+08 | 2.54E+08 | 2.36E+08 | 2.25E+08 | 2.57E+08 | 1.93E+08 | 1.83E+08 | 1.77E+08 | 2.25E+08 | 0.79 | -0.34 | 0.011782566 |
| P04075 | Fructose-bisphosphate aldolase A OS=Homo sapiens OX=9606 GN=ALDOA PE=1 SV=2 | 55 | 201 | 12 | 6.24E+07 | 5.46E+07 | 5.45E+07 | 6.10E+07 | 5.42E+07 | 5.59E+07 | 4.61E+07 | 4.94E+07 | 4.87E+07 | 5.05E+07 | 4.49E+07 | 4.03E+07 | 5.71E+07 | 4.67E+07 | 1.22 | 0.29 | 0.000613766 |
| P68400 | Casein kinase II subunit alpha OS=Homo sapiens OX=9606 GN=CSNK2A1 PE=1 SV=1 | 44 | 213 | 12 | 1.01E+08 | 9.81E+07 | 7.92E+07 | 8.27E+07 | 7.58E+07 | 8.39E+07 | 8.75E+07 | 7.54E+07 | 6.92E+07 | 6.40E+07 | 7.07E+07 | 6.99E+07 | 8.68E+07 | 7.28E+07 | 1.19 | 0.25 | 0.02688942 |
| P15880 | 40S ribosomal protein S2 OS=Homo sapiens OX=9606 GN=RPS2 PE=1 SV=2 | 41 | 382 | 12 | 2.99E+08 | 2.89E+08 | 2.95E+08 | 3.31E+08 | 2.88E+08 | 2.82E+08 | 1.98E+08 | 2.00E+08 | 1.94E+08 | 2.01E+08 | 1.84E+08 | 1.71E+08 | 2.97E+08 | 1.91E+08 | 1.55 | 0.64 | 8.86675E-07 |
| P00488 | Coagulation factor XIII A chain OS=Homo sapiens OX=9606 GN=F13A1 PE=1 SV=4 | 22 | 225 | 12 | 7.38E+07 | 5.69E+07 | 6.09E+07 | 6.05E+07 | 6.36E+07 | 6.21E+07 | 9.12E+07 | 7.53E+07 | 5.66E+07 | 6.21E+07 | 6.20E+07 | 1.41E+08 | 6.30E+07 | 8.13E+07 | 0.77 | -0.37 | 0.218186447 |
| Q08431 | Lactadherin OS=Homo sapiens OX=9606 GN=MFGE8 PE=1 SV=3 | 41 | 289 | 12 | 5.92E+07 | 5.27E+07 | 4.39E+07 | 4.99E+07 | 4.55E+07 | 4.56E+07 | 9.13E+07 | 1.01E+08 | 7.57E+07 | 7.90E+07 | 7.34E+07 | 6.73E+07 | 4.95E+07 | 8.13E+07 | 0.61 | -0.72 | 0.000741471 |
| Q9BQG0 | Myb-binding protein 1A OS=Homo sapiens OX=9606 GN=MYBBP1A PE=1 SV=2 | 13 | 132 | 12 | 2.29E+07 | 3.08E+07 | 2.44E+07 | 2.72E+07 | 2.82E+07 | 2.62E+07 | 1.61E+07 | 1.65E+07 | 1.19E+07 | 1.50E+07 | 1.37E+07 | 1.36E+07 | 2.66E+07 | 1.45E+07 | 1.84 | 0.88 | 1.30784E-05 |
| Q96CW1 | AP-2 complex subunit mu OS=Homo sapiens OX=9606 GN=AP2M1 PE=1 SV=2 | 37 | 237 | 12 | 8.04E+07 | 8.16E+07 | 6.98E+07 | 7.69E+07 | 7.36E+07 | 7.14E+07 | 5.47E+07 | 5.10E+07 | 4.86E+07 | 5.38E+07 | 4.74E+07 | 4.28E+07 | 7.56E+07 | 4.97E+07 | 1.52 | 0.61 | 2.2248E-06 |
| P26639 | Threonine--tRNA ligase 1, cytoplasmic OS=Homo sapiens OX=9606 GN=TARS1 PE=1 SV=3 | 23 | 222 | 12 | 4.32E+07 | 3.93E+07 | 4.30E+07 | 4.88E+07 | 4.61E+07 | 4.06E+07 | 3.86E+07 | 4.34E+07 | 4.10E+07 | 3.19E+07 | 4.05E+07 | 3.41E+07 | 4.35E+07 | 3.83E+07 | 1.14 | 0.19 | 0.045553176 |
| P15121 | Aldo-keto reductase family 1 member B1 OS=Homo sapiens OX=9606 GN=AKR1B1 PE=1 SV=3 | 49 | 268 | 12 | 8.76E+07 | 1.03E+08 | 8.75E+07 | 9.73E+07 | 9.47E+07 | 9.66E+07 | 9.82E+07 | 1.15E+08 | 8.47E+07 | 9.71E+07 | 9.87E+07 | 8.90E+07 | 9.45E+07 | 9.71E+07 | 0.97 | -0.04 | 0.61722682 |
| Q15008 | 26S proteasome non-ATPase regulatory subunit 6 OS=Homo sapiens OX=9606 GN=PSMD6 PE=1 SV=1 | 42 | 116 | 12 | 2.64E+07 | 3.90E+07 | 2.47E+07 | 3.05E+07 | 2.83E+07 | 2.53E+07 | 1.93E+07 | 1.36E+07 | 1.43E+07 | 1.54E+07 | 1.60E+07 | 1.19E+07 | 2.90E+07 | 1.51E+07 | 1.93 | 0.94 | 0.000603031 |
| P11413 | Glucose-6-phosphate 1-dehydrogenase OS=Homo sapiens OX=9606 GN=G6PD PE=1 SV=4 | 25 | 203 | 12 | 4.16E+07 | 4.05E+07 | 3.89E+07 | 4.38E+07 | 3.95E+07 | 3.70E+07 | 3.85E+07 | 3.37E+07 | 3.71E+07 | 3.46E+07 | 3.33E+07 | 3.05E+07 | 4.02E+07 | 3.46E+07 | 1.16 | 0.22 | 0.0041236 |
| P50995 | Annexin A11 OS=Homo sapiens OX=9606 GN=ANXA11 PE=1 SV=1 | 27 | 215 | 12 | 6.26E+07 | 5.86E+07 | 5.42E+07 | 5.74E+07 | 5.20E+07 | 5.20E+07 | 5.72E+07 | 5.51E+07 | 4.70E+07 | 4.53E+07 | 4.33E+07 | 4.74E+07 | 5.61E+07 | 4.92E+07 | 1.14 | 0.19 | 0.037287364 |
| P62269 | 40S ribosomal protein S18 OS=Homo sapiens OX=9606 GN=RPS18 PE=1 SV=3 | 51 | 302 | 12 | 9.69E+07 | 9.40E+07 | 7.63E+07 | 9.97E+07 | 9.21E+07 | 8.33E+07 | 7.80E+07 | 7.73E+07 | 7.84E+07 | 8.40E+07 | 6.94E+07 | 5.91E+07 | 9.04E+07 | 7.44E+07 | 1.22 | 0.28 | 0.01061015 |
| P60842 | Eukaryotic initiation factor 4A-I OS=Homo sapiens OX=9606 GN=EIF4A1 PE=1 SV=1 | 73 | 595 | 13 | 2.85E+08 | 3.39E+08 | 2.85E+08 | 3.00E+08 | 2.88E+08 | 3.15E+08 | 2.00E+08 | 2.10E+08 | 2.03E+08 | 2.02E+08 | 2.16E+08 | 2.25E+08 | 3.02E+08 | 2.09E+08 | 1.44 | 0.53 | 2.91951E-05 |
| P01889 | HLA class I histocompatibility antigen, B alpha chain OS=Homo sapiens OX=9606 GN=HLA-B PE=1 SV=3 | 57 | 672 | 13 | 4.13E+08 | 4.02E+08 | 3.71E+08 | 4.12E+08 | 3.90E+08 | 3.62E+08 | 1.84E+08 | 1.76E+08 | 1.64E+08 | 1.67E+08 | 1.61E+08 | 1.43E+08 | 3.92E+08 | 1.66E+08 | 2.36 | 1.24 | 7.51874E-09 |
| Q9H4M9 | EH domain-containing protein 1 OS=Homo sapiens OX=9606 GN=EHD1 PE=1 SV=2 | 54 | 379 | 13 | 1.16E+08 | 1.15E+08 | 1.08E+08 | 1.08E+08 | 1.08E+08 | 1.02E+08 | 1.23E+08 | 1.33E+08 | 1.11E+08 | 1.19E+08 | 1.14E+08 | 1.05E+08 | 1.10E+08 | 1.18E+08 | 0.93 | -0.10 | 0.116809906 |
| P63010 | AP-2 complex subunit beta OS=Homo sapiens OX=9606 GN=AP2B1 PE=1 SV=1 | 29 | 277 | 13 | 7.94E+07 | 7.43E+07 | 8.73E+07 | 7.45E+07 | 8.36E+07 | 7.78E+07 | 5.87E+07 | 4.70E+07 | 5.57E+07 | 4.15E+07 | 5.58E+07 | 4.36E+07 | 7.95E+07 | 5.04E+07 | 1.58 | 0.66 | 2.08862E-05 |
| Q9UMS4 | Pre-mRNA-processing factor 19 OS=Homo sapiens OX=9606 GN=PRPF19 PE=1 SV=1 | 44 | 208 | 13 | 5.98E+07 | 6.56E+07 | 7.82E+07 | 6.10E+07 | 6.77E+07 | 6.94E+07 | 4.45E+07 | 4.92E+07 | 4.27E+07 | 4.54E+07 | 5.11E+07 | 4.54E+07 | 6.69E+07 | 4.64E+07 | 1.44 | 0.53 | 0.000229779 |
| O00299 | Chloride intracellular channel protein 1 OS=Homo sapiens OX=9606 GN=CLIC1 PE=1 SV=4 | 69 | 326 | 13 | 1.06E+08 | 1.06E+08 | 7.33E+07 | 9.03E+07 | 8.58E+07 | 8.89E+07 | 1.04E+08 | 9.73E+07 | 7.42E+07 | 8.60E+07 | 7.60E+07 | 7.32E+07 | 9.18E+07 | 8.52E+07 | 1.08 | 0.11 | 0.391998483 |
| P27105 | Erythrocyte band 7 integral membrane protein OS=Homo sapiens OX=9606 GN=STOM PE=1 SV=3 | 61 | 353 | 13 | 1.05E+08 | 1.14E+08 | 9.16E+07 | 1.33E+08 | 1.07E+08 | 9.77E+07 | 9.19E+07 | 9.65E+07 | 9.31E+07 | 8.83E+07 | 9.58E+07 | 1.08E+08 | 1.08E+08 | 9.56E+07 | 1.13 | 0.18 | 0.094139126 |
| O14980 | Exportin-1 OS=Homo sapiens OX=9606 GN=XPO1 PE=1 SV=1 | 21 | 209 | 13 | 4.68E+07 | 4.47E+07 | 4.53E+07 | 4.57E+07 | 5.30E+07 | 4.11E+07 | 3.25E+07 | 2.32E+07 | 2.40E+07 | 2.56E+07 | 2.85E+07 | 2.17E+07 | 4.61E+07 | 2.59E+07 | 1.78 | 0.83 | 4.48877E-06 |
| Q08380 | Galectin-3-binding protein OS=Homo sapiens OX=9606 GN=LGALS3BP PE=1 SV=1 | 26 | 605 | 13 | 1.88E+08 | 1.93E+08 | 1.44E+08 | 1.56E+08 | 1.43E+08 | 1.57E+08 | 2.39E+08 | 2.15E+08 | 2.02E+08 | 1.87E+08 | 1.81E+08 | 1.79E+08 | 1.63E+08 | 2.01E+08 | 0.81 | -0.30 | 0.016795738 |
| P05121 | Plasminogen activator inhibitor 1 OS=Homo sapiens OX=9606 GN=SERPINE1 PE=1 SV=1 | 39 | 389 | 13 | 1.16E+08 | 1.21E+08 | 1.11E+08 | 1.04E+08 | 1.04E+08 | 1.13E+08 | 1.66E+08 | 1.60E+08 | 1.65E+08 | 1.35E+08 | 1.54E+08 | 1.10E+08 | 1.11E+08 | 1.48E+08 | 0.75 | -0.41 | 0.00813218 |
| P04792 | Heat shock protein beta-1 OS=Homo sapiens OX=9606 GN=HSPB1 PE=1 SV=2 | 75 | 371 | 13 | 1.22E+08 | 1.24E+08 | 9.38E+07 | 9.80E+07 | 8.79E+07 | 9.16E+07 | 8.13E+07 | 8.75E+07 | 6.49E+07 | 6.39E+07 | 6.08E+07 | 6.28E+07 | 1.03E+08 | 7.02E+07 | 1.47 | 0.55 | 0.002675706 |
| Q14254 | Flotillin-2 OS=Homo sapiens OX=9606 GN=FLOT2 PE=1 SV=2 | 35 | 169 | 13 | 2.30E+07 | 3.36E+07 | 3.18E+07 | 3.16E+07 | 2.73E+07 | 3.60E+07 | 4.88E+07 | 5.50E+07 | 4.68E+07 | 4.03E+07 | 4.07E+07 | 4.53E+07 | 3.05E+07 | 4.61E+07 | 0.66 | -0.60 | 0.000376662 |
| P12259 | Coagulation factor V OS=Homo sapiens OX=9606 GN=F5 PE=1 SV=4 | 6 | 583 | 13 | 2.64E+08 | 2.73E+08 | 2.70E+08 | 2.61E+08 | 2.96E+08 | 2.73E+08 | 3.12E+08 | 3.12E+08 | 2.66E+08 | 3.20E+08 | 3.04E+08 | 2.47E+08 | 2.73E+08 | 2.93E+08 | 0.93 | -0.10 | 0.16884312 |
| O00567 | Nucleolar protein 56 OS=Homo sapiens OX=9606 GN=NOP56 PE=1 SV=4 | 25 | 159 | 13 | 5.71E+07 | 5.30E+07 | 4.29E+07 | 5.53E+07 | 4.92E+07 | 5.08E+07 | 3.90E+07 | 3.44E+07 | 3.24E+07 | 3.04E+07 | 3.19E+07 | 2.87E+07 | 5.14E+07 | 3.28E+07 | 1.57 | 0.65 | 4.16516E-05 |
| P04114 | Apolipoprotein B-100 OS=Homo sapiens OX=9606 GN=APOB PE=1 SV=2 | 3 | 448 | 13 | 1.12E+08 | 1.24E+08 | 1.40E+08 | 1.30E+08 | 1.30E+08 | 1.10E+08 | 1.62E+08 | 1.56E+08 | 1.79E+08 | 1.77E+08 | 1.66E+08 | 1.40E+08 | 1.24E+08 | 1.63E+08 | 0.76 | -0.40 | 0.000466927 |

### Supplementary Table 4. List of all proteins identified by label free proteomics analysis.

| Accession | Description | Coverage [%] | # PSMs | # Unique Peptides | Amitriptyline 1 | Amitriptyline 2 | Amitriptyline 3 | Amitriptyline 4 | Amitriptyline 5 | Amitriptyline 6 | Vehicle 1 | Vehicle 2 | Vehicle 3 | Vehicle 4 | Vehicle 5 | Vehicle 6 | Average amitriptyline | Average vehicle | Ratio amitriptyline/ vehicle | Log 2 fold change (amitriptyline /vehicle) | P-value |
|---|---|---|---|---|---|---|---|---|---|---|---|---|---|---|---|---|---|---|---|---|---|
| Q9Y3I0 | tRNA-splicing ligase RtcB homolog OS=Homo sapiens OX=9606 GN=RTCB PE=1 SV=1 | 39 | 174 | 13 | 4.43E+07 | 4.57E+07 | 4.55E+07 | 4.29E+07 | 3.63E+07 | 4.20E+07 | 3.60E+07 | 3.78E+07 | 3.16E+07 | 2.88E+07 | 3.30E+07 | 2.99E+07 | 4.28E+07 | 3.28E+07 | 1.30 | 0.38 | 0.000582275 |
| O00622 | CCN family member 1 OS=Homo sapiens OX=9606 GN=CCN1 PE=1 SV=1 | 36 | 309 | 13 | 6.61E+07 | 5.16E+07 | 5.93E+07 | 6.32E+07 | 6.69E+07 | 6.31E+07 | 1.39E+08 | 1.39E+08 | 1.38E+08 | 1.49E+08 | 1.14E+08 | 1.28E+08 | 6.17E+07 | 1.35E+08 | 0.46 | -1.13 | 2.82828E-06 |
| O95373 | Importin-7 OS=Homo sapiens OX=9606 GN=IPO7 PE=1 SV=1 | 17 | 108 | 13 | 2.99E+07 | 3.47E+07 | 2.87E+07 | 2.90E+07 | 3.48E+07 | 3.36E+07 | 2.41E+07 | 1.89E+07 | 2.26E+07 | 1.96E+07 | 2.61E+07 | 2.17E+07 | 3.18E+07 | 2.22E+07 | 1.43 | 0.52 | 0.000149478 |
| O00231 | 26S proteasome non-ATPase regulatory subunit 11 OS=Homo sapiens OX=9606 GN=PSMD11 PE=1 SV=3 | 44 | 282 | 13 | 5.68E+07 | 5.65E+07 | 5.48E+07 | 5.72E+07 | 5.47E+07 | 5.32E+07 | 7.87E+07 | 4.10E+07 | 3.80E+07 | 3.92E+07 | 3.78E+07 | 3.19E+07 | 5.55E+07 | 4.44E+07 | 1.25 | 0.32 | 0.171866531 |
| Q15063 | Periostin OS=Homo sapiens OX=9606 GN=POSTN PE=1 SV=2 | 19 | 142 | 13 | 3.71E+07 | 3.23E+07 | 3.30E+07 | 2.89E+07 | 2.73E+07 | 3.27E+07 | 5.11E+07 | 4.86E+07 | 3.23E+07 | 3.92E+07 | 4.29E+07 | 3.78E+07 | 3.19E+07 | 4.20E+07 | 0.76 | -0.40 | 0.015345511 |
| Q99460 | 26S proteasome non-ATPase regulatory subunit 1 OS=Homo sapiens OX=9606 GN=PSMD1 PE=1 SV=2 | 21 | 166 | 13 | 4.70E+07 | 5.05E+07 | 4.69E+07 | 4.76E+07 | 4.21E+07 | 4.40E+07 | 3.77E+07 | 3.78E+07 | 2.97E+07 | 3.23E+07 | 3.17E+07 | 2.58E+07 | 4.64E+07 | 3.25E+07 | 1.43 | 0.51 | 0.000219888 |
| P55786 | Puromycin-sensitive aminopeptidase OS=Homo sapiens OX=9606 GN=NPEPPS PE=1 SV=2 | 21 | 168 | 13 | 3.89E+07 | 4.96E+07 | 3.77E+07 | 4.32E+07 | 3.92E+07 | 4.45E+07 | 4.48E+07 | 4.57E+07 | 4.64E+07 | 4.05E+07 | 4.73E+07 | 3.96E+07 | 4.22E+07 | 4.40E+07 | 0.96 | -0.06 | 0.430260256 |
| P12429 | Annexin A3 OS=Homo sapiens OX=9606 GN=ANXA3 PE=1 SV=3 | 44 | 233 | 13 | 7.70E+07 | 7.38E+07 | 6.16E+07 | 6.67E+07 | 7.53E+07 | 6.37E+07 | 6.64E+07 | 6.68E+07 | 6.38E+07 | 5.80E+07 | 5.76E+07 | 4.89E+07 | 6.97E+07 | 6.03E+07 | 1.16 | 0.21 | 0.035041928 |
| Q9Y230 | RuvB-like 2 OS=Homo sapiens OX=9606 GN=RUVBL2 PE=1 SV=3 | 30 | 152 | 13 | 4.60E+07 | 4.63E+07 | 4.38E+07 | 4.80E+07 | 5.04E+07 | 4.38E+07 | 3.20E+07 | 3.20E+07 | 2.90E+07 | 3.15E+07 | 2.89E+07 | 2.48E+07 | 4.64E+07 | 2.97E+07 | 1.56 | 0.64 | 7.89841E-07 |
| Q86UX7 | Fermitin family homolog 3 OS=Homo sapiens OX=9606 GN=FERMT3 PE=1 SV=1 | 22 | 216 | 13 | 5.65E+07 | 6.21E+07 | 5.95E+07 | 6.33E+07 | 5.82E+07 | 5.49E+07 | 8.40E+07 | 7.88E+07 | 7.89E+07 | 7.84E+07 | 7.93E+07 | 7.97E+07 | 5.91E+07 | 7.98E+07 | 0.74 | -0.43 | 5.14872E-07 |
| P60228 | Eukaryotic translation initiation factor 3 subunit E OS=Homo sapiens OX=9606 GN=EIF3E PE=1 SV=1 | 35 | 209 | 13 | 5.87E+07 | 6.31E+07 | 6.35E+07 | 6.71E+07 | 6.77E+07 | 5.83E+07 | 4.81E+07 | 3.68E+07 | 4.52E+07 | 4.42E+07 | 4.50E+07 | 4.00E+07 | 6.31E+07 | 4.32E+07 | 1.46 | 0.55 | 6.81537E-06 |
| P62906 | 60S ribosomal protein L10a OS=Homo sapiens OX=9606 GN=RPL10A PE=1 SV=2 | 62 | 358 | 14 | 2.20E+08 | 2.45E+08 | 2.28E+08 | 2.40E+08 | 2.31E+08 | 2.09E+08 | 1.80E+08 | 1.81E+08 | 1.54E+08 | 1.60E+08 | 1.59E+08 | 1.50E+08 | 2.29E+08 | 1.64E+08 | 1.40 | 0.48 | 7.01911E-06 |
| Q14974 | Importin subunit beta-1 OS=Homo sapiens OX=9606 GN=KPNB1 PE=1 SV=2 | 24 | 282 | 14 | 7.64E+07 | 7.15E+07 | 6.88E+07 | 7.08E+07 | 7.82E+07 | 6.72E+07 | 6.22E+07 | 4.97E+07 | 4.71E+07 | 4.24E+07 | 5.28E+07 | 3.88E+07 | 7.22E+07 | 4.88E+07 | 1.48 | 0.56 | 0.000349151 |
| P39023 | 60S ribosomal protein L3 OS=Homo sapiens OX=9606 GN=RPL3 PE=1 SV=2 | 43 | 274 | 14 | 1.15E+08 | 1.15E+08 | 1.16E+08 | 1.11E+08 | 1.16E+08 | 1.11E+08 | 7.18E+07 | 6.26E+07 | 5.41E+07 | 6.76E+07 | 5.72E+07 | 5.83E+07 | 1.14E+08 | 6.19E+07 | 1.84 | 0.88 | 1.36959E-06 |
| P11021 | Endoplasmic reticulum chaperone BiP OS=Homo sapiens OX=9606 GN=HSPA5 PE=1 SV=2 | 31 | 323 | 14 | 4.66E+07 | 4.68E+07 | 4.07E+07 | 3.68E+07 | 3.89E+07 | 3.58E+07 | 4.08E+07 | 3.79E+07 | 2.87E+07 | 3.07E+07 | 2.97E+07 | 3.20E+07 | 4.09E+07 | 3.33E+07 | 1.23 | 0.30 | 0.021029702 |
| P06493 | Cyclin-dependent kinase 1 OS=Homo sapiens OX=9606 GN=CDK1 PE=1 SV=3 | 56 | 238 | 14 | 8.99E+07 | 9.42E+07 | 8.03E+07 | 8.18E+07 | 9.00E+07 | 8.91E+07 | 4.63E+07 | 5.46E+07 | 4.57E+07 | 4.38E+07 | 5.08E+07 | 4.92E+07 | 8.75E+07 | 4.84E+07 | 1.81 | 0.85 | 1.3434E-07 |
| P62753 | 40S ribosomal protein S6 OS=Homo sapiens OX=9606 GN=RPS6 PE=1 SV=1 | 40 | 309 | 14 | 1.44E+08 | 1.61E+08 | 1.31E+08 | 1.36E+08 | 1.22E+08 | 1.37E+08 | 7.79E+07 | 9.17E+07 | 6.42E+07 | 7.97E+07 | 6.74E+07 | 6.11E+07 | 1.38E+08 | 7.37E+07 | 1.88 | 0.91 | 4.80964E-06 |
| P55060 | Exportin-2 OS=Homo sapiens OX=9606 GN=CSE1L PE=1 SV=3 | 22 | 168 | 14 | 4.61E+07 | 4.33E+07 | 4.85E+07 | 4.60E+07 | 5.06E+07 | 4.75E+07 | 3.38E+07 | 3.37E+07 | 3.00E+07 | 3.48E+07 | 3.39E+07 | 2.71E+07 | 4.70E+07 | 3.22E+07 | 1.46 | 0.54 | 3.9913E-06 |
| P20591 | Interferon-induced GTP-binding protein Mx1 OS=Homo sapiens OX=9606 GN=MX1 PE=1 SV=4 | 32 | 182 | 14 | 6.06E+07 | 5.34E+07 | 5.73E+07 | 5.24E+07 | 4.74E+07 | 5.23E+07 | 2.31E+07 | 2.33E+07 | 2.02E+07 | 1.88E+07 | 1.76E+07 | 2.21E+07 | 5.39E+07 | 2.08E+07 | 2.59 | 1.37 | 5.1142E-07 |
| P46777 | 60S ribosomal protein L5 OS=Homo sapiens OX=9606 GN=RPL5 PE=1 SV=3 | 47 | 322 | 14 | 1.86E+08 | 1.61E+08 | 1.38E+08 | 1.51E+08 | 1.49E+08 | 1.48E+08 | 1.13E+08 | 1.11E+08 | 9.69E+07 | 1.02E+08 | 9.48E+07 | 8.27E+07 | 1.56E+08 | 1.00E+08 | 1.55 | 0.64 | 8.30677E-05 |
| Q9H223 | EH domain-containing protein 4 OS=Homo sapiens OX=9606 GN=EHD4 PE=1 SV=1 | 42 | 198 | 14 | 5.51E+07 | 5.49E+07 | 4.75E+07 | 4.87E+07 | 4.77E+07 | 4.95E+07 | 3.91E+07 | 4.00E+07 | 3.70E+07 | 3.09E+07 | 3.21E+07 | 3.49E+07 | 5.05E+07 | 3.57E+07 | 1.42 | 0.50 | 3.27392E-05 |
| P38606 | V-type proton ATPase catalytic subunit A OS=Homo sapiens OX=9606 GN=ATP6V1A PE=1 SV=2 | 38 | 145 | 14 | 3.11E+07 | 3.34E+07 | 3.46E+07 | 3.17E+07 | 2.93E+07 | 3.07E+07 | 3.43E+07 | 3.31E+07 | 3.03E+07 | 2.71E+07 | 3.14E+07 | 2.75E+07 | 3.18E+07 | 3.06E+07 | 1.04 | 0.06 | 0.423305356 |
| Q92626 | Peroxidasin homolog OS=Homo sapiens OX=9606 GN=PXDN PE=1 SV=2 | 15 | 136 | 14 | 1.61E+07 | 1.48E+07 | 1.57E+07 | 1.62E+07 | 1.88E+07 | 1.84E+07 | 3.94E+07 | 2.93E+07 | 2.76E+07 | 2.46E+07 | 2.55E+07 | 2.81E+07 | 1.67E+07 | 2.91E+07 | 0.57 | -0.80 | 0.001656955 |
| O00410 | Importin-5 OS=Homo sapiens OX=9606 GN=IPO5 PE=1 SV=4 | 22 | 120 | 14 | 4.54E+07 | 3.60E+07 | 2.72E+07 | 3.32E+07 | 2.95E+07 | 2.81E+07 | 2.72E+07 | 1.52E+07 | 2.04E+07 | 1.63E+07 | 1.90E+07 | 1.51E+07 | 3.32E+07 | 1.89E+07 | 1.76 | 0.82 | 0.002178959 |
| P14625 | Endoplasmin OS=Homo sapiens OX=9606 GN=HSP90B1 PE=1 SV=1 | 22 | 317 | 14 | 4.28E+07 | 4.22E+07 | 3.45E+07 | 4.00E+07 | 3.74E+07 | 3.51E+07 | 4.55E+07 | 4.58E+07 | 4.23E+07 | 4.56E+07 | 4.30E+07 | 3.85E+07 | 3.87E+07 | 4.35E+07 | 0.89 | -0.17 | 0.027812594 |
| O43854 | EGF-like repeat and discoidin I-like domain-containing protein 3 OS=Homo sapiens OX=9606 GN=EDIL3 PE=1 SV=1 | 35 | 185 | 14 | 8.84E+07 | 9.20E+07 | 7.83E+07 | 9.00E+07 | 8.49E+07 | 7.63E+07 | 1.09E+08 | 1.12E+08 | 1.09E+08 | 1.15E+08 | 9.30E+07 | 8.98E+07 | 8.50E+07 | 1.05E+08 | 0.81 | -0.30 | 0.004109821 |
| P35998 | 26S proteasome regulatory subunit 7 OS=Homo sapiens OX=9606 GN=PSMC2 PE=1 SV=3 | 42 | 161 | 14 | 4.70E+07 | 4.55E+07 | 3.77E+07 | 4.42E+07 | 3.87E+07 | 3.77E+07 | 3.36E+07 | 2.94E+07 | 3.15E+07 | 3.09E+07 | 2.28E+07 | 1.94E+07 | 4.18E+07 | 2.80E+07 | 1.50 | 0.58 | 0.000815204 |
| Q5SSJ5 | Heterochromatin protein 1-binding protein 3 OS=Homo sapiens OX=9606 GN=HP1BP3 PE=1 SV=1 | 26 | 148 | 14 | 4.52E+07 | 4.14E+07 | 4.68E+07 | 4.68E+07 | 4.59E+07 | 4.16E+07 | 3.21E+07 | 3.06E+07 | 3.33E+07 | 3.32E+07 | 3.16E+07 | 2.17E+07 | 4.46E+07 | 3.04E+07 | 1.47 | 0.55 | 0.000125567 |
| P62280 | 40S ribosomal protein S11 OS=Homo sapiens OX=9606 GN=RPS11 PE=1 SV=3 | 63 | 350 | 14 | 1.70E+08 | 1.70E+08 | 1.69E+08 | 1.71E+08 | 1.75E+08 | 1.52E+08 | 1.23E+08 | 1.26E+08 | 1.24E+08 | 1.24E+08 | 1.11E+08 | 9.68E+07 | 1.68E+08 | 1.18E+08 | 1.43 | 0.51 | 1.09494E-05 |
| P11388 | DNA topoisomerase 2-alpha OS=Homo sapiens OX=9606 GN=TOP2A PE=1 SV=3 | 12 | 94 | 14 | 2.05E+07 | 2.16E+07 | 2.21E+07 | 2.14E+07 | 1.98E+07 | 1.91E+07 | 1.93E+07 | 1.37E+07 | 1.61E+07 | 1.73E+07 | 1.60E+07 | 1.03E+07 | 2.08E+07 | 1.54E+07 | 1.34 | 0.43 | 0.006887621 |
| P07942 | Laminin subunit beta-1 OS=Homo sapiens OX=9606 GN=LAMB1 PE=1 SV=2 | 10 | 133 | 14 | 2.07E+07 | 2.39E+07 | 2.72E+07 | 2.58E+07 | 2.45E+07 | 2.63E+07 | 3.64E+07 | 3.49E+07 | 3.43E+07 | 3.24E+07 | 3.08E+07 | 2.67E+07 | 2.47E+07 | 3.26E+07 | 0.76 | -0.40 | 0.00142394 |
| P62826 | GTP-binding nuclear protein Ran OS=Homo sapiens OX=9606 GN=RAN PE=1 SV=3 | 49 | 719 | 15 | 4.54E+08 | 5.14E+08 | 4.04E+08 | 4.17E+08 | 4.32E+08 | 4.24E+08 | 3.87E+08 | 4.25E+08 | 4.09E+08 | 3.57E+08 | 3.54E+08 | 3.14E+08 | 4.41E+08 | 3.74E+08 | 1.18 | 0.24 | 0.01683434 |
| Q16555 | Dihydropyrimidinase-related protein 2 OS=Homo sapiens OX=9606 GN=DPYSL2 PE=1 SV=1 | 46 | 280 | 15 | 6.41E+07 | 6.79E+07 | 6.25E+07 | 5.95E+07 | 5.68E+07 | 6.28E+07 | 6.77E+07 | 7.03E+07 | 7.12E+07 | 6.42E+07 | 5.79E+07 | 5.96E+07 | 6.23E+07 | 6.52E+07 | 0.96 | -0.07 | 0.318854539 |
| P62241 | 40S ribosomal protein S8 OS=Homo sapiens OX=9606 GN=RPS8 PE=1 SV=2 | 60 | 826 | 15 | 3.14E+08 | 3.23E+08 | 3.32E+08 | 3.12E+08 | 3.21E+08 | 3.45E+08 | 2.09E+08 | 2.20E+08 | 1.95E+08 | 2.02E+08 | 1.98E+08 | 1.75E+08 | 3.25E+08 | 2.00E+08 | 1.62 | 0.70 | 4.09199E-08 |
| P62424 | 60S ribosomal protein L7a OS=Homo sapiens OX=9606 GN=RPL7A PE=1 SV=2 | 51 | 367 | 15 | 1.63E+08 | 1.58E+08 | 1.57E+08 | 1.65E+08 | 1.50E+08 | 1.38E+08 | 9.09E+07 | 1.00E+08 | 8.18E+07 | 9.68E+07 | 8.98E+07 | 7.82E+07 | 1.55E+08 | 8.97E+07 | 1.73 | 0.79 | 2.79207E-07 |
| Q9NZM1 | Myoferlin OS=Homo sapiens OX=9606 GN=MYOF PE=1 SV=1 | 11 | 174 | 15 | 4.47E+07 | 4.29E+07 | 3.88E+07 | 4.10E+07 | 3.92E+07 | 3.94E+07 | 3.04E+07 | 2.42E+07 | 2.81E+07 | 2.69E+07 | 2.43E+07 | 2.10E+07 | 4.10E+07 | 2.58E+07 | 1.59 | 0.67 | 7.72441E-06 |
| Q9UHB9 | Signal recognition particle subunit SRP68 OS=Homo sapiens OX=9606 GN=SRP68 PE=1 SV=2 | 32 | 139 | 15 | 4.23E+07 | 4.33E+07 | 3.87E+07 | 3.92E+07 | 3.53E+07 | 3.89E+07 | 2.15E+07 | 1.72E+07 | 1.62E+07 | 1.78E+07 | 1.78E+07 | 1.53E+07 | 3.96E+07 | 1.76E+07 | 2.24 | 1.17 | 7.87903E-08 |
| O75955 | Flotillin-1 OS=Homo sapiens OX=9606 GN=FLOT1 PE=1 SV=3 | 38 | 175 | 15 | 3.40E+07 | 3.62E+07 | 3.73E+07 | 4.10E+07 | 4.21E+07 | 3.79E+07 | 5.37E+07 | 5.26E+07 | 5.08E+07 | 5.00E+07 | 4.72E+07 | 4.49E+07 | 3.81E+07 | 4.99E+07 | 0.76 | -0.39 | 7.60348E-05 |
| O75891 | Cytosolic 10-formyltetrahydrofolate dehydrogenase OS=Homo sapiens OX=9606 GN=ALDH1L1 PE=1 SV=2 | 19 | 425 | 15 | 1.97E+08 | 2.08E+08 | 1.74E+08 | 1.85E+08 | 1.70E+08 | 1.63E+08 | 2.51E+08 | 2.65E+08 | 2.09E+08 | 2.36E+08 | 2.15E+08 | 2.03E+08 | 1.83E+08 | 2.30E+08 | 0.80 | -0.33 | 0.004454169 |
| P02786 | Transferrin receptor protein 1 OS=Homo sapiens OX=9606 GN=TFRC PE=1 SV=2 | 24 | 313 | 15 | 7.51E+07 | 7.97E+07 | 7.55E+07 | 7.74E+07 | 6.29E+07 | 6.89E+07 | 8.12E+07 | 6.92E+07 | 9.12E+07 | 9.10E+07 | 9.38E+07 | 5.93E+07 | 7.32E+07 | 8.10E+07 | 0.90 | -0.14 | 0.25765062 |
| Q9Y6K5 | 2'-5'-oligoadenylate synthase 3 OS=Homo sapiens OX=9606 GN=OAS3 PE=1 SV=3 | 22 | 134 | 15 | 3.30E+07 | 3.44E+07 | 2.69E+07 | 3.10E+07 | 2.87E+07 | 2.98E+07 | 2.43E+07 | 2.10E+07 | 1.92E+07 | 1.83E+07 | 2.02E+07 | 1.56E+07 | 3.06E+07 | 1.98E+07 | 1.55 | 0.63 | 5.8463E-05 |
| Q9UNM6 | 26S proteasome non-ATPase regulatory subunit 13 OS=Homo sapiens OX=9606 GN=PSMD13 PE=1 SV=2 | 45 | 180 | 15 | 6.48E+07 | 6.22E+07 | 6.07E+07 | 6.30E+07 | 6.85E+07 | 5.67E+07 | 4.65E+07 | 4.48E+07 | 4.85E+07 | 5.50E+07 | 4.34E+07 | 3.85E+07 | 6.27E+07 | 4.61E+07 | 1.36 | 0.44 | 0.000198056 |
| Q15393 | Splicing factor 3B subunit 3 OS=Homo sapiens OX=9606 GN=SF3B3 PE=1 SV=4 | 15 | 180 | 15 | 7.23E+07 | 7.29E+07 | 6.35E+07 | 6.96E+07 | 6.52E+07 | 6.38E+07 | 4.27E+07 | 4.36E+07 | 4.69E+07 | 4.33E+07 | 3.89E+07 | 3.81E+07 | 6.79E+07 | 4.23E+07 | 1.61 | 0.68 | 6.47687E-07 |
| P14868 | Aspartate--tRNA ligase, cytoplasmic OS=Homo sapiens OX=9606 GN=DARS PE=1 SV=2 | 33 | 282 | 15 | 9.72E+07 | 1.06E+08 | 8.81E+07 | 8.51E+07 | 9.17E+07 | 8.53E+07 | 8.01E+07 | 6.84E+07 | 7.53E+07 | 6.55E+07 | 7.07E+07 | 5.23E+07 | 9.22E+07 | 6.87E+07 | 1.34 | 0.42 | 0.001086047 |
| O43242 | 26S proteasome non-ATPase regulatory subunit 3 OS=Homo sapiens OX=9606 GN=PSMD3 PE=1 SV=2 | 36 | 207 | 15 | 4.80E+07 | 5.86E+07 | 5.84E+07 | 4.98E+07 | 4.68E+07 | 5.16E+07 | 3.38E+07 | 3.84E+07 | 3.65E+07 | 3.87E+07 | 3.91E+07 | 3.31E+07 | 5.22E+07 | 3.66E+07 | 1.43 | 0.51 | 0.000234261 |
| P43686 | 26S proteasome regulatory subunit 6B OS=Homo sapiens OX=9606 GN=PSMC4 PE=1 SV=2 | 53 | 319 | 16 | 7.50E+07 | 7.84E+07 | 7.71E+07 | 7.54E+07 | 8.29E+07 | 7.85E+07 | 5.32E+07 | 5.85E+07 | 5.37E+07 | 5.11E+07 | 5.49E+07 | 4.76E+07 | 7.79E+07 | 5.32E+07 | 1.46 | 0.55 | 2.35414E-07 |
| P30153 | Serine/threonine-protein phosphatase 2A 65 kDa regulatory subunit A alpha isoform OS=Homo sapiens OX=9606 GN=PPP2R1A PE=1 SV=4 | 31 | 219 | 16 | 7.16E+07 | 5.21E+07 | 7.03E+07 | 6.15E+07 | 5.44E+07 | 6.33E+07 | 6.39E+07 | 5.85E+07 | 5.35E+07 | 5.37E+07 | 6.96E+07 | 5.29E+07 | 6.22E+07 | 5.87E+07 | 1.06 | 0.08 | 0.429990125 |
| P54136 | Arginine--tRNA ligase, cytoplasmic OS=Homo sapiens OX=9606 GN=RARS PE=1 SV=2 | 26 | 193 | 16 | 5.75E+07 | 6.11E+07 | 5.41E+07 | 5.27E+07 | 5.84E+07 | 5.20E+07 | 3.65E+07 | 3.79E+07 | 2.92E+07 | 3.25E+07 | 3.36E+07 | 2.71E+07 | 5.60E+07 | 3.28E+07 | 1.71 | 0.77 | 1.34381E-06 |
| P26038 | Moesin OS=Homo sapiens OX=9606 GN=MSN PE=1 SV=3 | 41 | 346 | 16 | 1.34E+08 | 1.31E+08 | 1.01E+08 | 1.05E+08 | 9.52E+07 | 1.01E+08 | 1.07E+08 | 1.06E+08 | 8.74E+07 | 8.81E+07 | 8.25E+07 | 8.84E+07 | 1.11E+08 | 9.32E+07 | 1.19 | 0.26 | 0.053169262 |
| Q01813 | ATP-dependent 6-phosphofructokinase, platelet type OS=Homo sapiens OX=9606 GN=PFKP PE=1 SV=2 | 26 | 199 | 16 | 7.24E+07 | 7.37E+07 | 6.31E+07 | 6.70E+07 | 5.99E+07 | 6.55E+07 | 5.45E+07 | 5.24E+07 | 4.80E+07 | 4.93E+07 | 5.65E+07 | 4.48E+07 | 6.69E+07 | 5.09E+07 | 1.31 | 0.39 | 0.000227945 |
| P41250 | Glycine--tRNA ligase OS=Homo sapiens OX=9606 GN=GARS1 PE=1 SV=3 | 29 | 132 | 16 | 4.50E+07 | 4.63E+07 | 4.81E+07 | 4.60E+07 | 4.20E+07 | 4.48E+07 | 3.43E+07 | 2.82E+07 | 2.75E+07 | 2.85E+07 | 2.77E+07 | 2.47E+07 | 4.53E+07 | 2.85E+07 | 1.59 | 0.67 | 2.40513E-06 |
| P23921 | Ribonucleoside-diphosphate reductase large subunit OS=Homo sapiens OX=9606 GN=RRM1 PE=1 SV=1 | 28 | 216 | 16 | 6.58E+07 | 6.47E+07 | 5.27E+07 | 5.71E+07 | 5.95E+07 | 5.04E+07 | 7.02E+07 | 6.32E+07 | 6.82E+07 | 6.70E+07 | 5.59E+07 | 6.68E+07 | 5.84E+07 | 6.52E+07 | 0.89 | -0.16 | 0.064327183 |
| P34897 | Serine hydroxymethyltransferase, mitochondrial OS=Homo sapiens OX=9606 GN=SHMT2 PE=1 SV=3 | 47 | 310 | 16 | 1.14E+08 | 1.28E+08 | 1.04E+08 | 1.07E+08 | 1.08E+08 | 1.10E+08 | 8.45E+07 | 9.95E+07 | 7.82E+07 | 8.56E+07 | 8.80E+07 | 7.78E+07 | 1.12E+08 | 8.56E+07 | 1.31 | 0.39 | 0.000280862 |
| P21980 | Protein-glutamine gamma-glutamyltransferase 2 OS=Homo sapiens OX=9606 GN=TGM2 PE=1 SV=2 | 33 | 275 | 16 | 7.16E+07 | 5.84E+07 | 5.17E+07 | 5.69E+07 | 5.78E+07 | 5.73E+07 | 4.59E+07 | 4.60E+07 | 3.57E+07 | 3.68E+07 | 4.94E+07 | 2.91E+07 | 5.90E+07 | 4.05E+07 | 1.46 | 0.54 | 0.001392039 |
| Q99613 | Eukaryotic translation initiation factor 3 subunit C OS=Homo sapiens OX=9606 GN=EIF3C PE=1 SV=1 | 18 | 288 | 16 | 7.68E+07 | 7.62E+07 | 7.21E+07 | 7.61E+07 | 7.00E+07 | 6.81E+07 | 4.90E+07 | 4.55E+07 | 4.10E+07 | 4.48E+07 | 3.76E+07 | 3.46E+07 | 7.32E+07 | 4.21E+07 | 1.74 | 0.80 | 1.14604E-06 |
| P49588 | Alanine--tRNA ligase, cytoplasmic OS=Homo sapiens OX=9606 GN=AARS PE=1 SV=2 | 22 | 231 | 16 | 6.97E+07 | 6.58E+07 | 7.05E+07 | 7.06E+07 | 7.12E+07 | 6.47E+07 | 4.60E+07 | 4.87E+07 | 4.47E+07 | 5.01E+07 | 4.85E+07 | 4.12E+07 | 6.87E+07 | 4.65E+07 | 1.48 | 0.56 | 2.26638E-07 |
| P24821 | Tenascin OS=Homo sapiens OX=9606 GN=TNC PE=1 SV=3 | 11 | 200 | 16 | 1.96E+07 | 1.70E+07 | 2.56E+07 | 3.14E+07 | 2.08E+07 | 2.54E+07 | 5.46E+07 | 5.10E+07 | 5.26E+07 | 4.48E+07 | 4.26E+07 | 4.44E+07 | 2.33E+07 | 4.83E+07 | 0.48 | -1.05 | 6.92534E-06 |
| P05362 | Intercellular adhesion molecule 1 OS=Homo sapiens OX=9606 GN=ICAM1 PE=1 SV=2 | 25 | 306 | 16 | 1.23E+08 | 1.26E+08 | 1.26E+08 | 1.30E+08 | 1.12E+08 | 1.21E+08 | 7.09E+07 | 8.23E+07 | 6.40E+07 | 6.27E+07 | 5.83E+07 | 5.82E+07 | 1.23E+08 | 6.61E+07 | 1.86 | 0.90 | 6.72191E-07 |
| O43776 | Asparagine--tRNA ligase, cytoplasmic OS=Homo sapiens OX=9606 GN=NARS1 PE=1 SV=1 | 30 | 187 | 16 | 6.19E+07 | 6.22E+07 | 5.71E+07 | 6.09E+07 | 6.18E+07 | 5.35E+07 | 5.16E+07 | 4.70E+07 | 4.68E+07 | 5.10E+07 | 4.30E+07 | 3.45E+07 | 5.96E+07 | 4.56E+07 | 1.31 | 0.38 | 0.001570899 |
| P29317 | Ephrin type-A receptor 2 OS=Homo sapiens OX=9606 GN=EPHA2 PE=1 SV=2 | 18 | 186 | 16 | 5.57E+07 | 5.87E+07 | 5.38E+07 | 6.14E+07 | 5.78E+07 | 5.49E+07 | 4.08E+07 | 4.15E+07 | 4.29E+07 | 4.53E+07 | 4.40E+07 | 3.62E+07 | 5.71E+07 | 4.18E+07 | 1.37 | 0.45 | 5.66128E-06 |
| P02545 | Prelamin-A/C OS=Homo sapiens OX=9606 GN=LMNA PE=1 SV=1 | 28 | 169 | 16 | 5.94E+07 | 6.30E+07 | 6.27E+07 | 7.17E+07 | 6.45E+07 | 5.95E+07 | 3.80E+07 | 3.95E+07 | 3.95E+07 | 4.47E+07 | 3.95E+07 | 3.14E+07 | 6.35E+07 | 3.88E+07 | 1.64 | 0.71 | 2.08857E-06 |
| P42224 | Signal transducer and activator of transcription 1-alpha/beta OS=Homo sapiens OX=9606 GN=STAT1 PE=1 SV=2 | 28 | 185 | 16 | 4.36E+07 | 4.76E+07 | 4.12E+07 | 3.82E+07 | 4.69E+07 | 3.63E+07 | 3.51E+07 | 3.53E+07 | 2.80E+07 | 2.81E+07 | 2.84E+07 | 2.33E+07 | 4.23E+07 | 2.97E+07 | 1.42 | 0.51 | 0.000833302 |
| P21589 | 5'-nucleotidase OS=Homo sapiens OX=9606 GN=NT5E PE=1 SV=1 | 41 | 398 | 17 | 1.63E+08 | 1.53E+08 | 1.30E+08 | 1.28E+08 | 1.15E+08 | 1.33E+08 | 1.08E+08 | 9.56E+07 | 7.62E+07 | 7.37E+07 | 8.00E+07 | 7.90E+07 | 1.37E+08 | 8.54E+07 | 1.60 | 0.68 | 0.000237758 |
| P36578 | 60S ribosomal protein L4 OS=Homo sapiens OX=9606 GN=RPL4 PE=1 SV=5 | 39 | 429 | 17 | 1.72E+08 | 1.76E+08 | 1.43E+08 | 1.54E+08 | 1.38E+08 | 1.57E+08 | 7.92E+07 | 8.11E+07 | 6.72E+07 | 6.71E+07 | 6.46E+07 | 7.49E+07 | 1.57E+08 | 7.24E+07 | 2.17 | 1.11 | 5.20448E-06 |
| P0C0L4 | Complement C4-A OS=Homo sapiens OX=9606 GN=C4A PE=1 SV=2 | 11 | 968 | 17 | 5.92E+08 | 6.03E+08 | 5.22E+08 | 5.78E+08 | 5.62E+08 | 4.73E+08 | 5.90E+08 | 6.16E+08 | 5.40E+08 | 5.93E+08 | 4.99E+08 | 5.61E+08 | 5.55E+08 | 5.66E+08 | 0.98 | -0.03 | 0.676630844 |
| P49915 | GMP synthase [glutamine-hydrolyzing] OS=Homo sapiens OX=9606 GN=GMPS PE=1 SV=1 | 30 | 207 | 17 | 5.12E+07 | 5.58E+07 | 4.69E+07 | 4.72E+07 | 4.65E+07 | 4.39E+07 | 3.42E+07 | 3.64E+07 | 3.36E+07 | 3.25E+07 | 3.48E+07 | 2.80E+07 | 4.86E+07 | 3.33E+07 | 1.46 | 0.55 | 4.9949E-05 |
| Q96QK1 | Vacuolar protein sorting-associated protein 35 OS=Homo sapiens OX=9606 GN=VPS35 PE=1 SV=2 | 29 | 192 | 17 | 4.67E+07 | 5.10E+07 | 4.89E+07 | 4.20E+07 | 4.93E+07 | 4.31E+07 | 4.66E+07 | 4.13E+07 | 4.28E+07 | 4.01E+07 | 4.14E+07 | 3.29E+07 | 4.68E+07 | 4.08E+07 | 1.15 | 0.20 | 0.029799799 |
| P26006 | Integrin alpha-3 OS=Homo sapiens OX=9606 GN=ITGA3 PE=1 SV=5 | 19 | 317 | 17 | 1.48E+08 | 1.44E+08 | 1.23E+08 | 1.29E+08 | 1.24E+08 | 1.21E+08 | 8.25E+07 | 7.90E+07 | 7.74E+07 | 8.81E+07 | 6.92E+07 | 6.20E+07 | 1.31E+08 | 7.64E+07 | 1.72 | 0.78 | 5.53512E-06 |
| Q9Y678 | Coatomer subunit gamma-1 OS=Homo sapiens OX=9606 GN=COPG1 PE=1 SV=1 | 32 | 163 | 17 | 5.11E+07 | 4.68E+07 | 4.79E+07 | 4.44E+07 | 4.54E+07 | 3.68E+07 | 2.86E+07 | 2.72E+07 | 2.42E+07 | 2.45E+07 | 2.47E+07 | 1.82E+07 | 4.54E+07 | 2.46E+07 | 1.85 | 0.89 | 1.17894E-05 |
| P12111 | Collagen alpha-3(VI) chain OS=Homo sapiens OX=9606 GN=COL6A3 PE=1 SV=5 | 6 | 293 | 17 | 5.72E+07 | 6.40E+07 | 5.34E+07 | 5.48E+07 | 4.79E+07 | 5.71E+07 | 8.72E+07 | 9.68E+07 | 8.09E+07 | 7.42E+07 | 7.49E+07 | 7.96E+07 | 5.57E+07 | 8.23E+07 | 0.68 | -0.56 | 0.00015959 |
| P49748 | Very long-chain specific acyl-CoA dehydrogenase, mitochondrial OS=Homo sapiens OX=9606 GN=ACADVL PE=1 SV=1 | 29 | 174 | 17 | 6.79E+07 | 6.19E+07 | 5.16E+07 | 5.07E+07 | 4.53E+07 | 4.67E+07 | 3.73E+07 | 3.75E+07 | 3.09E+07 | 3.14E+07 | 3.16E+07 | 2.76E+07 | 5.40E+07 | 3.27E+07 | 1.65 | 0.72 | 0.001139755 |
| P18124 | 60S ribosomal protein L7 OS=Homo sapiens OX=9606 GN=RPL7 PE=1 SV=1 | 57 | 391 | 17 | 1.59E+08 | 1.54E+08 | 1.26E+08 | 1.49E+08 | 1.37E+08 | 1.42E+08 | 8.82E+07 | 8.95E+07 | 9.47E+07 | 8.31E+07 | 8.01E+07 | 7.12E+07 | 1.44E+08 | 8.45E+07 | 1.71 | 0.77 | 3.6428E-06 |
| P04083 | Annexin A1 OS=Homo sapiens OX=9606 GN=ANXA1 PE=1 SV=2 | 53 | 523 | 18 | 2.61E+08 | 2.35E+08 | 2.30E+08 | 2.19E+08 | 2.16E+08 | 2.25E+08 | 2.32E+08 | 2.14E+08 | 2.01E+08 | 1.94E+08 | 1.92E+08 | 1.58E+08 | 2.31E+08 | 1.98E+08 | 1.16 | 0.22 | 0.02539966 |
| Q01518 | Adenylyl cyclase-associated protein 1 OS=Homo sapiens OX=9606 GN=CAP1 PE=1 SV=5 | 47 | 368 | 18 | 1.33E+08 | 1.15E+08 | 1.17E+08 | 1.30E+08 | 1.18E+08 | 1.25E+08 | 1.07E+08 | 1.10E+08 | 9.96E+07 | 1.06E+08 | 1.02E+08 | 9.41E+07 | 1.23E+08 | 1.03E+08 | 1.19 | 0.25 | 0.000571893 |
| P26641 | Elongation factor 1-gamma OS=Homo sapiens OX=9606 GN=EEF1G PE=1 SV=3 | 67 | 491 | 18 | 2.15E+08 | 2.12E+08 | 2.06E+08 | 2.04E+08 | 2.08E+08 | 1.98E+08 | 2.09E+08 | 2.10E+08 | 2.02E+08 | 2.22E+08 | 1.97E+08 | 1.74E+08 | 2.07E+08 | 2.02E+08 | 1.02 | 0.03 | 0.515144864 |
| P63244 | Receptor of activated protein C kinase 1 OS=Homo sapiens OX=9606 GN=RACK1 PE=1 SV=3 | 81 | 426 | 18 | 1.58E+08 | 1.39E+08 | 1.24E+08 | 1.19E+08 | 1.18E+08 | 1.36E+08 | 9.15E+07 | 8.32E+07 | 8.59E+07 | 7.43E+07 | 8.02E+07 | 8.36E+07 | 1.32E+08 | 8.31E+07 | 1.59 | 0.67 | 0.000235396 |
| P61158 | Actin-related protein 3 OS=Homo sapiens OX=9606 GN=ACTR3 PE=1 SV=3 | 59 | 274 | 18 | 8.41E+07 | 6.45E+07 | 5.76E+07 | 6.88E+07 | 7.29E+07 | 6.20E+07 | 6.67E+07 | 7.04E+07 | 6.46E+07 | 5.64E+07 | 5.58E+07 | 5.45E+07 | 6.83E+07 | 6.14E+07 | 1.11 | 0.15 | 0.174091944 |
| P61247 | 40S ribosomal protein S3a OS=Homo sapiens OX=9606 GN=RPS3A PE=1 SV=2 | 56 | 333 | 18 | 2.54E+08 | 2.74E+08 | 2.23E+08 | 2.39E+08 | 2.38E+08 | 2.26E+08 | 2.02E+08 | 2.25E+08 | 2.03E+08 | 1.95E+08 | 1.73E+08 | 1.67E+08 | 2.42E+08 | 1.94E+08 | 1.25 | 0.32 | 0.002070637 |
| O75083 | WD repeat-containing protein 1 OS=Homo sapiens OX=9606 GN=WDR1 PE=1 SV=4 | 46 | 529 | 19 | 2.27E+08 | 2.39E+08 | 2.07E+08 | 1.89E+08 | 1.91E+08 | 1.84E+08 | 2.53E+08 | 2.35E+08 | 2.15E+08 | 2.25E+08 | 2.22E+08 | 1.80E+08 | 2.06E+08 | 2.22E+08 | 0.93 | -0.10 | 0.276843875 |
| P00558 | Phosphoglycerate kinase 1 OS=Homo sapiens OX=9606 GN=PGK1 PE=1 SV=3 | 55 | 634 | 19 | 2.78E+08 | 2.88E+08 | 2.29E+08 | 2.47E+08 | 2.17E+08 | 2.49E+08 | 2.96E+08 | 3.25E+08 | 2.30E+08 | 2.42E+08 | 2.45E+08 | 2.44E+08 | 2.51E+08 | 2.64E+08 | 0.95 | -0.07 | 0.534270937 |
| Q00839 | Heterogeneous nuclear ribonucleoprotein U OS=Homo sapiens OX=9606 GN=HNRNPU PE=1 SV=6 | 29 | 335 | 19 | 1.53E+08 | 1.35E+08 | 1.24E+08 | 1.56E+08 | 1.37E+08 | 1.39E+08 | 7.50E+07 | 7.96E+07 | 6.56E+07 | 6.37E+07 | 6.21E+07 | 5.58E+07 | 1.41E+08 | 6.70E+07 | 2.10 | 1.07 | 5.55736E-07 |
| P56192 | Methionine--tRNA ligase, cytoplasmic OS=Homo sapiens OX=9606 GN=MARS PE=1 SV=2 | 33 | 264 | 19 | 8.06E+07 | 8.28E+07 | 7.22E+07 | 7.27E+07 | 7.33E+07 | 7.32E+07 | 4.71E+07 | 4.93E+07 | 3.60E+07 | 3.88E+07 | 3.80E+07 | 3.58E+07 | 7.58E+07 | 4.08E+07 | 1.86 | 0.89 | 7.18303E-07 |
| O60701 | UDP-glucose 6-dehydrogenase OS=Homo sapiens OX=9606 GN=UGDH PE=1 SV=1 | 52 | 330 | 19 | 8.34E+07 | 8.77E+07 | 8.54E+07 | 8.43E+07 | 8.85E+07 | 9.67E+07 | 1.06E+08 | 9.84E+07 | 8.79E+07 | 8.68E+07 | 1.07E+08 | 8.84E+07 | 8.77E+07 | 9.58E+07 | 0.92 | -0.13 | 0.097750282 |
| Q06210 | Glutamine--fructose-6-phosphate aminotransferase [isomerizing] 1 OS=Homo sapiens OX=9606 GN=GFPT1 PE=1 SV=3 | 36 | 282 | 19 | 7.70E+07 | 8.02E+07 | 6.49E+07 | 6.94E+07 | 6.86E+07 | 6.49E+07 | 7.82E+07 | 8.24E+07 | 7.38E+07 | 7.36E+07 | 7.13E+07 | 6.18E+07 | 7.08E+07 | 7.35E+07 | 0.96 | -0.05 | 0.503681628 |
| O00468 | Agrin OS=Homo sapiens OX=9606 GN=AGRN PE=1 SV=6 | 15 | 192 | 19 | 2.31E+07 | 2.74E+07 | 2.19E+07 | 2.45E+07 | 2.58E+07 | 2.27E+07 | 4.79E+07 | 4.69E+07 | 4.27E+07 | 3.81E+07 | 3.62E+07 | 3.69E+07 | 2.43E+07 | 4.15E+07 | 0.59 | -0.77 | 0.00016833 |
| P11387 | DNA topoisomerase 1 OS=Homo sapiens OX=9606 GN=TOP1 PE=1 SV=2 | 26 | 438 | 19 | 1.46E+08 | 1.52E+08 | 1.60E+08 | 1.47E+08 | 1.56E+08 | 1.38E+08 | 1.48E+08 | 1.57E+08 | 1.60E+08 | 1.41E+08 | 1.38E+08 | 1.23E+08 | 1.50E+08 | 1.45E+08 | 1.04 | 0.05 | 0.441249888 |
| P26640 | Valine--tRNA ligase OS=Homo sapiens OX=9606 GN=VARS PE=1 SV=4 | 22 | 193 | 19 | 3.58E+07 | 3.63E+07 | 3.99E+07 | 3.63E+07 | 3.79E+07 | 3.53E+07 | 2.61E+07 | 2.77E+07 | 2.82E+07 | 2.49E+07 | 2.38E+07 | 2.34E+07 | 3.69E+07 | 2.57E+07 | 1.44 | 0.52 | 1.25035E-06 |
| P62805 | Histone H4 OS=Homo sapiens OX=9606 GN=H4C1 PE=1 SV=2 | 65 | 9740 | 20 | 1.16E+10 | 1.10E+10 | 1.35E+10 | 1.27E+10 | 1.27E+10 | 1.27E+10 | 1.17E+10 | 1.08E+10 | 1.32E+10 | 1.31E+10 | 1.27E+10 | 1.27E+10 | 1.24E+10 | 1.24E+10 | 1.00 | 0.00 | 0.963183175 |
| P11216 | Glycogen phosphorylase, brain form OS=Homo sapiens OX=9606 GN=PYGB PE=1 SV=5 | 42 | 611 | 20 | 8.74E+07 | 9.30E+07 | 1.00E+08 | 9.44E+07 | 9.24E+07 | 9.41E+07 | 8.76E+07 | 9.36E+07 | 8.52E+07 | 8.79E+07 | 8.36E+07 | 7.14E+07 | 9.36E+07 | 8.49E+07 | 1.10 | 0.14 | 0.037576908 |
| O75367 | Core histone macro-H2A.1 OS=Homo sapiens OX=9606 GN=H2AFY PE=1 SV=4 | 60 | 714 | 20 | 3.84E+08 | 4.43E+08 | 3.23E+08 | 3.69E+08 | 3.55E+08 | 4.38E+08 | 4.56E+08 | 5.55E+08 | 3.89E+08 | 3.67E+08 | 4.70E+08 | 3.90E+08 | 3.85E+08 | 4.38E+08 | 0.88 | -0.18 | 0.16388985 |
| P08670 | Vimentin OS=Homo sapiens OX=9606 GN=VIM PE=1 SV=4 | 45 | 354 | 20 | 1.29E+08 | 1.35E+08 | 1.33E+08 | 1.46E+08 | 1.59E+08 | 1.33E+08 | 6.43E+07 | 6.37E+07 | 5.97E+07 | 6.08E+07 | 6.44E+07 | 5.39E+07 | 1.39E+08 | 6.11E+07 | 2.28 | 1.19 | 2.47387E-06 |
| P55884 | Eukaryotic translation initiation factor 3 subunit B OS=Homo sapiens OX=9606 GN=EIF3B PE=1 SV=3 | 31 | 257 | 20 | 6.42E+07 | 5.92E+07 | 6.81E+07 | 5.85E+07 | 6.40E+07 | 5.65E+07 | 5.34E+07 | 4.28E+07 | 4.26E+07 | 3.50E+07 | 4.06E+07 | 3.51E+07 | 6.18E+07 | 4.16E+07 | 1.48 | 0.57 | 0.000213992 |
| Q14697 | Neutral alpha-glucosidase AB OS=Homo sapiens OX=9606 GN=GANAB PE=1 SV=3 | 29 | 265 | 20 | 7.86E+07 | 7.47E+07 | 7.10E+07 | 7.26E+07 | 7.27E+07 | 7.55E+07 | 6.37E+07 | 6.30E+07 | 4.71E+07 | 4.76E+07 | 4.87E+07 | 4.74E+07 | 7.42E+07 | 5.29E+07 | 1.40 | 0.49 | 0.000834923 |
| P50990 | T-complex protein 1 subunit theta OS=Homo sapiens OX=9606 GN=CCT8 PE=1 SV=4 | 44 | 531 | 20 | 1.82E+08 | 1.90E+08 | 1.76E+08 | 1.77E+08 | 1.81E+08 | 1.62E+08 | 1.21E+08 | 1.33E+08 | 1.31E+08 | 1.18E+08 | 1.09E+08 | 1.20E+08 | 1.78E+08 | 1.22E+08 | 1.46 | 0.55 | 8.70307E-07 |
| P23526 | Adenosylhomocysteinase OS=Homo sapiens OX=9606 GN=AHCY PE=1 SV=4 | 45 | 1122 | 21 | 1.03E+09 | 1.07E+09 | 1.24E+09 | 1.24E+09 | 1.22E+09 | 1.09E+09 | 1.50E+09 | 1.36E+09 | 1.58E+09 | 1.56E+09 | 1.49E+09 | 1.26E+09 | 1.15E+09 | 1.46E+09 | 0.79 | -0.35 | 0.000722513 |
| Q13200 | 26S proteasome non-ATPase regulatory subunit 2 OS=Homo sapiens OX=9606 GN=PSMD2 PE=1 SV=3 | 33 | 309 | 21 | 1.01E+08 | 9.62E+07 | 8.49E+07 | 9.35E+07 | 9.36E+07 | 8.54E+07 | 6.72E+07 | 6.70E+07 | 6.32E+07 | 5.90E+07 | 5.91E+07 | 4.67E+07 | 9.25E+07 | 6.04E+07 | 1.53 | 0.61 | 1.49926E-05 |
| P12956 | X-ray repair cross-complementing protein 6 OS=Homo sapiens OX=9606 GN=XRCC6 PE=1 SV=2 | 43 | 450 | 21 | 1.02E+08 | 1.06E+08 | 1.08E+08 | 9.56E+07 | 1.07E+08 | 9.65E+07 | 9.81E+07 | 9.57E+07 | 1.09E+08 | 8.10E+07 | 9.81E+07 | 7.80E+07 | 1.03E+08 | 9.34E+07 | 1.10 | 0.13 | 0.126294519 |
| P62701 | 40S ribosomal protein S4, X isoform OS=Homo sapiens OX=9606 GN=RPS4X PE=1 SV=2 | 59 | 446 | 21 | 1.96E+08 | 2.05E+08 | 1.96E+08 | 2.16E+08 | 1.95E+08 | 1.84E+08 | 1.58E+08 | 1.59E+08 | 1.59E+08 | 1.58E+08 | 1.51E+08 | 1.27E+08 | 1.99E+08 | 1.52E+08 | 1.31 | 0.39 | 5.18458E-05 |
| Q9Y262 | Eukaryotic translation initiation factor 3 subunit L OS=Homo sapiens OX=9606 GN=EIF3L PE=1 SV=1 | 43 | 323 | 21 | 8.32E+07 | 9.09E+07 | 9.52E+07 | 7.49E+07 | 8.57E+07 | 7.17E+07 | 5.98E+07 | 5.83E+07 | 5.63E+07 | 5.73E+07 | 5.41E+07 | 4.42E+07 | 8.36E+07 | 5.50E+07 | 1.52 | 0.60 | 0.000144126 |

### Supplementary Table 4. List of all proteins identified by label free proteomics analysis.

| Accession | Description | Coverage [%] | # PSMs | # Unique Peptides | Amitriptyline 1 | Amitriptyline 2 | Amitriptyline 3 | Amitriptyline 4 | Amitriptyline 5 | Amitriptyline 6 | Vehicle 1 | Vehicle 2 | Vehicle 3 | Vehicle 4 | Vehicle 5 | Vehicle 6 | Average amitriptyline | Average vehicle | Ratio amitriptyline/ vehicle | Log 2 fold change (amitriptyline /vehicle) | P-value |
|---|---|---|---|---|---|---|---|---|---|---|---|---|---|---|---|---|---|---|---|---|---|
| Q86VP6 | Cullin-associated NEDD8-dissociated protein 1 OS=Homo sapiens OX=9606 GN=CAND1 PE=1 SV=2 | 24 | 316 | 21 | 9.30E+07 | 9.16E+07 | 8.64E+07 | 8.76E+07 | 8.37E+07 | 8.43E+07 | 9.53E+07 | 8.02E+07 | 6.86E+07 | 7.16E+07 | 6.46E+07 | 7.08E+07 | 8.78E+07 | 7.52E+07 | 1.17 | 0.22 | 0.038356109 |
| P36871 | Phosphoglucomutase-1 OS=Homo sapiens OX=9606 GN=PGM1 PE=1 SV=3 | 50 | 249 | 21 | 1.18E+08 | 1.10E+08 | 1.18E+08 | 1.26E+08 | 1.25E+08 | 1.14E+08 | 1.57E+08 | 1.57E+08 | 1.62E+08 | 1.49E+08 | 1.37E+08 | 1.05E+08 | 1.18E+08 | 1.45E+08 | 0.82 | -0.29 | 0.027021135 |
| P08473 | Neprilysin OS=Homo sapiens OX=9606 GN=MME PE=1 SV=2 | 40 | 278 | 21 | 7.90E+07 | 7.76E+07 | 9.87E+07 | 8.40E+07 | 9.29E+07 | 8.00E+07 | 6.14E+07 | 5.84E+07 | 7.07E+07 | 8.06E+07 | 5.84E+07 | 4.12E+07 | 8.54E+07 | 6.18E+07 | 1.38 | 0.47 | 0.005749762 |
| Q9P2J5 | Leucine--tRNA ligase, cytoplasmic OS=Homo sapiens OX=9606 GN=LARS PE=1 SV=2 | 23 | 258 | 21 | 7.95E+07 | 7.32E+07 | 7.08E+07 | 6.19E+07 | 7.17E+07 | 6.54E+07 | 5.73E+07 | 3.84E+07 | 4.38E+07 | 4.23E+07 | 4.18E+07 | 3.68E+07 | 7.04E+07 | 4.34E+07 | 1.62 | 0.70 | 4.74058E-05 |
| Q5T4S7 | E3 ubiquitin-protein ligase UBR4 OS=Homo sapiens OX=9606 GN=UBR4 PE=1 SV=1 | 7 | 102 | 21 | 2.18E+07 | 1.92E+07 | 2.20E+07 | 2.17E+07 | 2.23E+07 | 2.09E+07 | 1.63E+07 | 1.05E+07 | 1.34E+07 | 1.19E+07 | 1.42E+07 | 1.15E+07 | 2.13E+07 | 1.30E+07 | 1.65 | 0.72 | 3.28881E-05 |
| P23396 | 40S ribosomal protein S3 OS=Homo sapiens OX=9606 GN=RPS3 PE=1 SV=2 | 78 | 566 | 21 | 1.71E+08 | 1.81E+08 | 1.66E+08 | 1.47E+08 | 1.63E+08 | 1.46E+08 | 1.25E+08 | 1.34E+08 | 1.24E+08 | 1.16E+08 | 1.21E+08 | 1.01E+08 | 1.62E+08 | 1.20E+08 | 1.35 | 0.43 | 0.000208221 |
| Q15029 | 116 kDa U5 small nuclear ribonucleoprotein component OS=Homo sapiens OX=9606 GN=EFTUD2 PE=1 SV=1 | 33 | 201 | 21 | 5.50E+07 | 5.46E+07 | 5.28E+07 | 5.45E+07 | 5.85E+07 | 5.42E+07 | 3.82E+07 | 3.65E+07 | 3.86E+07 | 3.87E+07 | 4.25E+07 | 3.41E+07 | 5.49E+07 | 3.81E+07 | 1.44 | 0.53 | 7.64206E-07 |
| P08238 | Heat shock protein HSP 90-beta OS=Homo sapiens OX=9606 GN=HSP90AB1 PE=1 SV=4 | 52 | 1866 | 22 | 8.71E+08 | 9.37E+08 | 8.56E+08 | 8.81E+08 | 8.96E+08 | 8.14E+08 | 8.91E+08 | 9.15E+08 | 8.41E+08 | 8.06E+08 | 8.12E+08 | 7.02E+08 | 8.76E+08 | 8.28E+08 | 1.06 | 0.08 | 0.206432937 |
| P06737 | Glycogen phosphorylase, liver form OS=Homo sapiens OX=9606 GN=PYGL PE=1 SV=4 | 40 | 787 | 22 | 2.48E+08 | 2.43E+08 | 2.23E+08 | 2.23E+08 | 2.25E+08 | 2.26E+08 | 2.52E+08 | 2.53E+08 | 2.43E+08 | 2.24E+08 | 2.33E+08 | 1.95E+08 | 2.31E+08 | 2.33E+08 | 0.99 | -0.01 | 0.834485045 |
| P21399 | Cytoplasmic aconitate hydratase OS=Homo sapiens OX=9606 GN=ACO1 PE=1 SV=3 | 34 | 668 | 22 | 2.47E+08 | 2.44E+08 | 2.31E+08 | 7.36E+08 | 2.15E+08 | 2.33E+08 | 8.52E+08 | 3.18E+08 | 2.68E+08 | 2.66E+08 | 2.61E+08 | 2.51E+08 | 3.18E+08 | 3.69E+08 | 0.86 | -0.22 | 0.694913113 |
| P05023 | Sodium/potassium-transporting ATPase subunit alpha-1 OS=Homo sapiens OX=9606 GN=ATP1A1 PE=1 SV=1 | 26 | 335 | 22 | 1.31E+08 | 1.28E+08 | 1.21E+08 | 1.26E+08 | 1.24E+08 | 1.16E+08 | 1.02E+08 | 1.04E+08 | 9.44E+07 | 8.91E+07 | 8.88E+07 | 7.44E+07 | 1.24E+08 | 9.21E+07 | 1.35 | 0.43 | 0.000237934 |
| P35606 | Coatomer subunit beta' OS=Homo sapiens OX=9606 GN=COPB2 PE=1 SV=2 | 38 | 231 | 22 | 6.22E+07 | 7.29E+07 | 5.98E+07 | 6.90E+07 | 6.16E+07 | 6.42E+07 | 3.66E+07 | 3.92E+07 | 3.39E+07 | 3.69E+07 | 4.04E+07 | 3.17E+07 | 6.50E+07 | 3.64E+07 | 1.78 | 0.83 | 1.50556E-06 |
| P04406 | Glyceraldehyde-3-phosphate dehydrogenase OS=Homo sapiens OX=9606 GN=GAPDH PE=1 SV=3 | 82 | 4882 | 23 | 1.82E+09 | 1.71E+09 | 1.73E+09 | 1.64E+09 | 1.81E+09 | 1.91E+09 | 1.69E+09 | 1.72E+09 | 1.61E+09 | 1.60E+09 | 1.59E+09 | 1.55E+09 | 1.77E+09 | 1.63E+09 | 1.09 | 0.12 | 0.013562483 |
| P07900 | Heat shock protein HSP 90-alpha OS=Homo sapiens OX=9606 GN=HSP90AA1 PE=1 SV=5 | 50 | 1956 | 23 | 3.02E+08 | 2.93E+08 | 3.25E+08 | 3.36E+08 | 3.11E+08 | 2.85E+08 | 3.04E+08 | 3.40E+08 | 3.59E+08 | 3.17E+08 | 3.37E+08 | 2.82E+08 | 3.09E+08 | 3.23E+08 | 0.96 | -0.07 | 0.322908266 |
| P08195 | 4F2 cell-surface antigen heavy chain OS=Homo sapiens OX=9606 GN=SLC3A2 PE=1 SV=3 | 40 | 505 | 23 | 2.10E+08 | 2.14E+08 | 1.84E+08 | 1.79E+08 | 1.76E+08 | 1.88E+08 | 8.27E+07 | 8.29E+07 | 5.68E+07 | 6.45E+07 | 7.38E+07 | 6.18E+07 | 1.92E+08 | 7.04E+07 | 2.73 | 1.45 | 1.18049E-07 |
| Q9Y5B9 | FACT complex subunit SPT16 OS=Homo sapiens OX=9606 GN=SUPT16H PE=1 SV=1 | 29 | 295 | 23 | 7.51E+07 | 8.30E+07 | 8.24E+07 | 8.02E+07 | 6.67E+07 | 7.82E+07 | 3.79E+07 | 4.55E+07 | 3.80E+07 | 4.33E+07 | 3.80E+07 | 3.57E+07 | 7.76E+07 | 3.97E+07 | 1.95 | 0.97 | 7.78542E-07 |
| Q15149 | Plectin OS=Homo sapiens OX=9606 GN=PLEC PE=1 SV=3 | 7 | 143 | 23 | 3.00E+07 | 3.30E+07 | 3.08E+07 | 3.88E+07 | 3.33E+07 | 3.19E+07 | 2.19E+07 | 1.83E+07 | 2.27E+07 | 2.31E+07 | 2.02E+07 | 1.84E+07 | 3.30E+07 | 2.08E+07 | 1.59 | 0.67 | 2.85463E-05 |
| P06733 | Alpha-enolase OS=Homo sapiens OX=9606 GN=ENO1 PE=1 SV=2 | 72 | 1037 | 24 | 5.26E+08 | 6.02E+08 | 4.50E+08 | 4.65E+08 | 4.48E+08 | 5.30E+08 | 5.68E+08 | 6.82E+08 | 4.69E+08 | 5.25E+08 | 5.39E+08 | 5.18E+08 | 5.04E+08 | 5.50E+08 | 0.92 | -0.13 | 0.253643333 |
| P48643 | T-complex protein 1 subunit epsilon OS=Homo sapiens OX=9606 GN=CCT5 PE=1 SV=1 | 56 | 678 | 24 | 2.22E+08 | 2.23E+08 | 1.91E+08 | 2.00E+08 | 1.96E+08 | 2.06E+08 | 1.47E+08 | 1.56E+08 | 1.40E+08 | 1.26E+08 | 1.33E+08 | 1.09E+08 | 2.06E+08 | 1.35E+08 | 1.53 | 0.61 | 1.24491E-05 |
| P50991 | T-complex protein 1 subunit delta OS=Homo sapiens OX=9606 GN=CCT4 PE=1 SV=4 | 51 | 660 | 24 | 2.95E+08 | 2.90E+08 | 2.69E+08 | 2.72E+08 | 2.43E+08 | 2.83E+08 | 1.98E+08 | 1.90E+08 | 1.63E+08 | 1.81E+08 | 1.71E+08 | 1.57E+08 | 2.75E+08 | 1.77E+08 | 1.56 | 0.64 | 2.15189E-06 |
| P19338 | Nucleolin OS=Homo sapiens OX=9606 GN=NCL PE=1 SV=3 | 33 | 322 | 24 | 9.01E+07 | 8.93E+07 | 9.76E+07 | 1.31E+08 | 9.84E+07 | 1.05E+08 | 6.56E+07 | 6.26E+07 | 6.58E+07 | 5.77E+07 | 6.44E+07 | 5.27E+07 | 1.02E+08 | 6.15E+07 | 1.66 | 0.73 | 0.000859936 |
| P08758 | Annexin A5 OS=Homo sapiens OX=9606 GN=ANXA5 PE=1 SV=2 | 80 | 991 | 25 | 5.08E+08 | 4.55E+08 | 4.84E+08 | 4.77E+08 | 5.04E+08 | 4.41E+08 | 3.91E+08 | 3.55E+08 | 3.40E+08 | 3.38E+08 | 3.74E+08 | 2.94E+08 | 4.78E+08 | 3.49E+08 | 1.37 | 0.46 | 3.00974E-05 |
| P78371 | T-complex protein 1 subunit beta OS=Homo sapiens OX=9606 GN=CCT2 PE=1 SV=4 | 61 | 555 | 25 | 2.83E+08 | 2.85E+08 | 2.43E+08 | 2.51E+08 | 2.35E+08 | 2.30E+08 | 1.92E+08 | 2.02E+08 | 1.60E+08 | 1.69E+08 | 1.84E+08 | 1.51E+08 | 2.55E+08 | 1.76E+08 | 1.44 | 0.53 | 0.000124783 |
| P11142 | Heat shock cognate 71 kDa protein OS=Homo sapiens OX=9606 GN=HSPA8 PE=1 SV=1 | 66 | 2437 | 26 | 9.83E+08 | 9.82E+08 | 9.35E+08 | 1.01E+09 | 9.65E+08 | 9.54E+08 | 9.54E+08 | 1.01E+09 | 9.02E+08 | 9.39E+08 | 8.91E+08 | 8.46E+08 | 9.72E+08 | 9.23E+08 | 1.05 | 0.07 | 0.093247678 |
| P40227 | T-complex protein 1 subunit zeta OS=Homo sapiens OX=9606 GN=CCT6A PE=1 SV=3 | 53 | 709 | 26 | 2.97E+08 | 2.95E+08 | 2.92E+08 | 3.03E+08 | 3.06E+08 | 2.83E+08 | 2.21E+08 | 2.17E+08 | 2.05E+08 | 1.92E+08 | 2.01E+08 | 1.77E+08 | 2.96E+08 | 2.02E+08 | 1.46 | 0.55 | 3.16712E-06 |
| P13010 | X-ray repair cross-complementing protein 5 OS=Homo sapiens OX=9606 GN=XRCC5 PE=1 SV=3 | 53 | 374 | 26 | 1.10E+08 | 1.14E+08 | 1.09E+08 | 1.01E+08 | 1.12E+08 | 1.13E+08 | 7.94E+07 | 8.32E+07 | 7.40E+07 | 7.05E+07 | 6.78E+07 | 6.29E+07 | 1.10E+08 | 7.30E+07 | 1.51 | 0.59 | 4.82714E-06 |
| P17987 | T-complex protein 1 subunit alpha OS=Homo sapiens OX=9606 GN=TCP1 PE=1 SV=1 | 66 | 692 | 26 | 2.89E+08 | 3.16E+08 | 3.03E+08 | 2.56E+08 | 2.93E+08 | 2.80E+08 | 2.04E+08 | 2.37E+08 | 1.97E+08 | 2.10E+08 | 2.14E+08 | 1.72E+08 | 2.90E+08 | 2.05E+08 | 1.41 | 0.50 | 3.73317E-05 |
| Q8WUM4 | Programmed cell death 6-interacting protein OS=Homo sapiens OX=9606 GN=PDCD6IP PE=1 SV=1 | 41 | 379 | 26 | 8.91E+07 | 9.65E+07 | 9.81E+07 | 9.75E+07 | 9.05E+07 | 9.14E+07 | 8.57E+07 | 7.93E+07 | 8.23E+07 | 8.04E+07 | 8.38E+07 | 7.47E+07 | 9.38E+07 | 8.10E+07 | 1.16 | 0.21 | 0.000202288 |
| Q14152 | Eukaryotic translation initiation factor 3 subunit A OS=Homo sapiens OX=9606 GN=EIF3A PE=1 SV=1 | 21 | 374 | 26 | 1.17E+08 | 1.15E+08 | 1.11E+08 | 1.12E+08 | 8.94E+07 | 1.05E+08 | 6.75E+07 | 6.08E+07 | 6.15E+07 | 6.26E+07 | 6.03E+07 | 5.66E+07 | 1.08E+08 | 6.15E+07 | 1.76 | 0.82 | 3.26269E-05 |
| P02679 | Fibrinogen gamma chain OS=Homo sapiens OX=9606 GN=FGG PE=1 SV=3 | 67 | 603 | 27 | 6.97E+07 | 6.14E+07 | 6.84E+07 | 6.97E+07 | 6.63E+07 | 7.04E+07 | 3.38E+08 | 1.48E+08 | 9.16E+07 | 8.50E+07 | 7.93E+07 | 3.84E+09 | 6.77E+07 | 7.64E+08 | 0.09 | -3.50 | 0.310239774 |
| P0DMV9 | Heat shock 70 kDa protein 1B OS=Homo sapiens OX=9606 GN=HSPA1B PE=1 SV=1 | 63 | 974 | 27 | 2.82E+08 | 2.89E+08 | 2.39E+08 | 2.48E+08 | 2.24E+08 | 2.51E+08 | 2.79E+08 | 2.98E+08 | 2.46E+08 | 2.28E+08 | 2.38E+08 | 2.05E+08 | 2.55E+08 | 2.49E+08 | 1.03 | 0.04 | 0.712895103 |
| P49368 | T-complex protein 1 subunit gamma OS=Homo sapiens OX=9606 GN=CCT3 PE=1 SV=4 | 54 | 599 | 27 | 2.47E+08 | 2.40E+08 | 1.93E+08 | 1.83E+08 | 1.84E+08 | 2.15E+08 | 1.59E+08 | 1.58E+08 | 1.24E+08 | 1.40E+08 | 1.33E+08 | 1.34E+08 | 2.10E+08 | 1.41E+08 | 1.49 | 0.57 | 0.0008459 |
| P31939 | Bifunctional purine biosynthesis protein PURH OS=Homo sapiens OX=9606 GN=ATIC PE=1 SV=3 | 57 | 530 | 27 | 1.55E+08 | 1.54E+08 | 1.45E+08 | 1.47E+08 | 1.54E+08 | 1.35E+08 | 1.82E+08 | 1.85E+08 | 1.57E+08 | 1.53E+08 | 1.56E+08 | 1.37E+08 | 1.48E+08 | 1.62E+08 | 0.92 | -0.12 | 0.151560445 |
| Q99832 | T-complex protein 1 subunit eta OS=Homo sapiens OX=9606 GN=CCT7 PE=1 SV=2 | 61 | 772 | 28 | 3.51E+08 | 3.48E+08 | 3.08E+08 | 3.07E+08 | 3.34E+08 | 3.28E+08 | 2.33E+08 | 2.52E+08 | 2.20E+08 | 2.33E+08 | 2.34E+08 | 2.02E+08 | 3.29E+08 | 2.29E+08 | 1.44 | 0.52 | 2.22104E-06 |
| P05556 | Integrin beta-1 OS=Homo sapiens OX=9606 GN=ITGB1 PE=1 SV=2 | 38 | 775 | 28 | 3.82E+08 | 3.88E+08 | 3.73E+08 | 3.77E+08 | 3.64E+08 | 3.64E+08 | 2.69E+08 | 2.76E+08 | 2.38E+08 | 2.28E+08 | 2.21E+08 | 2.18E+08 | 3.75E+08 | 2.42E+08 | 1.55 | 0.63 | 1.13194E-05 |
| P11586 | C-1-tetrahydrofolate synthase, cytoplasmic OS=Homo sapiens OX=9606 GN=MTHFD1 PE=1 SV=3 | 36 | 445 | 28 | 1.37E+08 | 1.40E+08 | 1.31E+08 | 1.33E+08 | 1.21E+08 | 1.23E+08 | 1.23E+08 | 1.30E+08 | 1.31E+08 | 1.25E+08 | 1.09E+08 | 1.22E+08 | 1.31E+08 | 1.23E+08 | 1.06 | 0.08 | 0.123660787 |
| P09874 | Poly [ADP-ribose] polymerase 1 OS=Homo sapiens OX=9606 GN=PARP1 PE=1 SV=4 | 40 | 499 | 30 | 2.05E+08 | 1.95E+08 | 2.07E+08 | 2.09E+08 | 2.01E+08 | 1.95E+08 | 1.08E+08 | 1.04E+08 | 9.42E+07 | 9.77E+07 | 8.86E+07 | 8.98E+07 | 2.02E+08 | 9.70E+07 | 2.08 | 1.06 | 5.55454E-10 |
| P53621 | Coatomer subunit alpha OS=Homo sapiens OX=9606 GN=COPA PE=1 SV=2 | 34 | 420 | 30 | 2.86E+08 | 2.80E+08 | 3.24E+08 | 4.11E+08 | 4.45E+08 | 3.24E+08 | 3.67E+08 | 3.78E+08 | 4.07E+08 | 5.65E+08 | 5.21E+08 | 3.01E+08 | 3.45E+08 | 4.23E+08 | 0.82 | -0.29 | 0.148602083 |
| P41252 | Isoleucine--tRNA ligase, cytoplasmic OS=Homo sapiens OX=9606 GN=IARS PE=1 SV=2 | 32 | 497 | 30 | 1.33E+08 | 1.45E+08 | 1.15E+08 | 1.29E+08 | 1.26E+08 | 1.26E+08 | 8.39E+07 | 7.45E+07 | 7.54E+07 | 7.08E+07 | 7.22E+07 | 7.20E+07 | 1.29E+08 | 7.48E+07 | 1.72 | 0.79 | 4.89517E-06 |
| P98160 | Basement membrane-specific heparan sulfate proteoglycan core protein OS=Homo sapiens OX=9606 GN=HSPG2 PE=1 SV=4 | 10 | 269 | 30 | 9.29E+07 | 6.69E+07 | 5.83E+07 | 6.04E+07 | 5.34E+07 | 6.05E+07 | 9.00E+07 | 8.58E+07 | 5.08E+07 | 5.65E+07 | 5.69E+07 | 6.93E+07 | 6.54E+07 | 6.82E+07 | 0.96 | -0.06 | 0.758444528 |
| Q6P2Q9 | Pre-mRNA-processing-splicing factor 8 OS=Homo sapiens OX=9606 GN=PRPF8 PE=1 SV=2 | 19 | 225 | 30 | 6.08E+07 | 5.59E+07 | 5.37E+07 | 4.66E+07 | 5.16E+07 | 5.28E+07 | 3.53E+07 | 2.13E+07 | 2.35E+07 | 3.38E+07 | 2.73E+07 | 2.22E+07 | 5.36E+07 | 2.72E+07 | 1.97 | 0.98 | 1.10292E-05 |
| P35908 | Keratin, type II cytoskeletal 2 epidermal OS=Homo sapiens OX=9606 GN=KRT2 PE=1 SV=2 | 70 | 1456 | 31 | 4.85E+08 | 1.21E+09 | 2.35E+08 | 3.33E+08 | 4.06E+08 | 4.98E+08 | 1.23E+08 | 6.51E+08 | 2.20E+08 | 2.93E+08 | 3.74E+08 | 2.68E+08 | 5.28E+08 | 3.22E+08 | 1.64 | 0.72 | 0.236554245 |
| Q08211 | ATP-dependent RNA helicase A OS=Homo sapiens OX=9606 GN=DHX9 PE=1 SV=4 | 34 | 577 | 31 | 2.17E+08 | 2.10E+08 | 2.16E+08 | 2.24E+08 | 2.19E+08 | 2.16E+08 | 1.81E+08 | 1.85E+08 | 1.48E+08 | 1.65E+08 | 1.63E+08 | 1.49E+08 | 2.17E+08 | 1.65E+08 | 1.31 | 0.39 | 0.000259641 |
| O00159 | Unconventional myosin-Ic OS=Homo sapiens OX=9606 GN=MYO1C PE=1 SV=4 | 41 | 488 | 32 | 1.18E+08 | 1.07E+08 | 8.96E+07 | 8.97E+07 | 9.79E+07 | 9.14E+07 | 8.50E+07 | 8.28E+07 | 8.19E+07 | 7.51E+07 | 7.66E+07 | 6.03E+07 | 9.90E+07 | 7.70E+07 | 1.29 | 0.36 | 0.004669494 |
| P13645 | Keratin, type I cytoskeletal 10 OS=Homo sapiens OX=9606 GN=KRT10 PE=1 SV=6 | 58 | 1611 | 33 | 1.34E+09 | 2.38E+09 | 5.96E+08 | 8.69E+08 | 1.02E+09 | 1.21E+09 | 2.75E+08 | 1.47E+09 | 5.68E+08 | 7.63E+08 | 9.44E+08 | 6.51E+08 | 1.24E+09 | 7.78E+08 | 1.59 | 0.67 | 0.165045892 |
| P02671 | Fibrinogen alpha chain OS=Homo sapiens OX=9606 GN=FGA PE=1 SV=2 | 39 | 719 | 33 | 1.54E+07 | 1.65E+07 | 1.29E+07 | 1.45E+07 | 9.89E+06 | 1.58E+07 | 2.67E+08 | 1.03E+08 | 6.26E+07 | 6.13E+07 | 3.80E+07 | 3.26E+09 | 1.42E+07 | 6.32E+08 | 0.02 | -5.48 | 0.293603515 |
| P53396 | ATP-citrate synthase OS=Homo sapiens OX=9606 GN=ACLY PE=1 SV=3 | 36 | 770 | 33 | 2.01E+08 | 2.03E+08 | 1.84E+08 | 1.69E+08 | 1.68E+08 | 1.95E+08 | 1.73E+08 | 1.85E+08 | 1.55E+08 | 1.62E+08 | 1.68E+08 | 1.55E+08 | 1.87E+08 | 1.66E+08 | 1.12 | 0.17 | 0.029325133 |
| P35527 | Keratin, type I cytoskeletal 9 OS=Homo sapiens OX=9606 GN=KRT9 PE=1 SV=3 | 69 | 1049 | 34 | 7.49E+08 | 7.17E+08 | 3.45E+08 | 7.96E+08 | 6.78E+08 | 1.27E+09 | 3.70E+08 | 9.10E+08 | 3.49E+08 | 6.27E+08 | 8.38E+08 | 6.80E+08 | 7.59E+08 | 6.29E+08 | 1.21 | 0.27 | 0.419000351 |
| Q14764 | Major vault protein OS=Homo sapiens OX=9606 GN=MVP PE=1 SV=4 | 57 | 669 | 35 | 2.44E+08 | 2.41E+08 | 2.58E+08 | 2.62E+08 | 2.59E+08 | 2.45E+08 | 2.16E+08 | 2.31E+08 | 2.27E+08 | 2.00E+08 | 2.21E+08 | 1.74E+08 | 2.51E+08 | 2.12E+08 | 1.19 | 0.25 | 0.004458123 |
| Q6UVK1 | Chondroitin sulfate proteoglycan 4 OS=Homo sapiens OX=9606 GN=CSPG4 PE=1 SV=2 | 24 | 263 | 36 | 8.48E+07 | 8.39E+07 | 7.44E+07 | 7.44E+07 | 7.14E+07 | 7.15E+07 | 3.34E+07 | 3.14E+07 | 2.91E+07 | 2.56E+07 | 2.89E+07 | 2.61E+07 | 7.67E+07 | 2.91E+07 | 2.64 | 1.40 | 3.46182E-07 |
| P04264 | Keratin, type II cytoskeletal 1 OS=Homo sapiens OX=9606 GN=KRT1 PE=1 SV=6 | 62 | 2340 | 37 | 2.47E+09 | 2.77E+09 | 1.26E+09 | 2.08E+09 | 2.08E+09 | 3.06E+09 | 1.37E+09 | 2.81E+09 | 1.64E+09 | 2.06E+09 | 2.39E+09 | 1.85E+09 | 2.29E+09 | 2.02E+09 | 1.13 | 0.18 | 0.446670947 |
| P55072 | Transitional endoplasmic reticulum ATPase OS=Homo sapiens OX=9606 GN=VCP PE=1 SV=4 | 54 | 900 | 37 | 3.83E+08 | 3.70E+08 | 3.19E+08 | 3.25E+08 | 3.09E+08 | 3.69E+08 | 4.46E+08 | 3.92E+08 | 2.78E+08 | 2.68E+08 | 3.57E+08 | 3.22E+08 | 3.46E+08 | 3.44E+08 | 1.01 | 0.01 | 0.955918535 |
| P46940 | Ras GTPase-activating-like protein IQGAP1 OS=Homo sapiens OX=9606 GN=IQGAP1 PE=1 SV=1 | 35 | 378 | 37 | 8.21E+07 | 9.03E+07 | 8.61E+07 | 8.35E+07 | 7.60E+07 | 8.05E+07 | 7.51E+07 | 6.36E+07 | 6.87E+07 | 5.56E+07 | 6.55E+07 | 6.20E+07 | 8.31E+07 | 6.51E+07 | 1.28 | 0.35 | 0.000410721 |
| P07355 | Annexin A2 OS=Homo sapiens OX=9606 GN=ANXA2 PE=1 SV=2 | 78 | 1569 | 38 | 1.08E+09 | 1.05E+09 | 1.09E+09 | 9.28E+08 | 9.95E+08 | 1.02E+09 | 8.68E+08 | 8.87E+08 | 1.01E+09 | 8.21E+08 | 8.89E+08 | 7.72E+08 | 1.03E+09 | 8.74E+08 | 1.17 | 0.23 | 0.004093417 |
| P22314 | Ubiquitin-like modifier-activating enzyme 1 OS=Homo sapiens OX=9606 GN=UBA1 PE=1 SV=3 | 52 | 1085 | 40 | 3.25E+08 | 3.25E+08 | 2.84E+08 | 2.99E+08 | 2.72E+08 | 3.07E+08 | 3.14E+08 | 3.12E+08 | 2.71E+08 | 2.54E+08 | 2.79E+08 | 2.60E+08 | 3.02E+08 | 2.82E+08 | 1.07 | 0.10 | 0.168276897 |
| P07996 | Thrombospondin-1 OS=Homo sapiens OX=9606 GN=THBS1 PE=1 SV=2 | 45 | 1374 | 41 | 4.14E+08 | 4.27E+08 | 4.35E+08 | 4.16E+08 | 3.86E+08 | 4.18E+08 | 6.52E+08 | 6.48E+08 | 6.56E+08 | 6.24E+08 | 6.06E+08 | 5.96E+08 | 4.16E+08 | 6.31E+08 | 0.66 | -0.60 | 5.27845E-08 |
| P07814 | Bifunctional glutamate/proline--tRNA ligase OS=Homo sapiens OX=9606 GN=EPRS PE=1 SV=5 | 40 | 562 | 41 | 1.54E+08 | 1.45E+08 | 1.40E+08 | 1.58E+08 | 1.41E+08 | 1.47E+08 | 9.20E+07 | 8.36E+07 | 8.59E+07 | 8.29E+07 | 7.67E+07 | 7.46E+07 | 1.47E+08 | 8.26E+07 | 1.78 | 0.84 | 1.67041E-08 |
| P01024 | Complement C3 OS=Homo sapiens OX=9606 GN=C3 PE=1 SV=2 | 31 | 5444 | 42 | 3.85E+09 | 4.06E+09 | 4.88E+09 | 4.75E+09 | 4.91E+09 | 4.65E+09 | 5.43E+09 | 5.29E+09 | 6.73E+09 | 5.99E+09 | 7.00E+09 | 5.55E+09 | 4.52E+09 | 6.00E+09 | 0.75 | -0.41 | 0.002371545 |
| O75643 | U5 small nuclear ribonucleoprotein 200 kDa helicase OS=Homo sapiens OX=9606 GN=SNRNP200 PE=1 SV=2 | 29 | 405 | 42 | 9.26E+07 | 9.55E+07 | 8.26E+07 | 8.89E+07 | 8.61E+07 | 9.39E+07 | 6.74E+07 | 6.62E+07 | 5.79E+07 | 5.88E+07 | 5.47E+07 | 5.32E+07 | 8.99E+07 | 5.97E+07 | 1.51 | 0.59 | 2.77894E-06 |
| P14618 | Pyruvate kinase PKM OS=Homo sapiens OX=9606 GN=PKM PE=1 SV=4 | 76 | 2764 | 43 | 1.42E+09 | 1.58E+09 | 1.26E+09 | 1.33E+09 | 1.33E+09 | 1.49E+09 | 1.38E+09 | 1.56E+09 | 1.23E+09 | 1.27E+09 | 1.35E+09 | 1.24E+09 | 1.40E+09 | 1.34E+09 | 1.05 | 0.07 | 0.39160608 |
| P02675 | Fibrinogen beta chain OS=Homo sapiens OX=9606 GN=FGB PE=1 SV=2 | 76 | 920 | 44 | 2.55E+08 | 2.49E+08 | 2.60E+08 | 2.65E+08 | 2.44E+08 | 2.23E+08 | 6.61E+08 | 4.18E+08 | 3.60E+08 | 4.02E+08 | 2.81E+08 | 5.15E+09 | 2.49E+08 | 1.21E+09 | 0.21 | -2.28 | 0.276911003 |
| P13639 | Elongation factor 2 OS=Homo sapiens OX=9606 GN=EEF2 PE=1 SV=4 | 58 | 1610 | 45 | 7.71E+08 | 6.92E+08 | 7.21E+08 | 6.51E+08 | 6.65E+08 | 7.04E+08 | 7.10E+08 | 6.66E+08 | 6.21E+08 | 6.33E+08 | 6.38E+08 | 5.61E+08 | 7.01E+08 | 6.38E+08 | 1.10 | 0.13 | 0.042285128 |
| Q07954 | Prolow-density lipoprotein receptor-related protein 1 OS=Homo sapiens OX=9606 GN=LRP1 PE=1 SV=2 | 14 | 685 | 45 | 1.43E+08 | 1.50E+08 | 1.33E+08 | 1.44E+08 | 1.34E+08 | 1.43E+08 | 1.95E+08 | 1.97E+08 | 1.70E+08 | 1.80E+08 | 1.77E+08 | 1.55E+08 | 1.41E+08 | 1.79E+08 | 0.79 | -0.34 | 0.001256892 |
| P08133 | Annexin A6 OS=Homo sapiens OX=9606 GN=ANXA6 PE=1 SV=3 | 73 | 1213 | 48 | 4.77E+08 | 5.03E+08 | 4.71E+08 | 4.95E+08 | 4.79E+08 | 4.74E+08 | 3.95E+08 | 3.94E+08 | 3.68E+08 | 3.41E+08 | 3.85E+08 | 3.23E+08 | 4.83E+08 | 3.68E+08 | 1.31 | 0.39 | 6.75775E-05 |
| P78527 | DNA-dependent protein kinase catalytic subunit OS=Homo sapiens OX=9606 GN=PRKDC PE=1 SV=3 | 17 | 505 | 49 | 1.14E+08 | 1.22E+08 | 9.54E+07 | 1.17E+08 | 1.14E+08 | 1.03E+08 | 8.25E+07 | 7.09E+07 | 6.37E+07 | 7.06E+07 | 6.91E+07 | 5.91E+07 | 1.11E+08 | 6.93E+07 | 1.60 | 0.68 | 1.50396E-05 |
| O75369 | Filamin-B OS=Homo sapiens OX=9606 GN=FLNB PE=1 SV=2 | 37 | 870 | 54 | 1.92E+08 | 1.90E+08 | 1.68E+08 | 1.85E+08 | 1.59E+08 | 1.77E+08 | 1.44E+08 | 1.51E+08 | 1.33E+08 | 1.23E+08 | 1.23E+08 | 1.09E+08 | 1.78E+08 | 1.31E+08 | 1.37 | 0.45 | 0.000201024 |
| Q14315 | Filamin-C OS=Homo sapiens OX=9606 GN=FLNC PE=1 SV=3 | 33 | 902 | 55 | 2.18E+08 | 2.18E+08 | 2.05E+08 | 2.11E+08 | 1.97E+08 | 2.09E+08 | 1.50E+08 | 1.43E+08 | 1.36E+08 | 1.36E+08 | 1.24E+08 | 1.16E+08 | 2.10E+08 | 1.34E+08 | 1.56 | 0.64 | 9.32938E-07 |
| P49327 | Fatty acid synthase OS=Homo sapiens OX=9606 GN=FASN PE=1 SV=3 | 45 | 1668 | 80 | 6.59E+08 | 6.70E+08 | 6.05E+08 | 6.16E+08 | 6.02E+08 | 6.00E+08 | 5.70E+08 | 5.93E+08 | 4.93E+08 | 5.00E+08 | 5.19E+08 | 4.17E+08 | 6.25E+08 | 5.15E+08 | 1.21 | 0.28 | 0.00569581 |
| P35579 | Myosin-9 OS=Homo sapiens OX=9606 GN=MYH9 PE=1 SV=4 | 53 | 2773 | 89 | 8.51E+08 | 8.59E+08 | 8.02E+08 | 8.28E+08 | 7.72E+08 | 8.64E+08 | 9.69E+08 | 9.69E+08 | 8.65E+08 | 9.07E+08 | 9.15E+08 | 9.07E+08 | 8.29E+08 | 9.22E+08 | 0.90 | -0.15 | 0.001936962 |
| Q9Y490 | Talin-1 OS=Homo sapiens OX=9606 GN=TLN1 PE=1 SV=3 | 60 | 3010 | 94 | 1.31E+09 | 1.27E+09 | 1.14E+09 | 1.16E+09 | 1.17E+09 | 1.22E+09 | 1.49E+09 | 1.58E+09 | 1.38E+09 | 1.36E+09 | 1.37E+09 | 1.34E+09 | 1.21E+09 | 1.42E+09 | 0.85 | -0.23 | 0.001995176 |
| Q00610 | Clathrin heavy chain 1 OS=Homo sapiens OX=9606 GN=CLTC PE=1 SV=5 | 62 | 2680 | 96 | 1.08E+09 | 1.10E+09 | 1.07E+09 | 1.01E+09 | 1.05E+09 | 1.04E+09 | 7.58E+08 | 7.68E+08 | 7.06E+08 | 6.83E+08 | 7.89E+08 | 6.20E+08 | 1.06E+09 | 7.20E+08 | 1.47 | 0.56 | 3.84425E-06 |
| P02751 | Fibronectin OS=Homo sapiens OX=9606 GN=FN1 PE=1 SV=5 | 57 | 5321 | 97 | 2.38E+09 | 2.53E+09 | 2.46E+09 | 2.37E+09 | 2.31E+09 | 2.33E+09 | 3.82E+09 | 3.94E+09 | 3.35E+09 | 3.33E+09 | 3.23E+09 | 5.70E+09 | 2.40E+09 | 3.89E+09 | 0.62 | -0.70 | 0.010667989 |
| P21333 | Filamin-A OS=Homo sapiens OX=9606 GN=FLNA PE=1 SV=4 | 61 | 3031 | 105 | 1.45E+09 | 1.46E+09 | 1.39E+09 | 1.42E+09 | 1.34E+09 | 1.40E+09 | 1.06E+09 | 1.10E+09 | 9.80E+08 | 9.63E+08 | 8.95E+08 | 9.30E+08 | 1.41E+09 | 9.88E+08 | 1.43 | 0.51 | 3.16207E-06 |
| Q14204 | Cytoplasmic dynein 1 heavy chain 1 OS=Homo sapiens OX=9606 GN=DYNC1H1 PE=1 SV=5 | 35 | 1951 | 127 | 5.89E+08 | 5.63E+08 | 5.49E+08 | 5.70E+08 | 5.52E+08 | 5.19E+08 | 5.43E+08 | 4.82E+08 | 4.66E+08 | 4.44E+08 | 4.72E+08 | 3.93E+08 | 5.57E+08 | 4.67E+08 | 1.19 | 0.26 | 0.00454779 |
